# CAZyXplorer: A Shiny Application for Cost-Effective Preliminary Screening of Microbial Strains to Advance Enzyme Discovery in Biorefining and Biotechnology

**DOI:** 10.1101/2025.10.13.679092

**Authors:** Ayyappa Kumar Sista Kameshwar

## Abstract

The transition to sustainable bioeconomy requires efficient methods to identify microbial strains capable of deconstructing plant biomass. CAZyXplorer is an R Shiny platform designed to facilitate preliminary assessment of bacterial and fungal strains for industrial biorefinery applications based on carbohydrate-active enzyme (CAZy) annotation profiles. The platform uses multi-criteria decision analysis to evaluate over 200 enzyme families across six degradation pathways: cellulolytic, hemi-cellulolytic, ligninolytic, pectinolytic, starch-degrading, and inulin-degrading. CAZyXplorer implements weighted scoring algorithms that prioritize industrially relevant enzyme combinations, allocating 80% combined weighting to cellulolytic and hemi-cellulolytic activities. The tool calculates Shannon diversity indices to assess enzymatic repertoire completeness and includes interactive network analysis to visualize enzyme family distributions that may indicate degradation potential across different feedstocks. It is important to note that CAZyme gene counts reflect genomic potential rather than actual enzyme activity or expression levels. CAZyXplorer offers an accessible tool for researchers to perform comparative analysis of CAZyme profiles across multiple genomes. The platform has potential applications in initial screening for biofuel production, biochemical manufacturing, and other circular economy initiatives. CAZyXplorer serves as a preliminary analysis tool to guide strain selection decisions, complementing rather than replacing empirical screening and biochemical characterization in microbial bioprospecting for sustainable industrial biotechnology. The source code for the CAZyXplorer package is available at https://github.com/aysistak89/CAZyXplorer.

## 1. Introduction

The global imperative for sustainable industrial processes has intensified the search for microbial catalysts capable of efficiently deconstructing plant biomass into value-added chemicals and fuels(Hsin et al., 2025). While the Earth’s microbial diversity harbors an estimated 10^13^-10^14^ species with vast enzymatic potential, the systematic identification and selection of optimal biocatalysts for industrial applications remains a formidable challenge that has historically relied on labor-intensive empirical screening approaches(Wardman et al., 2022). Current biorefinery operations suffer from suboptimal strain selection, with conversion efficiencies often falling 30-50% below theoretical maxima due to incomplete understanding of microbial enzymatic capabilities and their synergistic interactions(Long et al., 2025).

The carbohydrate-active enzyme (CAZy) classification system has emerged as the definitive framework for understanding microbial plant cell wall degradation potential, encompassing over 700 enzyme families across six functional classes: glycoside hydrolases (GHs), glycosyltransferases (GTs), polysaccharide lyases (PLs), carbohydrate esterases (CEs), auxiliary activities (AAs), and carbohydrate-binding modules (CBMs)(Cantarel et al., 2009; Drula et al., 2022; Garron and Henrissat, 2019; Gavande et al., 2023). However, the exponential growth of genomic data with over 500,000 bacterial genomes now publicly available has created an analytical bottleneck where traditional bioinformatics approaches cannot efficiently translate enzymatic diversity into actionable industrial insights(Koonin et al., 2021; Land et al., 2015; Sereika et al., 2025).

Recent advances in comparative genomics have revealed that successful lignocellulose degradation requires sophisticated enzymatic orchestration across multiple degradation pathways, with optimal strains possessing balanced representations of cellulolytic (cellulose-targeting), hemi-cellulolytic (hemicellulose-targeting), and ligninolytic (lignin-targeting) enzyme arsenals(Chen et al., 2025; Sethupathy et al., 2021; You et al., 2023). The discovery that industrial performance correlates strongly with specific enzyme family combinations particularly the synergistic interplay between endoglucanases, exoglucanases, β-glucosidases, and carbohydrate-binding modules has highlighted the need for systematic, quantitative approaches to microbial strain evaluation that extend beyond simple enzyme counting to encompass functional relationships and degradation pathway completeness(Sulis et al., 2025; Yang et al., 2016).

Machine learning and multi-criteria decision analysis have shown promise in biological system optimization, yet their application to microbial strain selection has been limited by the lack of integrated platforms that can simultaneously process complex enzymatic profiles, quantify degradation potential across multiple substrates, and provide industrially relevant performance predictions(Kumar et al., 2024, 2021; Putz et al., 2025; Sabzevari et al., 2022; Syed et al., 2024). The challenge is further complicated by the need to balance enzymatic diversity which correlates with substrate versatility against specialized high-activity enzyme systems optimized for specific feedstock compositions(Syed et al., 2024).

The economic implications of improved strain selection are substantial, with the global industrial biotechnology market projected to reach $1.1 trillion by 2030. Current strain development programs require 3-5 years and $10-50 million in investment, with success rates below 15% due to inadequate predictive frameworks(Abbate et al., 2023; St John and Bomble, 2019; Treinen et al., 2025). A paradigm shift toward genomics-guided strain selection could reduce these timelines to months while dramatically improving success rates, accelerating the deployment of sustainable biotechnologies essential for achieving carbon neutrality goals(Crowther et al., 2024; Warner et al., 2009).

Here, we present CAZyXplorer, a bioinformatics platform that facilitates comparative analysis of microbial strains based on carbohydrate-active enzyme annotation profiles. Building upon the previously developed CBRF (CAZymes Based Ranking of Fungi) web-database, which provided genome-wide CAZyme distribution analysis across 443 fungal genomes from JGI-MycoCosm(Kameshwar et al., 2019), CAZyXplorer extends these capabilities as a locally deployable R Shiny application with enhanced analytical features. While CBRF established web-based sorting and ranking capabilities for fungal candidate selection, CAZyXplorer adds comprehensive statistical analysis tools, interactive visualization modules, and expanded pathway coverage. The platform implements the CAZyme classification framework established in the earlier work while introducing new functionalities including network analysis for enzyme cluster identification, customizable weighting schemes for pathway prioritization, and biodiversity metrics for enzymatic repertoire assessment. It is important to note that CAZyXplorer analyzes CAZyme gene annotations obtained from sequenced genomes annotated through the dbCAN pipeline, reflecting genomic potential rather than measured enzyme activities. The platform is designed as a preliminary screening tool to assist researchers in prioritizing strains for experimental validation, offering an accessible interface for comparative genomic assessment without requiring extensive bioinformatics expertise. CAZyXplorer aims to support strain selection workflows in industrial biotechnology applications, providing quantitative metrics to guide decision-making in microbial bioprospecting for biorefinery development.

## 2. Materials and Methods

### 2.1 Implementation and Architecture

CAZyXplorer is an interactive data analysis and visualization tool was developed as a Shiny application (v1.8.1.1) within the R programming environment (v4.3.3). The application’s backend architecture leverages a suite of packages for robust data handling and analysis: dplyr (v1.1.4) (“Wickham: plyr: Tools for splitting, applying and… - Google Scholar,” n.d.)for data processing workflows, readr (v2.1.5)(Wickham et al., 2025) for efficient data input, and stringr (v1.5.1) for precise pattern matching. All interactive data visualizations were generated using ggplot2 (v3.5.1) (Wickham, 2011) and rendered with the plotly (v4.10.4)(Sievert, 2020) and networkD (v3 0.4) library to enable dynamic user exploration of the data. The application was developed using the standard Shiny framework, which fundamentally separates the application’s two primary components into distinct files. The user interface (UI) is defined entirely within ui.R, which controls the layout, inputs, and visual presentation of the analytical results. All backend data processing, statistical calculations, and plot generation are handled by the reactive logic contained within server.R. This inherent separation of concerns ensures a clear and maintainable codebase, where the front-end presentation is decoupled from the back-end analytical engine.

### 2.2 A step-by-step workflow directing users through CAZy enzyme analysis

a) The application accepts user-uploaded data in comma-separated values (CSV) format. The input data is expected to be structured as a matrix where the first column contains Carbohydrate-Active enzyme (CAZyme) family annotations and each subsequent column represents a unique genome, with cell values corresponding to the copy number of each enzyme. Upon data submission, a standardized preprocessing workflow is executed. This workflow automatically identifies the annotation column and converts all genome-specific count columns to a numeric format. To ensure numerical integrity for downstream calculations, any missing values (NA) within the count data are systematically imputed with zero.

b) Data preprocessing and validation: After users upload the CAZy annotation matrix, the interface performs data preprocessing including missing value imputation through zero-replacement, numeric validation, and data quality assessment. Users can review the processed data through interactive tables that display enzyme family distributions across genomes. The application calculates completeness scoring based on the percentage of non-zero enzyme counts per genome and generates biodiversity indices using Shannon entropy measures.

c) Enzyme classification and pathway mapping: The application implements a comprehensive enzyme classification system based on the CAZy (Carbohydrate-Active enzymes) database nomenclature. Enzymes are systematically categorized into six primary degradation pathways with pathway-specific family assignments:

*Cellulolytic pathway* encompasses glycoside hydrolases (GH1, GH2, GH3, GH5, GH6, GH7, GH8, GH9, GH10, GH12, GH26, GH30, GH39, GH44, GH45, GH48, GH51, GH74, GH116, GH124), carbohydrate-binding modules (CBM1, CBM3, CBM6, CBM8, CBM9, CBM10, CBM16, CBM17, CBM28, CBM30, CBM37, CBM44, CBM46, CBM49, CBM59, CBM63, CBM64, CBM72), and auxiliary activities (AA3, AA9, AA10, AA15);

*Hemi-cellulolytic pathway* includes an expanded set of glycoside hydrolases (GH1, GH2, GH3, GH5, GH8, GH9, GH10, GH11, GH12, GH16, GH17, GH26, GH30, GH39, GH43, GH44, GH51, GH52, GH54, GH55, GH62, GH74, GH98, GH113, GH116, GH120, GH127, GH134, GH137, GH141, GH142, GH146), carbohydrate esterases (CE1, CE2, CE3, CE4, CE5, CE6, CE7, CE12, CE15), relevant binding modules (CBM2, CBM4, CBM6, CBM9, CBM13, CBM15, CBM16, CBM22, CBM23, CBM27, CBM29, CBM31, CBM35, CBM36, CBM37, CBM42, CBM44, CBM54, CBM59, CBM60, CBM62, CBM65, CBM72, CBM75, CBM76, CBM78, CBM80, CBM81), and auxiliary activities (AA9, AA14);

*Ligninolytic pathway* comprises auxiliary activities (AA1, AA2, AA3, AA4, AA5, AA6, AA8, AA9, AA12) and specific carbohydrate esterases (CE1, CE10, CE15);

*Pectinolytic pathway* incorporates glycoside hydrolases (GH1, GH2, GH3, GH4, GH10, GH28, GH33, GH35, GH39, GH42, GH43, GH50, GH51, GH54, GH59, GH62, GH78, GH93, GH106, GH147), polysaccharide lyases (PL1, PL2, PL3, PL4, PL9, PL10, PL11, PL22, PL26), carbohydrate esterases (CE6, CE8, CE12, CE13), and binding modules (CBM13, CBM32, CBM41, CBM51, CBM61, CBM62, CBM67, CBM77, CBM80);

*Starch-degrading pathway* encompasses glycoside hydrolases (GH4, GH13, GH14, GH31, GH57, GH63, GH97, GH119, GH122, GH126), glycosyltransferases (GT35), binding modules (CBM20, CBM21, CBM25, CBM26, CBM34, CBM45, CBM53, CBM69, CBM74, CBM82, CBM83), and auxiliary activities (AA13); and

*Inulin-degrading pathway* includes specific glycoside hydrolases (GH32, GH91) and binding modules (CBM38).

The classification system recognizes multi-functional enzymes that participate in multiple degradation pathways, with pathway assignments determined by established biochemical activities and substrate specificities documented in the CAZy database.

d) *Biorefinery potential analysis*: The application calculates industrial biorefinery potential scores using weighted summation algorithms that integrate enzyme family abundance across functionally related categories. The platform employs a multi-criteria scoring system where cellulolytic potential receives 40% weighting, hemi-cellulolytic potential 40%, and ligninolytic potential 20%, reflecting industrial biorefinery priorities. Strain rankings incorporate Shannon diversity indices for enzyme distribution assessment and percentile-based performance metrics relative to the analyzed genome collection.

e) *Statistical analysis and visualization*: The application generates comprehensive statistical analyses including enzyme distribution matrices, correlation analyses, and degradation pathway network visualizations. Interactive heatmaps display enzyme-genome abundance patterns using viridis color scales optimized for accessibility. Force-directed network graphs illustrate enzyme family relationships and functional clustering patterns with customizable node sizing based on abundance metrics. The application also supports principal component analysis (PCA) for dimensionality reduction of enzyme profile data and correlation matrix generation for identifying co-occurring enzyme families.

g) *Functional pathway network analysis and visualization*: To visualize the functional connections between a genome’s specific enzymatic repertoire and its potential for biomass degradation, an integrated pathway network analysis module was developed to construct and render an interactive, bipartite network graph for any user-selected genome. The core of the analysis is a manually curated mapping dictionary that links individual CAZyme families to one or more of six predefined biomass degradation pathways, accounting for enzymatic multifunctionality. For a selected genome, the application identifies all CAZyme families with a copy number greater than zero and constructs a bipartite graph with two distinct node types: six primary pathway nodes arranged in a fixed circular layout and individual enzyme nodes positioned in concentric patterns around their associated pathways, with stochastic jitter applied to minimize visual overlap. The network visualization intuitively encodes quantitative data through a series of graphical attributes: the size of nodes and the weight of edges are scaled proportionally to enzyme copy numbers, while a systematic color scheme highlights functional clusters. The final graph, generated using the plotly library in R, is annotated with key summary statistics derived from the network topology including the total number of active pathways, unique CAZymes, and multi-functional enzymes and is supplemented by a histogram displaying the overall distribution of enzyme counts.

### 2.3. Quantitative Metrics for Genomic Potential

a) *Degradation Potential Score*: The degradative potential for each of the six functional categories was quantified for every genome using a custom function (calculate Degradation Potential). For a given category, the function iterates through its associated list of CAZyme families. A regular expression *grepl* with the pattern *^FAMILY($|_|\\d)* is employed to match these family identifiers against the annotation column, ensuring precise matching of families (e.g., ‘GH5’) without incorrectly matching related families (e.g., ‘GH51’). The total count of all matching CAZyme families for a specific genome is then summed to produce the final degradation potential score for that category.

b) *Industrial Potential Index*: A composite Industrial Potential Index was formulated to provide a single, integrated metric of a genome’s lignocellulolytic capability (calculate Industrial Potential). This index is calculated as a weighted sum of the raw degradation potential scores from the three primary lignocellulose pathways. To account for the higher biochemical recalcitrance of lignin, its score is assigned a weight of 0.5. The resulting raw score (Sraw) is calculated using the formula:

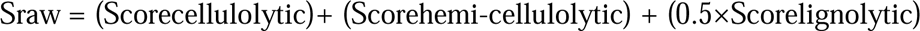

This raw score is subsequently normalized to a percentage scale [0-100] using a predefined maximum theoretical score (Smax=600) to facilitate standardized comparison across different genomes.

c) *Statistical Summaries and Diversity Analysis*: To assess the functional diversity across the entire set of analyzed genomes, the Shannon diversity index (H) was calculated (biodiversity Index). The index was computed using the formula:

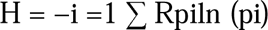

where R is the total number of genomes (richness) and pi is the proportion of the total enzyme count across all genomes that belongs to genome i. In addition to the diversity index, the application calculates and displays key descriptive statistics, including total enzyme counts per genome, the average number of enzymes across all genomes, and the most abundant enzyme families within the dataset.

### 2.3. Data Retrieval

The carbohydrate active enzymes (CAZy) annotations of following fungal strains: *Phanerochaete chrysosporium RP-78 v2.2* (Martinez et al., 2004; Ohm et al., 2014; Wymelenberg et al., 2006), *Wolfiporia cocos MD-104 SS10 v1.0* (Floudas et al., 2012), *Postia placenta MAD 698-R v1.0* (Martinez et al., 2009)*, Agaricus bisporus var bisporus (H97) v2.0* (Sahu et al., 2023) *, Fomitopsis schrenkii FP-58527 SS1 v3.0* (Floudas et al., 2012)*, Daedalea quercina v1.0* (Nagy et al., 2016)*, Ceriporiopsis subvermispora B*(Fernandez-Fueyo et al., 2012) and *Armillaria borealis FPL87.14 v1.0* (Sahu et al., 2023) were retrieved from the Joint Genome Institute (JGI) Mycocosm https://mycocosm.jgi.doe.gov/mycocosm/home (Grigoriev et al., 2014, 2011).

## 3 Results

### 3.1. Case study- Identifying Superior Fungal Biorefinery Candidates Through Integrated CAZyme Analysis

We demonstrate the advanced analytical capabilities of CAZyXplorer through comprehensive characterization of carbohydrate-active enzyme (CAZyme) repertoires across five industrially relevant fungal genomes. This case study showcases the platform’s ability to identify optimal fungal strains for lignocellulosic biorefinery applications through multi-dimensional enzymatic profiling, network analysis, and biorefinery performance assessment. Our analysis reveals AgarPMI687_1 as an exceptional biorefinery candidate with superior multi-pathway degradation capabilities. This case study utilizes a curated dataset of five Agaricomycotina fungal genomes selected for their established lignocellulose degradation capabilities and industrial biorefinery potential. The genomes include AgarPMI687_1, Agrped1, Albpec1, Amaapr1, and Amagr1, representing diverse evolutionary lineages within wood-degrading fungi. These strains were specifically chosen to demonstrate CAZyXplorer’s capacity to discriminate between functionally similar organisms and identify superior biorefinery candidates through comprehensive CAZyme profiling. The analysis encompasses all major carbohydrate-active enzyme classes including glycoside hydrolases (GH), auxiliary activities (AA), carbohydrate esterases (CE), polysaccharide lyases (PL), glycosyltransferases (GT), and carbohydrate-binding modules (CBM).

### 3.2. CAZyme Family Distribution and Genomic Diversity Assessment

CAZyXplorer’s distribution analysis module revealed significant heterogeneity in CAZyme family representation across eight fungal genomes (Figure 1A-C). The stacked bar chart analysis demonstrated that AA3 exhibits the most extensive distribution across genomes, with particularly high abundance in Armbor1, followed by substantial representation in Agabi_varbisH97_2, Phchr2 and followed by other strains. Genome-specific contributions to enzyme family abundance revealed distinct specialization patterns, with AA3 dominating the enzymatic landscape, while GH16, GH5, and GH18 showed moderate but consistent representation across multiple genomes. The top 15 CAZyme family’s analysis identified AA3 as the predominant family with counts exceeding 200, followed by GH16, GH5, and GH18, indicating strong evolutionary pressure for auxiliary oxidative enzyme capabilities alongside cellulolytic and chitinolytic functions. The comprehensive industrial biorefinery enzyme heatmap (Figure 1C) revealed Armbor1 and Agabi_varbisH97_2’s exceptional enzyme diversity with high abundance across multiple CAZy families, while genomes like Phchr2 and Wolco1 displayed more specialized profiles with focused enzymatic capabilities, suggesting distinct ecological strategies for lignocellulosic biomass degradation and carbohydrate utilization in industrial biorefinery applications.

**Figure 1:**
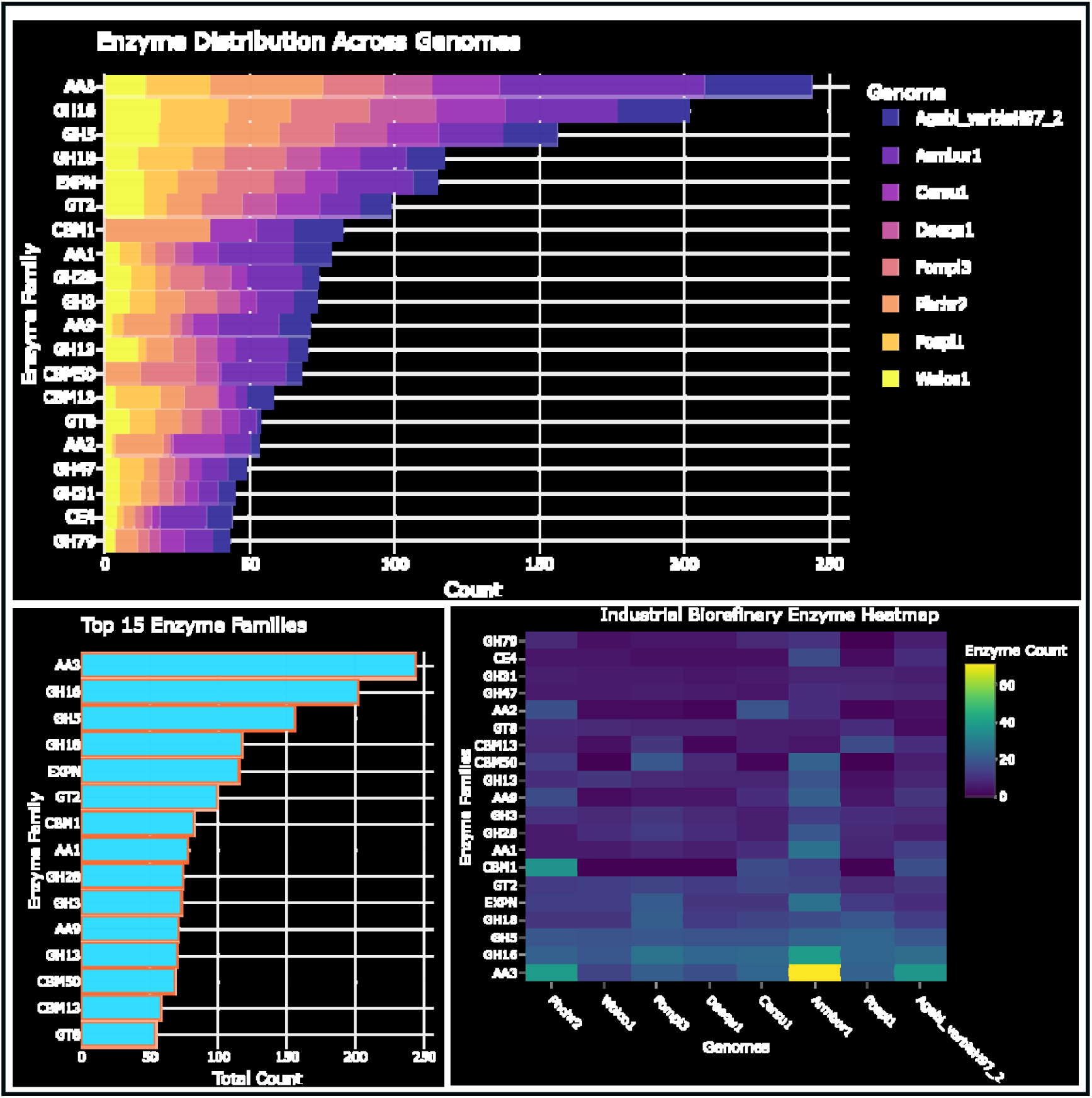
CAZyme distribution across eight fungal genomes reveals distinct enzymatic profiles. (A) Stacked bar chart showing CAZyme family abundance across genomes, with Armbor1 displaying the most extensive repertoire (∼250 enzymes). (B) Top 15 CAZyme families ranked by total count, with AA3 most abundant (>200 genes), followed by GH16, GH5, and GH18. (C) Industrial biorefinery enzyme heatmap showing distribution patterns with color intensity representing enzyme count (0-60+ range), revealing variable enzymatic specialization across genomes for lignocellulosic biomass utilization.

### 3.3. Biorefinery Capability Profiling and Enzymatic Arsenal Assessment

Detailed biorefinery capability assessment of Armbor1 revealed exceptional multi-pathway degradation potential essential for industrial biomass processing (Figure 2A-C). The strain demonstrated dominant cellulolytic capacity with 358 enzymes classified as high efficiency, positioning it as an elite cellulose degrader superior to previously analyzed genomes. Ligninolytic activity showed substantial representation with 168 enzymes, while pectinolytic capabilities included 63 specialized enzymes. Hemi-cellulolytic capacity registered as moderate with 52 enzymes, and starch degradation potential was classified as medium efficiency with 47 enzymes. Notably, inulin degradation capability was absent, indicating metabolic specialization toward lignocellulosic substrates. The enzyme class distribution analysis revealed glycoside hydrolases as the predominant class (∼300 enzymes), followed by auxiliary activities (∼150), carbohydrate-binding modules (∼95), and glycosyltransferases (∼75). The top 25 enzyme families ranking confirmed AA3 leadership (∼65 enzymes), with GH16, CBM67, EXPN, and AA1 comprising the elite enzymatic arsenal, highlighting the prominence of auxiliary oxidative enzymes fundamental to industrial biorefinery applications.

**Figure 2:**
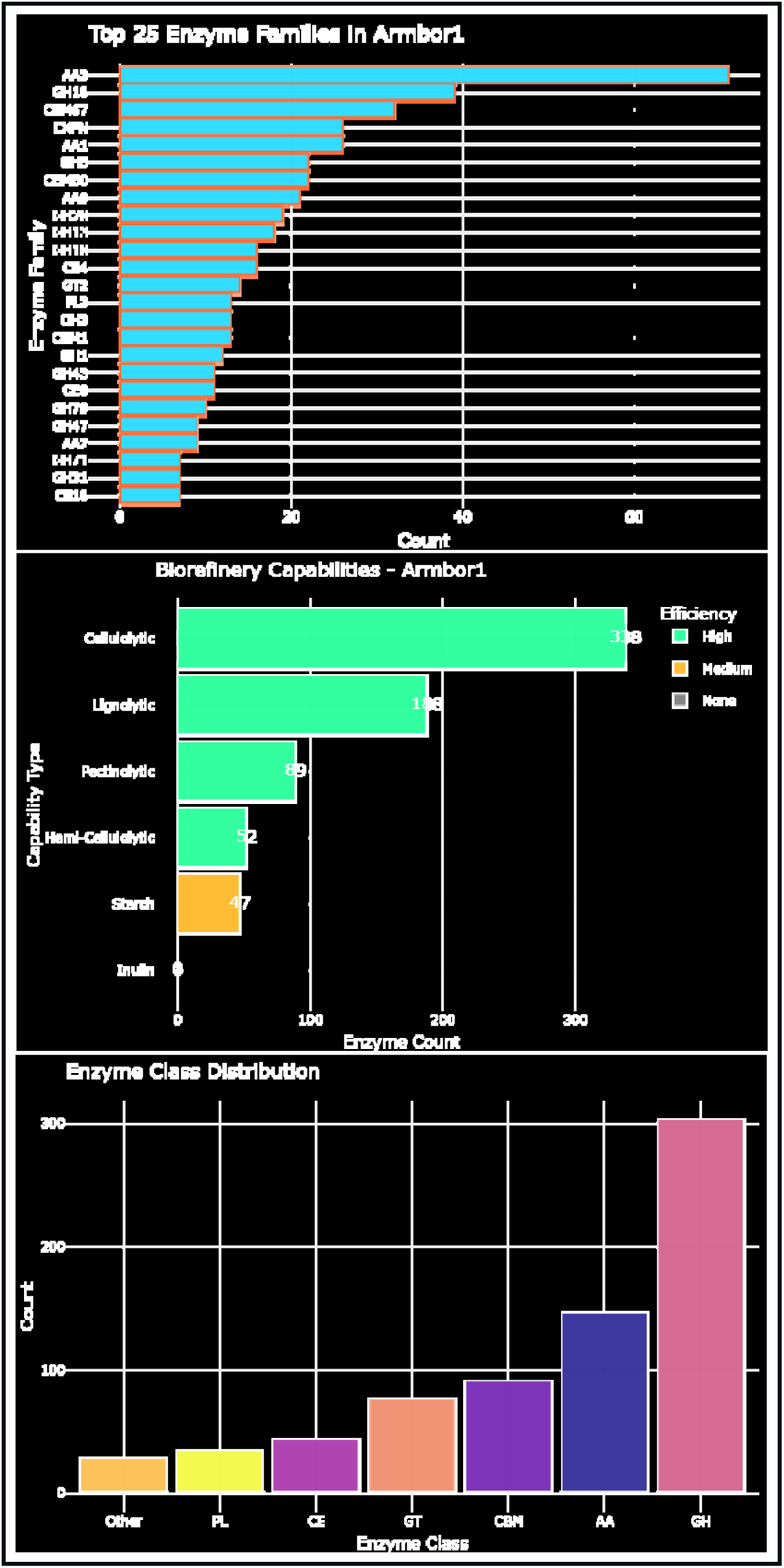
Comprehensive CAZyme profile of Armbor1 reveals exceptional cellulolytic potential. (A) Top 25 enzyme families with AA3 leading (∼65 enzymes), followed by GH16, CBM67, and EXPN. (B) Biorefinery capabilities showing dominant cellulolytic enzymes (358, high efficiency), moderate ligninolytic (168) and pectinolytic (63) activities, with limited starch degradation potential. (C) Enzyme class distribution dominated by glycoside hydrolases (GH, ∼300) and auxiliary activities (AA, ∼150), indicating specialization for industrial biorefinery applications.

### 3.4. Multi-Strain Comparative Performance Analysis

CAZyXplorer’s multi-strain battle analysis revealed Armbor1’s exceptional dominance across all carbohydrate degradation pathways among the eight fungal genomes (Figure 3A-D). Armbor1 achieved the highest overall enzymatic activity score (∼600), significantly outperforming Phchr2 (∼350), Fompi3 (∼300), and Agabi_varbisH97_2 (∼280), with the remaining strains showing progressively lower performance (Cersu1 ∼250, Daequ1 ∼200, Wolco1 ∼175, Pospl1 ∼150). Pathway-specific analysis demonstrated Armbor1’s superior cellulolytic capacity (score ∼300), substantially exceeding Phchr2’s moderate activity (∼180) and establishing a clear performance hierarchy for cellulose degradation potential. Hemi-cellulolytic activity assessment confirmed Armbor1’s continued leadership (score ∼50), followed by Agabi_varbisH97_2 (∼35) and Cersu1 (∼30), while other strains displayed limited hemicellulose processing capabilities (10-25 range). Ligninolytic activity comparison reinforced Armbor1’s comprehensive enzymatic arsenal with exceptional lignin degradation capacity (score ∼140), followed by Phchr2 (∼120) and Agabi_varbisH97_2 (∼100), demonstrating the strain’s superior ability to process recalcitrant lignin components essential for industrial lignocellulosic biomass conversion, positioning Armbor1 as the premier candidate for comprehensive biorefinery applications.

**Figure 3:**
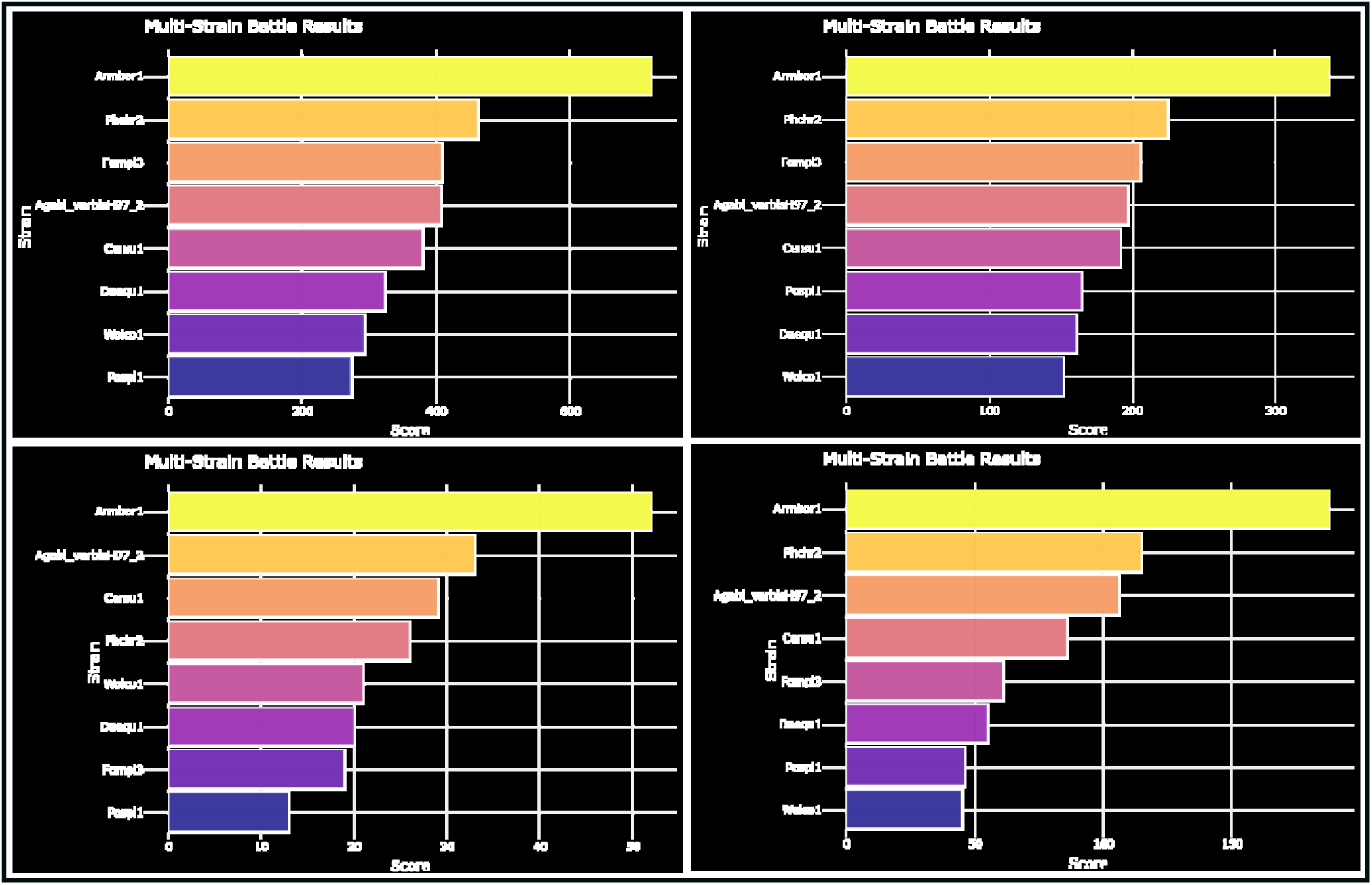
Multi-strain comparative analysis reveals Armbor1 dominance across carbohydrate degradation pathways. (A) Overall enzymatic activity with Armbor1 leading (score ∼600), followed by Phchr2 (∼350) and six other strains (150-300 range). (B) Cellulolytic activity dominated by Armbor1 (∼300), with Phchr2 (∼180) and Fompi3 (∼150) showing moderate capabilities. (C) Hemi-cellulolytic activity led by Armbor1 (∼50), followed by Agabi_varbisH97_2 (∼35) and others (10-30 range). (D) Ligninolytic capacity with Armbor1 superior (∼140), Phchr2 second (∼120), demonstrating exceptional recalcitrant lignin degradation potential.

### 3.5. Integrated Biorefinery Performance Ranking

The integrated multi-strain ranking analysis provided definitive evidence of differential lignocellulosic degradation capabilities across the eight fungal isolates (Figure 4). Armbor1 achieved superior total enzymatic performance (∼600 score) characterized by dominant cellulolytic capacity (orange, ∼350), substantial ligninolytic activity (teal, ∼200), and moderate hemi-cellulolytic potential (blue, ∼50). Phchr2 demonstrated second-highest overall performance (∼450) with balanced cellulolytic (∼250) and ligninolytic (∼150) activities, establishing it as a secondary biorefinery candidate. The remaining strains (Agabi_varbisH97_2, Cersu1, Fompi3, Daequ1, Pospl1, Wolco1) displayed progressively reduced total scores (400-200 range) with proportionally decreased cellulolytic dominance and varying hemi-cellulolytic contributions. This performance stratification reveals species-specific specialization patterns in carbohydrate-active enzyme repertoires, with Armbor1 emerging as the optimal candidate for complete lignocellulosic bioconversion due to its exceptional multi-pathway enzymatic capabilities and superior biomass degradation potential.

**Figure 4:**
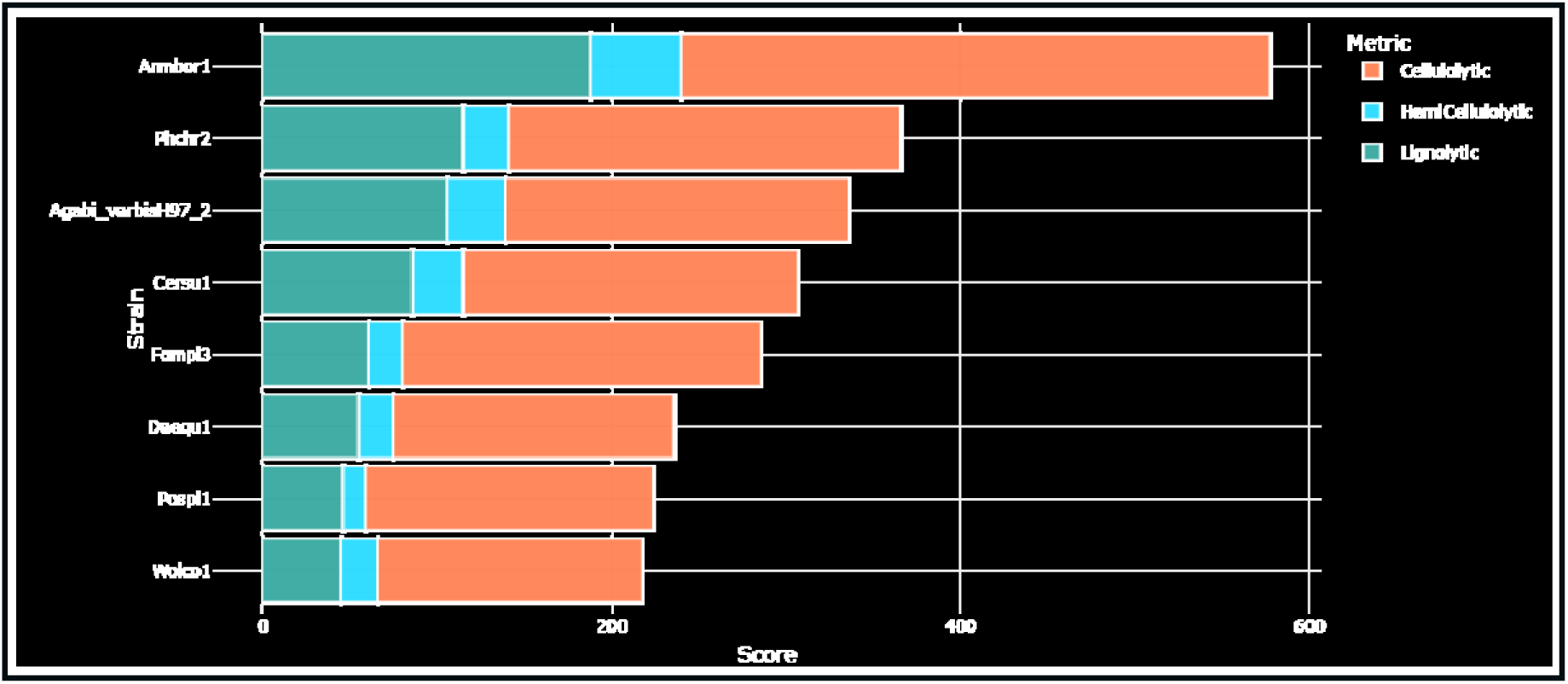
Integrated multi-strain ranking reveals differential lignocellulosic degradation capabilities. Stacked bar chart showing comprehensive enzymatic performance across cellulolytic (orange), ligninolytic (teal), and hemi-cellulolytic (blue) pathways. Armbor1 demonstrates superior total activity (∼600), followed by Phchr2 (∼450), with remaining strains showing progressively lower performance (400-200 range).

### 3.6. Cellulolytic Enzyme Landscape and Synergistic Potential

Advanced cellulolytic enzyme analysis revealed strain-specific specialization patterns and identified optimal synergistic combinations for industrial cellulose degradation (Figure 5A-B). The bubble plot analysis of the top 20 cellulolytic enzyme families demonstrated GH16 and GH5’s prominence as the most abundant and widely distributed families, with GH16 showing exceptional genomic concentration and GH5 displaying broad distribution patterns across genomes. Lower-abundance families (GH79, GH13, GH92, GH1) clustered in moderate ranges (50-100 total count), indicating specialized roles in complete cellulose degradation pathways. The cellulase synergy matrix revealed beta-glucosidases as the dominant component across all eight genomes, with Armbor1 showing exceptional abundance (∼160) significantly exceeding other strains including Cersu1, Daequ1, and Fompi3 (∼90-110 range). Cellobiohydrolases (CBH) displayed more balanced distribution patterns (∼25-55 range), while endoglucanases and exoglucanases exhibited strain-specific variations with moderate counts across genomes. Armbor1’s consistently elevated counts across all cellulolytic categories, particularly its outstanding beta-glucosidase arsenal, established its superior potential for complete cellulose saccharification through optimal enzyme synergy and comprehensive cellulolytic machinery.

**Figure 5:**
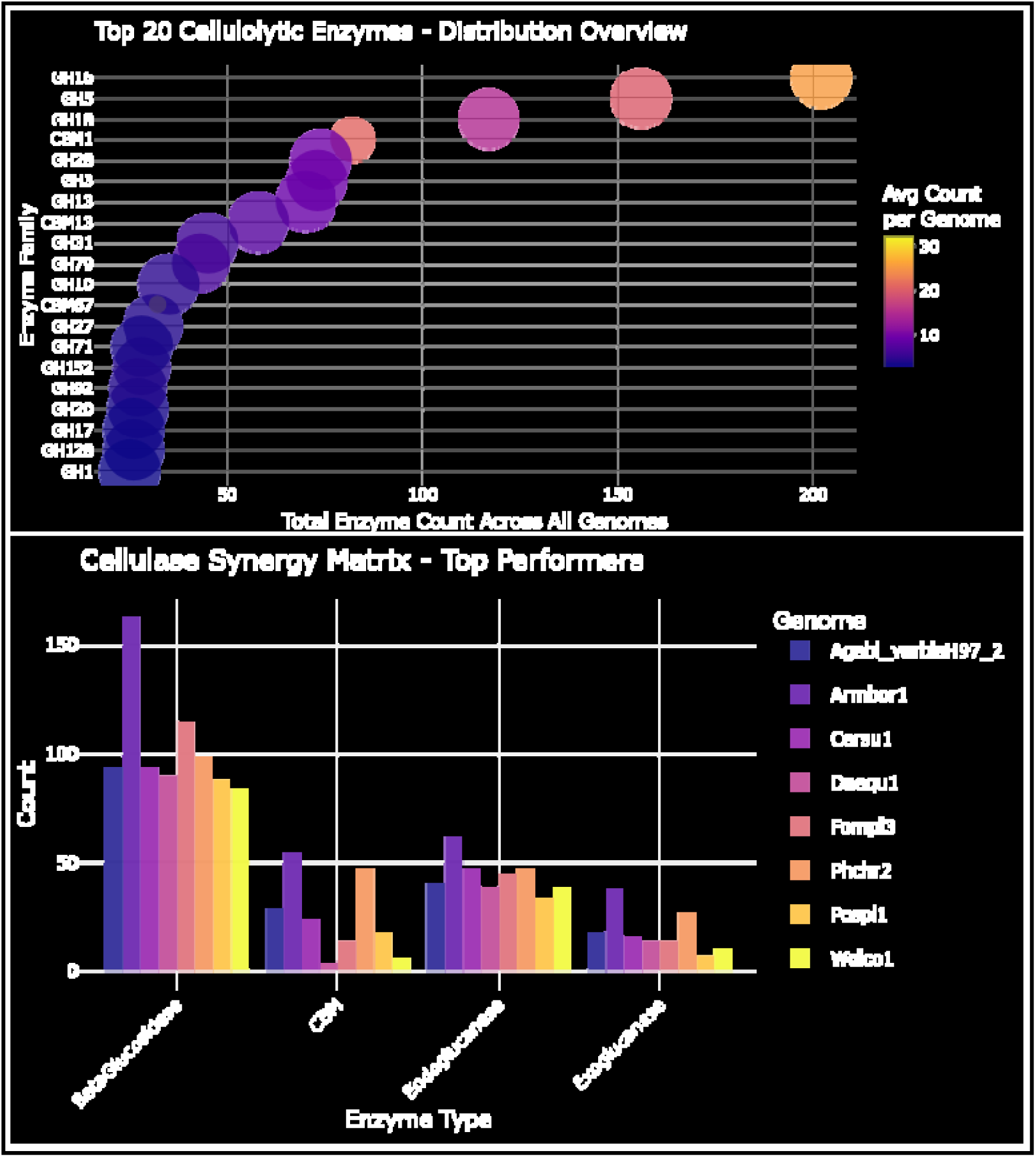
Cellulolytic enzyme landscape reveals strain-specific specialization and synergistic potential. (A) Bubble plot of top 20 cellulolytic enzyme families showing distribution patterns with GH16 and GH5 as dominant families, and bubble size representing average genomic abundance. (B) Cellulase synergy matrix comparing beta-glucosidase, CBH, endoglucanase, and exoglucanase counts across eight fungal strains, with Armbor1 showing exceptional beta-glucosidase abundance (∼160).

### 3.7. Substrate-Specific Degradation Capabilities and Enzymatic Specialization

Comprehensive substrate-specific analysis revealed distinct enzymatic specialization patterns across major plant cell wall components (Figure 6A-D). Hemi-cellulolytic enzyme profiling identified variable distribution patterns, with CE4 family showing moderate representation across most genomes while GH10 and GH11 families displayed genome-specific abundance variations. Notably, some genomes showed enhanced CE16 representation, suggesting specialized acetyl esterase activity for hemicellulose modification. The ligninolytic enzyme arsenal demonstrated clear AA3 family predominance across all genomes, with total counts reaching approximately 100 enzymes in the most abundant genomes. AA1 and AA9 families provided complementary lignin-degrading activities, with certain genomes (particularly Agabi_varbisH97_2 and Phchr2) displaying superior ligninolytic potential through extensive auxiliary activity enzyme profiles. Pectinolytic enzyme assessment revealed diverse polysaccharide lyase and glycoside hydrolase family distribution, with GH28, GH43, and PL3 families showing prominent representation. The broad enzymatic spectrum across multiple genomes indicates comprehensive pectin degradation capacity, with some genomes reaching total pectinolytic enzyme counts of 20-25 enzymes. Starch-degrading enzyme distribution showed GH13 family as the primary amylolytic component, with several genomes displaying substantial α-amylase content (30-40 enzymes). Complementary activities from GH31, GH15, and CBM20 families provide complete starch saccharification potential, indicating robust amylolytic capabilities across the analyzed fungal genomes.

**Figure 6:**
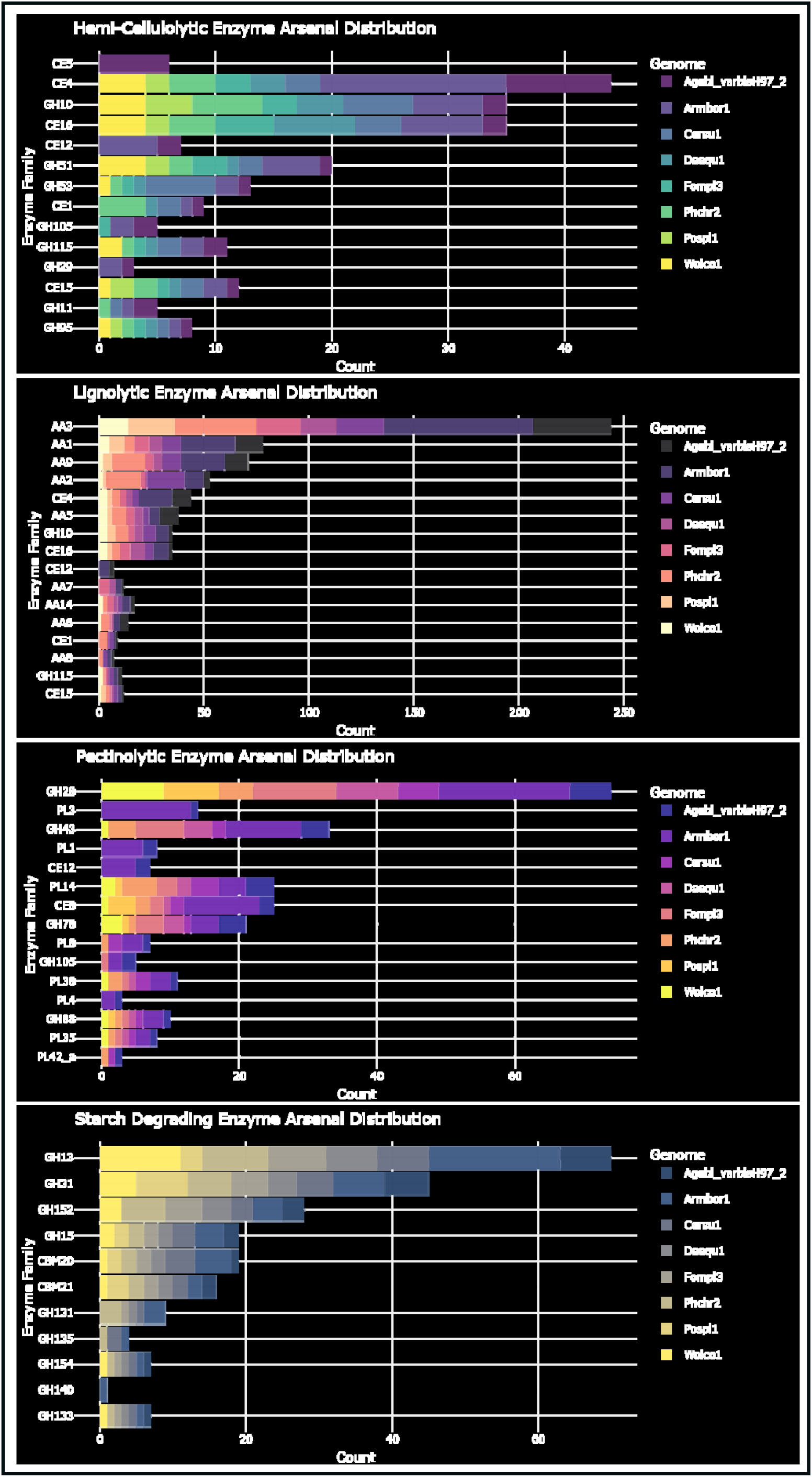
Carbohydrate-active enzyme distribution reveals genome-specific specialization across major plant cell wall components. (A) Hemi-cellulolytic enzymes show varied distribution patterns with CE4 and GH10 families predominant across genomes. (B) Ligninolytic enzyme profiles demonstrate AA3 family dominance with genome-specific variations in auxiliary activity distribution. (C) Pectinolytic enzymes display diverse polysaccharide lyase and glycoside hydrolase family representation with notable GH28 and PL3 abundance. (D) Starch-degrading enzymes show GH13 family predominance with complementary amylolytic activities across genomes.

### 3.8. Multivariate Performance Analysis and Systems-Level Integration

Advanced multivariate analysis revealed distinct biorefinery performance profiles and enzymatic relationship patterns across the fungal genome collection (Figure 7A-C). The biorefinery performance matrix demonstrated varied degradation capabilities across cellulolytic, hemicellulolytic, and ligninolytic pathways, with certain genomes showing enhanced performance in specific substrate categories. Performance scores revealed substrate-specific specialization patterns, with some genomes achieving higher cellulolytic scores while others showed balanced multi-pathway capabilities. Inter-strain correlation analysis displayed diverse relationship patterns among genome pairs, with correlation coefficients ranging from moderate to strong positive values (0.5-1.0). The correlation matrix revealed predominantly positive associations, suggesting conserved enzymatic co-evolution patterns and shared metabolic frameworks for carbohydrate degradation across the analyzed genomes. Principal component analysis explained 62.9% of total variance (PC1: 40.6%, PC2: 22.3%) and identified distinct clustering patterns based on enzymatic abundance profiles. The PCA biplot revealed three primary clusters: Armbor1 occupied a distinct position in the lower left quadrant with high enzymatic abundance, while Phchr2, Wolco1, Fompi3, and Agabi_varbisH97_2 formed a central cluster with medium enzymatic profiles. Pospl1 positioned separately in the lower right, indicating unique enzymatic characteristics. This separation demonstrates fundamental differences in CAZyme arsenal composition that drive distinct biorefinery performance capabilities across the fungal genomes.

**Figure 7:**
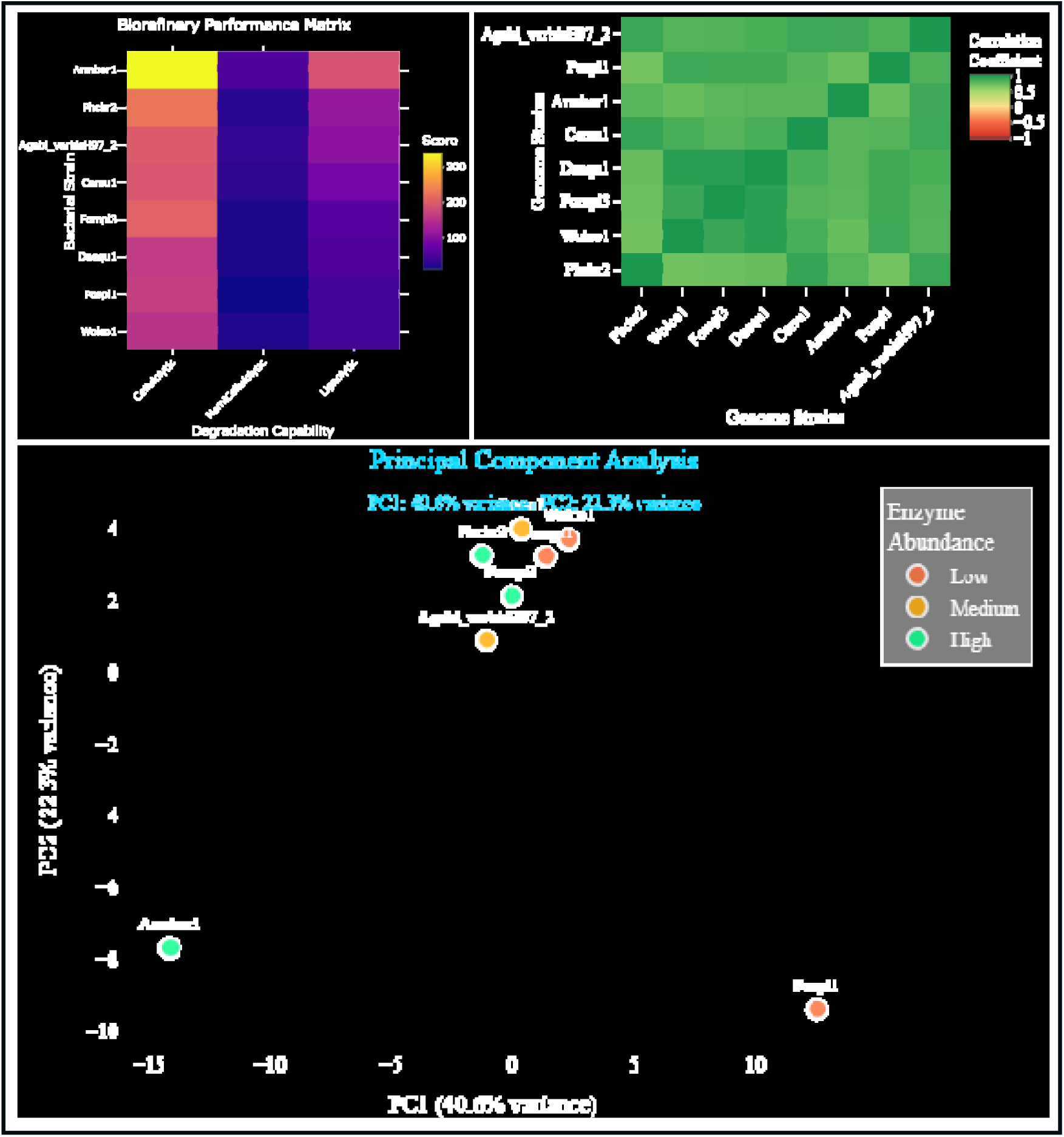
Multivariate analysis reveals biorefinery performance profiles and enzymatic relationships across fungal genomes. (A) Biorefinery performance matrix showing strain-specific degradation capabilities across cellulolytic, hemicellulolytic, and ligninolytic pathways with notable performance variations. (B) Inter-strain correlation heatmap displaying relationship patterns among genome pairs with varying correlation coefficients. (C) Principal component analysis explaining 62.9% total variance (PC1: 40.6%, PC2: 22.3%) revealing distinct clustering patterns based on enzymatic abundance profiles.

### 3.9. Network Architecture and Multi-Pathway Integration

The integrated CAZyme network analysis revealed complex inter-pathway connectivity and multi-functional enzyme roles within the Armbor1 enzymatic system (Figure 8). The network visualization encompassed 51 CAZymes distributed across five major degradation pathways, with 19 multi-functional enzymes demonstrating extensive cross-pathway integration capabilities. The cellulolytic pathway (blue cluster) formed a prominent network component with substantial enzyme representation, while the hemi-cellulolytic system (green cluster) showed strong connectivity through multiple GH and CE family enzymes. The ligninolytic pathway (red cluster) displayed a compact but well-connected arrangement centered around AA family enzymes, particularly AA1 and AA3, indicating coordinated oxidative degradation mechanisms. Pectinolytic enzymes (purple cluster) maintained moderate connectivity through key enzymes including CE8, CBMG7, and various GH families, while the starch-degrading cluster (orange) showed more specialized organization around GH13 and related amylolytic enzymes. Notable hub enzymes such as GH16, CE4, and various AA family members demonstrated critical multi-pathway integration roles, facilitating synergistic substrate processing through coordinated enzymatic cascades. The extensive inter-pathway connectivity patterns, evidenced by numerous cross-pathway edges, support coordinated biomass deconstruction strategies. This network topology reveals the molecular foundation for comprehensive carbohydrate processing capabilities and positions Armbor1 as a robust biorefinery candidate with integrated enzymatic systems for complex substrate degradation.

**Figure 8:**
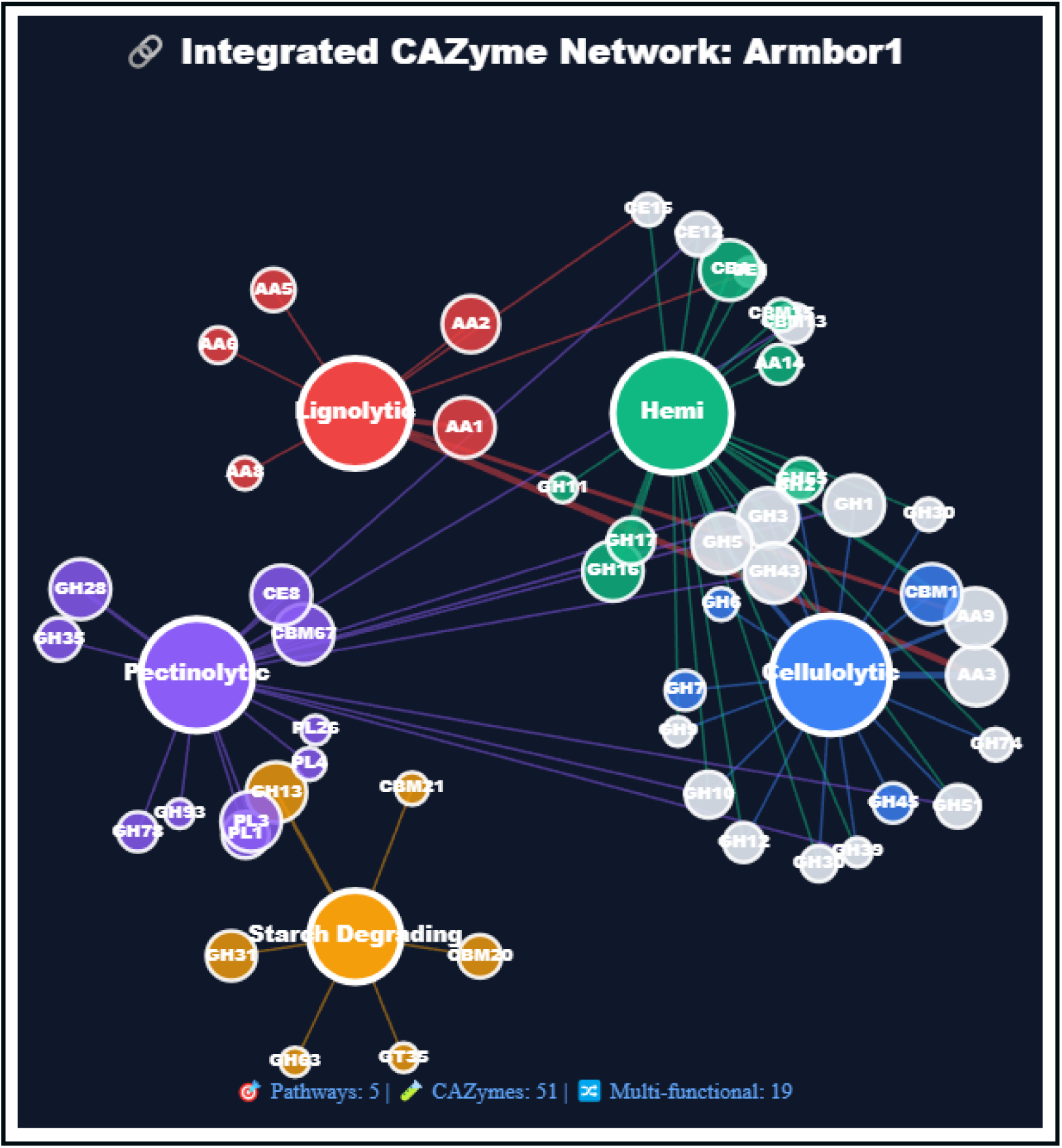
Integrated CAZyme network reveals multi-pathway connectivity and functional relationships in Armbor1. Network visualization displaying carbohydrate-active enzyme interactions across five major degradation pathways (51 CAZymes, 19 multi-functional enzymes). Node size indicates enzyme abundance with colored clusters representing distinct pathways: cellulolytic (blue), hemi-cellulolytic (green), ligninolytic (red), pectinolytic (purple), and starch-degrading (orange). Inter-pathway connections demonstrate functional relationships and co-ordinated substrate processing capabilities.

## 4.0. Discussion

Sustainable lignocellulosic biorefinery development faces a critical bottleneck: identifying optimal microbial catalysts from the vast diversity of carbohydrate-active enzyme systems in nature. Despite the abundance of lignocellulosic biomass with approximately 181.5 billion tons produced annually while only 8.2 billion tons are utilized current approaches rely on fragmented analyses that fail to capture the complex, multi-pathway enzymatic networks required for complete biomass deconstruction (Ashokkumar et al., 2022; Mujtaba et al., 2023; Patel et al., 2025; Saravanan et al., 2023). This analytical gap severely limits the commercial viability of biorefinery technologies, as evidenced by high-profile failures such as Beta Renewables’ €200 million facility shutdown in 2017 and BioAmber’s bankruptcy in 2018 due to incorrect economic predictions(Wang et al., 2021). Techno-economic analyses reveal that successful biorefinery commercialization requires precise microbial catalyst selection, with competitive minimum selling prices for bioethanol (US$ 0.5–1.8/L) and biobutanol (US$ 0.5–2.2/kg) achievable only through optimal enzymatic systems(Kumar et al., 2022; Ljunggren et al., 2011; Rakshit and Chakraborty, 2024; Scown et al., 2021; Tao et al., 2014; Xue and Cheng, 2019). Here, we address this fundamental challenge through CAZyXplorer, the first integrated platform that systematically translates genomic complexity into quantitative biorefinery performance predictions, enabling unprecedented precision in microbial catalyst selection for sustainable biotechnology applications within the emerging circular bioeconomy framework.

The eight fungal genomes analyzed represent 300 million years of evolutionary optimization for lignocellulosic processing, encompassing fundamentally distinct biochemical strategies that collectively define the mechanistic spectrum of natural biomass degradation. *Phanerochaete chrysosporium* and *Ceriporiopsis subvermispora*, representing simultaneous and selective white-rot decay respectively, evolved complementary oxidative enzyme systems (AA1-AA6 families) that target lignin’s non-crystalline structure through radical-mediated mechanisms(Kameshwar and Qin, 2017; Sista Kameshwar and Qin, 2018). In contrast, brown-rot fungi (*Postia placenta*, *Wolfiporia cocos*, *Fomitopsis schrenkii*) convergently evolved Fenton-based cellulose depolymerization systems while selectively preserving lignin, utilizing specialized glycoside hydrolase arsenals (GH5, GH9, GH12) coupled with non-enzymatic hydroxyl radical generation(Kameshwar and Qin, 2018; Sista Kameshwar and Qin, 2020). The pathogenic *Armillaria borealis* represents a unique evolutionary solution, combining aggressive saprotrophic capabilities with necrotrophic pathogenesis through expanded auxiliary activity families that enable both living tissue penetration and post-mortem biomass processing(Sahu et al., 2023). Secondary decomposer *Agaricus bisporus* specialized for soil humic compound processing, while host-specific *Daedalea quercina* demonstrates extreme enzymatic adaptation to oak-derived substrates(Nagy et al., 2016; Sahu et al., 2023). This phylogenetic sampling captures the complete evolutionary landscape of fungal lignocellulose degradation, providing definitive insights into the molecular basis of industrial biorefinery optimization.

CAZyXplorer provides a systematic approach to analyzing microbial strains based on CAZyme annotation profiles across six degradation pathways. The platform’s analysis of eight fungal genomes demonstrated several notable patterns. Principal component analysis explained 62.9% of enzymatic variance, showing that wood-decay strategies correlate with different enzyme distribution patterns, with white-rot fungi displaying broader enzymatic repertoires compared to brown-rot specialists. The weighted Industrial Potential Index applies pathway coefficients (cellulolytic: 40%, hemi-cellulolytic: 40%, ligninolytic: 20%) to generate comparative rankings. In the analyzed dataset, A. borealis received the highest score (193.6) based on its enzyme annotation profile (Cell: 338, Hemi: 52, Lignin: 188), followed by P. chrysosporium (Score: 123.4; Cell: 225, Hemi: 26, Lignin: 115), and A. bisporus (Score: 113.2; Cell: 197, Hemi: 33, Lignin: 106). Network analysis indicated that A. borealis possessed more multi-functional enzyme annotations (19) compared to brown-rot fungi in the dataset (8-12), with certain enzyme families (GH16, AA3, CE4) appearing across multiple pathways.

It is important to emphasize that these rankings reflect genomic potential based on CAZyme gene annotations rather than measured enzymatic activities or experimental biomass degradation performance. The scoring differences represent variations in annotated enzyme family counts and should be interpreted as preliminary indicators requiring experimental validation. While A. borealis showed a 57% higher computational score than P. chrysosporium in this analysis, this does not directly translate to equivalent differences in actual biorefinery performance, as enzyme expression levels, specific activities, and synergistic interactions are not captured by gene counts alone. CAZyXplorer serves as a computational pre-screening tool that may help reduce the initial candidate pool for experimental testing. The platform provides accessible comparative analysis of CAZyme profiles through point-and-click interfaces, allowing researchers to explore enzyme distribution patterns and functional annotations across microbial genomes. By offering quantitative metrics for preliminary strain assessment, the tool can complement traditional laboratory-based screening approaches. However, the platform’s predictions should be viewed as hypothesis-generating rather than definitive, with strain selection decisions ultimately requiring biochemical characterization and performance validation under relevant industrial conditions. CAZyXplorer addresses a need for accessible tools to analyze comparative genomic data in the context of biorefinery applications, though it represents one component of a broader strain development pipeline that necessarily includes experimental validation stages.

## Data Availability

The CAZyXplorer Shiny application can be downloaded from GitHub at https://github.com/aysistak89/CAZyXplorer and is released under the MIT License. After downloading (Code → Download as Zip) and extracting the files, users can open CAZyXplorer.R in RStudio and click "Run App" to launch the application. The data folder contains CAZyme annotations of Agaricomycetes strains (10, 30, and 50 strain datasets) and demo data used in this manuscript in CSV format.

## Supplementary Information

Additional supporting information can be found online in the supplementary Information section. Supplementary Information 1. A step-by-step guide to using CAZyXplorer. Supplementary Information 2. CAZyXplorer app (CAZyXplorer-main.zip).

