## Supplementary material for "CAZyXplorer: A Shiny Application for Cost-Effective Preliminary Screening of Microbial Strains to Advance Enzyme Discovery in Biorefining and Biotechnology": CAZxplorer.R

```
# CAZyXplorer main app 
# Advanced R Shiny Application for Industrial Biorefinery Research

# Load required libraries
library(shiny)
library(shinydashboard)
library(DT)
library(plotly)
library(dplyr)
library(ggplot2)
library(RColorBrewer)
library(shinycssloaders)
library(readr)
library(tidyr)
library(stringr)
library(viridis)
library(corrplot)
library(pheatmap)
library(scales)
library(shinyWidgets)
library(leaflet)
library(networkD3)
library(tibble)
library(DT)
library(rmarkdown)
library(knitr)
library(htmltools)

# Source UI and Server files
source("ui/ui.R")
source("server/server.R")

# Run the enhanced application
shinyApp(ui = ui, server = server)
```
