## Supplementary material for "CAZyXplorer: A Shiny Application for Cost-Effective Preliminary Screening of Microbial Strains to Advance Enzyme Discovery in Biorefining and Biotechnology": README.md

# CAZyxplorer: Advanced CAZy Enzyme Analysis Platform 

## 🧬 A comprehensive R Shiny application for bacterial plant cell wall degrading enzyme exploration and industrial biorefinery potential assessment
### Overview
CAZyxplorer is a modular, user-friendly R Shiny application designed for advanced analysis of Carbohydrate-Active enzymes (CAZy) in bacterial genomes. The platform enables researchers to evaluate industrial biorefinery potential through comprehensive enzyme classification, pathway mapping, and statistical analysis of CAZy annotation data.

Key Features
🔬 Comprehensive Enzyme Analysis

## Multi-pathway Classification: Automated categorization of enzymes into six primary degradation pathways:
Cellulolytic (40+ enzyme families including GH5-74, CBM1-72, AA9-15)
Hemi-cellulolytic (GH1-142, CBM2-81, CE1-15)
Ligninolytic (AA1-12, CE1/10/15)
Pectinolytic (GH1-147, PL1-26, CE6-13)
Starch-degrading (GH13-126, GT35, CBM20-83)
Inulin-degrading (GH32, GH91, CBM38)

## 🏭 Industrial Biorefinery Assessment

Weighted Scoring Algorithm: Multi-criteria evaluation system with optimized weightings (Cellulolytic 40%, Hemi-cellulolytic 40%, Ligninolytic 20%)
Shannon Diversity Integration: Enzyme distribution assessment using entropy measures
Percentile-based Rankings: Comparative performance metrics across genome collections
Industrial Synergy Indices: Holistic biorefinery potential combining core and auxiliary enzymatic capabilities

## 📊 Advanced Visualization & Statistics

Interactive Heatmaps: Enzyme-genome abundance patterns with accessibility-optimized viridis color scales
Force-directed Network Graphs: Functional clustering visualization with customizable node sizing
Principal Component Analysis (PCA): Dimensionality reduction for enzyme profile analysis
Correlation Matrix Analysis: Identification of co-occurring enzyme families
Biodiversity Indices: Shannon entropy-based diversity measurements

## 🔧 Robust Data Processing

Automated Input Validation: CSV format verification and genome column detection
Missing Value Imputation: Zero-replacement preprocessing with quality assessment
Completeness Scoring: Non-zero enzyme count percentage analysis
Interactive Data Tables: Real-time exploration of processed datasets

###Technical Architecture
###Component-based Design
Modular Shiny architecture promoting maintainability and expansion
Segregated UI components for enhanced reusability
Dedicated server logic subdirectories for analytical functions
Independent development support with seamless integration

### Technology Stack
Core Framework: R (≥4.1), Shiny v1.7.4, shinydashboard v0.7.2
Data Processing: data.table, dplyr, readr, tidyr
Visualization: ggplot2 v3.4.2, plotly v4.10.2, networkD3 v0.4
Statistical Analysis: Comprehensive PCA and correlation analysis packages
Reporting: rmarkdown, knitr for automated report generation

## Installation & Usage
###Prerequisites
R (version 4.1 or later)
RStudio (recommended)
Internet connection for initial package installation

Quick Start
r
# Clone the repository
git clone https://github.com/aysistak89/CAZyxplorer.git
cd CAZyxplorer

# Launch the application
R -e "shiny::runApp()"

## Automatic Dependency Management
Runtime package verification and installation
Reduced barriers for users with varying R experience
Local deployment for sensitive genomic data analysis

## Input Data Requirements
Format: CSV files with CAZy annotation matrices
Structure: Enzyme family identifiers in first column, genome abundance data in subsequent columns
Content: Quantitative enzyme family abundance across bacterial genomes
Validation: Automatic format checking and preprocessing

## Output Capabilities
## Comprehensive Reporting
Executive summary reports (HTML format)
Detailed scientific analyses with methodology
Processed enzyme matrices (CSV export)
Multi-format visualization exports
Industrial biorefinery potential assessments

## Export Options
Interactive tables and visualizations
Statistical analysis results
Network topology data
Comparative genome rankings

## Use Cases
Industrial Biotechnology: Strain selection for biorefinery applications
Comparative Genomics: Cross-genome enzyme profile analysis
Metabolic Engineering: Pathway optimization and enzyme discovery
Environmental Microbiology: Plant biomass degradation potential assessment
Synthetic Biology: Enzyme cocktail design and optimization

# License
This project is licensed under the MIT License - see the LICENSE file for details.

## Citation
If you use CAZyxplorer in your research, please cite:
bibtex@software{cazyxplorer,
  title={CAZyxplorer: Advanced CAZy Enzyme Analysis Platform for Industrial Biorefinery Research},
  author={[Ayyappa Kumar Sista Kameshwar]},
  year={2024},
  url={https://github.com/aysistak89/CAZyxplorer/}
}

### Support
Issues: Report bugs and feature requests via GitHub Issues
Documentation: Comprehensive user guide available in /docs
Community: Join discussions in the GitHub Discussions section

🚀 Accelerating biorefinery research through intelligent enzyme analysis
