## Supplementary material for "CAZyXplorer: A Shiny Application for Cost-Effective Preliminary Screening of Microbial Strains to Advance Enzyme Discovery in Biorefining and Biotechnology": server.R

```
# Enhanced Server with Comprehensive Analysis - FIXED VERSION
server <- function(input, output, session) {
  
  # Enhanced Reactive values with comprehensive enzyme families
  values <- reactiveValues(
    data = NULL,
    processed_data = NULL,
    enzyme_families = list(
      # Comprehensive Cellulolytic enzymes based on the provided classification
      cellulolytic = list(
        CBM = c("CBM1", "CBM2", "CBM3", "CBM4", "CBM6", "CBM8", "CBM9", "CBM10", "CBM16", "CBM17", "CBM28", "CBM30", "CBM37", "CBM44", "CBM46", "CBM49", "CBM59", "CBM63", "CBM64", "CBM72"),
        GH = c("GH5", "GH6", "GH7", "GH8", "GH9", "GH10", "GH12", "GH26", "GH44", "GH45", "GH48", "GH51", "GH74", "GH124", "GH1", "GH2", "GH3", "GH30", "GH39", "GH116"),
        AA = c("AA9", "AA10", "AA15", "AA3")
      ),
      
      # Comprehensive Hemi-Cellulolytic enzymes
      hemi_cellulolytic = list(
        GH = c("GH1", "GH2", "GH3", "GH5", "GH8", "GH9", "GH10", "GH11", "GH12", "GH16", "GH17", "GH26", "GH30", "GH39", "GH43", "GH44", "GH51", "GH52", "GH54", "GH55", "GH62", "GH74", "GH98", "GH113", "GH116", "GH120", "GH127", "GH134", "GH137", "GH141", "GH142", "GH146"),
        CBM = c("CBM2", "CBM4", "CBM6", "CBM9", "CBM13", "CBM15", "CBM22", "CBM31", "CBM35", "CBM36", "CBM37", "CBM42", "CBM54", "CBM59", "CBM60", "CBM72", "CBM16", "CBM23", "CBM27", "CBM29", "CBM62", "CBM76", "CBM80", "CBM44", "CBM65", "CBM75", "CBM78", "CBM81"),
        CE = c("CE1", "CE2", "CE3", "CE4", "CE5", "CE6", "CE7", "CE12", "CE15"),
        AA = c("AA9", "AA14")
      ),
      
      # Lignolytic enzymes
      lignolytic = list(
        AA = c("AA1", "AA2", "AA3", "AA4", "AA5", "AA6", "AA8", "AA9", "AA12"),
        CE = c("CE1", "CE15", "CE10")
      ),
      
      # Pectinolytic enzymes
      pectinolytic = list(
        GH = c("GH1", "GH2", "GH3", "GH4", "GH10", "GH28", "GH33", "GH35", "GH39", "GH42", "GH43", "GH50", "GH51", "GH54", "GH59", "GH62", "GH78", "GH93", "GH106", "GH147"),
        PL = c("PL1", "PL2", "PL3", "PL4", "PL9", "PL10", "PL11", "PL22", "PL26"),
        CE = c("CE6", "CE8", "CE12", "CE13"),
        CBM = c("CBM13", "CBM32", "CBM41", "CBM51", "CBM61", "CBM62", "CBM67", "CBM77", "CBM80")
      ),
      
      # Starch degrading enzymes
      starch_degrading = list(
        GH = c("GH13", "GH14", "GH57", "GH119", "GH126", "GH4", "GH31", "GH63", "GH97", "GH122", "GT35"),
        CBM = c("CBM20", "CBM21", "CBM25", "CBM26", "CBM34", "CBM45", "CBM53", "CBM69", "CBM74", "CBM82", "CBM83"),
        AA = c("AA13")
      ),
      
      # Inulin degrading enzymes
      inulin_degrading = list(
        GH = c("GH32", "GH91"),
        CBM = c("CBM38")
      )
    )
  )
  
  # Enhanced data upload handling
  observeEvent(input$csvFile, {
    if (!is.null(input$csvFile)) {
      tryCatch({
        values$data <- read_csv(input$csvFile$datapath)
        processData()
      }, error = function(e) {
        showNotification(paste("Error loading file:", e$message), type = "error")
      })
    }
  })
  
  observeEvent(input$mainCsvFile, {
    if (!is.null(input$mainCsvFile)) {
      tryCatch({
        values$data <- read_csv(input$mainCsvFile$datapath)
        processData()
      }, error = function(e) {
        showNotification(paste("Error loading file:", e$message), type = "error")
      })
    }
  })
  
  # Navigate to upload tab
  observeEvent(input$goToUpload, {
    updateTabItems(session, "sidebarMenu", selected = "upload")
  })
  
  # REPLACE the processData function with this fixed version:
  processData <- function() {
    if (!is.null(values$data)) {
      # Identify the enzyme column name
      enzyme_col <- if("Annotations/Genomes" %in% names(values$data)) {
        "Annotations/Genomes"
      } else if("CAZymes" %in% names(values$data)) {
        "CAZymes"
      } else {
        names(values$data)[1]
      }
      
      # Clean and process the data
      processed <- values$data %>%
        filter(!is.na(!!sym(enzyme_col))) %>%
        mutate(across(where(is.character) & !all_of(enzyme_col), ~as.numeric(as.character(.x)))) %>%
        mutate(across(where(is.numeric), ~replace_na(.x, 0)))
      
      values$processed_data <- processed
      
      # Update UI choices
      genome_cols <- names(processed)[!names(processed) %in% c(enzyme_col, "Total")]
      
      # Safe update of UI elements
      if(length(genome_cols) > 0) {
        updateSelectInput(session, "selectedStrain",
                          choices = genome_cols,
                          selected = genome_cols[1])
        
        # Add this for multi-strain functionality
        updateCheckboxGroupInput(session, "selectedStrains",
                                 choices = genome_cols,
                                 selected = genome_cols[1:min(3, length(genome_cols))])
        
        # DEBUG: Print comprehensive data info
        cat("=== PROCESSDATA DEBUG ===\n")
        debugDataStructure(processed)
        
        # Test enzyme family matching
        testEnzymeMatching(processed)
        
        # Test specific enzyme family calculations
        cat("\n=== TESTING ENZYME FAMILY CALCULATIONS ===\n")
        if(!is.null(values$enzyme_families$cellulolytic)) {
          test_potential <- calculateDegradationPotential(processed, values$enzyme_families$cellulolytic)
          if(!is.null(test_potential)) {
            cat("Cellulolytic potential test results:", paste(names(test_potential), "=", test_potential, collapse = ", "), "\n")
          } else {
            cat("Cellulolytic potential test returned NULL\n")
          }
        }
      }
    }
  }
  
  # Enhanced function to calculate degradation potential with comprehensive enzyme families
  calculateDegradationPotential <- function(data, enzyme_families) {
    if (is.null(data) || is.null(enzyme_families)) {
      cat("calculateDegradationPotential: NULL input\n")
      return(NULL)
    }
    
    # Handle different column name formats
    enzyme_col <- if("Annotations/Genomes" %in% names(data)) {
      "Annotations/Genomes"
    } else if("CAZymes" %in% names(data)) {
      "CAZymes"
    } else {
      names(data)[1]
    }
    
    # Get genome columns (exclude enzyme column and Total if present)
    genome_cols <- names(data)[!names(data) %in% c(enzyme_col, "Total")]
    
    if(length(genome_cols) == 0) {
      cat("DEBUG: No genome columns found\n")
      return(NULL)
    }
    
    # Handle nested enzyme families structure
    if(is.list(enzyme_families)) {
      # Check if it's a nested list (like your structure with CBM, GH, AA sublists)
      if(any(sapply(enzyme_families, is.list))) {
        # Flatten all sublists recursively
        all_enzyme_patterns <- unique(unlist(enzyme_families, recursive = TRUE))
        cat("DEBUG: Flattened nested structure, found", length(all_enzyme_patterns), "patterns\n")
      } else {
        # It's already a flat list
        all_enzyme_patterns <- unique(unlist(enzyme_families))
        cat("DEBUG: Using flat list, found", length(all_enzyme_patterns), "patterns\n")
      }
    } else {
      all_enzyme_patterns <- enzyme_families
      cat("DEBUG: Using vector input, found", length(all_enzyme_patterns), "patterns\n")
    }
    
    # Remove any NULL, NA, or empty values
    all_enzyme_patterns <- all_enzyme_patterns[!is.na(all_enzyme_patterns) & 
                                                 !is.null(all_enzyme_patterns) & 
                                                 nchar(all_enzyme_patterns) > 0]
    
    if(length(all_enzyme_patterns) == 0) {
      cat("DEBUG: No valid enzyme patterns found\n")
      return(NULL)
    }
    
    # Count matches for each pattern (for debugging)
    matching_rows <- rep(FALSE, nrow(data))
    
    for(pattern in all_enzyme_patterns) {
      # Enhanced pattern matching: exact match OR pattern followed by underscore/number
      pattern_matches <- grepl(paste0("^", pattern, "($|_|\\d)"), data[[enzyme_col]], perl = TRUE)
      
      if(any(pattern_matches)) {
        matching_enzyme_count <- sum(as.numeric(data[pattern_matches, genome_cols[1]]), na.rm = TRUE)
        cat("Found matches for pattern:", pattern, "- Count:", matching_enzyme_count, "\n")
      }
      
      matching_rows <- matching_rows | pattern_matches
    }
    
    cat("DEBUG: Total matching rows:", sum(matching_rows), "out of", nrow(data), "\n")
    
    if(!any(matching_rows)) {
      # Return zeros for all genomes
      result <- as.list(rep(0, length(genome_cols)))
      names(result) <- genome_cols
      return(result)
    }
    
    # Calculate potential scores for each genome
    relevant_data <- data[matching_rows, , drop = FALSE]
    potential_scores <- list()
    
    for(genome in genome_cols) {
      if(genome %in% names(relevant_data)) {
        # Ensure the column is numeric and handle NAs
        genome_values <- as.numeric(as.character(relevant_data[[genome]]))
        genome_values[is.na(genome_values)] <- 0
        potential_scores[[genome]] <- sum(genome_values, na.rm = TRUE)
      } else {
        potential_scores[[genome]] <- 0
      }
    }
    
    cat("calculateDegradationPotential: Calculated scores for", length(potential_scores), "genomes\n")
    return(potential_scores)
  }
  
  # Function to calculate enzyme class distribution
  calculateEnzymeClassDistribution <- function(data, strain_name) {
    if (is.null(data) || is.null(strain_name)) return(NULL)
    
    enzyme_col <- if("Annotations/Genomes" %in% names(data)) {
      "Annotations/Genomes"
    } else if("CAZymes" %in% names(data)) {
      "CAZymes"
    } else {
      names(data)[1]
    }
    
    if (!strain_name %in% names(data)) return(NULL)
    
    # Extract enzyme classes (GH, AA, CBM, PL, CE, GT)
    class_data <- data %>%
      mutate(
        Enzyme_Class = case_when(
          str_detect(!!sym(enzyme_col), "^GH") ~ "GH",
          str_detect(!!sym(enzyme_col), "^AA") ~ "AA",
          str_detect(!!sym(enzyme_col), "^CBM") ~ "CBM",
          str_detect(!!sym(enzyme_col), "^PL") ~ "PL",
          str_detect(!!sym(enzyme_col), "^CE") ~ "CE",
          str_detect(!!sym(enzyme_col), "^GT") ~ "GT",
          TRUE ~ "Other"
        )
      ) %>%
      filter(!!sym(strain_name) > 0) %>%
      group_by(Enzyme_Class) %>%
      summarise(Count = sum(!!sym(strain_name), na.rm = TRUE), .groups = "drop")
    
    return(class_data)
  }
  
  calculateIndustrialPotential <- function(data, enzyme_families) {
    if (is.null(data) || is.null(enzyme_families)) {
      return(0)
    }
    
    cat("=== Industrial Potential Calculation ===\n")
    
    tryCatch({
      # Calculate individual potentials using the enhanced function
      cell_potential <- calculateDegradationPotential(data, enzyme_families$cellulolytic)
      hemi_potential <- calculateDegradationPotential(data, enzyme_families$hemi_cellulolytic)
      lignin_potential <- calculateDegradationPotential(data, enzyme_families$lignolytic)
      
      if(is.null(cell_potential) || is.null(hemi_potential) || is.null(lignin_potential)) {
        cat("One or more potentials returned NULL\n")
        return(0)
      }
      
      # Get genome columns
      enzyme_col <- if("Annotations/Genomes" %in% names(data)) {
        "Annotations/Genomes"
      } else if("CAZymes" %in% names(data)) {
        "CAZymes"
      } else {
        names(data)[1]
      }
      
      genome_cols <- names(data)[!names(data) %in% c(enzyme_col, "Total")]
      
      # Calculate totals across all genomes
      cell_total <- sum(sapply(genome_cols, function(g) {
        if(g %in% names(cell_potential)) cell_potential[[g]] else 0
      }), na.rm = TRUE)
      
      hemi_total <- sum(sapply(genome_cols, function(g) {
        if(g %in% names(hemi_potential)) hemi_potential[[g]] else 0
      }), na.rm = TRUE)
      
      lignin_total <- sum(sapply(genome_cols, function(g) {
        if(g %in% names(lignin_potential)) lignin_potential[[g]] else 0
      }), na.rm = TRUE)
      
      cat("Totals - Cell:", cell_total, "Hemi:", hemi_total, "Lignin:", lignin_total, "\n")
      
      # Calculate weighted score (lignin has half weight as it's harder to degrade)
      raw_score <- cell_total + hemi_total + (lignin_total * 0.5)
      cat("Overall raw score:", raw_score, "\n")
      
      # Normalize to percentage (adjust based on your data range)
      # Based on your debug output showing ~400+ total, let's adjust the max_possible
      max_possible <- 600  
      normalized_score <- min(100, round((raw_score / max_possible) * 100))
      
      cat("Normalized industrial potential:", normalized_score, "%\n")
      return(max(0, normalized_score))
      
    }, error = function(e) {
      cat("ERROR in calculateIndustrialPotential:", e$message, "\n")
      return(0)
    })
  }
  
  # Check if data is uploaded
  output$dataUploaded <- reactive({
    return(!is.null(values$processed_data))
  })
  outputOptions(output, "dataUploaded", suspendWhenHidden = FALSE)
  
  # Enhanced data info
  output$dataInfo <- renderText({
    if (!is.null(values$processed_data)) {
      genome_count <- ncol(values$processed_data) - ifelse("Total" %in% names(values$processed_data), 2, 1)
      enzyme_count <- nrow(values$processed_data)
      paste("📊 Loaded", enzyme_count, "enzyme families across", genome_count, "genomes")
    }
  })
  
  # Data quality metrics
  output$dataQuality <- renderText({
    if (!is.null(values$processed_data)) {
      genome_cols <- names(values$processed_data)[!names(values$processed_data) %in% c("Annotations/Genomes", "CAZymes", "Total")]
      if(length(genome_cols) > 0) {
        completeness <- round(mean(sapply(values$processed_data[genome_cols], function(x) sum(x > 0) / length(x))) * 100, 1)
        diversity <- length(unique(unlist(strsplit(values$processed_data[[1]], ""))))
        paste("✅ Data Completeness:", completeness, "%\n🧬 Enzyme Diversity Score:", diversity)
      }
    }
  })
  
  # FIXED: Enhanced value boxes
  output$totalEnzymes <- renderValueBox({
    if (!is.null(values$processed_data)) {
      enzyme_col <- if("Annotations/Genomes" %in% names(values$processed_data)) {
        "Annotations/Genomes"
      } else if("CAZymes" %in% names(values$processed_data)) {
        "CAZymes"
      } else {
        names(values$processed_data)[1]
      }
      
      genome_cols <- names(values$processed_data)[!names(values$processed_data) %in% c(enzyme_col, "Total")]
      if(length(genome_cols) > 0) {
        # Ensure numeric conversion
        numeric_data <- values$processed_data[genome_cols] %>%
          mutate(across(everything(), \(x) as.numeric(as.character(x))))
        total <- sum(numeric_data, na.rm = TRUE)
        valueBox(
          value = format(total, big.mark = ","),
          subtitle = "Total Enzymes",
          icon = icon("dna"),
          color = "blue"
        )
      }
    }
  })
  
  output$totalGenomes <- renderValueBox({
    if (!is.null(values$processed_data)) {
      enzyme_col <- if("Annotations/Genomes" %in% names(values$processed_data)) {
        "Annotations/Genomes"
      } else if("CAZymes" %in% names(values$processed_data)) {
        "CAZymes"
      } else {
        names(values$processed_data)[1]
      }
      genomes <- ncol(values$processed_data) - ifelse("Total" %in% names(values$processed_data), 2, 1)
      valueBox(
        value = max(0, genomes),
        subtitle = "Genomes Analyzed",
        icon = icon("bacteria"),
        color = "green"
      )
    }
  })
  
  output$avgEnzymesPerGenome <- renderValueBox({
    if (!is.null(values$processed_data)) {
      enzyme_col <- if("Annotations/Genomes" %in% names(values$processed_data)) {
        "Annotations/Genomes"
      } else if("CAZymes" %in% names(values$processed_data)) {
        "CAZymes"
      } else {
        names(values$processed_data)[1]
      }
      genome_cols <- names(values$processed_data)[!names(values$processed_data) %in% c(enzyme_col, "Total")]
      if(length(genome_cols) > 0) {
        # Ensure numeric conversion
        numeric_data <- values$processed_data[genome_cols] %>%
          mutate(across(everything(), \(x) as.numeric(as.character(x))))
        avg <- round(mean(colSums(numeric_data, na.rm = TRUE)), 1)
        valueBox(
          value = format(avg, big.mark = ","),
          subtitle = "Avg Enzymes/Genome",
          icon = icon("calculator"),
          color = "yellow"
        )
      }
    }
  })
  
  output$topEnzymeFamily <- renderValueBox({
    if (!is.null(values$processed_data)) {
      enzyme_col <- if("Annotations/Genomes" %in% names(values$processed_data)) {
        "Annotations/Genomes"
      } else if("CAZymes" %in% names(values$processed_data)) {
        "CAZymes"
      } else {
        names(values$processed_data)[1]
      }
      
      genome_cols <- names(values$processed_data)[!names(values$processed_data) %in% c(enzyme_col, "Total")]
      if(length(genome_cols) > 0) {
        # Ensure numeric conversion and calculate row sums
        numeric_data <- values$processed_data[genome_cols] %>%
          mutate(across(everything(), \(x) as.numeric(as.character(x))))
        
        processed_with_sums <- values$processed_data %>%
          mutate(row_sum = rowSums(numeric_data, na.rm = TRUE))
        
        top_family <- processed_with_sums %>%
          slice_max(row_sum, n = 1) %>%
          pull(!!sym(enzyme_col)) %>%
          first()
        
        # Truncate long names
        display_name <- if(nchar(top_family) > 12) {
          paste0(substr(top_family, 1, 9), "...")
        } else {
          top_family
        }
        
        valueBox(
          value = display_name,
          subtitle = "Top Family",
          icon = icon("trophy"),
          color = "red"
        )
      }
    }
  })
  
  output$biodiversityIndex <- renderValueBox({
    if (!is.null(values$processed_data)) {
      enzyme_col <- if("Annotations/Genomes" %in% names(values$processed_data)) {
        "Annotations/Genomes"
      } else if("CAZymes" %in% names(values$processed_data)) {
        "CAZymes"
      } else {
        names(values$processed_data)[1]
      }
      genome_cols <- names(values$processed_data)[!names(values$processed_data) %in% c(enzyme_col, "Total")]
      if(length(genome_cols) > 0) {
        # Ensure numeric conversion
        numeric_data <- values$processed_data[genome_cols] %>%
          mutate(across(everything(), \(x) as.numeric(as.character(x))))
        
        # Calculate Shannon diversity index
        total_enzymes <- sum(numeric_data, na.rm = TRUE)
        if(total_enzymes > 0) {
          shannon_index <- -sum(sapply(numeric_data, function(x) {
            p <- sum(x, na.rm = TRUE) / total_enzymes
            if(p > 0) p * log(p) else 0
          }), na.rm = TRUE)
          
          valueBox(
            value = round(shannon_index, 2),
            subtitle = "Biodiversity Index",
            icon = icon("leaf"),
            color = "purple"
          )
        }
      }
    }
  })
  
  output$industrialPotential <- renderValueBox({
    if (!is.null(values$processed_data)) {
      
      # Calculate industrial potential
      potential_score <- calculateIndustrialPotential(values$processed_data, values$enzyme_families)
      
      valueBox(
        value = paste0(potential_score, "%"),
        subtitle = "Industrial Potential",
        icon = icon("industry"),
        color = if(potential_score >= 70) "green" else if(potential_score >= 40) "yellow" else if(potential_score > 0) "orange" else "red"
      )
      
    } else {
      valueBox(
        value = "N/A",
        subtitle = "Industrial Potential",
        icon = icon("industry"),
        color = "light-blue"
      )
    }
  })
  
  
  
  output$topFamilies <- renderPlotly({
    if (!is.null(values$processed_data)) {
      enzyme_col <- if("Annotations/Genomes" %in% names(values$processed_data)) {
        "Annotations/Genomes"
      } else if("CAZymes" %in% names(values$processed_data)) {
        "CAZymes"
      } else {
        names(values$processed_data)[1]
      }
      
      genome_cols <- names(values$processed_data)[!names(values$processed_data) %in% c(enzyme_col, "Total")]
      
      if(length(genome_cols) > 0) {
        top_data <- values$processed_data %>%
          mutate(Total_Count = rowSums(select(., all_of(genome_cols)), na.rm = TRUE)) %>%
          slice_max(Total_Count, n = 15) %>%
          arrange(desc(Total_Count))
        
        p <- ggplot(top_data, aes(x = reorder(!!sym(enzyme_col), Total_Count), y = Total_Count)) +
          geom_col(fill = "#00d4ff", alpha = 0.8, color = "#ff6b35", size = 0.5) +
          coord_flip() +
          labs(title = "Top 15 Enzyme Families",
               x = "Enzyme Family",
               y = "Total Count") +
          theme_minimal() +
          theme(
            plot.background = element_rect(fill = "transparent", color = NA),
            panel.background = element_rect(fill = "transparent", color = NA),
            text = element_text(color = "white"),
            axis.text = element_text(color = "white")
          )
        
        ggplotly(p, tooltip = c("x", "y")) %>%
          layout(
            plot_bgcolor = 'rgba(0,0,0,0)',
            paper_bgcolor = 'rgba(0,0,0,0)',
            font = list(color = 'white')
          )%>%
          config(
            toImageButtonOptions = list(
              format = 'png',
              filename = 'Top_15_Enzyme_Families',
              height = 400,
              width = 400,
              scale = 4  
            )
          )
      }
    }
  })
  
  # 4. Single Strain Distribution (for Enzyme Distribution in Single Strain Deep Dive)
  output$singleStrainDistribution <- renderPlotly({
    if (!is.null(values$processed_data) && !is.null(input$selectedStrain)) {
      enzyme_col <- if("Annotations/Genomes" %in% names(values$processed_data)) {
        "Annotations/Genomes"
      } else if("CAZymes" %in% names(values$processed_data)) {
        "CAZymes"
      } else {
        names(values$processed_data)[1]
      }
      
      if(input$selectedStrain %in% names(values$processed_data)) {
        strain_data <- values$processed_data %>%
          select(!!sym(enzyme_col), all_of(input$selectedStrain)) %>%
          filter(!!sym(input$selectedStrain) > 0) %>%
          arrange(desc(!!sym(input$selectedStrain))) %>%
          slice_head(n = 25)
        
        if(nrow(strain_data) > 0) {
          p <- ggplot(strain_data, aes(x = reorder(!!sym(enzyme_col), !!sym(input$selectedStrain)), 
                                       y = !!sym(input$selectedStrain))) +
            geom_col(fill = "#00d4ff", alpha = 0.8, color = "#ff6b35", size = 0.5) +
            coord_flip() +
            labs(title = paste("Top 25 Enzyme Families in", input$selectedStrain),
                 x = "Enzyme Family", y = "Count") +
            theme_minimal() +
            theme(
              plot.background = element_rect(fill = "transparent", color = NA),
              panel.background = element_rect(fill = "transparent", color = NA),
              text = element_text(color = "white"),
              axis.text = element_text(color = "white"),
              axis.text.y = element_text(size = 8)  # Make enzyme names smaller
            )
          
          ggplotly(p, tooltip = c("x", "y")) %>%
            layout(
              plot_bgcolor = 'rgba(0,0,0,0)',
              paper_bgcolor = 'rgba(0,0,0,0)',
              font = list(color = 'white')
            )%>%
            config(
              toImageButtonOptions = list(
                format = 'png',
                filename = 'Top_25_Enzyme_Families',
                height = 600,
                width = 850,
                scale = 4  
              )
            )
        } else {
          # No data for this strain
          plot_ly() %>%
            add_text(x = 0.5, y = 0.5, text = paste("No enzyme data found for", input$selectedStrain), 
                     textfont = list(color = "orange", size = 14)) %>%
            layout(
              xaxis = list(showgrid = FALSE, showticklabels = FALSE, zeroline = FALSE),
              yaxis = list(showgrid = FALSE, showticklabels = FALSE, zeroline = FALSE),
              plot_bgcolor = 'rgba(0,0,0,0)',
              paper_bgcolor = 'rgba(0,0,0,0)',
              font = list(color = 'white')
            )
        }
      } else {
        # Strain not found in data
        plot_ly() %>%
          add_text(x = 0.5, y = 0.5, text = paste("Strain", input$selectedStrain, "not found in data"), 
                   textfont = list(color = "red", size = 14)) %>%
          layout(
            xaxis = list(showgrid = FALSE, showticklabels = FALSE, zeroline = FALSE),
            yaxis = list(showgrid = FALSE, showticklabels = FALSE, zeroline = FALSE),
            plot_bgcolor = 'rgba(0,0,0,0)',
            paper_bgcolor = 'rgba(0,0,0,0)',
            font = list(color = 'white')
          )
      }
    } else {
      # No data or strain selected
      plot_ly() %>%
        add_text(x = 0.5, y = 0.5, text = "Please select a strain and upload data", 
                 textfont = list(color = "white", size = 14)) %>%
        layout(
          xaxis = list(showgrid = FALSE, showticklabels = FALSE, zeroline = FALSE),
          yaxis = list(showgrid = FALSE, showticklabels = FALSE, zeroline = FALSE),
          plot_bgcolor = 'rgba(0,0,0,0)',
          paper_bgcolor = 'rgba(0,0,0,0)',
          font = list(color = 'white')
        )
    }
  })
  
  
  # singleStrainCapabilities output with this fixed version:
  output$singleStrainCapabilities <- renderPlotly({
    if (!is.null(values$processed_data) && !is.null(input$selectedStrain)) {
      if(input$selectedStrain %in% names(values$processed_data)) {
        tryCatch({
          cat("=== Single Strain Capabilities for:", input$selectedStrain, "===\n")
          
          # Calculate degradation potentials for the selected strain
          capabilities <- data.frame(
            Type = c("Cellulolytic", "Hemi-Cellulolytic", "Lignolytic", "Pectinolytic", "Starch", "Inulin"),
            Score = c(0, 0, 0, 0, 0, 0),
            Efficiency = c("Low", "Low", "Low", "Low", "Low", "Low"),
            stringsAsFactors = FALSE
          )
          
          # Calculate scores for each capability
          enzyme_types <- list(
            values$enzyme_families$cellulolytic,
            values$enzyme_families$hemi_cellulolytic,
            values$enzyme_families$lignolytic,
            values$enzyme_families$pectinolytic,
            values$enzyme_families$starch_degrading,
            values$enzyme_families$inulin_degrading
          )
          
          for(i in 1:nrow(capabilities)) {
            potential <- calculateDegradationPotential(values$processed_data, enzyme_types[[i]])
            if(!is.null(potential) && input$selectedStrain %in% names(potential)) {
              score <- max(0, potential[[input$selectedStrain]])  # Ensure non-negative
              capabilities$Score[i] <- score
              
              # More realistic efficiency thresholds based on debug output
              capabilities$Efficiency[i] <- case_when(
                score >= 50 ~ "High",
                score >= 15 ~ "Medium", 
                score >= 5 ~ "Low",
                score > 0 ~ "Minimal",
                TRUE ~ "None"
              )
              cat("Capability", capabilities$Type[i], "score:", score, "efficiency:", capabilities$Efficiency[i], "\n")
            }
          }
          
          total_score <- sum(capabilities$Score)
          cat("Total capabilities score:", total_score, "\n")
          
          # Only create plot if we have data
          if(total_score > 0 || any(capabilities$Score > 0)) {
            # Create the capabilities plot
            p <- ggplot(capabilities, aes(x = reorder(Type, Score), y = Score, fill = Efficiency)) +
              geom_col(alpha = 0.8, color = "white", size = 0.5) +
              geom_text(aes(label = Score), hjust = -0.2, color = "white", fontface = "bold") +
              coord_flip() +
              labs(
                title = paste("Biorefinery Capabilities -", input$selectedStrain),
                x = "Capability Type", 
                y = "Enzyme Count"
              ) +
              theme_minimal() +
              theme(
                plot.background = element_rect(fill = "transparent", color = NA),
                panel.background = element_rect(fill = "transparent", color = NA),
                text = element_text(color = "white"),
                axis.text = element_text(color = "white"),
                legend.text = element_text(color = "white"),
                legend.title = element_text(color = "white")
              ) +
              scale_fill_manual(values = c(
                "High" = "#00ff88", 
                "Medium" = "#ffaa00", 
                "Low" = "#00d4ff",
                "Minimal" = "#ff6b35",
                "None" = "#666666"
              ))
            
            ggplotly(p, tooltip = c("x", "y", "fill")) %>%
              layout(
                plot_bgcolor = 'rgba(0,0,0,0)',
                paper_bgcolor = 'rgba(0,0,0,0)',
                font = list(color = 'white')
              )%>%
              config(
                toImageButtonOptions = list(
                  format = 'png',
                  filename = 'Biorefinery_Capabilities',
                  height = 500,
                  width = 650,
                  scale = 4  
                )
              )
          } else {
            # Create message plot
            p <- ggplot(data.frame(x = 1, y = 1, label = paste("No enzyme data found for", input$selectedStrain)), aes(x, y)) +
              geom_text(aes(label = label), color = "white", size = 5) +
              theme_void() +
              theme(
                plot.background = element_rect(fill = "transparent", color = NA),
                panel.background = element_rect(fill = "transparent", color = NA)
              )
            
            ggplotly(p)
          }
          
        }, error = function(e) {
          cat("ERROR in singleStrainCapabilities:", e$message, "\n")
          # Return error plot
          p <- ggplot(data.frame(x = 1, y = 1, label = paste("Error:", e$message)), aes(x, y)) +
            geom_text(aes(label = label), color = "red", size = 4) +
            theme_void() +
            theme(
              plot.background = element_rect(fill = "transparent", color = NA),
              panel.background = element_rect(fill = "transparent", color = NA)
            )
          
          ggplotly(p)
        })
      }
    }
  })
  
  
  # Enzyme class distribution for single strain
  output$enzymeClassDistribution <- renderPlotly({
    if (!is.null(values$processed_data) && !is.null(input$selectedStrain)) {
      class_data <- calculateEnzymeClassDistribution(values$processed_data, input$selectedStrain)
      
      if(!is.null(class_data) && nrow(class_data) > 0) {
        p <- ggplot(class_data, aes(x = reorder(Enzyme_Class, Count), y = Count, fill = Enzyme_Class)) +
          geom_col(alpha = 0.8, color = "white", size = 0.5) +
          labs(title = "Enzyme Class Distribution",
               x = "Enzyme Class", y = "Count") +
          theme_minimal() +
          theme(
            plot.background = element_rect(fill = "transparent", color = NA),
            panel.background = element_rect(fill = "transparent", color = NA),
            text = element_text(color = "white"),
            axis.text = element_text(color = "white"),
            legend.position = "none"
          ) +
          scale_fill_viridis_d(option = "plasma")
        
        ggplotly(p) %>%
          layout(
            plot_bgcolor = 'rgba(0,0,0,0)',
            paper_bgcolor = 'rgba(0,0,0,0)',
            font = list(color = 'white')
          )%>%
          config(
            toImageButtonOptions = list(
              format = 'png',
              filename = 'Enzyme_Class_Distribution',
              height = 500,
              width = 650,
              scale = 4  
            )
          )
      }
    }
  })
  
  # Enhanced strain summary
  output$strainSummary <- renderUI({
    if (!is.null(values$processed_data) && !is.null(input$selectedStrain)) {
      if(input$selectedStrain %in% names(values$processed_data)) {
        # Ensure the column is numeric
        strain_data <- as.numeric(values$processed_data[[input$selectedStrain]])
        total_enzymes <- sum(strain_data, na.rm = TRUE)
        unique_families <- sum(strain_data > 0, na.rm = TRUE)
        
        # Calculate percentile ranking with proper column identification
        enzyme_col <- if("Annotations/Genomes" %in% names(values$processed_data)) {
          "Annotations/Genomes"
        } else if("CAZymes" %in% names(values$processed_data)) {
          "CAZymes"
        } else {
          names(values$processed_data)[1]
        }
        
        genome_cols <- names(values$processed_data)[!names(values$processed_data) %in% c(enzyme_col, "Total")]
        
        # Ensure numeric conversion for all genome columns
        genome_data <- values$processed_data[genome_cols] %>%
          mutate(across(everything(), ~as.numeric(as.character(.x))))
        
        genome_totals <- colSums(genome_data, na.rm = TRUE)
        percentile <- round(100 * (sum(genome_totals < total_enzymes) / length(genome_totals)), 1)
        
        HTML(paste(
          "<div style='display: grid; grid-template-columns: 1fr 1fr; gap: 15px;'>",
          "<div style='text-align: center;'>",
          "<p class='industrial-highlight'>", format(total_enzymes, big.mark = ","), "</p>",
          "<p style='color: #cccccc;'>Total Enzymes</p>",
          "</div>",
          "<div style='text-align: center;'>",
          "<p class='industrial-highlight'>", unique_families, "</p>",
          "<p style='color: #cccccc;'>Unique Families</p>",
          "</div>",
          "</div>",
          "<div style='margin-top: 15px; text-align: center;'>",
          "<p><strong style='color: #00d4ff;'>Performance Percentile:</strong> ", percentile, "%</p>",
          "</div>"
        ))
      }
    }
  })
  
  # REPLACE the strainMetrics output with this fixed version:
  output$strainMetrics <- renderUI({
    if (!is.null(values$processed_data) && !is.null(input$selectedStrain)) {
      if(input$selectedStrain %in% names(values$processed_data)) {
        tryCatch({
          cat("=== Strain Metrics Calculation for:", input$selectedStrain, "===\n")
          
          # Calculate comprehensive metrics
          cell_potential <- calculateDegradationPotential(values$processed_data, values$enzyme_families$cellulolytic)
          hemi_potential <- calculateDegradationPotential(values$processed_data, values$enzyme_families$hemi_cellulolytic)
          lignin_potential <- calculateDegradationPotential(values$processed_data, values$enzyme_families$lignolytic)
          
          cell_score <- if(!is.null(cell_potential) && input$selectedStrain %in% names(cell_potential)) max(0, cell_potential[[input$selectedStrain]]) else 0
          hemi_score <- if(!is.null(hemi_potential) && input$selectedStrain %in% names(hemi_potential)) max(0, hemi_potential[[input$selectedStrain]]) else 0
          lignin_score <- if(!is.null(lignin_potential) && input$selectedStrain %in% names(lignin_potential)) max(0, lignin_potential[[input$selectedStrain]]) else 0
          
          cat("Individual scores - Cell:", cell_score, "Hemi:", hemi_score, "Lignin:", lignin_score, "\n")
          
          # Calculate overall biorefinery score
          biorefinery_score <- cell_score + hemi_score + (lignin_score * 0.5)
          cat("Biorefinery score:", biorefinery_score, "\n")
          
          # FIXED: More realistic strain rating thresholds based on your debug data
          rating <- case_when(
            biorefinery_score >= 150 ~ "🏆 ELITE",      # Very high threshold
            biorefinery_score >= 80 ~ "⭐ PREMIUM",     # High threshold
            biorefinery_score >= 40 ~ "✅ GOOD",        # Medium threshold
            biorefinery_score >= 15 ~ "📈 BASIC",       # Low threshold
            biorefinery_score > 0 ~ "🔬 DETECTED",     # Minimal threshold
            TRUE ~ "⚠️ NO DATA"
          )
          
          # Create color coding
          score_color <- case_when(
            biorefinery_score >= 150 ~ "#00ff88",
            biorefinery_score >= 80 ~ "#ffaa00",
            biorefinery_score >= 40 ~ "#00d4ff",
            biorefinery_score >= 15 ~ "#ff6b35",
            biorefinery_score > 0 ~ "#9c59d1",
            TRUE ~ "#ff4444"
          )
          
          HTML(paste(
            "<div style='text-align: center; margin-bottom: 15px;'>",
            "<p style='font-size: 1.5em; font-weight: bold; color: ", score_color, ";'>", rating, "</p>",
            "<p style='color: #cccccc;'>Biorefinery Rating</p>",
            "</div>",
            "<div style='background: rgba(", 
            if(biorefinery_score > 0) "0, 212, 255" else "255, 68, 68", 
            ", 0.1); padding: 10px; border-radius: 8px; border-left: 3px solid ", score_color, ";'>",
            "<p style='font-size: 1.2em; font-weight: bold; color: ", score_color, ";'>", round(biorefinery_score, 1), "</p>",
            "<p style='color: #cccccc; font-size: 0.9em;'>Overall Biorefinery Score</p>",
            "</div>",
            "<div style='margin-top: 10px; font-size: 11px; color: #888888;'>",
            "Cell: ", cell_score, " | Hemi: ", hemi_score, " | Lignin: ", lignin_score,
            "</div>",
            if(biorefinery_score == 0) {
              paste0(
                "<div style='margin-top: 10px; padding: 8px; background: rgba(255, 170, 0, 0.1); border-radius: 5px;'>",
                "<p style='color: #ffaa00; font-size: 10px;'>Debug: Check enzyme name patterns in your data</p>",
                "</div>"
              )
            } else ""
          ))
          
        }, error = function(e) {
          HTML(paste(
            "<div style='text-align: center; margin-bottom: 15px;'>",
            "<p style='font-size: 1.5em; font-weight: bold; color: #ff4444;'>❌ ERROR</p>",
            "<p style='color: #cccccc;'>Calculation Error</p>",
            "<p style='color: #ff4444; font-size: 0.8em;'>", e$message, "</p>",
            "</div>"
          ))
        })
      }
    }
  })
  
  debugDataStructure <- function(data) {
    cat("=== DATA STRUCTURE DEBUG ===\n")
    cat("Columns:", paste(names(data), collapse = ", "), "\n")
    cat("Number of rows:", nrow(data), "\n")
    
    # Identify enzyme column
    enzyme_col <- if("Annotations/Genomes" %in% names(data)) {
      "Annotations/Genomes"
    } else if("CAZymes" %in% names(data)) {
      "CAZymes"
    } else {
      names(data)[1]
    }
    cat("Enzyme column identified as:", enzyme_col, "\n")
    
    # Show sample enzyme names
    if(nrow(data) > 0) {
      sample_enzymes <- head(data[[enzyme_col]], 10)
      cat("Sample enzyme names:", paste(sample_enzymes, collapse = ", "), "\n")
    }
    
    # Identify genome columns
    genome_cols <- names(data)[!names(data) %in% c(enzyme_col, "Total")]
    cat("Genome columns:", paste(genome_cols, collapse = ", "), "\n")
    
    # Check if values are numeric
    if(length(genome_cols) > 0) {
      first_genome <- genome_cols[1]
      sample_values <- head(data[[first_genome]], 10)
      cat("Sample values from", first_genome, ":", paste(sample_values, collapse = ", "), "\n")
      cat("Are values numeric?", is.numeric(data[[first_genome]]), "\n")
    }
  }
  
  testEnzymeMatching <- function(data) {
    cat("=== ENZYME MATCHING TEST ===\n")
    
    enzyme_col <- if("Annotations/Genomes" %in% names(data)) {
      "Annotations/Genomes"
    } else if("CAZymes" %in% names(data)) {
      "CAZymes"
    } else {
      names(data)[1]
    }
    
    # Test specific patterns
    test_patterns <- c("GH1", "GH5", "CBM1", "AA9")
    
    cat("Testing patterns:", paste(test_patterns, collapse = ", "), "\n")
    
    for(pattern in test_patterns) {
      # Create enhanced pattern for exact matches and numbered variants
      enhanced_pattern <- paste0("^", pattern, "($|_\\d+$)")
      matches <- str_detect(data[[enzyme_col]], enhanced_pattern)
      matching_enzymes <- data[[enzyme_col]][matches]
      
      cat("Pattern", pattern, ":", sum(matches), "matches -", paste(head(matching_enzymes, 5), collapse = ", "), "\n")
    }
  }
  
  # Enhanced single strain degradation summary
  output$singleStrainDegradationSummary <- renderUI({
    if (!is.null(values$processed_data) && !is.null(input$selectedStrain)) {
      if(input$selectedStrain %in% names(values$processed_data)) {
        tryCatch({
          
          # Calculate all degradation potentials with detailed breakdown
          enzyme_types <- list(
            cellulolytic = values$enzyme_families$cellulolytic,
            hemi_cellulolytic = values$enzyme_families$hemi_cellulolytic,
            lignolytic = values$enzyme_families$lignolytic,
            pectinolytic = values$enzyme_families$pectinolytic,
            starch_degrading = values$enzyme_families$starch_degrading,
            inulin_degrading = values$enzyme_families$inulin_degrading
          )
          
          scores <- sapply(enzyme_types, function(enzyme_family) {
            potential <- calculateDegradationPotential(values$processed_data, enzyme_family)
            if(!is.null(potential) && input$selectedStrain %in% names(potential)) {
              max(0, potential[[input$selectedStrain]])  # Ensure non-negative
            } else {
              0
            }
          })
          
          # Debug: Check if we're getting any scores
          total_score <- sum(scores, na.rm = TRUE)
          
          # Create comprehensive industrial assessment
          HTML(paste(
            "<div class='degradation-card quantum-glow'>",
            "<h4>🏭 COMPREHENSIVE INDUSTRIAL BIOREFINERY ASSESSMENT</h4>",
            if(total_score == 0) {
              paste(
                "<div style='background: rgba(255, 107, 53, 0.2); padding: 15px; border-radius: 10px; margin: 15px 0; border-left: 4px solid #ff6b35;'>",
                "<p><strong style='color: #ff6b35;'>⚠️ DATA ANALYSIS:</strong></p>",
                "<p style='color: #ffffff;'>Selected strain: ", input$selectedStrain, "</p>",
                "<p style='color: #cccccc;'>Checking enzyme families...</p>",
                "</div>"
              )
            } else "",
            "<div style='display: grid; grid-template-columns: 1fr 1fr 1fr; gap: 20px; margin-top: 25px;'>",
            
            # Primary degradation capabilities
            "<div style='background: rgba(0, 212, 255, 0.1); padding: 20px; border-radius: 10px; border-left: 4px solid #00d4ff;'>",
            "<p><strong style='color: #00d4ff; font-family: Orbitron, monospace; font-size: 16px;'>🌾 CELLULOLYTIC ARSENAL:</strong></p>",
            "<p style='color: #ffffff; font-family: Orbitron, sans-serif; font-size: 28px; font-weight: bold;'>", scores[["cellulolytic"]], "</p>",
            "<p style='color: #cccccc; font-family: Orbitron, sans-serif;'>Cellulose degradation enzymes</p>",
            "<div style='margin-top: 10px; font-size: 12px; color: #aaaaaa;'>",
            "GH families + CBM modules + AA oxidases",
            "</div>",
            "</div>",
            
            "<div style='background: rgba(56, 239, 125, 0.1); padding: 20px; border-radius: 10px; border-left: 4px solid #38ef7d;'>",
            "<p><strong style='color: #38ef7d; font-family: Orbitron, monospace; font-size: 16px;'>🌿 HEMI-CELLULOLYTIC POWER:</strong></p>",
            "<p style='color: #ffffff; font-family: Orbitron, sans-serif; font-size: 28px; font-weight: bold;'>", scores[["hemi_cellulolytic"]], "</p>",
            "<p style='color: #cccccc; font-family: Orbitron, sans-serif;'>Hemicellulose processing capacity</p>",
            "<div style='margin-top: 10px; font-size: 12px; color: #aaaaaa;'>",
            "Xylanases + Mannanases + CE esterases",
            "</div>",
            "</div>",
            
            "<div style='background: rgba(255, 107, 53, 0.1); padding: 20px; border-radius: 10px; border-left: 4px solid #ff6b35;'>",
            "<p><strong style='color: #ff6b35; font-family: Orbitron, monospace; font-size: 16px;'>🌳 LIGNOLYTIC POTENTIAL:</strong></p>",
            "<p style='color: #ffffff; font-family: Orbitron, sans-serif; font-size: 28px; font-weight: bold;'>", scores[["lignolytic"]], "</p>",
            "<p style='color: #cccccc; font-family: Orbitron, sans-serif;'>Lignin modification enzymes</p>",
            "<div style='margin-top: 10px; font-size: 12px; color: #aaaaaa;'>",
            "AA oxidases + Peroxidases + Laccases",
            "</div>",
            "</div>",
            
            "</div>",
            
            # Secondary capabilities
            "<div style='display: grid; grid-template-columns: 1fr 1fr 1fr; gap: 20px; margin-top: 20px;'>",
            
            "<div style='background: rgba(156, 89, 209, 0.1); padding: 15px; border-radius: 8px; border-left: 3px solid #9c59d1;'>",
            "<p><strong style='color: #9c59d1; font-family: Orbitron, monospace; font-size: 14px;'>🍎 PECTINOLYTIC:</strong></p>",
            "<p style='color: #ffffff; font-family: Orbitron, sans-serif; font-size: 22px; font-weight: bold;'>", scores[["pectinolytic"]], "</p>",
            "</div>",
            
            "<div style='background: rgba(247, 147, 30, 0.1); padding: 15px; border-radius: 8px; border-left: 3px solid #f7931e;'>",
            "<p><strong style='color: #f7931e; font-family: Orbitron, monospace; font-size: 14px;'>🌽 STARCH PROCESSING:</strong></p>",
            "<p style='color: #ffffff; font-family: Orbitron, sans-serif; font-size: 22px; font-weight: bold;'>", scores[["starch_degrading"]], "</p>",
            "</div>",
            
            "<div style='background: rgba(26, 188, 156, 0.1); padding: 15px; border-radius: 8px; border-left: 3px solid #1abc9c;'>",
            "<p><strong style='color: #1abc9c; font-family: Orbitron, monospace; font-size: 14px;'>🥔 INULIN DEGRADATION:</strong></p>",
            "<p style='color: #ffffff; font-family: Orbitron, sans-serif; font-size: 22px; font-weight: bold;'>", scores[["inulin_degrading"]], "</p>",
            "</div>",
            
            "</div>",
            
            # Debug information (remove in production)
            if(total_score == 0) {
              paste(
                "<div style='margin-top: 20px; padding: 10px; background: rgba(255, 255, 0, 0.1); border-radius: 5px;'>",
                "<p style='color: #ffff00; font-size: 12px;'>Debug: Total score = ", total_score, "</p>",
                "<p style='color: #ffff00; font-size: 12px;'>Available columns: ", paste(names(values$processed_data), collapse = ", "), "</p>",
                "</div>"
              )
            } else "",
            
            "</div>"
          ))
          
        }, error = function(e) {
          HTML(paste(
            "<div class='degradation-card quantum-glow'>",
            "<h4>❌ ERROR IN ASSESSMENT</h4>",
            "<p style='color: #ff4444;'>Error: ", e$message, "</p>",
            "<p style='color: #cccccc;'>Please check your data format and try again.</p>",
            "</div>"
          ))
        })
      }
    }
  })
  
  # Multi-strain battle outputs
  output$multiBattleChart <- renderPlotly({
    if (!is.null(values$processed_data) && !is.null(input$selectedStrains) && length(input$selectedStrains) > 1) {
      
      enzyme_col <- if("Annotations/Genomes" %in% names(values$processed_data)) {
        "Annotations/Genomes"
      } else if("CAZymes" %in% names(values$processed_data)) {
        "CAZymes"
      } else {
        names(values$processed_data)[1]
      }
      
      battle_data <- data.frame(
        Strain = input$selectedStrains,
        Score = numeric(length(input$selectedStrains)),
        stringsAsFactors = FALSE
      )
      
      for(i in 1:nrow(battle_data)) {
        strain <- battle_data$Strain[i]
        if(input$battleMetric == "total") {
          battle_data$Score[i] <- sum(as.numeric(values$processed_data[[strain]]), na.rm = TRUE)
        } else {
          potential <- calculateDegradationPotential(values$processed_data, values$enzyme_families[[input$battleMetric]])
          battle_data$Score[i] <- if(!is.null(potential) && strain %in% names(potential)) potential[[strain]] else 0
        }
      }
      
      battle_data <- battle_data %>% arrange(desc(Score))
      
      p <- ggplot(battle_data, aes(x = reorder(Strain, Score), y = Score, fill = reorder(Strain, Score))) +
        geom_col(alpha = 0.8, color = "white", size = 0.5) +
        coord_flip() +
        labs(title = "Multi-Strain Battle Results",
             x = "Strain", y = "Score") +
        theme_minimal() +
        theme(
          plot.background = element_rect(fill = "transparent", color = NA),
          panel.background = element_rect(fill = "transparent", color = NA),
          text = element_text(color = "white"),
          axis.text = element_text(color = "white"),
          legend.position = "none"
        ) +
        scale_fill_viridis_d(option = "plasma")
      
      ggplotly(p) %>%
        layout(
          plot_bgcolor = 'rgba(0,0,0,0)',
          paper_bgcolor = 'rgba(0,0,0,0)',
          font = list(color = 'white')
        )%>%
        config(
          toImageButtonOptions = list(
            format = 'png',
            filename = 'Multi_Strain_Battle_Results',
            height = 500,
            width = 750,
            scale = 4  
          )
        )
      
    }
  })
  
  # Multi-strain battle comparison table
  output$multiBattleTable <- DT::renderDataTable({
    if (!is.null(values$processed_data) && !is.null(input$selectedStrains) && length(input$selectedStrains) > 1) {
      
      # Identify the enzyme column name
      enzyme_col <- if("Annotations/Genomes" %in% names(values$processed_data)) {
        "Annotations/Genomes"
      } else if("CAZymes" %in% names(values$processed_data)) {
        "CAZymes"
      } else {
        names(values$processed_data)[1]
      }
      
      # Create comparison matrix
      comparison_data <- data.frame(
        Enzyme_Family = values$processed_data[[enzyme_col]],
        stringsAsFactors = FALSE
      )
      
      # Add columns for each selected strain
      for(strain in input$selectedStrains) {
        if(strain %in% names(values$processed_data)) {
          comparison_data[[strain]] <- as.numeric(as.character(values$processed_data[[strain]]))
        }
      }
      
      # Remove rows where all strain values are 0
      strain_cols <- input$selectedStrains[input$selectedStrains %in% names(comparison_data)]
      comparison_data <- comparison_data[rowSums(comparison_data[strain_cols], na.rm = TRUE) > 0, ]
      
      # Add difference columns for comparison
      if(length(strain_cols) >= 2) {
        for(i in 1:(length(strain_cols)-1)) {
          for(j in (i+1):length(strain_cols)) {
            col_name <- paste0(strain_cols[i], "_vs_", strain_cols[j])
            comparison_data[[col_name]] <- comparison_data[[strain_cols[i]]] - comparison_data[[strain_cols[j]]]
          }
        }
      }
      
      DT::datatable(comparison_data,
                    options = list(
                      pageLength = 15,
                      scrollX = TRUE,
                      order = list(list(1, 'desc')),
                      initComplete = JS(
                        "function(settings, json) {",
                        "$(this.api().table().header()).css({'background-color': '#1a1a2e', 'color': '#fff'});",
                        "}"
                      )
                    ),
                    class = 'cell-border stripe hover',
                    rownames = FALSE) %>%
        formatStyle(columns = 1:ncol(comparison_data), 
                    backgroundColor = 'rgba(26, 26, 46, 0.9)', 
                    color = 'white') %>%
        formatStyle(strain_cols,
                    background = styleColorBar(c(0, max(comparison_data[strain_cols], na.rm = TRUE)), '#00d4ff'),
                    backgroundSize = '100% 90%',
                    backgroundRepeat = 'no-repeat',
                    backgroundPosition = 'center')
    } else {
      # Return empty table with message
      DT::datatable(data.frame(Message = "Please select at least 2 strains for comparison"),
                    options = list(dom = 't'),
                    rownames = FALSE)
    }
  })
  
  
  # Strain ranking table
  output$strainRankingTable <- DT::renderDataTable({
    if (!is.null(values$processed_data)) {
      
      enzyme_col <- if("Annotations/Genomes" %in% names(values$processed_data)) {
        "Annotations/Genomes"
      } else if("CAZymes" %in% names(values$processed_data)) {
        "CAZymes" 
      } else {
        names(values$processed_data)[1]
      }
      
      genome_cols <- names(values$processed_data)[!names(values$processed_data) %in% c(enzyme_col, "Total")]
      
      ranking_data <- data.frame(
        Strain = genome_cols,
        TotalEnzymes = numeric(length(genome_cols)),
        Cellulolytic = numeric(length(genome_cols)),
        HemiCellulolytic = numeric(length(genome_cols)),
        Lignolytic = numeric(length(genome_cols)),
        stringsAsFactors = FALSE
      )
      
      # Calculate scores for ranking
      for(i in 1:nrow(ranking_data)) {
        strain <- ranking_data$Strain[i]
        
        # Total enzymes
        ranking_data$TotalEnzymes[i] <- sum(as.numeric(values$processed_data[[strain]]), na.rm = TRUE)
        
        # Degradation potentials
        cell_pot <- calculateDegradationPotential(values$processed_data, values$enzyme_families$cellulolytic)
        hemi_pot <- calculateDegradationPotential(values$processed_data, values$enzyme_families$hemi_cellulolytic) 
        lignin_pot <- calculateDegradationPotential(values$processed_data, values$enzyme_families$lignolytic)
        
        ranking_data$Cellulolytic[i] <- if(!is.null(cell_pot) && strain %in% names(cell_pot)) cell_pot[[strain]] else 0
        ranking_data$HemiCellulolytic[i] <- if(!is.null(hemi_pot) && strain %in% names(hemi_pot)) hemi_pot[[strain]] else 0
        ranking_data$Lignolytic[i] <- if(!is.null(lignin_pot) && strain %in% names(lignin_pot)) lignin_pot[[strain]] else 0
      }
      
      # Calculate overall score and rank
      ranking_data <- ranking_data %>%
        mutate(
          OverallScore = Cellulolytic + HemiCellulolytic + (Lignolytic * 0.5),
          Rank = row_number(desc(OverallScore))
        ) %>%
        arrange(Rank)
      
      DT::datatable(ranking_data,
                    options = list(
                      pageLength = 20,
                      order = list(list(6, 'asc')),
                      initComplete = JS(
                        "function(settings, json) {",
                        "$(this.api().table().header()).css({'background-color': '#1a1a2e', 'color': '#fff'});",
                        "}"
                      )
                    ),
                    class = 'cell-border stripe hover',
                    rownames = FALSE) %>%
        formatStyle(columns = 1:7, 
                    backgroundColor = 'rgba(26, 26, 46, 0.9)', 
                    color = 'white') %>%
        formatStyle('OverallScore',
                    background = styleColorBar(range(ranking_data$OverallScore), '#00d4ff'),
                    backgroundSize = '100% 90%',
                    backgroundRepeat = 'no-repeat',
                    backgroundPosition = 'center')
    }
  })
  
  # Top performers visualization for strain ranking
  output$topPerformersChart <- renderPlotly({
    if (!is.null(values$processed_data)) {
      
      # Identify the enzyme column name
      enzyme_col <- if("Annotations/Genomes" %in% names(values$processed_data)) {
        "Annotations/Genomes"
      } else if("CAZymes" %in% names(values$processed_data)) {
        "CAZymes"
      } else {
        names(values$processed_data)[1]
      }
      
      genome_cols <- names(values$processed_data)[!names(values$processed_data) %in% c(enzyme_col, "Total")]
      
      # Calculate performance metrics
      performance_data <- data.frame(
        Strain = genome_cols,
        TotalEnzymes = numeric(length(genome_cols)),
        Cellulolytic = numeric(length(genome_cols)),
        HemiCellulolytic = numeric(length(genome_cols)),
        Lignolytic = numeric(length(genome_cols)),
        stringsAsFactors = FALSE
      )
      
      # Calculate scores
      for(i in 1:nrow(performance_data)) {
        strain <- performance_data$Strain[i]
        
        # Total enzymes
        performance_data$TotalEnzymes[i] <- sum(as.numeric(as.character(values$processed_data[[strain]])), na.rm = TRUE)
        
        # Degradation potentials (using simplified calculation if enzyme_families not available)
        if(!is.null(values$enzyme_families)) {
          cell_pot <- calculateDegradationPotential(values$processed_data, values$enzyme_families$cellulolytic)
          hemi_pot <- calculateDegradationPotential(values$processed_data, values$enzyme_families$hemi_cellulolytic)
          lignin_pot <- calculateDegradationPotential(values$processed_data, values$enzyme_families$lignolytic)
          
          performance_data$Cellulolytic[i] <- if(!is.null(cell_pot) && strain %in% names(cell_pot)) cell_pot[[strain]] else 0
          performance_data$HemiCellulolytic[i] <- if(!is.null(hemi_pot) && strain %in% names(hemi_pot)) hemi_pot[[strain]] else 0
          performance_data$Lignolytic[i] <- if(!is.null(lignin_pot) && strain %in% names(lignin_pot)) lignin_pot[[strain]] else 0
        } else {
          # Fallback: use total enzymes as proxy
          performance_data$Cellulolytic[i] <- performance_data$TotalEnzymes[i] * 0.3
          performance_data$HemiCellulolytic[i] <- performance_data$TotalEnzymes[i] * 0.3
          performance_data$Lignolytic[i] <- performance_data$TotalEnzymes[i] * 0.2
        }
      }
      
      # Get top 10 performers
      top_performers <- performance_data %>%
        mutate(OverallScore = Cellulolytic + HemiCellulolytic + (Lignolytic * 0.5)) %>%
        arrange(desc(OverallScore)) %>%
        head(10)
      
      # Reshape data for plotting
      plot_data <- top_performers %>%
        select(Strain, Cellulolytic, HemiCellulolytic, Lignolytic) %>%
        pivot_longer(cols = c(Cellulolytic, HemiCellulolytic, Lignolytic),
                     names_to = "Metric", values_to = "Score")
      
      # Create stacked bar chart
      p <- ggplot(plot_data, aes(x = reorder(Strain, Score), y = Score, fill = Metric)) +
        geom_col(position = "stack", alpha = 0.8, color = "white", size = 0.3) +
        coord_flip() +
        labs(title = "Top Strain Performers",
             subtitle = "Degradation Potential by Category",
             x = "Strain", y = "Score") +
        theme_minimal() +
        theme(
          plot.background = element_rect(fill = "transparent", color = NA),
          panel.background = element_rect(fill = "transparent", color = NA),
          text = element_text(color = "white"),
          axis.text = element_text(color = "white"),
          legend.text = element_text(color = "white"),
          legend.title = element_text(color = "white"),
          plot.title = element_text(color = "white", size = 16, face = "bold"),
          plot.subtitle = element_text(color = "white", size = 12)
        ) +
        scale_fill_manual(values = c("Cellulolytic" = "#ff6b35", 
                                     "HemiCellulolytic" = "#00d4ff", 
                                     "Lignolytic" = "#11998e"))
      
      ggplotly(p) %>%
        layout(
          plot_bgcolor = 'rgba(0,0,0,0)',
          paper_bgcolor = 'rgba(0,0,0,0)',
          font = list(color = 'white'),
          legend = list(font = list(color = 'white'))
        )%>%
        config(
          toImageButtonOptions = list(
            format = 'png',
            filename = 'Top_Strain_Performers',
            height = 500,
            width = 750,
            scale = 4  
          )
        )
    } else {
      # Return empty plot
      plot_ly() %>%
        layout(
          title = list(text = "No data available", font = list(color = 'white')),
          plot_bgcolor = 'rgba(0,0,0,0)',
          paper_bgcolor = 'rgba(0,0,0,0)'
        )
    }
  })
  
  # Enhanced function to calculate degradation potential with comprehensive enzyme families
  calculateDegradationPotential <- function(data, enzyme_families) {
    if (is.null(data) || is.null(enzyme_families)) {
      cat("calculateDegradationPotential: NULL input\n")
      return(NULL)
    }
    
    # Handle different column name formats
    enzyme_col <- if("Annotations/Genomes" %in% names(data)) {
      "Annotations/Genomes"
    } else if("CAZymes" %in% names(data)) {
      "CAZymes"
    } else {
      names(data)[1]
    }
    
    # Get genome columns (exclude enzyme column and Total if present)
    genome_cols <- names(data)[!names(data) %in% c(enzyme_col, "Total")]
    
    if(length(genome_cols) == 0) {
      cat("calculateDegradationPotential: No genome columns found\n")
      return(NULL)
    }
    
    # Process enzyme families
    if(is.list(enzyme_families)) {
      if(any(sapply(enzyme_families, is.list))) {
        all_enzyme_patterns <- unique(unlist(enzyme_families, recursive = TRUE))
      } else {
        all_enzyme_patterns <- unique(unlist(enzyme_families))
      }
    } else {
      all_enzyme_patterns <- enzyme_families
    }
    
    all_enzyme_patterns <- all_enzyme_patterns[!is.na(all_enzyme_patterns) & nchar(all_enzyme_patterns) > 0]
    
    if(length(all_enzyme_patterns) == 0) {
      cat("calculateDegradationPotential: No enzyme patterns found\n")
      # Return zeros for all genomes
      result <- as.list(rep(0, length(genome_cols)))
      names(result) <- genome_cols
      return(result)
    }
    
    # Enhanced pattern matching
    matching_rows <- rep(FALSE, nrow(data))
    
    for(pattern in all_enzyme_patterns) {
      # Match exact pattern or pattern with underscore/numbers (e.g., AA1, AA1_2, GH5_1)
      pattern_matches <- grepl(paste0("^", pattern, "($|_|\\d)"), data[[enzyme_col]], perl = TRUE)
      matching_rows <- matching_rows | pattern_matches
      
      if(any(pattern_matches)) {
        cat("Found matches for pattern:", pattern, "- Count:", sum(pattern_matches), "\n")
      }
    }
    
    if(!any(matching_rows)) {
      cat("calculateDegradationPotential: No matches found for any patterns\n")
      # Return zeros for all genomes
      result <- as.list(rep(0, length(genome_cols)))
      names(result) <- genome_cols
      return(result)
    }
    
    # Calculate potential scores for each genome
    relevant_data <- data[matching_rows, , drop = FALSE]
    potential_scores <- list()
    
    for(genome in genome_cols) {
      if(genome %in% names(relevant_data)) {
        genome_values <- as.numeric(as.character(relevant_data[[genome]]))
        genome_values[is.na(genome_values)] <- 0
        potential_scores[[genome]] <- sum(genome_values, na.rm = TRUE)
      } else {
        potential_scores[[genome]] <- 0
      }
    }
    
    cat("calculateDegradationPotential: Calculated scores for", length(potential_scores), "genomes\n")
    return(potential_scores)
  }
  
  #Enhanced industrial potential calculation with debugging
  calculateIndustrialPotential <- function(data, enzyme_families) {
    if (is.null(data) || is.null(enzyme_families)) {
      cat("DEBUG Industrial Potential: NULL data or enzyme families\n")
      return(0)
    }
    
    tryCatch({
      cat("=== Industrial Potential Calculation ===\n")
      
      # Calculate individual potentials
      cell_potential <- calculateDegradationPotential(data, enzyme_families$cellulolytic)
      hemi_potential <- calculateDegradationPotential(data, enzyme_families$hemi_cellulolytic)
      lignin_potential <- calculateDegradationPotential(data, enzyme_families$lignolytic)
      
      if(is.null(cell_potential) || is.null(hemi_potential) || is.null(lignin_potential)) {
        cat("DEBUG Industrial Potential: One or more potentials is NULL\n")
        return(0)
      }
      
      # Get all genome columns
      enzyme_col <- if("Annotations/Genomes" %in% names(data)) {
        "Annotations/Genomes"
      } else if("CAZymes" %in% names(data)) {
        "CAZymes"
      } else {
        names(data)[1]
      }
      
      genome_cols <- names(data)[!names(data) %in% c(enzyme_col, "Total")]
      
      # Calculate totals across all genomes
      cell_total <- sum(sapply(genome_cols, function(g) {
        if(g %in% names(cell_potential)) cell_potential[[g]] else 0
      }), na.rm = TRUE)
      
      hemi_total <- sum(sapply(genome_cols, function(g) {
        if(g %in% names(hemi_potential)) hemi_potential[[g]] else 0
      }), na.rm = TRUE)
      
      lignin_total <- sum(sapply(genome_cols, function(g) {
        if(g %in% names(lignin_potential)) lignin_potential[[g]] else 0
      }), na.rm = TRUE)
      
      cat("Totals - Cell:", cell_total, "Hemi:", hemi_total, "Lignin:", lignin_total, "\n")
      
      # Calculate raw score with weighted components
      raw_score <- cell_total + hemi_total + (lignin_total * 0.5)
      cat("Overall raw score:", raw_score, "\n")
      
      # Normalize to percentage (adjust max_possible based on your data range)
      max_possible <- 600  # Adjust this based on your typical maximum scores
      normalized_score <- min(100, round((raw_score / max_possible) * 100))
      
      cat("Normalized industrial potential:", normalized_score, "%\n")
      return(normalized_score)
      
    }, error = function(e) {
      cat("ERROR in calculateIndustrialPotential:", e$message, "\n")
      return(0)
    })
  }
 
  # Industrial Biorefinery Heatmap
  output$enzymeHeatmap <- renderPlotly({
    if (!is.null(values$processed_data)) {
      enzyme_col <- if("Annotations/Genomes" %in% names(values$processed_data)) {
        "Annotations/Genomes"
      } else if("CAZymes" %in% names(values$processed_data)) {
        "CAZymes"
      } else {
        names(values$processed_data)[1]
      }
      
      # Prepare data for heatmap
      genome_cols <- names(values$processed_data)[!names(values$processed_data) %in% c(enzyme_col, "Total")]
      
      if(length(genome_cols) > 0) {
        # Ensure numeric conversion
        numeric_data <- values$processed_data[genome_cols] %>%
          mutate(across(everything(), \(x) as.numeric(as.character(x))))
        
        # Get top 20 families by total count
        heatmap_data <- values$processed_data %>%
          mutate(Total_Count = rowSums(numeric_data, na.rm = TRUE)) %>%
          slice_max(Total_Count, n = 20) %>%
          select(all_of(c(enzyme_col, genome_cols)))
        
        # Convert genome columns to numeric for heatmap
        heatmap_data[genome_cols] <- lapply(heatmap_data[genome_cols], function(x) as.numeric(as.character(x)))
        
        # Convert to matrix properly
        row_names <- heatmap_data[[enzyme_col]]
        heatmap_matrix <- heatmap_data %>%
          select(-all_of(enzyme_col)) %>%
          as.matrix()
        
        # Set row names
        rownames(heatmap_matrix) <- row_names
        
        # Handle any remaining NAs
        heatmap_matrix[is.na(heatmap_matrix)] <- 0
        
        plot_ly(
          z = ~heatmap_matrix,
          x = ~colnames(heatmap_matrix),
          y = ~rownames(heatmap_matrix),
          type = "heatmap",
          colorscale = "Viridis",
          hovertemplate = "Genome: %{x}<br>Family: %{y}<br>Count: %{z}<extra></extra>",
          colorbar = list(title = "Enzyme Count")
        ) %>%
          layout(
            title = "Industrial Biorefinery Enzyme Heatmap",
            xaxis = list(title = "Genomes", tickangle = 45),
            yaxis = list(title = "Enzyme Families"),
            plot_bgcolor = 'rgba(0,0,0,0)',
            paper_bgcolor = 'rgba(0,0,0,0)',
            font = list(color = 'white')
          )%>%
          config(
            toImageButtonOptions = list(
              format = 'png',
              filename = 'Industrial_Biorefinery_Enzyme_Heatmap',
              height = 550,
              width = 650,
              scale = 4 
            )
          )
      }
    }
  })
  
  # Network visualization for enzyme relationships (optional - if you want this too)
  output$enzymeNetwork <- renderForceNetwork({
    if (!is.null(values$processed_data)) {
      # Create a simple network showing enzyme family relationships
      # This is a simplified example - in practice, you'd want more sophisticated network analysis
      
      enzyme_col <- if("Annotations/Genomes" %in% names(values$processed_data)) {
        "Annotations/Genomes"
      } else if("CAZymes" %in% names(values$processed_data)) {
        "CAZymes"
      } else {
        names(values$processed_data)[1]
      }
      
      # Create nodes and links for top enzyme families
      top_families <- values$processed_data %>%
        slice_head(n = 20) %>%
        pull(!!sym(enzyme_col))
      
      # Simple network structure
      nodes <- data.frame(
        name = top_families,
        group = case_when(
          str_detect(top_families, "GH") ~ 1,
          str_detect(top_families, "AA") ~ 2,
          str_detect(top_families, "CBM") ~ 3,
          str_detect(top_families, "PL") ~ 4,
          str_detect(top_families, "CE") ~ 5,
          TRUE ~ 6
        ),
        size = runif(length(top_families), 10, 30)
      )
      
      # Create some random links for demonstration
      num_links <- min(30, nrow(nodes))
      if(nrow(nodes) > 1) {
        links <- data.frame(
          source = sample(0:(nrow(nodes)-1), num_links, replace = TRUE),
          target = sample(0:(nrow(nodes)-1), num_links, replace = TRUE),
          value = runif(num_links, 1, 10)
        )
        # Ensure no self-loops and valid indices
        links <- links[links$source != links$target, ]
        # Ensure indices are valid
        links <- links[links$source >= 0 & links$source < nrow(nodes) & 
                         links$target >= 0 & links$target < nrow(nodes), ]
        if(nrow(links) == 0 && nrow(nodes) > 1) {
          links <- data.frame(source = 0, target = 1, value = 5)
        }
      } else {
        links <- data.frame(source = numeric(0), target = numeric(0), value = numeric(0))
      }
      
      forceNetwork(
        Links = links,
        Nodes = nodes,
        Source = "source",
        Target = "target",
        Value = "value",
        NodeID = "name",
        Group = "group",
        opacity = 0.8,
        zoom = TRUE,
        fontSize = 12,
        fontFamily = "Orbitron"
      )
    }
  })
  
  output$enzymeDistribution <- renderPlotly({
    if (!is.null(values$processed_data)) {
      enzyme_col <- if("Annotations/Genomes" %in% names(values$processed_data)) {
        "Annotations/Genomes"
      } else if("CAZymes" %in% names(values$processed_data)) {
        "CAZymes"
      } else {
        names(values$processed_data)[1]
      }
      
      # Get genome columns
      genome_cols <- names(values$processed_data)[!names(values$processed_data) %in% c(enzyme_col, "Total")]
      
      if(length(genome_cols) > 0) {
        # Ensure numeric conversion for genome columns
        numeric_data <- values$processed_data[genome_cols] %>%
          mutate(across(everything(), \(x) as.numeric(as.character(x))))
        
        # Calculate row sums for top families
        plot_data <- values$processed_data %>%
          mutate(row_sum = rowSums(numeric_data, na.rm = TRUE)) %>%
          slice_max(row_sum, n = 20) %>%
          select(all_of(c(enzyme_col, genome_cols)))
        
        # Convert genome columns to numeric for the plot data
        plot_data[genome_cols] <- lapply(plot_data[genome_cols], function(x) as.numeric(as.character(x)))
        
        # Pivot for plotting
        plot_data_long <- plot_data %>%
          pivot_longer(cols = all_of(genome_cols), names_to = "Genome", values_to = "Count")
        
        p <- ggplot(plot_data_long, aes(x = reorder(!!sym(enzyme_col), Count), y = Count, fill = Genome)) +
          geom_bar(stat = "identity", alpha = 0.8) +
          coord_flip() +
          labs(title = "Enzyme Distribution Across Genomes",
               x = "Enzyme Family",
               y = "Count") +
          theme_minimal() +
          theme(
            plot.background = element_rect(fill = "transparent", color = NA),
            panel.background = element_rect(fill = "transparent", color = NA),
            text = element_text(color = "white"),
            axis.text = element_text(color = "white"),
            legend.text = element_text(color = "white"),
            legend.title = element_text(color = "white")
          ) +
          scale_fill_viridis_d(option = "plasma")
        
        ggplotly(p, tooltip = c("x", "y", "fill")) %>%
          layout(
            plot_bgcolor = 'rgba(0,0,0,0)',
            paper_bgcolor = 'rgba(0,0,0,0)',
            font = list(color = 'white')
          )%>%
          config(
            toImageButtonOptions = list(
              format = 'png',
              filename = 'Enzyme_Distribution_Across_Genomes',
              height = 500,
              width = 1000,
              scale = 4  
            )
          )
      }
    }
  })
  
 
  # Define enzyme families (add this early in your server function)
  values$enzyme_families <- list(
    cellulolytic = c("GH1", "GH2", "GH3", "GH5", "GH6", "GH7", "GH8", "GH9", "GH10", 
                     "GH11", "GH12", "GH45", "GH48", "GH74", "GH124", "CBM1", "CBM3", "CBM6"),
    hemi_cellulolytic = c("GH10", "GH11", "GH26", "GH29", "GH51", "GH53", "GH62", "GH67", 
                          "GH95", "GH115", "CE1", "CE2", "CE3", "CE4", "CE5", "CE6", "CE15", "CE16"),
    lignolytic = c("AA1", "AA2", "AA3", "AA4", "AA5", "AA6", "AA7", "AA8", "AA9", "AA14", 
                   "CE1", "CE4", "CE10", "GH10", "GH11"),
    pectinolytic = c("GH28", "GH43", "GH78", "GH88", "GH105", "GH106", "PL1", "PL2", "PL3", 
                     "PL4", "PL6", "PL7", "PL8", "PL9", "PL10", "PL11", "CE8", "CE12"),
    starch_degrading = c("GH13", "GH14", "GH15", "GH31", "GH57", "GH77", "GH119", "CBM20", 
                         "CBM21", "CBM25", "CBM26", "CBM34", "CBM41", "CBM45"),
    inulin_degrading = c("GH32", "GH91", "GH134")
  )
  
  # 3. COMPLETELY REWRITTEN: Industrial Biorefinery Heatmap
  output$industrialHeatmap <- renderPlotly({
    cat("=== Starting Industrial Heatmap Generation ===\n")
    
    if (is.null(values$processed_data)) {
      cat("No processed data available for heatmap\n")
      return(NULL)
    }
    
    tryCatch({
      # Identify columns
      enzyme_col <- if("Annotations/Genomes" %in% names(values$processed_data)) {
        "Annotations/Genomes"
      } else if("CAZymes" %in% names(values$processed_data)) {
        "CAZymes"
      } else {
        names(values$processed_data)[1]
      }
      
      genome_cols <- names(values$processed_data)[!names(values$processed_data) %in% c(enzyme_col, "Total")]
      
      cat("Heatmap: Found", length(genome_cols), "genome columns\n")
      
      if(length(genome_cols) == 0) {
        return(plot_ly() %>%
                 add_text(x = 0.5, y = 0.5, text = "No genome data available for heatmap", 
                          textfont = list(color = "white", size = 16)) %>%
                 layout(
                   xaxis = list(showgrid = FALSE, showticklabels = FALSE, zeroline = FALSE),
                   yaxis = list(showgrid = FALSE, showticklabels = FALSE, zeroline = FALSE),
                   plot_bgcolor = 'rgba(0,0,0,0)',
                   paper_bgcolor = 'rgba(0,0,0,0)',
                   font = list(color = 'white')
                 ))
      }
      
      # Define enzyme types and their families
      enzyme_types <- c("Cellulolytic", "Hemi-Cellulolytic", "Lignolytic", "Pectinolytic", "Starch", "Inulin")
      enzyme_families_list <- list(
        values$enzyme_families$cellulolytic,
        values$enzyme_families$hemi_cellulolytic,
        values$enzyme_families$lignolytic,
        values$enzyme_families$pectinolytic,
        values$enzyme_families$starch_degrading,
        values$enzyme_families$inulin_degrading
      )
      
      # Create heatmap matrix
      heatmap_matrix <- matrix(0, nrow = length(genome_cols), ncol = length(enzyme_types))
      rownames(heatmap_matrix) <- genome_cols
      colnames(heatmap_matrix) <- enzyme_types
      
      # Calculate scores for each combination
      for(i in seq_along(enzyme_types)) {
        cat("Calculating", enzyme_types[i], "potential...\n")
        potential <- calculateDegradationPotential(values$processed_data, enzyme_families_list[[i]])
        
        if(!is.null(potential)) {
          for(j in seq_along(genome_cols)) {
            genome <- genome_cols[j]
            if(genome %in% names(potential)) {
              heatmap_matrix[j, i] <- max(0, potential[[genome]])
            }
          }
        }
      }
      
      cat("Heatmap matrix created:\n")
      print(heatmap_matrix)
      
      if(all(heatmap_matrix == 0)) {
        return(plot_ly() %>%
                 add_text(x = 0.5, y = 0.5, text = "No degradation potential detected", 
                          textfont = list(color = "orange", size = 16)) %>%
                 layout(
                   xaxis = list(showgrid = FALSE, showticklabels = FALSE, zeroline = FALSE),
                   yaxis = list(showgrid = FALSE, showticklabels = FALSE, zeroline = FALSE),
                   plot_bgcolor = 'rgba(0,0,0,0)',
                   paper_bgcolor = 'rgba(0,0,0,0)',
                   font = list(color = 'white')
                 ))
      }
      
      # Create the heatmap
      plot_ly(
        z = ~heatmap_matrix,
        x = colnames(heatmap_matrix),
        y = rownames(heatmap_matrix),
        type = "heatmap",
        colorscale = list(
          c(0, "#0d1421"),
          c(0.2, "#1e3a8a"),
          c(0.4, "#3b82f6"),
          c(0.6, "#f59e0b"),
          c(0.8, "#ef4444"),
          c(1, "#dc2626")
        ),
        hovertemplate = "Genome: %{y}<br>Type: %{x}<br>Score: %{z}<extra></extra>",
        colorbar = list(
          title = "Enzyme Count",
          titlefont = list(color = "white"),
          tickfont = list(color = "white")
        )
      ) %>%
        layout(
          title = list(
            text = "Industrial Biorefinery Enzyme Heatmap",
            font = list(color = "white", size = 16)
          ),
          xaxis = list(
            title = "Degradation Capability",
            tickangle = 45,
            color = "white",
            tickfont = list(color = "white")
          ),
          yaxis = list(
            title = "Genome",
            color = "white",
            tickfont = list(color = "white")
          ),
          plot_bgcolor = 'rgba(0,0,0,0)',
          paper_bgcolor = 'rgba(0,0,0,0)',
          font = list(color = 'white')
        )
      
    }, error = function(e) {
      cat("ERROR in industrialHeatmap:", e$message, "\n")
      print(traceback())
      
      plot_ly() %>%
        add_text(x = 0.5, y = 0.5, text = paste("Heatmap Error:", e$message), 
                 textfont = list(color = "red", size = 14)) %>%
        layout(
          xaxis = list(showgrid = FALSE, showticklabels = FALSE, zeroline = FALSE),
          yaxis = list(showgrid = FALSE, showticklabels = FALSE, zeroline = FALSE),
          plot_bgcolor = 'rgba(0,0,0,0)',
          paper_bgcolor = 'rgba(0,0,0,0)',
          font = list(color = 'white')
        )
    })
  })
  
  # 3. FIXED: Degradation Pathway Network
  output$degradationNetwork <- renderPlotly({
    cat("=== Starting Degradation Network Generation ===\n")
    
    if (is.null(values$processed_data)) {
      cat("No processed data available for network\n")
      return(NULL)
    }
    
    tryCatch({
      # Identify columns
      enzyme_col <- if("Annotations/Genomes" %in% names(values$processed_data)) {
        "Annotations/Genomes"
      } else if("CAZymes" %in% names(values$processed_data)) {
        "CAZymes"
      } else {
        names(values$processed_data)[1]
      }
      
      genome_cols <- names(values$processed_data)[!names(values$processed_data) %in% c(enzyme_col, "Total")]
      
      cat("Network: Found", length(genome_cols), "genome columns\n")
      
      if(length(genome_cols) == 0) {
        return(plot_ly() %>%
                 add_text(x = 0.5, y = 0.5, text = "No genome data for network", 
                          textfont = list(color = "white", size = 16)) %>%
                 layout(
                   xaxis = list(showgrid = FALSE, showticklabels = FALSE, zeroline = FALSE),
                   yaxis = list(showgrid = FALSE, showticklabels = FALSE, zeroline = FALSE),
                   plot_bgcolor = 'rgba(0,0,0,0)',
                   paper_bgcolor = 'rgba(0,0,0,0)',
                   font = list(color = 'white')
                 ))
      }
      
      # Define pathways (reduced to avoid empty categories)
      pathways <- c("Cellulose", "Hemicellulose", "Lignin", "Starch")
      pathway_families <- list(
        values$enzyme_families$cellulolytic,
        values$enzyme_families$hemi_cellulolytic,
        values$enzyme_families$lignolytic,
        values$enzyme_families$starch_degrading
      )
      
      # Create network data
      network_data <- data.frame()
      
      for(i in seq_along(pathways)) {
        cat("Calculating pathway strength for", pathways[i], "\n")
        potential <- calculateDegradationPotential(values$processed_data, pathway_families[[i]])
        
        if(!is.null(potential)) {
          for(genome in genome_cols) {
            if(genome %in% names(potential)) {
              strength <- max(0, potential[[genome]])
              
              network_data <- rbind(network_data, data.frame(
                Pathway = pathways[i],
                Genome = genome,
                Strength = strength,
                stringsAsFactors = FALSE
              ))
            }
          }
        }
      }
      
      cat("Network data created with", nrow(network_data), "connections\n")
      
      if(nrow(network_data) == 0 || all(network_data$Strength == 0)) {
        return(plot_ly() %>%
                 add_text(x = 0.5, y = 0.5, text = "No degradation pathways detected", 
                          textfont = list(color = "orange", size = 16)) %>%
                 layout(
                   xaxis = list(showgrid = FALSE, showticklabels = FALSE, zeroline = FALSE),
                   yaxis = list(showgrid = FALSE, showticklabels = FALSE, zeroline = FALSE),
                   plot_bgcolor = 'rgba(0,0,0,0)',
                   paper_bgcolor = 'rgba(0,0,0,0)',
                   font = list(color = 'white')
                 ))
      }
      
      # Filter out zero strength connections for cleaner visualization
      network_data <- network_data[network_data$Strength > 0, ]
      
      if(nrow(network_data) == 0) {
        return(plot_ly() %>%
                 add_text(x = 0.5, y = 0.5, text = "No active degradation pathways", 
                          textfont = list(color = "orange", size = 16)) %>%
                 layout(
                   xaxis = list(showgrid = FALSE, showticklabels = FALSE, zeroline = FALSE),
                   yaxis = list(showgrid = FALSE, showticklabels = FALSE, zeroline = FALSE),
                   plot_bgcolor = 'rgba(0,0,0,0)',
                   paper_bgcolor = 'rgba(0,0,0,0)',
                   font = list(color = 'white')
                 ))
      }
      
      # Create network plot as a scatter plot
      plot_ly(
        data = network_data,
        x = ~Pathway,
        y = ~Genome,
        size = ~Strength,
        color = ~Strength,
        colors = c("#1e40af", "#3b82f6", "#60a5fa", "#f59e0b", "#f97316", "#dc2626"),
        type = "scatter",
        mode = "markers",
        text = ~paste("Pathway:", Pathway, "<br>Genome:", Genome, "<br>Strength:", Strength),
        hovertemplate = "%{text}<extra></extra>",
        marker = list(
          sizemode = "diameter",
          sizeref = if(max(network_data$Strength) > 0) max(network_data$Strength) / 50 else 1,
          opacity = 0.8,
          line = list(color = "white", width = 1)
        )
      ) %>%
        layout(
          title = list(
            text = "Degradation Pathway Network",
            font = list(color = "white", size = 16)
          ),
          xaxis = list(
            title = "Degradation Pathway",
            tickangle = 45,
            color = "white",
            tickfont = list(color = "white")
          ),
          yaxis = list(
            title = "Genome",
            color = "white",
            tickfont = list(color = "white")
          ),
          plot_bgcolor = 'rgba(0,0,0,0)',
          paper_bgcolor = 'rgba(0,0,0,0)',
          font = list(color = 'white'),
          showlegend = FALSE
        )
      
    }, error = function(e) {
      cat("ERROR in degradationNetwork:", e$message, "\n")
      print(traceback())
      
      plot_ly() %>%
        add_text(x = 0.5, y = 0.5, text = paste("Network Error:", e$message), 
                 textfont = list(color = "red", size = 14)) %>%
        layout(
          xaxis = list(showgrid = FALSE, showticklabels = FALSE, zeroline = FALSE),
          yaxis = list(showgrid = FALSE, showticklabels = FALSE, zeroline = FALSE),
          plot_bgcolor = 'rgba(0,0,0,0)',
          paper_bgcolor = 'rgba(0,0,0,0)',
          font = list(color = 'white')
        )%>%
        config(
          toImageButtonOptions = list(
            format = 'png',
            filename = 'Degradation_Pathway_Network',
            height = 1200,
            width = 1200,
            scale = 4  
          )
        )
    })
  })
  
  
  observe({
    if(!is.null(values$enzyme_families)) {
      cat("=== ENZYME FAMILY STRUCTURE DEBUG ===\n")
      cat("Cellulolytic structure:\n")
      print(str(values$enzyme_families$cellulolytic, max.level = 2))
      
      # Test flattening
      flattened <- unlist(values$enzyme_families$cellulolytic, recursive = TRUE)
      cat("Flattened cellulolytic patterns:", length(flattened), "\n")
      cat("Sample patterns:", paste(head(flattened, 10), collapse = ", "), "\n")
    }
  })
  
  # Enhanced cellulolytic analysis with informative dot plots
  output$cellulaseDistribution <- renderPlotly({
    if (!is.null(values$processed_data)) {
      enzyme_col <- if("Annotations/Genomes" %in% names(values$processed_data)) {
        "Annotations/Genomes"
      } else {
        names(values$processed_data)[1]
      }
      
      # Get all cellulolytic enzymes
      all_cellulolytic <- unique(unlist(values$enzyme_families$cellulolytic))
      enzyme_pattern <- paste(all_cellulolytic, collapse = "|")
      
      cellulase_data <- values$processed_data %>%
        filter(str_detect(!!sym(enzyme_col), enzyme_pattern))
      
      if (nrow(cellulase_data) > 0) {
        genome_cols <- names(cellulase_data)[!names(cellulase_data) %in% c(enzyme_col, "Total")]
        
        if(length(genome_cols) > 0) {
          # Create summary statistics for dot plot
          enzyme_summary <- cellulase_data %>%
            select(all_of(c(enzyme_col, genome_cols))) %>%
            pivot_longer(cols = all_of(genome_cols), names_to = "Genome", values_to = "Count") %>%
            filter(Count > 0) %>%
            group_by(!!sym(enzyme_col)) %>%
            summarise(
              Total_Count = sum(Count, na.rm = TRUE),
              Genome_Count = n_distinct(Genome),
              Avg_Count = mean(Count, na.rm = TRUE),
              Max_Count = max(Count, na.rm = TRUE),
              .groups = 'drop'
            ) %>%
            arrange(desc(Total_Count)) %>%
            head(20)  # Top 20 enzymes
          
          # Create dot plot
          p <- ggplot(enzyme_summary, aes(x = Total_Count, y = reorder(!!sym(enzyme_col), Total_Count))) +
            geom_point(aes(size = Genome_Count, color = Avg_Count), alpha = 0.8) +
            scale_size_continuous(name = "# Genomes\nwith Enzyme", range = c(3, 12)) +
            scale_color_viridis_c(name = "Avg Count\nper Genome", option = "plasma") +
            labs(
              title = "Top 20 Cellulolytic Enzymes - Distribution Overview",
              subtitle = "Bubble size = number of genomes, Color = average count per genome",
              x = "Total Enzyme Count Across All Genomes", 
              y = "Enzyme Family"
            ) +
            theme_minimal() +
            theme(
              plot.background = element_rect(fill = "transparent", color = NA),
              panel.background = element_rect(fill = "transparent", color = NA),
              text = element_text(color = "white"),
              axis.text = element_text(color = "white"),
              legend.text = element_text(color = "white"),
              legend.title = element_text(color = "white"),
              panel.grid.major = element_line(color = "grey30"),
              panel.grid.minor = element_line(color = "grey20")
            )
          
          ggplotly(p, tooltip = c("x", "y", "size", "colour")) %>%
            layout(
              plot_bgcolor = 'rgba(0,0,0,0)',
              paper_bgcolor = 'rgba(0,0,0,0)',
              font = list(color = 'white')
            )%>%
            config(
              toImageButtonOptions = list(
                format = 'png',
                filename = 'Top_20_Cellulolytic_Enzymes',
                height = 600,
                width = 800,
                scale = 4  
              )
            )
        }
      }
    }
  })
  
  # New: Genome-level cellulolytic potential dot plot
  output$genomeCellulolytic <- renderPlotly({
    if (!is.null(values$processed_data)) {
      tryCatch({
        enzyme_col <- if("Annotations/Genomes" %in% names(values$processed_data)) {
          "Annotations/Genomes"
        } else {
          names(values$processed_data)[1]
        }
        
        genome_cols <- names(values$processed_data)[!names(values$processed_data) %in% c(enzyme_col, "Total")]
        
        if(length(genome_cols) > 0) {
          # Calculate comprehensive cellulolytic metrics per genome
          genome_metrics <- data.frame(
            Genome = genome_cols,
            Total_Cellulases = 0,
            Enzyme_Diversity = 0,
            Endoglucanases = 0,
            Exoglucanases = 0,
            BetaGlucosidases = 0,
            CBM_Count = 0,
            stringsAsFactors = FALSE
          )
          
          # Define enzyme categories
          endoglucanases <- c("GH5", "GH8", "GH9", "GH12", "GH44", "GH45", "GH74")
          exoglucanases <- c("GH6", "GH7", "GH48")
          beta_glucosidases <- c("GH1", "GH3", "GH116")
          
          # Get cellulolytic enzymes
          all_cellulolytic <- unique(unlist(values$enzyme_families$cellulolytic))
          cellulase_pattern <- paste(all_cellulolytic, collapse = "|")
          cellulase_rows <- str_detect(values$processed_data[[enzyme_col]], cellulase_pattern)
          
          # CBM modules
          if(!is.null(values$enzyme_families) && !is.null(values$enzyme_families$cellulolytic)) {
            if(is.list(values$enzyme_families$cellulolytic) && "CBM" %in% names(values$enzyme_families$cellulolytic)) {
              cbm_modules <- values$enzyme_families$cellulolytic$CBM
            } else {
              all_patterns <- unlist(values$enzyme_families$cellulolytic, recursive = TRUE)
              cbm_modules <- all_patterns[grepl("^CBM", all_patterns)]
            }
          } else {
            cbm_modules <- c("CBM1", "CBM3", "CBM6")
          }
          
          for(i in 1:nrow(genome_metrics)) {
            genome <- genome_metrics$Genome[i]
            if(genome %in% names(values$processed_data)) {
              genome_data <- values$processed_data[[genome]]
              
              # Convert to numeric
              if(is.list(genome_data)) genome_data <- unlist(genome_data)
              genome_data <- suppressWarnings(as.numeric(as.character(genome_data)))
              genome_data[is.na(genome_data)] <- 0
              
              # Calculate metrics
              genome_metrics$Total_Cellulases[i] <- sum(genome_data[cellulase_rows], na.rm = TRUE)
              genome_metrics$Enzyme_Diversity[i] <- sum(genome_data[cellulase_rows] > 0, na.rm = TRUE)
              
              # Specific enzyme counts
              endo_pattern <- paste0("^(", paste(endoglucanases, collapse = "|"), ")($|_|\\d)")
              exo_pattern <- paste0("^(", paste(exoglucanases, collapse = "|"), ")($|_|\\d)")
              beta_pattern <- paste0("^(", paste(beta_glucosidases, collapse = "|"), ")($|_|\\d)")
              cbm_pattern <- paste0("^(", paste(cbm_modules, collapse = "|"), ")($|_|\\d)")
              
              genome_metrics$Endoglucanases[i] <- sum(genome_data[str_detect(values$processed_data[[enzyme_col]], endo_pattern)], na.rm = TRUE)
              genome_metrics$Exoglucanases[i] <- sum(genome_data[str_detect(values$processed_data[[enzyme_col]], exo_pattern)], na.rm = TRUE)
              genome_metrics$BetaGlucosidases[i] <- sum(genome_data[str_detect(values$processed_data[[enzyme_col]], beta_pattern)], na.rm = TRUE)
              genome_metrics$CBM_Count[i] <- sum(genome_data[str_detect(values$processed_data[[enzyme_col]], cbm_pattern)], na.rm = TRUE)
            }
          }
          
          # Filter and rank genomes
          top_genomes <- genome_metrics %>%
            filter(Total_Cellulases > 0) %>%
            mutate(
              Synergy_Score = Endoglucanases + Exoglucanases + BetaGlucosidases + CBM_Count,
              Genome_Short = if_else(nchar(Genome) > 12, paste0(substr(Genome, 1, 12), "..."), Genome)
            ) %>%
            arrange(desc(Total_Cellulases)) %>%
            head(25)  # Top 25 genomes
          
          if(nrow(top_genomes) > 0) {
            p <- ggplot(top_genomes, aes(x = Total_Cellulases, y = reorder(Genome_Short, Total_Cellulases))) +
              geom_point(aes(size = Enzyme_Diversity, color = Synergy_Score), alpha = 0.8) +
              scale_size_continuous(name = "Enzyme\nDiversity", range = c(3, 10)) +
              scale_color_viridis_c(name = "Synergy\nScore", option = "plasma") +
              labs(
                title = "Top 25 Cellulolytic Genomes - Comprehensive Analysis",
                subtitle = "Bubble size = enzyme diversity, Color = synergy score",
                x = "Total Cellulolytic Enzymes", 
                y = "Genome"
              ) +
              theme_minimal() +
              theme(
                plot.background = element_rect(fill = "transparent", color = NA),
                panel.background = element_rect(fill = "transparent", color = NA),
                text = element_text(color = "white"),
                axis.text = element_text(color = "white"),
                legend.text = element_text(color = "white"),
                legend.title = element_text(color = "white"),
                panel.grid.major = element_line(color = "grey30"),
                panel.grid.minor = element_line(color = "grey20")
              )
            
            ggplotly(p, tooltip = c("x", "y", "size", "colour")) %>%
              layout(
                plot_bgcolor = 'rgba(0,0,0,0)',
                paper_bgcolor = 'rgba(0,0,0,0)',
                font = list(color = 'white')
              )%>%
              config(
                toImageButtonOptions = list(
                  format = 'png',
                  filename = 'Total_Cellulolytic_Enzymes',
                  height = 500,
                  width = 700,
                  scale = 4  
                )
              )
          } else {
            plot_ly() %>%
              add_text(x = 0.5, y = 0.5, text = "No cellulolytic genomes found", 
                       textfont = list(color = "orange", size = 16)) %>%
              layout(
                xaxis = list(showgrid = FALSE, showticklabels = FALSE, zeroline = FALSE),
                yaxis = list(showgrid = FALSE, showticklabels = FALSE, zeroline = FALSE),
                plot_bgcolor = 'rgba(0,0,0,0)',
                paper_bgcolor = 'rgba(0,0,0,0)',
                font = list(color = 'white')
              )
          }
        }
      }, error = function(e) {
        cat("ERROR in genomeCellulolytic:", e$message, "\n")
        plot_ly() %>%
          add_text(x = 0.5, y = 0.5, text = paste("Error:", e$message), 
                   textfont = list(color = "red", size = 14)) %>%
          layout(
            xaxis = list(showgrid = FALSE, showticklabels = FALSE, zeroline = FALSE),
            yaxis = list(showgrid = FALSE, showticklabels = FALSE, zeroline = FALSE),
            plot_bgcolor = 'rgba(0,0,0,0)',
            paper_bgcolor = 'rgba(0,0,0,0)',
            font = list(color = 'white')
          )
      })
    }
  })
  
  
  # Enhanced cellulolytic value boxes
  output$cellulaseCount <- renderValueBox({
    if (!is.null(values$processed_data)) {
      tryCatch({
        enzyme_col <- if("Annotations/Genomes" %in% names(values$processed_data)) {
          "Annotations/Genomes"
        } else if("CAZymes" %in% names(values$processed_data)) {
          "CAZymes"
        } else {
          names(values$processed_data)[1]
        }
        
        # Count GH families specifically - FIXED: Proper list access
        if(!is.null(values$enzyme_families) && !is.null(values$enzyme_families$cellulolytic)) {
          # Handle nested structure properly
          if(is.list(values$enzyme_families$cellulolytic) && "GH" %in% names(values$enzyme_families$cellulolytic)) {
            gh_cellulolytic <- values$enzyme_families$cellulolytic$GH
          } else {
            # If it's a flat list, get GH patterns
            all_patterns <- unlist(values$enzyme_families$cellulolytic, recursive = TRUE)
            gh_cellulolytic <- all_patterns[grepl("^GH", all_patterns)]
          }
          
          if(length(gh_cellulolytic) > 0) {
            gh_pattern <- paste0("^(", paste(gh_cellulolytic, collapse = "|"), ")($|_|\\d)")
            
            cellulase_families <- values$processed_data %>%
              filter(str_detect(!!sym(enzyme_col), gh_pattern))
            
            count <- nrow(cellulase_families)
          } else {
            count <- 0
          }
        } else {
          count <- 0
        }
        
        valueBox(
          value = count,
          subtitle = "GH Cellulase Families",
          icon = icon("seedling"),
          color = "green"
        )
      }, error = function(e) {
        cat("ERROR in cellulaseCount:", e$message, "\n")
        valueBox(
          value = "Error",
          subtitle = "GH Cellulase Families",
          icon = icon("seedling"),
          color = "red"
        )
      })
    } else {
      valueBox(
        value = "N/A",
        subtitle = "GH Cellulase Families",
        icon = icon("seedling"),
        color = "light-blue"
      )
    }
  })
  
  
  output$celluloseBMCount <- renderValueBox({
    if (!is.null(values$processed_data)) {
      tryCatch({
        enzyme_col <- if("Annotations/Genomes" %in% names(values$processed_data)) {
          "Annotations/Genomes"
        } else if("CAZymes" %in% names(values$processed_data)) {
          "CAZymes"
        } else {
          names(values$processed_data)[1]
        }
        
        # Count CBM families - FIXED: Proper list access
        if(!is.null(values$enzyme_families) && !is.null(values$enzyme_families$cellulolytic)) {
          # Handle nested structure properly
          if(is.list(values$enzyme_families$cellulolytic) && "CBM" %in% names(values$enzyme_families$cellulolytic)) {
            cbm_cellulolytic <- values$enzyme_families$cellulolytic$CBM
          } else {
            # If it's a flat list, get CBM patterns
            all_patterns <- unlist(values$enzyme_families$cellulolytic, recursive = TRUE)
            cbm_cellulolytic <- all_patterns[grepl("^CBM", all_patterns)]
          }
          
          if(length(cbm_cellulolytic) > 0) {
            cbm_pattern <- paste0("^(", paste(cbm_cellulolytic, collapse = "|"), ")($|_|\\d)")
            
            cbm_families <- values$processed_data %>%
              filter(str_detect(!!sym(enzyme_col), cbm_pattern))
            
            count <- nrow(cbm_families)
          } else {
            count <- 0
          }
        } else {
          count <- 0
        }
        
        valueBox(
          value = count,
          subtitle = "CBM Binding Modules",
          icon = icon("link"),
          color = "blue"
        )
      }, error = function(e) {
        cat("ERROR in celluloseBMCount:", e$message, "\n")
        valueBox(
          value = "Error",
          subtitle = "CBM Binding Modules",
          icon = icon("link"),
          color = "red"
        )
      })
    } else {
      valueBox(
        value = "N/A",
        subtitle = "CBM Binding Modules",
        icon = icon("link"),
        color = "light-blue"
      )
    }
  })
  
  
  output$topCellulaseStrain <- renderValueBox({
    if (!is.null(values$processed_data)) {
      tryCatch({
        cellulase_potential <- calculateDegradationPotential(values$processed_data, values$enzyme_families$cellulolytic)
        
        if (!is.null(cellulase_potential) && length(cellulase_potential) > 0) {
          # FIXED: Proper handling of list results
          if(is.list(cellulase_potential)) {
            potential_values <- unlist(cellulase_potential)
            if(length(potential_values) > 0 && max(potential_values) > 0) {
              top_strain <- names(potential_values)[which.max(potential_values)]
              display_name <- if(nchar(top_strain) > 8) paste0(substr(top_strain, 1, 8), "...") else top_strain
            } else {
              display_name <- "None"
            }
          } else {
            display_name <- "Error"
          }
          
          valueBox(
            value = display_name,
            subtitle = "Top Cellulase Producer",
            icon = icon("crown"),
            color = "purple"
          )
        } else {
          valueBox(
            value = "N/A",
            subtitle = "Top Cellulase Producer",
            icon = icon("crown"),
            color = "light-blue"
          )
        }
      }, error = function(e) {
        cat("ERROR in topCellulaseStrain:", e$message, "\n")
        valueBox(
          value = "Error",
          subtitle = "Top Cellulase Producer",
          icon = icon("crown"),
          color = "red"
        )
      })
    } else {
      valueBox(
        value = "N/A",
        subtitle = "Top Cellulase Producer",
        icon = icon("crown"),
        color = "light-blue"
      )
    }
  })
  
  output$cellulaseEfficiency <- renderValueBox({
    if (!is.null(values$processed_data)) {
      tryCatch({
        # Calculate efficiency based on enzyme diversity and count
        cellulase_potential <- calculateDegradationPotential(values$processed_data, values$enzyme_families$cellulolytic)
        
        if (!is.null(cellulase_potential) && length(cellulase_potential) > 0) {
          # FIXED: Proper handling of list results
          if(is.list(cellulase_potential)) {
            potential_values <- unlist(cellulase_potential)
            if(length(potential_values) > 0) {
              max_score <- max(potential_values, na.rm = TRUE)
              avg_score <- mean(potential_values, na.rm = TRUE)
            } else {
              max_score <- 0
              avg_score <- 0
            }
          } else {
            max_score <- 0
            avg_score <- 0
          }
          
          efficiency <- case_when(
            max_score >= 100 ~ "ELITE",
            max_score >= 50 ~ "HIGH", 
            max_score >= 20 ~ "MEDIUM",
            max_score > 0 ~ "LOW",
            TRUE ~ "NONE"
          )
          
          color <- case_when(
            efficiency == "ELITE" ~ "purple",
            efficiency == "HIGH" ~ "green",
            efficiency == "MEDIUM" ~ "yellow",
            efficiency == "LOW" ~ "orange",
            TRUE ~ "red"
          )
          
          valueBox(
            value = efficiency,
            subtitle = "Max Efficiency Rating",
            icon = icon("star"),
            color = color
          )
        } else {
          valueBox(
            value = "N/A",
            subtitle = "Efficiency Rating",
            icon = icon("star"),
            color = "light-blue"
          )
        }
      }, error = function(e) {
        cat("ERROR in cellulaseEfficiency:", e$message, "\n")
        valueBox(
          value = "Error",
          subtitle = "Efficiency Rating",
          icon = icon("star"),
          color = "red"
        )
      })
    } else {
      valueBox(
        value = "N/A",
        subtitle = "Efficiency Rating",
        icon = icon("star"),
        color = "light-blue"
      )
    }
  })
  
  output$endoglucanaseCount <- renderValueBox({
    if (!is.null(values$processed_data)) {
      tryCatch({
        enzyme_col <- if("Annotations/Genomes" %in% names(values$processed_data)) {
          "Annotations/Genomes"
        } else if("CAZymes" %in% names(values$processed_data)) {
          "CAZymes"
        } else {
          names(values$processed_data)[1]
        }
        
        # Endoglucanases: GH5, GH8, GH9, GH12, GH44, GH45, GH74
        endoglucanases <- c("GH5", "GH8", "GH9", "GH12", "GH44", "GH45", "GH74")
        endo_pattern <- paste0("^(", paste(endoglucanases, collapse = "|"), ")($|_|\\d)")
        
        endo_families <- values$processed_data %>%
          filter(str_detect(!!sym(enzyme_col), endo_pattern))
        
        count <- nrow(endo_families)
        
        valueBox(
          value = count,
          subtitle = "Endoglucanase Families",
          icon = icon("cut"),
          color = "red"
        )
      }, error = function(e) {
        cat("ERROR in endoglucanaseCount:", e$message, "\n")
        valueBox(
          value = "Error",
          subtitle = "Endoglucanase Families",
          icon = icon("cut"),
          color = "red"
        )
      })
    } else {
      valueBox(
        value = "N/A",
        subtitle = "Endoglucanase Families",
        icon = icon("cut"),
        color = "light-blue"
      )
    }
  })
  
  output$exoglucanaseCount <- renderValueBox({
    if (!is.null(values$processed_data)) {
      tryCatch({
        enzyme_col <- if("Annotations/Genomes" %in% names(values$processed_data)) {
          "Annotations/Genomes"
        } else if("CAZymes" %in% names(values$processed_data)) {
          "CAZymes"
        } else {
          names(values$processed_data)[1]
        }
        
        # Exoglucanases: GH6, GH7, GH48
        exoglucanases <- c("GH6", "GH7", "GH48")
        exo_pattern <- paste0("^(", paste(exoglucanases, collapse = "|"), ")($|_|\\d)")
        
        exo_families <- values$processed_data %>%
          filter(str_detect(!!sym(enzyme_col), exo_pattern))
        
        count <- nrow(exo_families)
        
        valueBox(
          value = count,
          subtitle = "Exoglucanase Families",
          icon = icon("scissors"),
          color = "teal"
        )
      }, error = function(e) {
        cat("ERROR in exoglucanaseCount:", e$message, "\n")
        valueBox(
          value = "Error",
          subtitle = "Exoglucanase Families",
          icon = icon("scissors"),
          color = "red"
        )
      })
    } else {
      valueBox(
        value = "N/A",
        subtitle = "Exoglucanase Families",
        icon = icon("scissors"),
        color = "light-blue"
      )
    }
  })
  
  # FIXED: Cellulase synergy analysis
  output$cellulaseSynergy <- renderPlotly({
    if (!is.null(values$processed_data)) {
      tryCatch({
        enzyme_col <- if("Annotations/Genomes" %in% names(values$processed_data)) {
          "Annotations/Genomes"
        } else if("CAZymes" %in% names(values$processed_data)) {
          "CAZymes"
        } else {
          names(values$processed_data)[1]
        }
        
        genome_cols <- names(values$processed_data)[!names(values$processed_data) %in% c(enzyme_col, "Total")]
        
        if(length(genome_cols) > 0) {
          # Calculate synergy scores based on presence of complementary enzymes
          synergy_data <- data.frame(
            Genome = genome_cols,
            Endoglucanase = 0,
            Exoglucanase = 0,
            BetaGlucosidase = 0,
            CBM = 0,
            stringsAsFactors = FALSE
          )
          
          # Define enzyme groups with proper pattern matching
          endoglucanases <- c("GH5", "GH8", "GH9", "GH12", "GH44", "GH45", "GH74")
          exoglucanases <- c("GH6", "GH7", "GH48")
          beta_glucosidases <- c("GH1", "GH3", "GH116")
          
          # FIXED: Get CBM modules properly
          if(!is.null(values$enzyme_families) && !is.null(values$enzyme_families$cellulolytic)) {
            if(is.list(values$enzyme_families$cellulolytic) && "CBM" %in% names(values$enzyme_families$cellulolytic)) {
              cbm_modules <- values$enzyme_families$cellulolytic$CBM
            } else {
              all_patterns <- unlist(values$enzyme_families$cellulolytic, recursive = TRUE)
              cbm_modules <- all_patterns[grepl("^CBM", all_patterns)]
            }
          } else {
            cbm_modules <- c("CBM1", "CBM3", "CBM6")  # Default CBM modules
          }
          
          for(i in 1:nrow(synergy_data)) {
            genome <- synergy_data$Genome[i]
            if(genome %in% names(values$processed_data)) {
              # Calculate scores for each enzyme group with proper pattern matching
              endo_pattern <- paste0("^(", paste(endoglucanases, collapse = "|"), ")($|_|\\d)")
              exo_pattern <- paste0("^(", paste(exoglucanases, collapse = "|"), ")($|_|\\d)")
              beta_pattern <- paste0("^(", paste(beta_glucosidases, collapse = "|"), ")($|_|\\d)")
              cbm_pattern <- paste0("^(", paste(cbm_modules, collapse = "|"), ")($|_|\\d)")
              
              # FIXED: Proper handling of data conversion
              genome_data <- values$processed_data[[genome]]
              
              # Convert to numeric, handling any list or character issues
              if(is.list(genome_data)) {
                genome_data <- unlist(genome_data)
              }
              genome_data <- suppressWarnings(as.numeric(as.character(genome_data)))
              genome_data[is.na(genome_data)] <- 0
              
              # Calculate enzyme counts
              endo_matches <- str_detect(values$processed_data[[enzyme_col]], endo_pattern)
              exo_matches <- str_detect(values$processed_data[[enzyme_col]], exo_pattern)
              beta_matches <- str_detect(values$processed_data[[enzyme_col]], beta_pattern)
              cbm_matches <- str_detect(values$processed_data[[enzyme_col]], cbm_pattern)
              
              synergy_data$Endoglucanase[i] <- sum(genome_data[endo_matches], na.rm = TRUE)
              synergy_data$Exoglucanase[i] <- sum(genome_data[exo_matches], na.rm = TRUE)
              synergy_data$BetaGlucosidase[i] <- sum(genome_data[beta_matches], na.rm = TRUE)
              synergy_data$CBM[i] <- sum(genome_data[cbm_matches], na.rm = TRUE)
            }
          }
          
          # Filter out genomes with no data and take top performers
          synergy_data <- synergy_data %>%
            filter(Endoglucanase + Exoglucanase + BetaGlucosidase + CBM > 0) %>%
            mutate(TotalScore = Endoglucanase + Exoglucanase + BetaGlucosidase + CBM) %>%
            arrange(desc(TotalScore)) %>%
            head(10) %>%
            select(-TotalScore)
          
          if(nrow(synergy_data) > 0) {
            # Create synergy chart data
            plot_data <- synergy_data %>%
              pivot_longer(cols = c("Endoglucanase", "Exoglucanase", "BetaGlucosidase", "CBM"), 
                           names_to = "EnzymeType", values_to = "Count")
            
            p <- ggplot(plot_data, aes(x = EnzymeType, y = Count, fill = Genome)) +
              geom_bar(stat = "identity", position = "dodge", alpha = 0.8) +
              labs(title = "Cellulase Synergy Matrix - Top Performers",
                   x = "Enzyme Type", y = "Count") +
              theme_minimal() +
              theme(
                plot.background = element_rect(fill = "transparent", color = NA),
                panel.background = element_rect(fill = "transparent", color = NA),
                text = element_text(color = "white"),
                axis.text = element_text(color = "white"),
                axis.text.x = element_text(angle = 45, hjust = 1),
                legend.text = element_text(color = "white"),
                legend.title = element_text(color = "white")
              ) +
              scale_fill_viridis_d(option = "plasma")
            
            ggplotly(p) %>%
              layout(
                plot_bgcolor = 'rgba(0,0,0,0)',
                paper_bgcolor = 'rgba(0,0,0,0)',
                font = list(color = 'white')
              )%>%
              config(
                toImageButtonOptions = list(
                  format = 'png',
                  filename = 'Cellulase_Synergy_Matrix',
                  height = 400,
                  width = 600,
                  scale = 4  
                )
              )
          } else {
            # No data available
            plot_ly() %>%
              add_text(x = 0.5, y = 0.5, text = "No cellulase synergy data available", 
                       textfont = list(color = "orange", size = 16)) %>%
              layout(
                xaxis = list(showgrid = FALSE, showticklabels = FALSE, zeroline = FALSE),
                yaxis = list(showgrid = FALSE, showticklabels = FALSE, zeroline = FALSE),
                plot_bgcolor = 'rgba(0,0,0,0)',
                paper_bgcolor = 'rgba(0,0,0,0)',
                font = list(color = 'white')
              )
          }
        } else {
          # No genome data
          plot_ly() %>%
            add_text(x = 0.5, y = 0.5, text = "No genome data available", 
                     textfont = list(color = "red", size = 16)) %>%
            layout(
              xaxis = list(showgrid = FALSE, showticklabels = FALSE, zeroline = FALSE),
              yaxis = list(showgrid = FALSE, showticklabels = FALSE, zeroline = FALSE),
              plot_bgcolor = 'rgba(0,0,0,0)',
              paper_bgcolor = 'rgba(0,0,0,0)',
              font = list(color = 'white')
            )
        }
      }, error = function(e) {
        cat("ERROR in cellulaseSynergy:", e$message, "\n")
        plot_ly() %>%
          add_text(x = 0.5, y = 0.5, text = paste("Synergy Error:", e$message), 
                   textfont = list(color = "red", size = 14)) %>%
          layout(
            xaxis = list(showgrid = FALSE, showticklabels = FALSE, zeroline = FALSE),
            yaxis = list(showgrid = FALSE, showticklabels = FALSE, zeroline = FALSE),
            plot_bgcolor = 'rgba(0,0,0,0)',
            paper_bgcolor = 'rgba(0,0,0,0)',
            font = list(color = 'white')
          )
      })
    } else {
      plot_ly() %>%
        add_text(x = 0.5, y = 0.5, text = "Please upload data first", 
                 textfont = list(color = "white", size = 16)) %>%
        layout(
          xaxis = list(showgrid = FALSE, showticklabels = FALSE, zeroline = FALSE),
          yaxis = list(showgrid = FALSE, showticklabels = FALSE, zeroline = FALSE),
          plot_bgcolor = 'rgba(0,0,0,0)',
          paper_bgcolor = 'rgba(0,0,0,0)',
          font = list(color = 'white')
        )
    }
  })
  
  # New: Enzyme category distribution dot plot
  output$enzymeCategoryDist <- renderPlotly({
    if (!is.null(values$processed_data)) {
      tryCatch({
        enzyme_col <- if("Annotations/Genomes" %in% names(values$processed_data)) {
          "Annotations/Genomes"
        } else {
          names(values$processed_data)[1]
        }
        
        genome_cols <- names(values$processed_data)[!names(values$processed_data) %in% c(enzyme_col, "Total")]
        
        if(length(genome_cols) > 0) {
          # Define enzyme categories
          enzyme_categories <- list(
            "Endoglucanases" = c("GH5", "GH8", "GH9", "GH12", "GH44", "GH45", "GH74"),
            "Exoglucanases" = c("GH6", "GH7", "GH48"),
            "Beta-Glucosidases" = c("GH1", "GH3", "GH116"),
            "Hemicellulases" = c("GH10", "GH11", "GH26", "GH43"),
            "Pectinases" = c("GH28", "PL1", "PL3", "PL4"),
            "CBM Modules" = c("CBM1", "CBM3", "CBM6", "CBM10")
          )
          
          category_summary <- data.frame(
            Category = names(enzyme_categories),
            Total_Enzymes = 0,
            Avg_per_Genome = 0,
            Max_per_Genome = 0,
            Genomes_with_Category = 0,
            stringsAsFactors = FALSE
          )
          
          for(i in 1:nrow(category_summary)) {
            category <- category_summary$Category[i]
            enzymes <- enzyme_categories[[category]]
            
            if(length(enzymes) > 0) {
              pattern <- paste0("^(", paste(enzymes, collapse = "|"), ")($|_|\\d)")
              matching_rows <- str_detect(values$processed_data[[enzyme_col]], pattern)
              
              if(sum(matching_rows) > 0) {
                category_data <- values$processed_data[matching_rows, genome_cols]
                
                # Convert to numeric
                numeric_data <- as.matrix(category_data)
                numeric_data <- suppressWarnings(apply(numeric_data, 2, as.numeric))
                numeric_data[is.na(numeric_data)] <- 0
                
                genome_totals <- colSums(numeric_data, na.rm = TRUE)
                
                category_summary$Total_Enzymes[i] <- sum(numeric_data, na.rm = TRUE)
                category_summary$Avg_per_Genome[i] <- mean(genome_totals, na.rm = TRUE)
                category_summary$Max_per_Genome[i] <- max(genome_totals, na.rm = TRUE)
                category_summary$Genomes_with_Category[i] <- sum(genome_totals > 0, na.rm = TRUE)
              }
            }
          }
          
          category_summary <- category_summary %>%
            filter(Total_Enzymes > 0) %>%
            arrange(desc(Total_Enzymes))
          
          if(nrow(category_summary) > 0) {
            p <- ggplot(category_summary, aes(x = Avg_per_Genome, y = reorder(Category, Total_Enzymes))) +
              geom_point(aes(size = Total_Enzymes, color = Genomes_with_Category), alpha = 0.8) +
              scale_size_continuous(name = "Total\nEnzymes", range = c(4, 15)) +
              scale_color_viridis_c(name = "Genomes with\nCategory", option = "plasma") +
              labs(
                title = "Enzyme Category Distribution Analysis",
                subtitle = "Bubble size = total enzymes, Color = genome prevalence",
                x = "Average Enzymes per Genome", 
                y = "Enzyme Category"
              ) +
              theme_minimal() +
              theme(
                plot.background = element_rect(fill = "transparent", color = NA),
                panel.background = element_rect(fill = "transparent", color = NA),
                text = element_text(color = "white"),
                axis.text = element_text(color = "white"),
                legend.text = element_text(color = "white"),
                legend.title = element_text(color = "white"),
                panel.grid.major = element_line(color = "grey30"),
                panel.grid.minor = element_line(color = "grey20")
              )
            
            ggplotly(p, tooltip = c("x", "y", "size", "colour")) %>%
              layout(
                plot_bgcolor = 'rgba(0,0,0,0)',
                paper_bgcolor = 'rgba(0,0,0,0)',
                font = list(color = 'white')
              )%>%
              config(
                toImageButtonOptions = list(
                  format = 'png',
                  filename = 'Enzyme_Category_Distribution_Analysis',
                  height = 500,
                  width = 700,
                  scale = 4  
                )
              )
          } else {
            plot_ly() %>%
              add_text(x = 0.5, y = 0.5, text = "No enzyme categories found", 
                       textfont = list(color = "orange", size = 16)) %>%
              layout(
                xaxis = list(showgrid = FALSE, showticklabels = FALSE, zeroline = FALSE),
                yaxis = list(showgrid = FALSE, showticklabels = FALSE, zeroline = FALSE),
                plot_bgcolor = 'rgba(0,0,0,0)',
                paper_bgcolor = 'rgba(0,0,0,0)',
                font = list(color = 'white')
              )
          }
        }
      }, error = function(e) {
        cat("ERROR in enzymeCategoryDist:", e$message, "\n")
        plot_ly() %>%
          add_text(x = 0.5, y = 0.5, text = paste("Category Error:", e$message), 
                   textfont = list(color = "red", size = 14)) %>%
          layout(
            xaxis = list(showgrid = FALSE, showticklabels = FALSE, zeroline = FALSE),
            yaxis = list(showgrid = FALSE, showticklabels = FALSE, zeroline = FALSE),
            plot_bgcolor = 'rgba(0,0,0,0)',
            paper_bgcolor = 'rgba(0,0,0,0)',
            font = list(color = 'white')
          )
      })
    }
  })
  
  
  # Hemi-cellulolytic analysis outputs
  output$hemiCellulaseDistribution <- renderPlotly({
    if (!is.null(values$processed_data)) {
      enzyme_col <- if("Annotations/Genomes" %in% names(values$processed_data)) {
        "Annotations/Genomes"
      } else {
        names(values$processed_data)[1]
      }
      
      # Get all hemi-cellulolytic enzymes
      all_hemi <- unique(unlist(values$enzyme_families$hemi_cellulolytic))
      enzyme_pattern <- paste(all_hemi, collapse = "|")
      
      hemi_data <- values$processed_data %>%
        filter(str_detect(!!sym(enzyme_col), enzyme_pattern))
      
      if (nrow(hemi_data) > 0) {
        genome_cols <- names(hemi_data)[!names(hemi_data) %in% c(enzyme_col, "Total")]
        
        if(length(genome_cols) > 0) {
          # Calculate total counts and get top families
          hemi_data <- hemi_data %>%
            mutate(Total_Count = rowSums(select(., all_of(genome_cols)), na.rm = TRUE)) %>%
            slice_max(Total_Count, n = 20)
          
          plot_data <- hemi_data %>%
            select(all_of(c(enzyme_col, genome_cols))) %>%
            pivot_longer(cols = all_of(genome_cols), names_to = "Genome", values_to = "Count") %>%
            filter(Count > 0)
          
          p <- ggplot(plot_data, aes(x = reorder(!!sym(enzyme_col), Count), y = Count, fill = Genome)) +
            geom_bar(stat = "identity", position = "stack", alpha = 0.8) +
            coord_flip() +
            labs(title = "Hemi-Cellulolytic Enzyme Arsenal Distribution",
                 x = "Enzyme Family", y = "Count") +
            theme_minimal() +
            theme(
              plot.background = element_rect(fill = "transparent", color = NA),
              panel.background = element_rect(fill = "transparent", color = NA),
              text = element_text(color = "white"),
              axis.text = element_text(color = "white"),
              legend.text = element_text(color = "white"),
              legend.title = element_text(color = "white")
            ) +
            scale_fill_viridis_d(option = "viridis")
          
          ggplotly(p) %>%
            layout(
              plot_bgcolor = 'rgba(0,0,0,0)',
              paper_bgcolor = 'rgba(0,0,0,0)',
              font = list(color = 'white')
            )%>%
            config(
              toImageButtonOptions = list(
                format = 'png',
                filename = 'Hemi_Cellulolytic_Enzyme_Arsenal_Distribution',
                height = 300,
                width = 700,
                scale = 4  
              )
            )
        }
      }
    }
  })
  
  # Hemi-cellulolytic value boxes
  output$hemiCellulaseCount <- renderValueBox({
    if (!is.null(values$processed_data)) {
      enzyme_col <- if("Annotations/Genomes" %in% names(values$processed_data)) {
        "Annotations/Genomes"
      } else {
        names(values$processed_data)[1]
      }
      
      all_hemi <- unique(unlist(values$enzyme_families$hemi_cellulolytic))
      hemi_pattern <- paste(all_hemi, collapse = "|")
      
      hemi_families <- values$processed_data %>%
        filter(str_detect(!!sym(enzyme_col), hemi_pattern))
      
      count <- nrow(hemi_families)
      
      valueBox(
        value = count,
        subtitle = "Hemi-Cellulase Families",
        icon = icon("leaf"),
        color = "green"
      )
    }
  })
  
  output$xylanaseCount <- renderValueBox({
    if (!is.null(values$processed_data)) {
      enzyme_col <- if("Annotations/Genomes" %in% names(values$processed_data)) {
        "Annotations/Genomes"
      } else {
        names(values$processed_data)[1]
      }
      
      xylanases <- c("GH10", "GH11", "GH43")
      xylanase_pattern <- paste(xylanases, collapse = "|")
      
      xylanase_families <- values$processed_data %>%
        filter(str_detect(!!sym(enzyme_col), xylanase_pattern))
      
      count <- nrow(xylanase_families)
      
      valueBox(
        value = count,
        subtitle = "Xylanase Families",
        icon = icon("tree"),
        color = "blue"
      )
    }
  })
  
  output$mannanaseCount <- renderValueBox({
    if (!is.null(values$processed_data)) {
      enzyme_col <- if("Annotations/Genomes" %in% names(values$processed_data)) {
        "Annotations/Genomes"
      } else {
        names(values$processed_data)[1]
      }
      
      mannanases <- c("GH5", "GH26", "GH113")
      mannanase_pattern <- paste(mannanases, collapse = "|")
      
      mannanase_families <- values$processed_data %>%
        filter(str_detect(!!sym(enzyme_col), mannanase_pattern))
      
      count <- nrow(mannanase_families)
      
      valueBox(
        value = count,
        subtitle = "Mannanase Families",
        icon = icon("seedling"),
        color = "yellow"
      )
    }
  })
  
  output$topHemiStrain <- renderValueBox({
    if (!is.null(values$processed_data)) {
      hemi_potential <- calculateDegradationPotential(values$processed_data, values$enzyme_families$hemi_cellulolytic)
      
      if (!is.null(hemi_potential)) {
        top_strain <- names(hemi_potential)[which.max(hemi_potential)]
        display_name <- if(nchar(top_strain) > 8) paste0(substr(top_strain, 1, 8), "...") else top_strain
        
        valueBox(
          value = display_name,
          subtitle = "Top Hemi Producer",
          icon = icon("crown"),
          color = "purple"
        )
      } else {
        valueBox(
          value = "N/A",
          subtitle = "Top Hemi Producer",
          icon = icon("crown"),
          color = "purple"
        )
      }
    }
  })
  
  output$hemiEfficiency <- renderValueBox({
    if (!is.null(values$processed_data)) {
      hemi_potential <- calculateDegradationPotential(values$processed_data, values$enzyme_families$hemi_cellulolytic)
      
      if (!is.null(hemi_potential)) {
        max_score <- max(unlist(hemi_potential))
        
        efficiency <- case_when(
          max_score >= 150 ~ "ELITE",
          max_score >= 75 ~ "HIGH",
          max_score >= 30 ~ "MEDIUM",
          TRUE ~ "LOW"
        )
        
        valueBox(
          value = efficiency,
          subtitle = "Efficiency Rating",
          icon = icon("star"),
          color = "red"
        )
      } else {
        valueBox(
          value = "N/A",
          subtitle = "Efficiency Rating",
          icon = icon("star"),
          color = "red"
        )
      }
    }
  })
  
  output$hemiDiversity <- renderValueBox({
    if (!is.null(values$processed_data)) {
      enzyme_col <- if("Annotations/Genomes" %in% names(values$processed_data)) {
        "Annotations/Genomes"
      } else {
        names(values$processed_data)[1]
      }
      
      # Count unique enzyme types in hemi-cellulolytic category
      all_hemi <- unique(unlist(values$enzyme_families$hemi_cellulolytic))
      hemi_pattern <- paste(all_hemi, collapse = "|")
      
      diversity_score <- values$processed_data %>%
        filter(str_detect(!!sym(enzyme_col), hemi_pattern)) %>%
        nrow()
      
      valueBox(
        value = diversity_score,
        subtitle = "Enzyme Diversity",
        icon = icon("diversity"),
        color = "teal"
      )
    }
  })
  
  # ============================================================================
  # LIGNOLYTIC POTENTIAL SERVER FUNCTIONS
  # ============================================================================
  
  # Lignolytic value boxes
  output$ligninaseCount <- renderValueBox({
    if (!is.null(values$processed_data)) {
      enzyme_col <- if("Annotations/Genomes" %in% names(values$processed_data)) {
        "Annotations/Genomes"
      } else {
        names(values$processed_data)[1]
      }
      
      lignin_enzymes <- unique(unlist(values$enzyme_families$lignolytic))
      lignin_pattern <- paste(lignin_enzymes, collapse = "|")
      
      lignin_families <- values$processed_data %>%
        filter(str_detect(!!sym(enzyme_col), lignin_pattern))
      
      count <- nrow(lignin_families)
      
      valueBox(
        value = count,
        subtitle = "Lignolytic Families",
        icon = icon("tree"),
        color = "purple"
      )
    }
  })
  
  output$peroxidaseCount <- renderValueBox({
    if (!is.null(values$processed_data)) {
      enzyme_col <- if("Annotations/Genomes" %in% names(values$processed_data)) {
        "Annotations/Genomes"
      } else {
        names(values$processed_data)[1]
      }
      
      peroxidases <- c("AA2", "AA3")
      peroxidase_pattern <- paste(peroxidases, collapse = "|")
      
      peroxidase_families <- values$processed_data %>%
        filter(str_detect(!!sym(enzyme_col), peroxidase_pattern))
      
      count <- nrow(peroxidase_families)
      
      valueBox(
        value = count,
        subtitle = "Peroxidase Families",
        icon = icon("burn"),
        color = "red"
      )
    }
  })
  
  output$laccaseCount <- renderValueBox({
    if (!is.null(values$processed_data)) {
      enzyme_col <- if("Annotations/Genomes" %in% names(values$processed_data)) {
        "Annotations/Genomes"
      } else {
        names(values$processed_data)[1]
      }
      
      laccases <- c("AA1")
      laccase_pattern <- paste(laccases, collapse = "|")
      
      laccase_families <- values$processed_data %>%
        filter(str_detect(!!sym(enzyme_col), laccase_pattern))
      
      count <- nrow(laccase_families)
      
      valueBox(
        value = count,
        subtitle = "Laccase Families",
        icon = icon("atom"),
        color = "blue"
      )
    }
  })
  
  output$topLigninStrain <- renderValueBox({
    if (!is.null(values$processed_data)) {
      lignin_potential <- calculateDegradationPotential(values$processed_data, values$enzyme_families$lignolytic)
      
      if (!is.null(lignin_potential) && length(lignin_potential) > 0) {
        top_strain <- names(lignin_potential)[which.max(lignin_potential)]
        display_name <- if(nchar(top_strain) > 8) paste0(substr(top_strain, 1, 8), "...") else top_strain
        
        valueBox(
          value = display_name,
          subtitle = "Top Lignin Producer",
          icon = icon("crown"),
          color = "yellow"
        )
      } else {
        valueBox(
          value = "N/A",
          subtitle = "Top Lignin Producer",
          icon = icon("crown"),
          color = "yellow"
        )
      }
    }
  })
  
  output$ligninEfficiency <- renderValueBox({
    if (!is.null(values$processed_data)) {
      lignin_potential <- calculateDegradationPotential(values$processed_data, values$enzyme_families$lignolytic)
      
      if (!is.null(lignin_potential) && length(lignin_potential) > 0) {
        max_score <- max(unlist(lignin_potential))
        
        efficiency <- case_when(
          max_score >= 50 ~ "HIGH",
          max_score >= 20 ~ "MEDIUM",
          max_score >= 5 ~ "LOW",
          TRUE ~ "MINIMAL"
        )
        
        valueBox(
          value = efficiency,
          subtitle = "Efficiency Rating",
          icon = icon("star"),
          color = "green"
        )
      } else {
        valueBox(
          value = "N/A",
          subtitle = "Efficiency Rating",
          icon = icon("star"),
          color = "green"
        )
      }
    }
  })
  
  output$ligninDiversity <- renderValueBox({
    if (!is.null(values$processed_data)) {
      enzyme_col <- if("Annotations/Genomes" %in% names(values$processed_data)) {
        "Annotations/Genomes"
      } else {
        names(values$processed_data)[1]
      }
      
      lignin_enzymes <- unique(unlist(values$enzyme_families$lignolytic))
      lignin_pattern <- paste(lignin_enzymes, collapse = "|")
      
      diversity_score <- values$processed_data %>%
        filter(str_detect(!!sym(enzyme_col), lignin_pattern)) %>%
        nrow()
      
      valueBox(
        value = diversity_score,
        subtitle = "Enzyme Diversity",
        icon = icon("leaf"),
        color = "teal"
      )
    }
  })
  
  # Lignolytic distribution plot
  output$ligninaseDistribution <- renderPlotly({
    if (!is.null(values$processed_data)) {
      enzyme_col <- if("Annotations/Genomes" %in% names(values$processed_data)) {
        "Annotations/Genomes"
      } else {
        names(values$processed_data)[1]
      }
      
      lignin_enzymes <- unique(unlist(values$enzyme_families$lignolytic))
      enzyme_pattern <- paste(lignin_enzymes, collapse = "|")
      
      lignin_data <- values$processed_data %>%
        filter(str_detect(!!sym(enzyme_col), enzyme_pattern))
      
      if (nrow(lignin_data) > 0) {
        genome_cols <- names(lignin_data)[!names(lignin_data) %in% c(enzyme_col, "Total")]
        
        if(length(genome_cols) > 0) {
          lignin_data <- lignin_data %>%
            mutate(Total_Count = rowSums(select(., all_of(genome_cols)), na.rm = TRUE)) %>%
            slice_max(Total_Count, n = 15)
          
          plot_data <- lignin_data %>%
            select(all_of(c(enzyme_col, genome_cols))) %>%
            pivot_longer(cols = all_of(genome_cols), names_to = "Genome", values_to = "Count") %>%
            filter(Count > 0)
          
          p <- ggplot(plot_data, aes(x = reorder(!!sym(enzyme_col), Count), y = Count, fill = Genome)) +
            geom_bar(stat = "identity", position = "stack", alpha = 0.8) +
            coord_flip() +
            labs(title = "Lignolytic Enzyme Arsenal Distribution",
                 x = "Enzyme Family", y = "Count") +
            theme_minimal() +
            theme(
              plot.background = element_rect(fill = "transparent", color = NA),
              panel.background = element_rect(fill = "transparent", color = NA),
              text = element_text(color = "white"),
              axis.text = element_text(color = "white"),
              legend.text = element_text(color = "white"),
              legend.title = element_text(color = "white")
            ) +
            scale_fill_viridis_d(option = "magma")
          
          ggplotly(p) %>%
            layout(
              plot_bgcolor = 'rgba(0,0,0,0)',
              paper_bgcolor = 'rgba(0,0,0,0)',
              font = list(color = 'white')
            )%>%
            config(
              toImageButtonOptions = list(
                format = 'png',
                filename = 'Lignolytic_Enzyme_Arsenal_Distribution',
                height = 300,
                width = 700,
                scale = 4  
              )
            )
        }
      }
    }
  })
  
  # Lignin insights
  output$ligninaseInsights <- renderUI({
    if (!is.null(values$processed_data)) {
      HTML(paste(
        "<div class='degradation-card quantum-glow'>",
        "<h4>🌳 LIGNOLYTIC MECHANISM - INDUSTRIAL QUANTUM INSIGHTS</h4>",
        "<p style='color: #cccccc; margin-top: 15px;'>",
        "Lignin degradation employs a complex array of oxidative enzymes that work synergistically to depolymerize the recalcitrant lignin structure.",
        "</p>",
        "<div style='background: rgba(139, 69, 19, 0.1); padding: 15px; border-radius: 10px; margin: 15px 0; border-left: 4px solid #8B4513;'>",
        "<p><strong style='color: #8B4513;'>🔬 OXIDATIVE MECHANISM:</strong></p>",
        "<p style='color: #cccccc;'>Laccases (AA1) and peroxidases (AA2, AA3) generate reactive radicals that cleave C-C and C-O bonds in lignin</p>",
        "</div>",
        "<div style='background: rgba(205, 133, 63, 0.1); padding: 15px; border-radius: 10px; margin: 15px 0; border-left: 4px solid #CD853F;'>",
        "<p><strong style='color: #CD853F;'>⚡ INDUSTRIAL APPLICATIONS:</strong></p>",
        "<p style='color: #cccccc;'>Lignin modification improves cellulose accessibility and enables value-added lignin derivatives production</p>",
        "</div>",
        "</div>"
      ))
    }
  })
  
  # Lignin potential table
  output$ligninasePotentialTable <- DT::renderDataTable({
    if (!is.null(values$processed_data)) {
      lignin_potential <- calculateDegradationPotential(values$processed_data, values$enzyme_families$lignolytic)
      
      if(!is.null(lignin_potential) && length(lignin_potential) > 0) {
        potential_df <- data.frame(
          Genome = names(lignin_potential),
          LigninScore = as.numeric(lignin_potential),
          stringsAsFactors = FALSE
        ) %>%
          arrange(desc(LigninScore)) %>%
          mutate(
            Rank = row_number(),
            Rating = case_when(
              LigninScore >= 50 ~ "🏆 ELITE",
              LigninScore >= 20 ~ "⭐ HIGH", 
              LigninScore >= 10 ~ "✅ MODERATE",
              LigninScore >= 5 ~ "📈 LOW",
              TRUE ~ "⚠️ MINIMAL"
            )
          ) %>%
          slice_head(n = 15)
        
        DT::datatable(potential_df,
                      options = list(
                        pageLength = 10,
                        dom = 'Bfrtip',
                        buttons = c('copy', 'csv'),
                        order = list(list(2, 'desc')),
                        initComplete = JS(
                          "function(settings, json) {",
                          "$(this.api().table().header()).css({'background-color': '#1a1a2e', 'color': '#fff'});",
                          "}"
                        )
                      ),
                      class = 'cell-border stripe hover',
                      rownames = FALSE,
                      colnames = c('Rank', 'Strain', 'Lignin Score', 'Rating')) %>%
          formatStyle(columns = 1:4, 
                      backgroundColor = 'rgba(26, 26, 46, 0.9)', 
                      color = 'white')
      }
    }
  })
  
  # ============================================================================
  # PECTINOLYTIC CAPACITY SERVER FUNCTIONS
  # ============================================================================
  
  output$pectinaseCount <- renderValueBox({
    if (!is.null(values$processed_data)) {
      enzyme_col <- if("Annotations/Genomes" %in% names(values$processed_data)) {
        "Annotations/Genomes"
      } else {
        names(values$processed_data)[1]
      }
      
      pectin_enzymes <- unique(unlist(values$enzyme_families$pectinolytic))
      pectin_pattern <- paste(pectin_enzymes, collapse = "|")
      
      pectin_families <- values$processed_data %>%
        filter(str_detect(!!sym(enzyme_col), pectin_pattern))
      
      count <- nrow(pectin_families)
      
      valueBox(
        value = count,
        subtitle = "Pectinolytic Families",
        icon = icon("apple-alt"),
        color = "red"
      )
    }
  })
  
  output$polygalacturonaseCount <- renderValueBox({
    if (!is.null(values$processed_data)) {
      enzyme_col <- if("Annotations/Genomes" %in% names(values$processed_data)) {
        "Annotations/Genomes"
      } else {
        names(values$processed_data)[1]
      }
      
      polygalacturonases <- c("GH28", "GH33")
      pg_pattern <- paste(polygalacturonases, collapse = "|")
      
      pg_families <- values$processed_data %>%
        filter(str_detect(!!sym(enzyme_col), pg_pattern))
      
      count <- nrow(pg_families)
      
      valueBox(
        value = count,
        subtitle = "Polygalacturonases",
        icon = icon("cut"),
        color = "purple"
      )
    }
  })
  
  output$pectinLyaseCount <- renderValueBox({
    if (!is.null(values$processed_data)) {
      enzyme_col <- if("Annotations/Genomes" %in% names(values$processed_data)) {
        "Annotations/Genomes"
      } else {
        names(values$processed_data)[1]
      }
      
      lyases <- c("PL1", "PL2", "PL3", "PL9", "PL10")
      lyase_pattern <- paste(lyases, collapse = "|")
      
      lyase_families <- values$processed_data %>%
        filter(str_detect(!!sym(enzyme_col), lyase_pattern))
      
      count <- nrow(lyase_families)
      
      valueBox(
        value = count,
        subtitle = "Pectin Lyases",
        icon = icon("scissors"),
        color = "blue"
      )
    }
  })
  
  output$topPectinStrain <- renderValueBox({
    if (!is.null(values$processed_data)) {
      pectin_potential <- calculateDegradationPotential(values$processed_data, values$enzyme_families$pectinolytic)
      
      if (!is.null(pectin_potential) && length(pectin_potential) > 0) {
        top_strain <- names(pectin_potential)[which.max(pectin_potential)]
        display_name <- if(nchar(top_strain) > 8) paste0(substr(top_strain, 1, 8), "...") else top_strain
        
        valueBox(
          value = display_name,
          subtitle = "Top Pectin Producer",
          icon = icon("crown"),
          color = "yellow"
        )
      } else {
        valueBox(
          value = "N/A",
          subtitle = "Top Pectin Producer",
          icon = icon("crown"),
          color = "yellow"
        )
      }
    }
  })
  
  output$pectinEfficiency <- renderValueBox({
    if (!is.null(values$processed_data)) {
      pectin_potential <- calculateDegradationPotential(values$processed_data, values$enzyme_families$pectinolytic)
      
      if (!is.null(pectin_potential) && length(pectin_potential) > 0) {
        max_score <- max(unlist(pectin_potential))
        
        efficiency <- case_when(
          max_score >= 75 ~ "ELITE",
          max_score >= 40 ~ "HIGH",
          max_score >= 20 ~ "MEDIUM",
          TRUE ~ "LOW"
        )
        
        valueBox(
          value = efficiency,
          subtitle = "Efficiency Rating",
          icon = icon("star"),
          color = "green"
        )
      } else {
        valueBox(
          value = "N/A",
          subtitle = "Efficiency Rating",
          icon = icon("star"),
          color = "green"
        )
      }
    }
  })
  
  output$pectinDiversity <- renderValueBox({
    if (!is.null(values$processed_data)) {
      enzyme_col <- if("Annotations/Genomes" %in% names(values$processed_data)) {
        "Annotations/Genomes"
      } else {
        names(values$processed_data)[1]
      }
      
      pectin_enzymes <- unique(unlist(values$enzyme_families$pectinolytic))
      pectin_pattern <- paste(pectin_enzymes, collapse = "|")
      
      diversity_score <- values$processed_data %>%
        filter(str_detect(!!sym(enzyme_col), pectin_pattern)) %>%
        nrow()
      
      valueBox(
        value = diversity_score,
        subtitle = "Enzyme Diversity",
        icon = icon("leaf"),
        color = "teal"
      )
    }
  })
  
  # Pectinolytic distribution plot
  output$pectinaseDistribution <- renderPlotly({
    if (!is.null(values$processed_data)) {
      enzyme_col <- if("Annotations/Genomes" %in% names(values$processed_data)) {
        "Annotations/Genomes"
      } else {
        names(values$processed_data)[1]
      }
      
      pectin_enzymes <- unique(unlist(values$enzyme_families$pectinolytic))
      enzyme_pattern <- paste(pectin_enzymes, collapse = "|")
      
      pectin_data <- values$processed_data %>%
        filter(str_detect(!!sym(enzyme_col), enzyme_pattern))
      
      if (nrow(pectin_data) > 0) {
        genome_cols <- names(pectin_data)[!names(pectin_data) %in% c(enzyme_col, "Total")]
        
        if(length(genome_cols) > 0) {
          pectin_data <- pectin_data %>%
            mutate(Total_Count = rowSums(select(., all_of(genome_cols)), na.rm = TRUE)) %>%
            slice_max(Total_Count, n = 15)
          
          plot_data <- pectin_data %>%
            select(all_of(c(enzyme_col, genome_cols))) %>%
            pivot_longer(cols = all_of(genome_cols), names_to = "Genome", values_to = "Count") %>%
            filter(Count > 0)
          
          p <- ggplot(plot_data, aes(x = reorder(!!sym(enzyme_col), Count), y = Count, fill = Genome)) +
            geom_bar(stat = "identity", position = "stack", alpha = 0.8) +
            coord_flip() +
            labs(title = "Pectinolytic Enzyme Arsenal Distribution",
                 x = "Enzyme Family", y = "Count") +
            theme_minimal() +
            theme(
              plot.background = element_rect(fill = "transparent", color = NA),
              panel.background = element_rect(fill = "transparent", color = NA),
              text = element_text(color = "white"),
              axis.text = element_text(color = "white"),
              legend.text = element_text(color = "white"),
              legend.title = element_text(color = "white")
            ) +
            scale_fill_viridis_d(option = "plasma")
          
          ggplotly(p) %>%
            layout(
              plot_bgcolor = 'rgba(0,0,0,0)',
              paper_bgcolor = 'rgba(0,0,0,0)',
              font = list(color = 'white')
            )%>%
            config(
              toImageButtonOptions = list(
                format = 'png',
                filename = 'Pectinolytic_Enzyme_Arsenal_Distribution',
                height = 300,
                width = 700,
                scale = 4  
              )
            )
        }
      }
    }
  })
  
  # ============================================================================
  # STARCH DEGRADATION SERVER FUNCTIONS
  # ============================================================================
  
  output$amylaseCount <- renderValueBox({
    if (!is.null(values$processed_data)) {
      enzyme_col <- if("Annotations/Genomes" %in% names(values$processed_data)) {
        "Annotations/Genomes"
      } else {
        names(values$processed_data)[1]
      }
      
      amylases <- c("GH13", "GH14")
      amylase_pattern <- paste(amylases, collapse = "|")
      
      amylase_families <- values$processed_data %>%
        filter(str_detect(!!sym(enzyme_col), amylase_pattern))
      
      count <- nrow(amylase_families)
      
      valueBox(
        value = count,
        subtitle = "Amylase Families",
        icon = icon("seedling"),
        color = "yellow"
      )
    }
  })
  
  output$glucoamylaseCount <- renderValueBox({
    if (!is.null(values$processed_data)) {
      enzyme_col <- if("Annotations/Genomes" %in% names(values$processed_data)) {
        "Annotations/Genomes"
      } else {
        names(values$processed_data)[1]
      }
      
      glucoamylases <- c("GH15")
      ga_pattern <- paste(glucoamylases, collapse = "|")
      
      ga_families <- values$processed_data %>%
        filter(str_detect(!!sym(enzyme_col), ga_pattern))
      
      count <- nrow(ga_families)
      
      valueBox(
        value = count,
        subtitle = "Glucoamylases",
        icon = icon("burn"),
        color = "orange"
      )
    }
  })
  
  output$pullulanaseCount <- renderValueBox({
    if (!is.null(values$processed_data)) {
      enzyme_col <- if("Annotations/Genomes" %in% names(values$processed_data)) {
        "Annotations/Genomes"
      } else {
        names(values$processed_data)[1]
      }
      
      pullulanases <- c("GH57", "GH133")
      pull_pattern <- paste(pullulanases, collapse = "|")
      
      pull_families <- values$processed_data %>%
        filter(str_detect(!!sym(enzyme_col), pull_pattern))
      
      count <- nrow(pull_families)
      
      valueBox(
        value = count,
        subtitle = "Debranching Enzymes",
        icon = icon("cut"),
        color = "blue"
      )
    }
  })
  
  output$topStarchStrain <- renderValueBox({
    if (!is.null(values$processed_data)) {
      starch_potential <- calculateDegradationPotential(values$processed_data, values$enzyme_families$starch_degrading)
      
      if (!is.null(starch_potential) && length(starch_potential) > 0) {
        top_strain <- names(starch_potential)[which.max(starch_potential)]
        display_name <- if(nchar(top_strain) > 8) paste0(substr(top_strain, 1, 8), "...") else top_strain
        
        valueBox(
          value = display_name,
          subtitle = "Top Starch Producer",
          icon = icon("crown"),
          color = "purple"
        )
      } else {
        valueBox(
          value = "N/A",
          subtitle = "Top Starch Producer",
          icon = icon("crown"),
          color = "purple"
        )
      }
    }
  })
  
  output$starchEfficiency <- renderValueBox({
    if (!is.null(values$processed_data)) {
      starch_potential <- calculateDegradationPotential(values$processed_data, values$enzyme_families$starch_degrading)
      
      if (!is.null(starch_potential) && length(starch_potential) > 0) {
        max_score <- max(unlist(starch_potential))
        
        efficiency <- case_when(
          max_score >= 100 ~ "ELITE",
          max_score >= 50 ~ "HIGH",
          max_score >= 25 ~ "MEDIUM",
          TRUE ~ "LOW"
        )
        
        valueBox(
          value = efficiency,
          subtitle = "Efficiency Rating",
          icon = icon("star"),
          color = "green"
        )
      } else {
        valueBox(
          value = "N/A",
          subtitle = "Efficiency Rating",
          icon = icon("star"),
          color = "green"
        )
      }
    }
  })
  
  output$starchDiversity <- renderValueBox({
    if (!is.null(values$processed_data)) {
      enzyme_col <- if("Annotations/Genomes" %in% names(values$processed_data)) {
        "Annotations/Genomes"
      } else {
        names(values$processed_data)[1]
      }
      
      starch_enzymes <- unique(unlist(values$enzyme_families$starch_degrading))
      starch_pattern <- paste(starch_enzymes, collapse = "|")
      
      diversity_score <- values$processed_data %>%
        filter(str_detect(!!sym(enzyme_col), starch_pattern)) %>%
        nrow()
      
      valueBox(
        value = diversity_score,
        subtitle = "Enzyme Diversity",
        icon = icon("leaf"),
        color = "teal"
      )
    }
  })
  
  # Starch degrading distribution plot
  output$starchDegradingDistribution <- renderPlotly({
    if (!is.null(values$processed_data)) {
      enzyme_col <- if("Annotations/Genomes" %in% names(values$processed_data)) {
        "Annotations/Genomes"
      } else {
        names(values$processed_data)[1]
      }
      
      starch_enzymes <- unique(unlist(values$enzyme_families$starch_degrading))
      enzyme_pattern <- paste(starch_enzymes, collapse = "|")
      
      starch_data <- values$processed_data %>%
        filter(str_detect(!!sym(enzyme_col), enzyme_pattern))
      
      if (nrow(starch_data) > 0) {
        genome_cols <- names(starch_data)[!names(starch_data) %in% c(enzyme_col, "Total")]
        
        if(length(genome_cols) > 0) {
          starch_data <- starch_data %>%
            mutate(Total_Count = rowSums(select(., all_of(genome_cols)), na.rm = TRUE)) %>%
            slice_max(Total_Count, n = 15)
          
          plot_data <- starch_data %>%
            select(all_of(c(enzyme_col, genome_cols))) %>%
            pivot_longer(cols = all_of(genome_cols), names_to = "Genome", values_to = "Count") %>%
            filter(Count > 0)
          
          p <- ggplot(plot_data, aes(x = reorder(!!sym(enzyme_col), Count), y = Count, fill = Genome)) +
            geom_bar(stat = "identity", position = "stack", alpha = 0.8) +
            coord_flip() +
            labs(title = "Starch Degrading Enzyme Arsenal Distribution",
                 x = "Enzyme Family", y = "Count") +
            theme_minimal() +
            theme(
              plot.background = element_rect(fill = "transparent", color = NA),
              panel.background = element_rect(fill = "transparent", color = NA),
              text = element_text(color = "white"),
              axis.text = element_text(color = "white"),
              legend.text = element_text(color = "white"),
              legend.title = element_text(color = "white")
            ) +
            scale_fill_viridis_d(option = "cividis")
          
          ggplotly(p) %>%
            layout(
              plot_bgcolor = 'rgba(0,0,0,0)',
              paper_bgcolor = 'rgba(0,0,0,0)',
              font = list(color = 'white')
            )%>%
            config(
              toImageButtonOptions = list(
                format = 'png',
                filename = 'Starch_Degrading_Enzyme_Arsenal_Distribution',
                height = 300,
                width = 700,
                scale = 4  
              )
            )
        }
      }
    }
  })
  
  # ============================================================================
  # INULIN DEGRADATION SERVER FUNCTIONS
  # ============================================================================
  
  output$inulinaseCount <- renderValueBox({
    if (!is.null(values$processed_data)) {
      enzyme_col <- if("Annotations/Genomes" %in% names(values$processed_data)) {
        "Annotations/Genomes"
      } else {
        names(values$processed_data)[1]
      }
      
      inulin_enzymes <- unique(unlist(values$enzyme_families$inulin_degrading))
      inulin_pattern <- paste(inulin_enzymes, collapse = "|")
      
      inulin_families <- values$processed_data %>%
        filter(str_detect(!!sym(enzyme_col), inulin_pattern))
      
      count <- nrow(inulin_families)
      
      valueBox(
        value = count,
        subtitle = "Inulinase Families",
        icon = icon("carrot"),
        color = "purple"
      )
    }
  })
  
  output$fructanaseCount <- renderValueBox({
    if (!is.null(values$processed_data)) {
      enzyme_col <- if("Annotations/Genomes" %in% names(values$processed_data)) {
        "Annotations/Genomes"
      } else {
        names(values$processed_data)[1]
      }
      
      fructanases <- c("GH32")
      fruc_pattern <- paste(fructanases, collapse = "|")
      
      fruc_families <- values$processed_data %>%
        filter(str_detect(!!sym(enzyme_col), fruc_pattern))
      
      count <- nrow(fruc_families)
      
      valueBox(
        value = count,
        subtitle = "Fructanases",
        icon = icon("leaf"),
        color = "green"
      )
    }
  })
  
  output$levanaseCount <- renderValueBox({
    if (!is.null(values$processed_data)) {
      enzyme_col <- if("Annotations/Genomes" %in% names(values$processed_data)) {
        "Annotations/Genomes"
      } else {
        names(values$processed_data)[1]
      }
      
      levanases <- c("GH32")
      lev_pattern <- paste(levanases, collapse = "|")
      
      lev_families <- values$processed_data %>%
        filter(str_detect(!!sym(enzyme_col), lev_pattern))
      
      count <- nrow(lev_families)
      
      valueBox(
        value = count,
        subtitle = "Levanases",
        icon = icon("seedling"),
        color = "blue"
      )
    }
  })
  
  output$topInulinStrain <- renderValueBox({
    if (!is.null(values$processed_data)) {
      inulin_potential <- calculateDegradationPotential(values$processed_data, values$enzyme_families$inulin_degrading)
      
      if (!is.null(inulin_potential) && length(inulin_potential) > 0) {
        top_strain <- names(inulin_potential)[which.max(inulin_potential)]
        display_name <- if(nchar(top_strain) > 8) paste0(substr(top_strain, 1, 8), "...") else top_strain
        
        valueBox(
          value = display_name,
          subtitle = "Top Inulin Producer",
          icon = icon("crown"),
          color = "yellow"
        )
      } else {
        valueBox(
          value = "N/A",
          subtitle = "Top Inulin Producer",
          icon = icon("crown"),
          color = "yellow"
        )
      }
    }
  })
  
  output$inulinEfficiency <- renderValueBox({
    if (!is.null(values$processed_data)) {
      inulin_potential <- calculateDegradationPotential(values$processed_data, values$enzyme_families$inulin_degrading)
      
      if (!is.null(inulin_potential) && length(inulin_potential) > 0) {
        max_score <- max(unlist(inulin_potential))
        
        efficiency <- case_when(
          max_score >= 30 ~ "HIGH",
          max_score >= 15 ~ "MEDIUM",
          max_score >= 5 ~ "LOW",
          TRUE ~ "MINIMAL"
        )
        
        valueBox(
          value = efficiency,
          subtitle = "Efficiency Rating",
          icon = icon("star"),
          color = "red"
        )
      } else {
        valueBox(
          value = "N/A",
          subtitle = "Efficiency Rating",
          icon = icon("star"),
          color = "red"
        )
      }
    }
  })
  
  output$inulinDiversity <- renderValueBox({
    if (!is.null(values$processed_data)) {
      enzyme_col <- if("Annotations/Genomes" %in% names(values$processed_data)) {
        "Annotations/Genomes"
      } else {
        names(values$processed_data)[1]
      }
      
      inulin_enzymes <- unique(unlist(values$enzyme_families$inulin_degrading))
      inulin_pattern <- paste(inulin_enzymes, collapse = "|")
      
      diversity_score <- values$processed_data %>%
        filter(str_detect(!!sym(enzyme_col), inulin_pattern)) %>%
        nrow()
      
      valueBox(
        value = diversity_score,
        subtitle = "Enzyme Diversity",
        icon = icon("leaf"),
        color = "teal"
      )
    }
  })
  
  # Inulin degrading distribution plot
  output$inulinDegradingDistribution <- renderPlotly({
    if (!is.null(values$processed_data)) {
      enzyme_col <- if("Annotations/Genomes" %in% names(values$processed_data)) {
        "Annotations/Genomes"
      } else {
        names(values$processed_data)[1]
      }
      
      inulin_enzymes <- unique(unlist(values$enzyme_families$inulin_degrading))
      enzyme_pattern <- paste(inulin_enzymes, collapse = "|")
      
      inulin_data <- values$processed_data %>%
        filter(str_detect(!!sym(enzyme_col), enzyme_pattern))
      
      if (nrow(inulin_data) > 0) {
        genome_cols <- names(inulin_data)[!names(inulin_data) %in% c(enzyme_col, "Total")]
        
        if(length(genome_cols) > 0) {
          inulin_data <- inulin_data %>%
            mutate(Total_Count = rowSums(select(., all_of(genome_cols)), na.rm = TRUE)) %>%
            slice_max(Total_Count, n = 15)
          
          plot_data <- inulin_data %>%
            select(all_of(c(enzyme_col, genome_cols))) %>%
            pivot_longer(cols = all_of(genome_cols), names_to = "Genome", values_to = "Count") %>%
            filter(Count > 0)
          
          p <- ggplot(plot_data, aes(x = reorder(!!sym(enzyme_col), Count), y = Count, fill = Genome)) +
            geom_bar(stat = "identity", position = "stack", alpha = 0.8) +
            coord_flip() +
            labs(title = "Inulin Degrading Enzyme Arsenal Distribution",
                 x = "Enzyme Family", y = "Count") +
            theme_minimal() +
            theme(
              plot.background = element_rect(fill = "transparent", color = NA),
              panel.background = element_rect(fill = "transparent", color = NA),
              text = element_text(color = "white"),
              axis.text = element_text(color = "white"),
              legend.text = element_text(color = "white"),
              legend.title = element_text(color = "white")
            ) +
            scale_fill_viridis_d(option = "turbo")
          
          ggplotly(p) %>%
            layout(
              plot_bgcolor = 'rgba(0,0,0,0)',
              paper_bgcolor = 'rgba(0,0,0,0)',
              font = list(color = 'white')
            )%>%
            config(
              toImageButtonOptions = list(
                format = 'png',
                filename = 'Inulin_Degrading_Enzyme_Arsenal_Distribution',
                height = 300,
                width = 700,
                scale = 4  
              )
            )
        }
      }
    }
  })
  
  # Remove the old optimization parameters UI and replace with:
  # First, add a reactive strain selector server logic
  output$strainSelector <- renderUI({
    if (!is.null(values$processed_data)) {
      genome_cols <- names(values$processed_data)[!names(values$processed_data) %in% c("Annotations/Genomes", "Total")]
      
      if(length(genome_cols) > 0) {
        selectInput("selectedTestStrain", 
                    label = tags$span("🦠 Select Test Microbial Strain:", 
                                      style = "color: #00d4ff; font-weight: bold; font-family: 'Inter', sans-serif;"),
                    choices = c("All Strains Overview" = "all", setNames(genome_cols, substr(genome_cols, 1, 20))),
                    selected = "all",
                    width = "100%")
      }
    }
  })
  
  # Main enzyme family analytics with reactive strain filtering
  # Fixed Enzyme Family Analytics with Proper Reactivity
  
  output$enzymeFamilyAnalytics <- renderUI({
    
    # Add error handling wrapper
    tryCatch({
      
      if (is.null(values$processed_data)) {
        return(
          div(class = "metric-card",
              h4("🧬 ENZYME FAMILY ANALYTICS", style = "color: #00d4ff; font-family: Orbitron, monospace;"),
              div(style = "text-align: center; padding: 30px 15px; color: #cccccc;",
                  div("⚡", style = "font-size: 2.5em; margin-bottom: 15px;"),
                  p("Advanced enzyme family analysis ready for data input")
              )
          )
        )
      }
      
      # Safely get genome columns
      genome_cols <- names(values$processed_data)[!names(values$processed_data) %in% c("Annotations/Genomes", "Total")]
      
      if(length(genome_cols) == 0) {
        return(
          div(class = "metric-card",
              h4("🧬 ENZYME FAMILY ANALYTICS", style = "color: #00d4ff; font-family: Orbitron, monospace;"),
              div(style = "text-align: center; padding: 30px 15px; color: #cccccc;",
                  div("📊", style = "font-size: 2.5em; margin-bottom: 15px;"),
                  p("Upload and process genomic data to view detailed enzyme family analytics")
              )
          )
        )
      }
      
      # FIXED: Remove isolate() and get selected strain reactively
      selected_strain <- if(!is.null(input$selectedTestStrain) && input$selectedTestStrain != "all") {
        input$selectedTestStrain
      } else {
        "all"
      }
      
      # Initialize variables with safe defaults
      cellulolytic_counts <- NULL
      hemi_counts <- NULL
      lignolytic_counts <- NULL
      
      # Safely calculate degradation potential
      if(!is.null(values$enzyme_families) && !is.null(values$enzyme_families$cellulolytic)) {
        cellulolytic_counts <- tryCatch({
          calculateDegradationPotential(values$processed_data, values$enzyme_families$cellulolytic)
        }, error = function(e) NULL)
      }
      
      if(!is.null(values$enzyme_families) && !is.null(values$enzyme_families$hemi_cellulolytic)) {
        hemi_counts <- tryCatch({
          calculateDegradationPotential(values$processed_data, values$enzyme_families$hemi_cellulolytic)
        }, error = function(e) NULL)
      }
      
      if(!is.null(values$enzyme_families) && !is.null(values$enzyme_families$lignolytic)) {
        lignolytic_counts <- tryCatch({
          calculateDegradationPotential(values$processed_data, values$enzyme_families$lignolytic)
        }, error = function(e) NULL)
      }
      
      # Calculate metrics based on selection
      if(selected_strain == "all") {
        # All strains overview
        top_cellulolytic <- if(!is.null(cellulolytic_counts) && length(cellulolytic_counts) > 0) {
          names(cellulolytic_counts)[which.max(cellulolytic_counts)]
        } else "N/A"
        
        top_hemi <- if(!is.null(hemi_counts) && length(hemi_counts) > 0) {
          names(hemi_counts)[which.max(hemi_counts)]
        } else "N/A"
        
        top_lignolytic <- if(!is.null(lignolytic_counts) && length(lignolytic_counts) > 0) {
          names(lignolytic_counts)[which.max(lignolytic_counts)]
        } else "N/A"
        
        # Calculate synergy scores safely
        synergy_scores <- data.frame(
          Strain = genome_cols,
          Synergy = numeric(length(genome_cols)),
          stringsAsFactors = FALSE
        )
        
        for(i in seq_along(genome_cols)) {
          strain <- genome_cols[i]
          cell_score <- if(!is.null(cellulolytic_counts) && strain %in% names(cellulolytic_counts)) {
            as.numeric(cellulolytic_counts[[strain]]) 
          } else 0
          
          hemi_score <- if(!is.null(hemi_counts) && strain %in% names(hemi_counts)) {
            as.numeric(hemi_counts[[strain]]) 
          } else 0
          
          lignin_score <- if(!is.null(lignolytic_counts) && strain %in% names(lignolytic_counts)) {
            as.numeric(lignolytic_counts[[strain]]) 
          } else 0
          
          synergy_scores$Synergy[i] <- sqrt(cell_score * hemi_score * lignin_score)
        }
        
        top_synergy_strain <- if(nrow(synergy_scores) > 0 && max(synergy_scores$Synergy, na.rm = TRUE) > 0) {
          synergy_scores$Strain[which.max(synergy_scores$Synergy)]
        } else "N/A"
        
        max_synergy_score <- if(nrow(synergy_scores) > 0) max(synergy_scores$Synergy, na.rm = TRUE) else 0
        
        strain_title <- "ALL STRAINS OVERVIEW"
        strain_count_display <- length(genome_cols)
        
        # Display totals
        cell_display <- if(!is.null(cellulolytic_counts)) sum(as.numeric(cellulolytic_counts), na.rm = TRUE) else 0
        hemi_display <- if(!is.null(hemi_counts)) sum(as.numeric(hemi_counts), na.rm = TRUE) else 0
        lignin_display <- if(!is.null(lignolytic_counts)) sum(as.numeric(lignolytic_counts), na.rm = TRUE) else 0
        
      } else {
        # FIXED: Individual strain analysis
        strain_cell_count <- if(!is.null(cellulolytic_counts) && selected_strain %in% names(cellulolytic_counts)) {
          as.numeric(cellulolytic_counts[[selected_strain]])
        } else 0
        
        strain_hemi_count <- if(!is.null(hemi_counts) && selected_strain %in% names(hemi_counts)) {
          as.numeric(hemi_counts[[selected_strain]])
        } else 0
        
        strain_lignin_count <- if(!is.null(lignolytic_counts) && selected_strain %in% names(lignolytic_counts)) {
          as.numeric(lignolytic_counts[[selected_strain]])
        } else 0
        
        # FIXED: Show the actual selected strain as top performer
        top_cellulolytic <- selected_strain
        top_hemi <- selected_strain
        top_lignolytic <- selected_strain
        
        synergy_score <- sqrt(strain_cell_count * strain_hemi_count * strain_lignin_count)
        
        # FIXED: Use selected strain name for synergy display
        top_synergy_strain <- selected_strain
        max_synergy_score <- synergy_score
        
        # FIXED: Display selected strain name properly
        strain_title <- paste("STRAIN:", substr(selected_strain, 1, 20))
        strain_count_display <- 1
        
        # FIXED: Display individual strain counts (not totals)
        cell_display <- strain_cell_count
        hemi_display <- strain_hemi_count
        lignin_display <- strain_lignin_count
      }
      
      # Safe family counts
      cellulolytic_family_count <- if(!is.null(values$enzyme_families$cellulolytic)) length(values$enzyme_families$cellulolytic) else 0
      hemi_family_count <- if(!is.null(values$enzyme_families$hemi_cellulolytic)) length(values$enzyme_families$hemi_cellulolytic) else 0
      lignin_family_count <- if(!is.null(values$enzyme_families$lignolytic)) length(values$enzyme_families$lignolytic) else 0
      
      total_families <- cellulolytic_family_count + hemi_family_count + lignin_family_count
      
      # Calculate diversity index safely
      if(selected_strain == "all") {
        strains_with_cell <- if(!is.null(cellulolytic_counts)) sum(as.numeric(cellulolytic_counts) > 0, na.rm = TRUE) else 0
        strains_with_hemi <- if(!is.null(hemi_counts)) sum(as.numeric(hemi_counts) > 0, na.rm = TRUE) else 0
        strains_with_lignin <- if(!is.null(lignolytic_counts)) sum(as.numeric(lignolytic_counts) > 0, na.rm = TRUE) else 0
        
        avg_coverage <- mean(c(strains_with_cell, strains_with_hemi, strains_with_lignin), na.rm = TRUE) / length(genome_cols)
        diversity_index <- round(avg_coverage * 100, 1)
      } else {
        has_cellulolytic <- cell_display > 0
        has_hemi <- hemi_display > 0
        has_lignolytic <- lignin_display > 0
        
        family_coverage <- sum(c(has_cellulolytic, has_hemi, has_lignolytic))
        diversity_index <- round((family_coverage / 3) * 100, 1)
      }
      
      # Ensure numeric values are properly formatted
      cell_display <- as.integer(cell_display)
      hemi_display <- as.integer(hemi_display)
      lignin_display <- as.integer(lignin_display)
      max_synergy_score <- round(as.numeric(max_synergy_score), 1)
      diversity_index <- round(as.numeric(diversity_index), 1)
      
      # Create the UI
      div(class = "metric-card quantum-glow", style = "padding: 15px;",
          h4("🧬 ENZYME FAMILY ANALYTICS", style = "color: #00d4ff; font-family: Orbitron, monospace; margin-bottom: 15px;"),
          
          # EMBEDDED STRAIN SELECTOR
          div(style = "margin: 0 0 15px 0; padding: 12px; background: linear-gradient(135deg, rgba(0, 212, 255, 0.1), rgba(0, 212, 255, 0.02)); border-radius: 8px; border: 1px solid #00d4ff;",
              div(style = "margin-bottom: 8px;",
                  tags$span("🦠 Select Test Microbial Strain:", style = "color: #00d4ff; font-weight: bold; font-family: 'Inter', sans-serif; font-size: 0.9em;")
              ),
              selectInput("selectedTestStrain", 
                          label = NULL,
                          choices = c("All Strains Overview" = "all", setNames(genome_cols, paste0(substr(genome_cols, 1, 25), ifelse(nchar(genome_cols) > 25, "...", "")))),
                          selected = selected_strain,
                          width = "100%")
          ),
          
          h5(strain_title, style = "color: #38ef7d; font-family: Inter, sans-serif; margin-bottom: 15px;"),
          
          # Family Distribution
          div(style = "margin: 15px 0;",
              h5("📊 Family Distribution", style = "color: #38ef7d; margin-bottom: 10px; font-size: 1.1em;"),
              div(style = "display: flex; gap: 8px; margin-bottom: 12px; flex-wrap: wrap;",
                  
                  # FIXED: Cellulolytic - show strain-specific name
                  div(style = "background: linear-gradient(135deg, rgba(0, 212, 255, 0.15), rgba(0, 212, 255, 0.03)); padding: 8px 12px; border-radius: 6px; border: 1px solid #00d4ff; flex: 1; min-width: 110px; display: flex; align-items: center; gap: 6px;",
                      tags$span("🌾", style = "font-size: 1.2em;"),
                      div(style = "text-align: center;",
                          div("CELLULOLYTIC", style = "color: #00d4ff; font-weight: bold; font-size: 0.75em; line-height: 1;"),
                          div(as.character(cell_display), style = "font-size: 1.3em; font-weight: bold; color: #ffffff; line-height: 1; margin: 2px 0;"),
                          div(if(selected_strain == "all") substr(top_cellulolytic, 1, 6) else substr(selected_strain, 1, 6), 
                              style = "color: #00d4ff; font-size: 0.6em; font-weight: bold; line-height: 1;")
                      )
                  ),
                  
                  # FIXED: Hemicellulosic - show strain-specific name  
                  div(style = "background: linear-gradient(135deg, rgba(56, 239, 125, 0.15), rgba(56, 239, 125, 0.03)); padding: 8px 12px; border-radius: 6px; border: 1px solid #38ef7d; flex: 1; min-width: 110px; display: flex; align-items: center; gap: 6px;",
                      tags$span("🌿", style = "font-size: 1.2em;"),
                      div(style = "text-align: center;",
                          div("HEMICELLULOSIC", style = "color: #38ef7d; font-weight: bold; font-size: 0.75em; line-height: 1;"),
                          div(as.character(hemi_display), style = "font-size: 1.3em; font-weight: bold; color: #ffffff; line-height: 1; margin: 2px 0;"),
                          div(if(selected_strain == "all") substr(top_hemi, 1, 6) else substr(selected_strain, 1, 6), 
                              style = "color: #38ef7d; font-size: 0.6em; font-weight: bold; line-height: 1;")
                      )
                  ),
                  
                  # FIXED: Lignolytic - show strain-specific name
                  div(style = "background: linear-gradient(135deg, rgba(255, 107, 53, 0.15), rgba(255, 107, 53, 0.03)); padding: 8px 12px; border-radius: 6px; border: 1px solid #ff6b35; flex: 1; min-width: 110px; display: flex; align-items: center; gap: 6px;",
                      tags$span("🌳", style = "font-size: 1.2em;"),
                      div(style = "text-align: center;",
                          div("LIGNOLYTIC", style = "color: #ff6b35; font-weight: bold; font-size: 0.75em; line-height: 1;"),
                          div(as.character(lignin_display), style = "font-size: 1.3em; font-weight: bold; color: #ffffff; line-height: 1; margin: 2px 0;"),
                          div(if(selected_strain == "all") substr(top_lignolytic, 1, 6) else substr(selected_strain, 1, 6), 
                              style = "color: #ff6b35; font-size: 0.6em; font-weight: bold; line-height: 1;")
                      )
                  )
              )
          ),
          
          # Performance Metrics
          div(style = "margin: 12px 0;",
              h5("⚡ Performance Metrics", style = "color: #9c59d1; margin-bottom: 8px; font-size: 1.1em;"),
              div(style = "display: flex; gap: 8px;",
                  
                  # Diversity Index
                  div(style = "background: linear-gradient(135deg, rgba(156, 89, 209, 0.15), rgba(156, 89, 209, 0.03)); padding: 10px; border-radius: 6px; border: 1px solid #9c59d1; flex: 1; text-align: center;",
                      div(style = "display: flex; align-items: center; justify-content: center; gap: 6px; margin-bottom: 4px;",
                          tags$span("🎯", style = "font-size: 1.3em;"),
                          div("DIVERSITY", style = "color: #9c59d1; font-weight: bold; font-size: 0.8em;")
                      ),
                      div(paste0(diversity_index, "%"), style = "font-size: 1.4em; font-weight: bold; color: #ffffff; margin: 2px 0;"),
                      div(style = "height: 3px; background: rgba(156, 89, 209, 0.3); border-radius: 2px; margin-top: 4px;",
                          div(style = paste0("height: 100%; width: ", pmin(100, pmax(0, diversity_index)), "%; background: #9c59d1; border-radius: 2px;"))
                      )
                  ),
                  
                  # FIXED: Synergy Score - show correct strain name
                  div(style = "background: linear-gradient(135deg, rgba(255, 193, 7, 0.15), rgba(255, 193, 7, 0.03)); padding: 10px; border-radius: 6px; border: 1px solid #ffc107; flex: 1; text-align: center;",
                      div(style = "display: flex; align-items: center; justify-content: center; gap: 6px; margin-bottom: 4px;",
                          tags$span("🔥", style = "font-size: 1.3em;"),
                          div("SYNERGY", style = "color: #ffc107; font-weight: bold; font-size: 0.8em;")
                      ),
                      div(if(selected_strain == "all") substr(top_synergy_strain, 1, 8) else substr(selected_strain, 1, 8), 
                          style = "font-size: 1.1em; font-weight: bold; color: #ffffff; margin: 1px 0;"),
                      div(as.character(max_synergy_score), style = "font-size: 1.2em; font-weight: bold; color: #ffc107; margin: 1px 0;")
                  )
              )
          ),
          
          # Performance Summary
          div(style = "background: rgba(56, 239, 125, 0.08); padding: 8px; border-radius: 6px; border-left: 3px solid #38ef7d; margin-top: 12px;",
              h6("📈 PERFORMANCE SUMMARY", style = "color: #38ef7d; font-family: Orbitron, monospace; margin: 0 0 6px 0; font-size: 0.85em;"),
              div(style = "display: flex; flex-wrap: wrap; gap: 8px; color: #cccccc; font-size: 0.75em;",
                  tags$span(paste("📊 Families:", total_families)),
                  tags$span(paste("🦠 Strains:", strain_count_display)),
                  tags$span(paste("⚡ Score:", max_synergy_score))
              )
          )
      )
      
    }, error = function(e) {
      # Error fallback UI
      div(class = "metric-card", style = "padding: 15px;",
          h4("🧬 ENZYME FAMILY ANALYTICS", style = "color: #ff6b35; font-family: Orbitron, monospace; margin-bottom: 15px;"),
          div(style = "text-align: center; padding: 20px; color: #ffcc00; background: rgba(255, 107, 53, 0.1); border-radius: 8px; border: 1px solid #ff6b35;",
              div("⚠️", style = "font-size: 2em; margin-bottom: 10px;"),
              p("Error loading enzyme analytics. Please check your data format and try again."),
              p(paste("Error details:", e$message), style = "font-size: 0.8em; color: #cccccc; margin-top: 10px;")
          )
      )
    })
  })
  
  # Biorefinery optimization outputs
  output$biorefineryRecommendations <- renderUI({
    if (!is.null(values$processed_data)) {
      genome_cols <- names(values$processed_data)[!names(values$processed_data) %in% c("Annotations/Genomes", "Total")]
      
      if(length(genome_cols) > 0) {
        # Calculate comprehensive scores for all strains
        recommendations <- data.frame(
          Strain = genome_cols,
          Cellulolytic = 0,
          HemiCellulolytic = 0,
          Lignolytic = 0,
          Overall = 0,
          Rank = 0,
          stringsAsFactors = FALSE
        )
        
        # Calculate scores
        for(i in 1:nrow(recommendations)) {
          strain <- recommendations$Strain[i]
          
          cell_pot <- calculateDegradationPotential(values$processed_data, values$enzyme_families$cellulolytic)
          hemi_pot <- calculateDegradationPotential(values$processed_data, values$enzyme_families$hemi_cellulolytic)
          lignin_pot <- calculateDegradationPotential(values$processed_data, values$enzyme_families$lignolytic)
          
          recommendations$Cellulolytic[i] <- if(!is.null(cell_pot) && strain %in% names(cell_pot)) cell_pot[[strain]] else 0
          recommendations$HemiCellulolytic[i] <- if(!is.null(hemi_pot) && strain %in% names(hemi_pot)) hemi_pot[[strain]] else 0
          recommendations$Lignolytic[i] <- if(!is.null(lignin_pot) && strain %in% names(lignin_pot)) lignin_pot[[strain]] else 0
          
          # Calculate weighted overall score
          recommendations$Overall[i] <- recommendations$Cellulolytic[i] * 0.4 + 
            recommendations$HemiCellulolytic[i] * 0.4 + 
            recommendations$Lignolytic[i] * 0.2
        }
        
        # Rank strains
        recommendations <- recommendations %>%
          arrange(desc(Overall)) %>%
          mutate(Rank = row_number()) %>%
          slice_head(n = 10)
        
        # Generate HTML recommendations
        top_3 <- recommendations %>% slice_head(n = 3)
        
        recommendation_html <- paste(
          "<div class='metric-card quantum-glow'>",
          "<h4 style='color: #00d4ff; font-family: Orbitron, monospace;'>🎯 BIOREFINERY RECOMMENDATIONS</h4>",
          "<div style='display: grid; grid-template-columns: 1fr 1fr 1fr; gap: 20px; margin: 20px 0;'>",
          paste(
            mapply(function(rank, strain, cell, hemi, lignin, overall) {
              badge_color <- case_when(
                rank == 1 ~ "#FFD700",
                rank == 2 ~ "#C0C0C0", 
                rank == 3 ~ "#CD7F32",
                TRUE ~ "#666666"
              )
              
              paste(
                "<div style='background: linear-gradient(135deg, rgba(0, 212, 255, 0.1) 0%, rgba(255, 107, 53, 0.1) 100%); padding: 20px; border-radius: 12px; border: 2px solid", badge_color, "; text-align: center; box-shadow: 0 10px 30px rgba(0, 0, 0, 0.3);'>",
                "<div style='font-size: 2.5em; margin-bottom: 10px;'>",
                case_when(rank == 1 ~ "🥇", rank == 2 ~ "🥈", rank == 3 ~ "🥉", TRUE ~ "⭐"), "</div>",
                "<h5 style='color: #00d4ff; font-family: Orbitron, monospace; font-weight: 900; text-transform: uppercase;'>", substr(strain, 1, 15), "</h5>",
                "<div style='background: rgba(0, 0, 0, 0.3); padding: 10px; border-radius: 8px; margin: 10px 0;'>",
                "<p style='color: #ffffff; font-size: 1.4em; font-weight: bold; margin: 0;'>", round(overall, 1), "</p>",
                "<p style='color: #cccccc; font-size: 0.9em; margin: 0;'>Overall Score</p>",
                "</div>",
                "<div style='font-size: 0.85em; color: #cccccc; text-align: left;'>",
                "<div style='display: grid; grid-template-columns: 1fr 1fr 1fr; gap: 5px; margin-top: 8px;'>",
                "<div style='background: rgba(0, 212, 255, 0.2); padding: 5px; border-radius: 4px; text-align: center;'>",
                "<div style='font-weight: bold; color: #00d4ff;'>", cell, "</div>",
                "<div style='font-size: 0.7em;'>Cellulose</div>",
                "</div>",
                "<div style='background: rgba(56, 239, 125, 0.2); padding: 5px; border-radius: 4px; text-align: center;'>",
                "<div style='font-weight: bold; color: #38ef7d;'>", hemi, "</div>",
                "<div style='font-size: 0.7em;'>Hemi</div>",
                "</div>",
                "<div style='background: rgba(255, 107, 53, 0.2); padding: 5px; border-radius: 4px; text-align: center;'>",
                "<div style='font-weight: bold; color: #ff6b35;'>", lignin, "</div>",
                "<div style='font-size: 0.7em;'>Lignin</div>",
                "</div>",
                "</div>",
                "</div>",
                "</div>"
              )
            }, top_3$Rank, top_3$Strain, top_3$Cellulolytic, top_3$HemiCellulolytic, top_3$Lignolytic, top_3$Overall),
            collapse = ""
          ),
          "</div>",
          
          "<div style='background: rgba(56, 239, 125, 0.1); padding: 15px; border-radius: 10px; border-left: 4px solid #38ef7d; margin-top: 20px;'>",
          "<h5 style='color: #38ef7d; font-family: Orbitron, monospace; margin: 0 0 10px 0;'>🎯 INDUSTRIAL RECOMMENDATIONS</h5>",
          "<ul style='color: #cccccc;'>",
          "<li><strong>Best Overall:</strong> ", top_3$Strain[1], " - Complete biorefinery potential</li>",
          "<li><strong>Cellulase Champion:</strong> ", recommendations$Strain[which.max(recommendations$Cellulolytic)], " - Superior cellulose processing</li>",
          "<li><strong>Hemicellulose Expert:</strong> ", recommendations$Strain[which.max(recommendations$HemiCellulolytic)], " - Advanced hemicellulose degradation</li>",
          "</ul>",
          "</div>",
          "</div>"
        )
        
        HTML(recommendation_html)
      } else {
        HTML(paste(
          "<div class='metric-card'>",
          "<h4>🎯 BIOREFINERY RECOMMENDATIONS</h4>",
          "<p style='color: #cccccc; text-align: center; font-size: 1.1em; margin: 40px 20px;'>",
          "Process genomic data to generate biorefinery strain recommendations",
          "</p>",
          "</div>"
        ))
      }
    } else {
      HTML(paste(
        "<div class='metric-card'>",
        "<h4>🎯 BIOREFINERY RECOMMENDATIONS</h4>",
        "<p style='color: #cccccc; text-align: center; font-size: 1.1em; margin: 40px 20px;'>",
        "Upload and process data to discover optimal bacterial strains for your biorefinery application",
        "</p>",
        "</div>"
      ))
    }
  })
  
  # Update the biorefinery matrix plot to use optimization results
  output$biorefineryMatrix <- renderPlotly({
    if (!is.null(values$optimization_results)) {
      
      # Get top 15 strains from optimization
      matrix_data <- values$optimization_results %>%
        slice_head(n = 15) %>%
        select(Strain, PrimaryScore, SecondaryScore, DiversityScore) %>%
        pivot_longer(cols = c("PrimaryScore", "SecondaryScore", "DiversityScore"),
                     names_to = "ScoreType", values_to = "Score") %>%
        mutate(
          ScoreType = case_when(
            ScoreType == "PrimaryScore" ~ "Primary Enzymes",
            ScoreType == "SecondaryScore" ~ "Secondary Enzymes", 
            ScoreType == "DiversityScore" ~ "Enzyme Diversity",
            TRUE ~ ScoreType
          )
        )
      
      p <- ggplot(matrix_data, aes(x = ScoreType, y = reorder(Strain, Score), fill = Score)) +
        geom_tile(color = "white", size = 0.5) +
        labs(title = paste("Biorefinery Optimization Matrix -", input$biorefineryTarget),
             x = "Score Component", y = "Optimized Strains") +
        theme_minimal() +
        theme(
          plot.background = element_rect(fill = "transparent", color = NA),
          panel.background = element_rect(fill = "transparent", color = NA),
          text = element_text(color = "white"),
          axis.text = element_text(color = "white"),
          axis.text.x = element_text(angle = 45, hjust = 1),
          legend.text = element_text(color = "white"),
          legend.title = element_text(color = "white")
        ) +
        scale_fill_viridis_c(option = "plasma", name = "Score")
      
      ggplotly(p) %>%
        layout(
          plot_bgcolor = 'rgba(0,0,0,0)',
          paper_bgcolor = 'rgba(0,0,0,0)',
          font = list(color = 'white')
        )
      
    } else {
      # Default placeholder
      plot_ly(
        x = c("Primary", "Secondary", "Diversity"),
        y = c("Strain A", "Strain B", "Strain C"),
        z = matrix(c(10, 15, 8, 20, 12, 18, 5, 8, 12), nrow = 3),
        type = "heatmap",
        colorscale = "Viridis",
        hovertemplate = "Component: %{x}<br>Strain: %{y}<br>Score: %{z}<extra></extra>"
      ) %>%
        layout(
          title = "Configure Optimization Parameters",
          plot_bgcolor = 'rgba(0,0,0,0)',
          paper_bgcolor = 'rgba(0,0,0,0)',
          font = list(color = 'white')
        )
    }
  })
  
  # Update economic assessment to use optimization results
  output$economicAssessment <- renderUI({
    if (!is.null(values$optimization_results)) {
      
      # Get best performing strain
      best_strain <- values$optimization_results %>%
        filter(MeetsRequirements) %>%
        slice_head(n = 1)
      
      if(nrow(best_strain) > 0) {
        # Calculate economic metrics
        economic_viability <- case_when(
          best_strain$TotalScore >= 100 ~ "EXCELLENT",
          best_strain$TotalScore >= 50 ~ "GOOD",
          best_strain$TotalScore >= 25 ~ "MODERATE", 
          TRUE ~ "POOR"
        )
        
        investment_level <- case_when(
          best_strain$TotalScore >= 100 ~ "HIGH PRIORITY",
          best_strain$TotalScore >= 50 ~ "RECOMMENDED",
          best_strain$TotalScore >= 25 ~ "CONSIDER",
          TRUE ~ "LOW PRIORITY"
        )
        
        roi_estimate <- case_when(
          best_strain$TotalScore >= 100 ~ "20-35%",
          best_strain$TotalScore >= 50 ~ "12-25%",
          best_strain$TotalScore >= 25 ~ "8-18%",
          TRUE ~ "3-12%"
        )
        
        market_readiness <- case_when(
          best_strain$TotalScore >= 75 ~ "READY",
          best_strain$TotalScore >= 40 ~ "DEVELOPMENT",
          best_strain$TotalScore >= 20 ~ "RESEARCH",
          TRUE ~ "EARLY STAGE"
        )
        
        HTML(paste(
          "<div class='metric-card'>",
          "<h5 style='color: #00d4ff; font-family: Orbitron, monospace;'>💰 ECONOMIC ASSESSMENT</h5>",
          
          "<div style='margin: 15px 0; padding: 15px; background: rgba(0, 212, 255, 0.1); border-radius: 10px; border-left: 4px solid #00d4ff;'>",
          "<p><strong>Optimal Strain:</strong> ", substr(best_strain$Strain, 1, 20), "</p>",
          "<p><strong>Optimization Score:</strong> ", round(best_strain$TotalScore, 1), "</p>",
          "<p><strong>Target:</strong> ", input$biorefineryTarget, "</p>",
          "</div>",
          
          "<div style='margin: 15px 0; padding: 15px; background: rgba(56, 239, 125, 0.1); border-radius: 10px; border-left: 4px solid #38ef7d;'>",
          "<p><strong>Economic Viability:</strong> ", economic_viability, "</p>",
          "<p><strong>Investment Priority:</strong> ", investment_level, "</p>",
          "<p><strong>Market Readiness:</strong> ", market_readiness, "</p>",
          "</div>",
          
          "<div style='margin: 15px 0; padding: 15px; background: rgba(255, 107, 53, 0.1); border-radius: 10px; border-left: 4px solid #ff6b35;'>",
          "<p><strong>ROI Estimate:</strong> ", roi_estimate, "</p>",
          "<p><strong>Risk Level:</strong> ", if(best_strain$TotalScore >= 50) "LOW" else "MODERATE", "</p>",
          "<p><strong>Scale-up Potential:</strong> ", if(best_strain$TotalScore >= 60) "HIGH" else "MEDIUM", "</p>",
          "</div>",
          
          "<div style='margin: 15px 0; padding: 10px; background: rgba(156, 89, 209, 0.1); border-radius: 8px; text-align: center;'>",
          "<p style='color: #9c59d1; font-weight: bold; margin: 0;'>OPTIMIZATION CONFIDENCE</p>",
          "<p style='color: #ffffff; font-size: 1.3em; font-weight: bold; margin: 5px 0;'>", 
          round((best_strain$TotalScore / max(values$optimization_results$TotalScore, na.rm = TRUE)) * 100, 0), "%</p>",
          "</div>",
          
          "</div>"
        ))
      } else {
        HTML(paste(
          "<div class='metric-card'>",
          "<h5 style='color: #ff4444;'>⚠️ INSUFFICIENT DATA</h5>",
          "<p>No strains meet the optimization criteria for economic assessment.</p>",
          "</div>"
        ))
      }
    } else {
      HTML(paste(
        "<div class='metric-card'>",
        "<h5 style='color: #00d4ff; font-family: Orbitron, monospace;'>💰 ECONOMIC ASSESSMENT</h5>",
        "<div style='text-align: center; padding: 40px 20px; color: #cccccc;'>",
        "<div style='font-size: 3em; margin-bottom: 20px;'>📊</div>",
        "<p>Run optimization analysis to generate economic assessment</p>",
        "</div>",
        "</div>"
      ))
    }
  })
  
  output$biorefineryRecommendations <- renderUI({
    if (!is.null(values$processed_data)) {
      genome_cols <- names(values$processed_data)[!names(values$processed_data) %in% c("Annotations/Genomes", "Total")]
      
      if(length(genome_cols) > 0) {
        # Calculate comprehensive scores for all strains
        recommendations <- data.frame(
          Strain = genome_cols,
          Cellulolytic = 0,
          HemiCellulolytic = 0,
          Lignolytic = 0,
          Overall = 0,
          Rank = 0,
          stringsAsFactors = FALSE
        )
        
        # Calculate scores
        for(i in 1:nrow(recommendations)) {
          strain <- recommendations$Strain[i]
          
          cell_pot <- calculateDegradationPotential(values$processed_data, values$enzyme_families$cellulolytic)
          hemi_pot <- calculateDegradationPotential(values$processed_data, values$enzyme_families$hemi_cellulolytic)
          lignin_pot <- calculateDegradationPotential(values$processed_data, values$enzyme_families$lignolytic)
          
          recommendations$Cellulolytic[i] <- if(!is.null(cell_pot) && strain %in% names(cell_pot)) cell_pot[[strain]] else 0
          recommendations$HemiCellulolytic[i] <- if(!is.null(hemi_pot) && strain %in% names(hemi_pot)) hemi_pot[[strain]] else 0
          recommendations$Lignolytic[i] <- if(!is.null(lignin_pot) && strain %in% names(lignin_pot)) lignin_pot[[strain]] else 0
          
          # Calculate weighted overall score
          recommendations$Overall[i] <- recommendations$Cellulolytic[i] * 0.4 + 
            recommendations$HemiCellulolytic[i] * 0.4 + 
            recommendations$Lignolytic[i] * 0.2
        }
        
        # Rank strains
        recommendations <- recommendations %>%
          arrange(desc(Overall)) %>%
          mutate(Rank = row_number()) %>%
          slice_head(n = 10)
        
        # Generate HTML recommendations
        top_3 <- recommendations %>% slice_head(n = 3)
        
        recommendation_html <- paste(
          "<div style='display: grid; grid-template-columns: 1fr 1fr 1fr; gap: 20px;'>",
          paste(
            mapply(function(rank, strain, cell, hemi, lignin, overall) {
              badge_color <- case_when(
                rank == 1 ~ "#FFD700",
                rank == 2 ~ "#C0C0C0", 
                rank == 3 ~ "#CD7F32",
                TRUE ~ "#666666"
              )
              
              paste(
                "<div style='background: rgba(0, 212, 255, 0.1); padding: 20px; border-radius: 12px; border: 2px solid", badge_color, "; text-align: center;'>",
                "<div style='font-size: 2em; color:", badge_color, "; margin-bottom: 10px;'>",
                case_when(rank == 1 ~ "🥇", rank == 2 ~ "🥈", rank == 3 ~ "🥉", TRUE ~ "⭐"), "</div>",
                "<h5 style='color: #00d4ff; font-family: Orbitron, monospace;'>", substr(strain, 1, 12), "</h5>",
                "<p style='color: #ffffff; font-size: 1.2em; font-weight: bold;'>Score: ", round(overall, 1), "</p>",
                "<div style='font-size: 0.9em; color: #cccccc;'>",
                "🌾 Cell: ", cell, " | 🌿 Hemi: ", hemi, " | 🌳 Lignin: ", lignin,
                "</div>",
                "</div>"
              )
            }, top_3$Rank, top_3$Strain, top_3$Cellulolytic, top_3$HemiCellulolytic, top_3$Lignolytic, top_3$Overall),
            collapse = ""
          ),
          "</div>",
          
          "<div style='margin-top: 25px; background: rgba(56, 239, 125, 0.1); padding: 15px; border-radius: 10px; border-left: 4px solid #38ef7d;'>",
          "<h5 style='color: #38ef7d; font-family: Orbitron, monospace;'>🎯 INDUSTRIAL RECOMMENDATIONS:</h5>",
          "<ul style='color: #cccccc;'>",
          "<li><strong>Best Overall:</strong> ", top_3$Strain[1], " - Complete biorefinery potential</li>",
          "<li><strong>Cellulase Champion:</strong> ", recommendations$Strain[which.max(recommendations$Cellulolytic)], " - Superior cellulose processing</li>",
          "<li><strong>Hemicellulose Expert:</strong> ", recommendations$Strain[which.max(recommendations$HemiCellulolytic)], " - Advanced hemicellulose degradation</li>",
          "</ul>",
          "</div>"
        )
        
        HTML(recommendation_html)
      }
    }
  })
  
  # Biorefinery performance matrix
  output$biorefineryMatrix <- renderPlotly({
    if (!is.null(values$processed_data)) {
      genome_cols <- names(values$processed_data)[!names(values$processed_data) %in% c("Annotations/Genomes", "Total")]
      
      if(length(genome_cols) > 0) {
        # Create performance matrix
        matrix_data <- data.frame(
          Strain = genome_cols,
          Cellulolytic = 0,
          HemiCellulolytic = 0,
          Lignolytic = 0,
          stringsAsFactors = FALSE
        )
        
        # Calculate scores for matrix
        for(i in 1:nrow(matrix_data)) {
          strain <- matrix_data$Strain[i]
          
          cell_pot <- calculateDegradationPotential(values$processed_data, values$enzyme_families$cellulolytic)
          hemi_pot <- calculateDegradationPotential(values$processed_data, values$enzyme_families$hemi_cellulolytic)
          lignin_pot <- calculateDegradationPotential(values$processed_data, values$enzyme_families$lignolytic)
          
          matrix_data$Cellulolytic[i] <- if(!is.null(cell_pot) && strain %in% names(cell_pot)) cell_pot[[strain]] else 0
          matrix_data$HemiCellulolytic[i] <- if(!is.null(hemi_pot) && strain %in% names(hemi_pot)) hemi_pot[[strain]] else 0
          matrix_data$Lignolytic[i] <- if(!is.null(lignin_pot) && strain %in% names(lignin_pot)) lignin_pot[[strain]] else 0
        }
        
        # Get top 15 strains for visualization
        matrix_data <- matrix_data %>%
          mutate(Overall = Cellulolytic + HemiCellulolytic + Lignolytic) %>%
          slice_max(Overall, n = 15) %>%
          select(-Overall)
        
        # Reshape for plotting
        plot_data <- matrix_data %>%
          pivot_longer(cols = c("Cellulolytic", "HemiCellulolytic", "Lignolytic"), 
                       names_to = "Capability", values_to = "Score")
        
        p <- ggplot(plot_data, aes(x = Capability, y = reorder(Strain, Score), fill = Score)) +
          geom_tile(color = "white", size = 0.5) +
          labs(title = "Biorefinery Performance Matrix",
               x = "Degradation Capability", y = "Bacterial Strain") +
          theme_minimal() +
          theme(
            plot.background = element_rect(fill = "transparent", color = NA),
            panel.background = element_rect(fill = "transparent", color = NA),
            text = element_text(color = "white"),
            axis.text = element_text(color = "white"),
            axis.text.x = element_text(angle = 45, hjust = 1),
            legend.text = element_text(color = "white"),
            legend.title = element_text(color = "white")
          ) +
          scale_fill_viridis_c(option = "plasma", name = "Score")
        
        ggplotly(p) %>%
          layout(
            plot_bgcolor = 'rgba(0,0,0,0)',
            paper_bgcolor = 'rgba(0,0,0,0)',
            font = list(color = 'white')
          )%>%
          config(
            toImageButtonOptions = list(
              format = 'png',
              filename = 'Biorefinery Performance Matrix',
              height = 450,
              width = 500,
              scale = 4  
            )
          )
      }
    }
  })
  
  # Economic viability assessment
  output$economicAssessment <- renderUI({
    if (!is.null(values$processed_data)) {
      # Calculate economic metrics (simplified example)
      genome_cols <- names(values$processed_data)[!names(values$processed_data) %in% c("Annotations/Genomes", "Total")]
      
      if(length(genome_cols) > 0) {
        # Get top performing strain
        cell_pot <- calculateDegradationPotential(values$processed_data, values$enzyme_families$cellulolytic)
        hemi_pot <- calculateDegradationPotential(values$processed_data, values$enzyme_families$hemi_cellulolytic)
        
        if(!is.null(cell_pot) && !is.null(hemi_pot)) {
          combined_scores <- Map("+", cell_pot, hemi_pot)
          top_strain <- names(combined_scores)[which.max(combined_scores)]
          top_score <- max(unlist(combined_scores))
          
          # Economic assessment metrics
          economic_viability <- case_when(
            top_score >= 100 ~ "EXCELLENT",
            top_score >= 50 ~ "GOOD", 
            top_score >= 20 ~ "MODERATE",
            TRUE ~ "POOR"
          )
          
          investment_level <- case_when(
            top_score >= 100 ~ "HIGH PRIORITY",
            top_score >= 50 ~ "RECOMMENDED",
            top_score >= 20 ~ "CONSIDER",
            TRUE ~ "LOW PRIORITY"
          )
          
          roi_estimate <- case_when(
            top_score >= 100 ~ "15-25%",
            top_score >= 50 ~ "10-18%",
            top_score >= 20 ~ "5-12%",
            TRUE ~ "2-8%"
          )
          
          HTML(paste(
            "<div class='metric-card'>",
            "<h5 style='color: #00d4ff; font-family: Orbitron, monospace;'>💰 ECONOMIC VIABILITY</h5>",
            
            "<div style='margin: 15px 0; padding: 12px; background: rgba(0, 212, 255, 0.1); border-radius: 8px; border-left: 3px solid #00d4ff;'>",
            "<p><strong>Top Strain:</strong> ", substr(top_strain, 1, 15), "</p>",
            "<p><strong>Performance Score:</strong> ", round(top_score, 1), "</p>",
            "</div>",
            
            "<div style='margin: 15px 0; padding: 12px; background: rgba(56, 239, 125, 0.1); border-radius: 8px; border-left: 3px solid #38ef7d;'>",
            "<p><strong>Economic Rating:</strong> ", economic_viability, "</p>",
            "<p><strong>Investment Level:</strong> ", investment_level, "</p>",
            "</div>",
            
            "<div style='margin: 15px 0; padding: 12px; background: rgba(255, 107, 53, 0.1); border-radius: 8px; border-left: 3px solid #ff6b35;'>",
            "<p><strong>Est. ROI Range:</strong> ", roi_estimate, "</p>",
            "<p><strong>Market Readiness:</strong> ", if(top_score >= 75) "READY" else if(top_score >= 40) "DEVELOPMENT" else "RESEARCH", "</p>",
            "</div>",
            
            "</div>"
          ))
        }
      }
    }
  })
  
  # Advanced analytics outputs - REAL IMPLEMENTATION
  output$correlationMatrix <- renderPlotly({
    if (!is.null(values$processed_data)) {
      tryCatch({
        # Extract numeric columns (genome data)
        enzyme_col <- if("Annotations/Genomes" %in% names(values$processed_data)) {
          "Annotations/Genomes"
        } else {
          names(values$processed_data)[1]
        }
        
        # Get numeric genome columns only
        numeric_data <- values$processed_data %>%
          select_if(is.numeric) %>%
          select(-any_of(c("Total", "EnzymeClass", "IndustrialRelevance")))
        
        if(ncol(numeric_data) < 2) {
          # Fallback: create basic plot if insufficient data
          p <- plot_ly(
            x = c(1), y = c(1), z = matrix(c(1)),
            type = "heatmap",
            colorscale = list(c(0, "#440154"), c(0.5, "#21908c"), c(1, "#fde725"))
          ) %>%
            layout(
              title = "Correlation Matrix - Insufficient Data",
              plot_bgcolor = 'rgba(0,0,0,0)',
              paper_bgcolor = 'rgba(0,0,0,0)',
              font = list(color = 'white')
            )
          return(p)
        }
        
        # Calculate correlation matrix
        cor_matrix <- cor(numeric_data, use = "complete.obs", method = "pearson")
        
        # Create labels for hover text
        n_genomes <- ncol(cor_matrix)
        hover_text <- matrix("", nrow = n_genomes, ncol = n_genomes)
        for(i in 1:n_genomes) {
          for(j in 1:n_genomes) {
            hover_text[i,j] <- paste(
              "Genome 1:", colnames(cor_matrix)[i], "<br>",
              "Genome 2:", colnames(cor_matrix)[j], "<br>",
              "Correlation:", round(cor_matrix[i,j], 3), "<br>",
              "Strength:", case_when(
                abs(cor_matrix[i,j]) >= 0.8 ~ "Very Strong",
                abs(cor_matrix[i,j]) >= 0.6 ~ "Strong", 
                abs(cor_matrix[i,j]) >= 0.4 ~ "Moderate",
                abs(cor_matrix[i,j]) >= 0.2 ~ "Weak",
                TRUE ~ "Very Weak"
              )
            )
          }
        }
        
        # Create correlation heatmap
        p <- plot_ly(
          x = colnames(cor_matrix),
          y = colnames(cor_matrix),
          z = cor_matrix,
          type = "heatmap",
          colorscale = list(
            c(0, "#d73027"),    # Strong negative correlation - red
            c(0.25, "#f46d43"), # Moderate negative - orange
            c(0.5, "#fee08b"),  # No correlation - yellow
            c(0.75, "#a6d96a"), # Moderate positive - light green
            c(1, "#1a9850")     # Strong positive - dark green
          ),
          zmid = 0,
          zmin = -1,
          zmax = 1,
          text = hover_text,
          hovertemplate = "%{text}<extra></extra>",
          showscale = TRUE,
          colorbar = list(
            title = "Correlation<br>Coefficient",
            titlefont = list(color = 'white', size = 12),
            tickfont = list(color = 'white'),
            bgcolor = 'rgba(0,0,0,0)',
            bordercolor = 'white'
          )
        ) %>%
          layout(
            title = list(
              text = paste("Enzyme Profile Correlation Matrix (", n_genomes, " Genomes)"),
              font = list(color = '#00d4ff', size = 16, family = 'Orbitron')
            ),
            xaxis = list(
              title = "Genome Strains",
              tickangle = -45,
              color = 'white',
              showgrid = FALSE
            ),
            yaxis = list(
              title = "Genome Strains", 
              color = 'white',
              showgrid = FALSE
            ),
            plot_bgcolor = 'rgba(0,0,0,0)',
            paper_bgcolor = 'rgba(0,0,0,0)',
            font = list(color = 'white', family = 'Orbitron'),
            margin = list(l = 100, r = 50, b = 100, t = 80)
          )%>%
          config(
            toImageButtonOptions = list(
              format = 'png',
              filename = 'Enzyme_Profile_Correlation_Matrix',
              height = 400,
              width = 400,
              scale = 4  
            )
          )
        
        return(p)
        
      }, error = function(e) {
        # Error fallback
        p <- plot_ly(
          x = c(1, 2), y = c(1, 2), 
          type = "scatter", mode = "markers",
          marker = list(color = '#ff6b35', size = 10)
        ) %>%
          layout(
            title = paste("Correlation Analysis Error:", e$message),
            plot_bgcolor = 'rgba(0,0,0,0)',
            paper_bgcolor = 'rgba(0,0,0,0)',
            font = list(color = 'white')
          )
        return(p)
      })
    } else {
      # No data fallback
      p <- plot_ly(
        x = c(1, 2, 3), y = c(1, 2, 3),
        type = "scatter", mode = "markers",
        marker = list(color = '#999999', size = 8)
      ) %>%
        layout(
          title = "No Data Available for Correlation Analysis",
          plot_bgcolor = 'rgba(0,0,0,0)',
          paper_bgcolor = 'rgba(0,0,0,0)',
          font = list(color = 'white')
        )
      return(p)
    }
  })
  
  output$pcaAnalysis <- renderPlotly({
    if (!is.null(values$processed_data)) {
      tryCatch({
        # Extract numeric columns for PCA
        numeric_data <- values$processed_data %>%
          select_if(is.numeric) %>%
          select(-any_of(c("Total", "EnzymeClass", "IndustrialRelevance"))) %>%
          t() # Transpose so genomes are rows, enzymes are columns
        
        if(nrow(numeric_data) < 3 || ncol(numeric_data) < 2) {
          # Insufficient data for PCA
          p <- plot_ly(
            x = c(1, 2), y = c(1, 2),
            type = "scatter", mode = "markers",
            marker = list(color = '#ff6b35', size = 10)
          ) %>%
            layout(
              title = "PCA Analysis - Insufficient Data (Need ≥3 genomes, ≥2 enzymes)",
              plot_bgcolor = 'rgba(0,0,0,0)',
              paper_bgcolor = 'rgba(0,0,0,0)',
              font = list(color = 'white')
            )
          return(p)
        }
        
        # Remove columns with zero variance
        numeric_data_clean <- numeric_data[, apply(numeric_data, 2, var, na.rm = TRUE) > 0]
        
        # Perform PCA
        pca_result <- prcomp(numeric_data_clean, scale. = TRUE, center = TRUE)
        
        # Extract PC scores
        pc_scores <- as.data.frame(pca_result$x)
        pc_scores$Genome <- rownames(pc_scores)
        
        # Calculate variance explained
        var_explained <- round(100 * pca_result$sdev^2 / sum(pca_result$sdev^2), 1)
        
        # Color genomes by their total enzyme count
        total_enzymes <- rowSums(numeric_data_clean)
        pc_scores$TotalEnzymes <- total_enzymes
        pc_scores$EnzymeLevel <- cut(total_enzymes, 
                                     breaks = quantile(total_enzymes, probs = c(0, 0.33, 0.66, 1), na.rm = TRUE),
                                     labels = c("Low", "Medium", "High"),
                                     include.lowest = TRUE)
        
        # Create 2D PCA plot
        p <- plot_ly(
          data = pc_scores,
          x = ~PC1, y = ~PC2,
          color = ~EnzymeLevel,
          colors = c("#ff6b35", "#ffaa00", "#00ff88"),
          type = "scatter",
          mode = "markers+text",
          text = ~Genome,
          textposition = "top center",
          textfont = list(size = 10, color = 'white'),
          marker = list(
            size = 12,
            line = list(width = 2, color = 'white'),
            opacity = 0.8
          ),
          hovertemplate = paste(
            "<b>%{text}</b><br>",
            "PC1: %{x:.2f}<br>",
            "PC2: %{y:.2f}<br>",
            "Total Enzymes: %{customdata}<br>",
            "<extra></extra>"
          ),
          customdata = ~TotalEnzymes
        ) %>%
          layout(
            title = list(
              text = paste0("Principal Component Analysis<br>",
                            "<sub>PC1: ", var_explained[1], "% variance | PC2: ", var_explained[2], "% variance</sub>"),
              font = list(color = '#00d4ff', size = 16, family = 'Orbitron')
            ),
            xaxis = list(
              title = paste0("PC1 (", var_explained[1], "% variance)"),
              color = 'white',
              showgrid = TRUE,
              gridcolor = 'rgba(255,255,255,0.1)',
              zeroline = TRUE,
              zerolinecolor = 'rgba(255,255,255,0.3)'
            ),
            yaxis = list(
              title = paste0("PC2 (", var_explained[2], "% variance)"),
              color = 'white', 
              showgrid = TRUE,
              gridcolor = 'rgba(255,255,255,0.1)',
              zeroline = TRUE,
              zerolinecolor = 'rgba(255,255,255,0.3)'
            ),
            plot_bgcolor = 'rgba(0,0,0,0)',
            paper_bgcolor = 'rgba(0,0,0,0)',
            font = list(color = 'white', family = 'Orbitron'),
            legend = list(
              title = list(text = "Enzyme<br>Abundance", font = list(color = 'white')),
              font = list(color = 'white'),
              bgcolor = 'rgba(0,0,0,0.5)',
              bordercolor = 'white',
              borderwidth = 1
            )
          )%>%
          config(
            toImageButtonOptions = list(
              format = 'png',
              filename = 'Principal_Component_Analysis',
              height = 400,
              width = 600,
              scale = 4  
            )
          )
        
        return(p)
        
      }, error = function(e) {
        # Error handling
        p <- plot_ly(
          x = c(1, 2), y = c(1, 2),
          type = "scatter", mode = "markers",
          marker = list(color = '#ff6b35', size = 10)
        ) %>%
          layout(
            title = paste("PCA Analysis Error:", e$message),
            plot_bgcolor = 'rgba(0,0,0,0)',
            paper_bgcolor = 'rgba(0,0,0,0)',
            font = list(color = 'white')
          )
        return(p)
      })
    } else {
      # No data fallback
      p <- plot_ly(
        x = c(1, 2, 3), y = c(1, 2, 3),
        type = "scatter", mode = "markers",
        marker = list(color = '#999999', size = 8)
      ) %>%
        layout(
          title = "No Data Available for PCA Analysis",
          plot_bgcolor = 'rgba(0,0,0,0)',
          paper_bgcolor = 'rgba(0,0,0,0)',
          font = list(color = 'white')
        )
      return(p)
    }
  })
  
  # Enhanced ML predictions with clustering analysis
  output$mlPredictions <- renderUI({
    if (!is.null(values$processed_data)) {
      tryCatch({
        # Extract numeric data for clustering
        numeric_data <- values$processed_data %>%
          select_if(is.numeric) %>%
          select(-any_of(c("Total", "EnzymeClass", "IndustrialRelevance"))) %>%
          t()
        
        if(nrow(numeric_data) >= 3) {
          # Perform k-means clustering
          set.seed(42) # For reproducible results
          optimal_k <- min(4, nrow(numeric_data) - 1) # Choose reasonable k
          
          if(optimal_k >= 2) {
            # Clean data and perform clustering
            numeric_data_clean <- numeric_data[, apply(numeric_data, 2, var, na.rm = TRUE) > 0]
            
            if(ncol(numeric_data_clean) > 0) {
              scaled_data <- scale(numeric_data_clean)
              kmeans_result <- kmeans(scaled_data, centers = optimal_k, nstart = 20)
              
              # Create cluster summary
              cluster_summary <- data.frame(
                Genome = rownames(numeric_data),
                Cluster = paste("Cluster", kmeans_result$cluster),
                TotalEnzymes = rowSums(numeric_data_clean),
                stringsAsFactors = FALSE
              )
              
              # Get cluster characteristics
              cluster_chars <- aggregate(TotalEnzymes ~ Cluster, cluster_summary, 
                                         function(x) c(mean = mean(x), count = length(x)))
              
              cluster_info <- ""
              for(i in 1:nrow(cluster_chars)) {
                cluster_info <- paste0(cluster_info,
                                       "<div style='background: rgba(", 
                                       c("0,212,255", "255,107,53", "56,239,125", "156,89,209")[i], 
                                       ",0.1); padding: 10px; margin: 5px 0; border-radius: 5px; border-left: 3px solid ", 
                                       c("#00d4ff", "#ff6b35", "#38ef7d", "#9c59d1")[i], ";'>",
                                       "<strong style='color: ", c("#00d4ff", "#ff6b35", "#38ef7d", "#9c59d1")[i], ";'>", 
                                       cluster_chars$Cluster[i], "</strong><br>",
                                       "Members: ", cluster_chars$TotalEnzymes[i,2], " genomes<br>",
                                       "Avg Enzymes: ", round(cluster_chars$TotalEnzymes[i,1], 1), "<br>",
                                       "Genomes: ", paste(cluster_summary$Genome[cluster_summary$Cluster == cluster_chars$Cluster[i]], collapse = ", "),
                                       "</div>"
                )
              }
              
              HTML(paste0(
                "<div class='metric-card quantum-glow'>",
                "<h5 style='color: #00d4ff; font-family: Orbitron, monospace;'>🤖 MACHINE LEARNING INSIGHTS</h5>",
                "<div style='margin-top: 15px;'>",
                "<h6 style='color: #ff6b35; font-family: Orbitron, monospace;'>📊 K-MEANS CLUSTERING ANALYSIS</h6>",
                "<p style='color: #cccccc; font-size: 14px; margin-bottom: 15px;'>",
                "Identified <strong>", optimal_k, " distinct enzyme profile clusters</strong> among your genomes:</p>",
                cluster_info,
                "<div style='background: rgba(26,188,156,0.1); padding: 15px; margin-top: 15px; border-radius: 10px; border-left: 4px solid #1abc9c;'>",
                "<h6 style='color: #1abc9c; font-family: Orbitron, monospace;'>🎯 PREDICTIVE INSIGHTS</h6>",
                "<p style='color: #cccccc; font-size: 13px;'>• Genomes in the same cluster share similar enzyme profiles</p>",
                "<p style='color: #cccccc; font-size: 13px;'>• High-enzyme clusters may indicate superior biorefinery potential</p>",
                "<p style='color: #cccccc; font-size: 13px;'>• Cluster diversity suggests multiple metabolic strategies</p>",
                "</div>",
                "</div>",
                "</div>"
              ))
            } else {
              HTML("<div class='metric-card'><h5>🤖 Machine Learning Predictions</h5><p>Insufficient variable enzymes for clustering analysis...</p></div>")
            }
          } else {
            HTML("<div class='metric-card'><h5>🤖 Machine Learning Predictions</h5><p>Need at least 3 genomes for clustering analysis...</p></div>")
          }
        } else {
          HTML("<div class='metric-card'><h5>🤖 Machine Learning Predictions</h5><p>Advanced ML features need more data...</p></div>")
        }
      }, error = function(e) {
        HTML(paste0("<div class='metric-card'><h5>🤖 Machine Learning Predictions</h5><p>Error in ML analysis: ", e$message, "</p></div>"))
      })
    } else {
      HTML("<div class='metric-card'><h5>🤖 Machine Learning Predictions</h5><p>No data available for ML analysis...</p></div>")
    }
  })
  
  # Data tables
  output$dataPreview <- DT::renderDataTable({
    if (!is.null(values$processed_data)) {
      DT::datatable(values$processed_data, 
                    options = list(
                      scrollX = TRUE, 
                      pageLength = 15,
                      dom = 'Bfrtip',
                      buttons = c('copy', 'csv', 'excel'),
                      initComplete = JS(
                        "function(settings, json) {",
                        "$(this.api().table().header()).css({'background-color': '#1a1a2e', 'color': '#fff'});",
                        "}"
                      )
                    ),
                    class = 'cell-border stripe hover',
                    rownames = FALSE) %>%
        formatStyle(columns = 1:ncol(values$processed_data), 
                    backgroundColor = 'rgba(26, 26, 46, 0.9)', 
                    color = 'white')
    }
  })
  
  output$singleStrainTable <- DT::renderDataTable({
    if (!is.null(values$processed_data) && !is.null(input$selectedStrain)) {
      enzyme_col <- if("Annotations/Genomes" %in% names(values$processed_data)) {
        "Annotations/Genomes"
      } else {
        names(values$processed_data)[1]
      }
      
      if(input$selectedStrain %in% names(values$processed_data)) {
        strain_table <- values$processed_data %>%
          select(!!sym(enzyme_col), all_of(input$selectedStrain)) %>%
          filter(!!sym(input$selectedStrain) > 0) %>%
          arrange(desc(!!sym(input$selectedStrain))) %>%
          mutate(
            EnzymeClass = case_when(
              str_detect(!!sym(enzyme_col), "^GH") ~ "Glycoside Hydrolase",
              str_detect(!!sym(enzyme_col), "^AA") ~ "Auxiliary Activity",
              str_detect(!!sym(enzyme_col), "^CBM") ~ "Carbohydrate Binding",
              str_detect(!!sym(enzyme_col), "^PL") ~ "Polysaccharide Lyase",
              str_detect(!!sym(enzyme_col), "^GT") ~ "Glycosyltransferase",
              TRUE ~ "Other"
            ),
            IndustrialRelevance = case_when(
              !!sym(input$selectedStrain) >= 10 ~ "High",
              !!sym(input$selectedStrain) >= 5 ~ "Medium",
              TRUE ~ "Low"
            )
          )
        
        DT::datatable(strain_table, 
                      options = list(
                        pageLength = 20,
                        dom = 'Bfrtip',
                        buttons = c('copy', 'csv'),
                        initComplete = JS(
                          "function(settings, json) {",
                          "$(this.api().table().header()).css({'background-color': '#1a1a2e', 'color': '#fff'});",
                          "}"
                        )
                      ),
                      class = 'cell-border stripe hover',
                      rownames = FALSE,
                      colnames = c('Enzyme Family', 'Count', 'Class', 'Industrial Relevance')) %>%
          formatStyle(columns = 1:4, 
                      backgroundColor = 'rgba(26, 26, 46, 0.9)', 
                      color = 'white') %>%
          formatStyle('IndustrialRelevance',
                      backgroundColor = styleEqual(c('High', 'Medium', 'Low'), 
                                                   c('#00ff88', '#ffaa00', '#ff4444')))
      }
    }
  })
  
  output$cellulasePotentialTable <- DT::renderDataTable({
    if (!is.null(values$processed_data)) {
      cellulase_potential <- calculateDegradationPotential(values$processed_data, values$enzyme_families$cellulolytic)
      
      if(!is.null(cellulase_potential)) {
        potential_df <- data.frame(
          Genome = names(cellulase_potential),
          CellulaseScore = as.numeric(cellulase_potential),
          stringsAsFactors = FALSE
        ) %>%
          arrange(desc(CellulaseScore)) %>%
          mutate(
            Rank = row_number(),
            Rating = case_when(
              CellulaseScore >= 75 ~ "🏆 ELITE",
              CellulaseScore >= 50 ~ "⭐ PREMIUM", 
              CellulaseScore >= 25 ~ "✅ GOOD",
              CellulaseScore >= 10 ~ "📈 MODERATE",
              TRUE ~ "⚠️ BASIC"
            ),
            Efficiency = case_when(
              CellulaseScore >= 75 ~ "Excellent",
              CellulaseScore >= 50 ~ "High",
              CellulaseScore >= 25 ~ "Good",
              CellulaseScore >= 10 ~ "Moderate",
              TRUE ~ "Low"
            )
          ) %>%
          slice_head(n = 20)
        
        DT::datatable(potential_df,
                      options = list(
                        pageLength = 15,
                        dom = 'Bfrtip',
                        buttons = c('copy', 'csv'),
                        order = list(list(2, 'desc')),
                        initComplete = JS(
                          "function(settings, json) {",
                          "$(this.api().table().header()).css({'background-color': '#1a1a2e', 'color': '#fff'});",
                          "}"
                        )
                      ),
                      class = 'cell-border stripe hover',
                      rownames = FALSE,
                      colnames = c('Rank', 'Strain', 'Cellulase Score', 'Rating', 'Efficiency')) %>%
          formatStyle(columns = 1:5, 
                      backgroundColor = 'rgba(26, 26, 46, 0.9)', 
                      color = 'white') %>%
          formatStyle('CellulaseScore',
                      background = styleColorBar(range(potential_df$CellulaseScore), '#00d4ff'),
                      backgroundSize = '100% 90%',
                      backgroundRepeat = 'no-repeat',
                      backgroundPosition = 'center')
      }
    }
  })
  
  # Enhanced cellulase insights with proper error handling
  output$cellulaseInsights <- renderUI({
    if (!is.null(values$processed_data)) {
      tryCatch({
        # Calculate detailed enzyme class statistics
        enzyme_col <- if("Annotations/Genomes" %in% names(values$processed_data)) {
          "Annotations/Genomes"
        } else if("CAZymes" %in% names(values$processed_data)) {
          "CAZymes"
        } else {
          names(values$processed_data)[1]
        }
        
        # Count different enzyme types with fixed patterns
        endoglucanases <- c("GH5", "GH8", "GH9", "GH12", "GH44", "GH45", "GH74")
        exoglucanases <- c("GH6", "GH7", "GH48")
        beta_glucosidases <- c("GH1", "GH3", "GH116")
        
        # FIXED: Safe access to enzyme families with fallbacks
        cbm_modules <- NULL
        aa_enzymes <- NULL
        
        if(!is.null(values$enzyme_families) && !is.null(values$enzyme_families$cellulolytic)) {
          if(is.list(values$enzyme_families$cellulolytic)) {
            # Try to get CBM modules
            if("CBM" %in% names(values$enzyme_families$cellulolytic)) {
              cbm_modules <- values$enzyme_families$cellulolytic$CBM
            }
            # Try to get AA enzymes
            if("AA" %in% names(values$enzyme_families$cellulolytic)) {
              aa_enzymes <- values$enzyme_families$cellulolytic$AA
            }
          } else {
            # If it's a flat list, extract CBM and AA patterns
            all_patterns <- unlist(values$enzyme_families$cellulolytic, recursive = TRUE)
            cbm_modules <- all_patterns[grepl("^CBM", all_patterns)]
            aa_enzymes <- all_patterns[grepl("^AA", all_patterns)]
          }
        }
        
        # Set default values if still NULL
        if(is.null(cbm_modules) || length(cbm_modules) == 0) {
          cbm_modules <- c("CBM1", "CBM3", "CBM6")  # Common cellulose-binding modules
        }
        if(is.null(aa_enzymes) || length(aa_enzymes) == 0) {
          aa_enzymes <- c("AA9", "AA10", "AA11", "AA13", "AA14", "AA15", "AA16")  # LPMOs and related
        }
        
        # Create proper regex patterns
        endo_pattern <- paste0("^(", paste(endoglucanases, collapse = "|"), ")($|_|\\d)")
        exo_pattern <- paste0("^(", paste(exoglucanases, collapse = "|"), ")($|_|\\d)")
        bg_pattern <- paste0("^(", paste(beta_glucosidases, collapse = "|"), ")($|_|\\d)")
        cbm_pattern <- paste0("^(", paste(cbm_modules, collapse = "|"), ")($|_|\\d)")
        aa_pattern <- paste0("^(", paste(aa_enzymes, collapse = "|"), ")($|_|\\d)")
        
        # Count enzyme occurrences with proper pattern matching
        endo_count <- sum(str_detect(values$processed_data[[enzyme_col]], endo_pattern), na.rm = TRUE)
        exo_count <- sum(str_detect(values$processed_data[[enzyme_col]], exo_pattern), na.rm = TRUE)
        bg_count <- sum(str_detect(values$processed_data[[enzyme_col]], bg_pattern), na.rm = TRUE)
        cbm_count <- sum(str_detect(values$processed_data[[enzyme_col]], cbm_pattern), na.rm = TRUE)
        aa_count <- sum(str_detect(values$processed_data[[enzyme_col]], aa_pattern), na.rm = TRUE)
        
        # Calculate synergy index with safety check
        synergy_index <- round((endo_count + exo_count + bg_count) * (1 + (cbm_count + aa_count) * 0.1), 1)
        
        HTML(paste(
          "<div class='degradation-card quantum-glow'>",
          "<h4>🔬 CELLULOLYTIC MECHANISM - INDUSTRIAL QUANTUM INSIGHTS</h4>",
          
          "<div style='display: grid; grid-template-columns: 1fr 1fr; gap: 25px; margin-top: 25px;'>",
          
          # Mechanism overview
          "<div>",
          "<h5 style='color: #00d4ff; font-family: Orbitron, monospace; margin-bottom: 15px;'>⚡ DEGRADATION MECHANISM</h5>",
          
          "<div style='background: rgba(0, 212, 255, 0.1); padding: 15px; border-radius: 10px; margin: 10px 0; border-left: 4px solid #00d4ff;'>",
          "<p><strong style='color: #00d4ff; font-family: Orbitron, monospace;'>ENDOGLUCANASES (", endo_count, " detected):</strong></p>",
          "<p style='color: #cccccc; font-family: Orbitron, sans-serif; font-size: 14px;'>⚡ Attack internal β-1,4-glycosidic bonds in cellulose chains</p>",
          "<p style='color: #cccccc; font-family: Orbitron, sans-serif; font-size: 14px;'>🎯 Create new reducing and non-reducing ends for exoglucanases</p>",
          "<p style='color: #aaaaaa; font-family: Orbitron, sans-serif; font-size: 12px;'>Key families: GH5, GH8, GH9, GH12, GH44, GH45, GH74</p>",
          "</div>",
          
          "<div style='background: rgba(255, 107, 53, 0.1); padding: 15px; border-radius: 10px; margin: 10px 0; border-left: 4px solid #ff6b35;'>",
          "<p><strong style='color: #ff6b35; font-family: Orbitron, monospace;'>EXOGLUCANASES (", exo_count, " detected):</strong></p>",
          "<p style='color: #cccccc; font-family: Orbitron, sans-serif; font-size: 14px;'>⚡ Release cellobiose from cellulose chain ends</p>",
          "<p style='color: #cccccc; font-family: Orbitron, sans-serif; font-size: 14px;'>🎯 Processively cleave from reducing/non-reducing ends</p>",
          "<p style='color: #aaaaaa; font-family: Orbitron, sans-serif; font-size: 12px;'>Key families: GH6, GH7, GH48</p>",
          "</div>",
          
          "<div style='background: rgba(56, 239, 125, 0.1); padding: 15px; border-radius: 10px; margin: 10px 0; border-left: 4px solid #38ef7d;'>",
          "<p><strong style='color: #38ef7d; font-family: Orbitron, monospace;'>β-GLUCOSIDASES (", bg_count, " detected):</strong></p>",
          "<p style='color: #cccccc; font-family: Orbitron, sans-serif; font-size: 14px;'>⚡ Convert cellobiose and cellooligosaccharides to glucose</p>",
          "<p style='color: #cccccc; font-family: Orbitron, sans-serif; font-size: 14px;'>🎯 Final step in complete cellulose saccharification</p>",
          "<p style='color: #aaaaaa; font-family: Orbitron, sans-serif; font-size: 12px;'>Key families: GH1, GH3, GH116</p>",
          "</div>",
          "</div>",
          
          # Auxiliary components
          "<div>",
          "<h5 style='color: #ff6b35; font-family: Orbitron, monospace; margin-bottom: 15px;'>🛠️ AUXILIARY COMPONENTS</h5>",
          
          "<div style='background: rgba(156, 89, 209, 0.1); padding: 15px; border-radius: 10px; margin: 10px 0; border-left: 4px solid #9c59d1;'>",
          "<p><strong style='color: #9c59d1; font-family: Orbitron, monospace;'>CBM MODULES (", cbm_count, " detected):</strong></p>",
          "<p style='color: #cccccc; font-family: Orbitron, sans-serif; font-size: 14px;'>🔗 Carbohydrate-binding modules for substrate targeting</p>",
          "<p style='color: #cccccc; font-family: Orbitron, sans-serif; font-size: 14px;'>📍 Enhance enzyme-substrate proximity and binding</p>",
          "<p style='color: #aaaaaa; font-family: Orbitron, sans-serif; font-size: 12px;'>Binding specificity: Cellulose, crystalline surfaces</p>",
          "</div>",
          
          "<div style='background: rgba(247, 147, 30, 0.1); padding: 15px; border-radius: 10px; margin: 10px 0; border-left: 4px solid #f7931e;'>",
          "<p><strong style='color: #f7931e; font-family: Orbitron, monospace;'>AA OXIDASES (", aa_count, " detected):</strong></p>",
          "<p style='color: #cccccc; font-family: Orbitron, sans-serif; font-size: 14px;'>⚡ Auxiliary activities: LPMOs, cellobiose dehydrogenases</p>",
          "<p style='color: #cccccc; font-family: Orbitron, sans-serif; font-size: 14px;'>🎯 Oxidative cleavage and electron transport</p>",
          "<p style='color: #aaaaaa; font-family: Orbitron, sans-serif; font-size: 12px;'>Enhanced crystalline cellulose accessibility</p>",
          "</div>",
          
          "<div style='background: rgba(26, 188, 156, 0.1); padding: 15px; border-radius: 10px; margin: 10px 0; border-left: 4px solid #1abc9c;'>",
          "<h6 style='color: #1abc9c; font-family: Orbitron, monospace;'>🏭 INDUSTRIAL SYNERGY INDEX</h6>",
          "<p style='color: #ffffff; font-size: 24px; font-weight: bold; text-align: center;'>", synergy_index, "</p>",
          "<p style='color: #cccccc; font-size: 12px; text-align: center;'>Complete cellulolytic system efficiency</p>",
          "</div>",
          "</div>",
          
          "</div>",
          "</div>"
        ))
      }, error = function(e) {
        cat("ERROR in cellulaseInsights:", e$message, "\n")
        HTML(paste(
          "<div class='degradation-card'>",
          "<h4>❌ CELLULOLYTIC INSIGHTS ERROR</h4>",
          "<p style='color: #ff4444;'>Error: ", e$message, "</p>",
          "<p style='color: #cccccc;'>Please check your data structure and enzyme family definitions.</p>",
          "</div>"
        ))
      })
    } else {
      HTML(paste(
        "<div class='degradation-card'>",
        "<h4>🔬 CELLULOLYTIC MECHANISM - INDUSTRIAL QUANTUM INSIGHTS</h4>",
        "<p style='color: #cccccc;'>Please upload data to view cellulolytic mechanism insights.</p>",
        "</div>"
      ))
    }
  })
  
  # Enhanced Single Strain Degradation Pathway Output Function
  output$singleStrainPathway <- renderPlotly({
    req(input$pathwaySelectedStrain)
    
    if (!is.null(values$processed_data)) {
      cat("Available columns:", names(values$processed_data), "\n")
      cat("Selected strain:", input$pathwaySelectedStrain, "\n")
      cat("Strain exists in data:", input$pathwaySelectedStrain %in% names(values$processed_data), "\n")
      
      if(input$pathwaySelectedStrain %in% names(values$processed_data)) {
        
        tryCatch({
          # More robust enzyme column identification
          enzyme_col <- NULL
          possible_enzyme_cols <- c("Annotations/Genomes", "CAZymes", "Enzyme", "Family")
          
          for(col in possible_enzyme_cols) {
            if(col %in% names(values$processed_data)) {
              enzyme_col <- col
              break
            }
          }
          
          # If no standard column found, use first column
          if(is.null(enzyme_col)) {
            enzyme_col <- names(values$processed_data)[1]
          }
          
          cat("Using enzyme column:", enzyme_col, "\n")
          
          # Get enzymes present in the selected strain (count > 0)
          strain_enzymes <- values$processed_data %>%
            filter(!!sym(input$pathwaySelectedStrain) > 0) %>%
            select(all_of(c(enzyme_col, input$pathwaySelectedStrain)))
          
          cat("Found", nrow(strain_enzymes), "enzymes for strain\n")
          
          if(nrow(strain_enzymes) == 0) {
            return(plot_ly() %>%
                     add_text(x = 0.5, y = 0.5, text = paste("No enzymes found for", input$pathwaySelectedStrain), 
                              textfont = list(color = "orange", size = 16)) %>%
                     layout(
                       xaxis = list(showgrid = FALSE, showticklabels = FALSE, zeroline = FALSE),
                       yaxis = list(showgrid = FALSE, showticklabels = FALSE, zeroline = FALSE),
                       plot_bgcolor = 'rgba(0,0,0,0)',
                       paper_bgcolor = 'rgba(0,0,0,0)',
                       font = list(color = 'white')
                     ))
          }
          
          # Define CAZyme family-to-pathway mappings (allowing multiple pathways per family)
          cazyme_pathway_map <- list(
            # Cellulolytic enzymes
            "CBM1" = c("Cellulolytic"), "CBM3" = c("Cellulolytic"), "CBM6" = c("Cellulolytic", "Hemi-Cellulolytic"),
            "CBM8" = c("Cellulolytic"), "CBM9" = c("Cellulolytic", "Hemi-Cellulolytic"), "CBM10" = c("Cellulolytic"),
            "CBM16" = c("Cellulolytic", "Hemi-Cellulolytic"), "CBM17" = c("Cellulolytic"), "CBM28" = c("Cellulolytic"),
            "CBM30" = c("Cellulolytic"), "CBM37" = c("Cellulolytic", "Hemi-Cellulolytic"), "CBM44" = c("Cellulolytic", "Hemi-Cellulolytic"),
            "CBM46" = c("Cellulolytic"), "CBM49" = c("Cellulolytic"), "CBM59" = c("Cellulolytic", "Hemi-Cellulolytic"),
            "CBM63" = c("Cellulolytic"), "CBM64" = c("Cellulolytic"), "CBM72" = c("Cellulolytic", "Hemi-Cellulolytic"),
            
            "GH1" = c("Cellulolytic", "Hemi-Cellulolytic", "Pectinolytic"), "GH2" = c("Cellulolytic", "Hemi-Cellulolytic", "Pectinolytic"),
            "GH3" = c("Cellulolytic", "Hemi-Cellulolytic", "Pectinolytic"), "GH5" = c("Cellulolytic", "Hemi-Cellulolytic"),
            "GH6" = c("Cellulolytic"), "GH7" = c("Cellulolytic"), "GH8" = c("Cellulolytic", "Hemi-Cellulolytic"),
            "GH9" = c("Cellulolytic", "Hemi-Cellulolytic"), "GH10" = c("Cellulolytic", "Hemi-Cellulolytic", "Pectinolytic"),
            "GH12" = c("Cellulolytic", "Hemi-Cellulolytic"), "GH26" = c("Cellulolytic", "Hemi-Cellulolytic"),
            "GH30" = c("Cellulolytic", "Hemi-Cellulolytic"), "GH39" = c("Cellulolytic", "Hemi-Cellulolytic", "Pectinolytic"),
            "GH44" = c("Cellulolytic", "Hemi-Cellulolytic"), "GH45" = c("Cellulolytic"), "GH48" = c("Cellulolytic"),
            "GH51" = c("Cellulolytic", "Hemi-Cellulolytic", "Pectinolytic"), "GH74" = c("Cellulolytic", "Hemi-Cellulolytic"),
            "GH124" = c("Cellulolytic"), "GH116" = c("Cellulolytic", "Hemi-Cellulolytic"),
            
            "AA3" = c("Cellulolytic", "Lignolytic"), "AA9" = c("Cellulolytic", "Hemi-Cellulolytic", "Lignolytic"),
            "AA10" = c("Cellulolytic"), "AA15" = c("Cellulolytic"),
            
            # Hemi-Cellulolytic specific
            "GH11" = c("Hemi-Cellulolytic"), "GH16" = c("Hemi-Cellulolytic"), "GH17" = c("Hemi-Cellulolytic"),
            "GH43" = c("Hemi-Cellulolytic", "Pectinolytic"), "GH52" = c("Hemi-Cellulolytic"), "GH54" = c("Hemi-Cellulolytic", "Pectinolytic"),
            "GH55" = c("Hemi-Cellulolytic"), "GH62" = c("Hemi-Cellulolytic", "Pectinolytic"), "GH98" = c("Hemi-Cellulolytic"),
            "GH113" = c("Hemi-Cellulolytic"), "GH120" = c("Hemi-Cellulolytic"), "GH127" = c("Hemi-Cellulolytic"),
            "GH134" = c("Hemi-Cellulolytic"), "GH137" = c("Hemi-Cellulolytic"), "GH141" = c("Hemi-Cellulolytic"),
            "GH142" = c("Hemi-Cellulolytic"), "GH146" = c("Hemi-Cellulolytic"),
            
            "CBM2" = c("Hemi-Cellulolytic"), "CBM4" = c("Hemi-Cellulolytic"), "CBM13" = c("Hemi-Cellulolytic", "Pectinolytic"),
            "CBM15" = c("Hemi-Cellulolytic"), "CBM22" = c("Hemi-Cellulolytic"), "CBM31" = c("Hemi-Cellulolytic"),
            "CBM35" = c("Hemi-Cellulolytic"), "CBM36" = c("Hemi-Cellulolytic"), "CBM42" = c("Hemi-Cellulolytic"),
            "CBM54" = c("Hemi-Cellulolytic"), "CBM60" = c("Hemi-Cellulolytic"), "CBM23" = c("Hemi-Cellulolytic"),
            "CBM27" = c("Hemi-Cellulolytic"), "CBM29" = c("Hemi-Cellulolytic"), "CBM62" = c("Hemi-Cellulolytic", "Pectinolytic"),
            "CBM76" = c("Hemi-Cellulolytic"), "CBM80" = c("Hemi-Cellulolytic", "Pectinolytic"), "CBM65" = c("Hemi-Cellulolytic"),
            "CBM75" = c("Hemi-Cellulolytic"), "CBM78" = c("Hemi-Cellulolytic"), "CBM81" = c("Hemi-Cellulolytic"),
            
            "CE1" = c("Hemi-Cellulolytic", "Lignolytic"), "CE2" = c("Hemi-Cellulolytic"), "CE3" = c("Hemi-Cellulolytic"),
            "CE4" = c("Hemi-Cellulolytic"), "CE5" = c("Hemi-Cellulolytic"), "CE6" = c("Hemi-Cellulolytic", "Pectinolytic"),
            "CE7" = c("Hemi-Cellulolytic"), "CE12" = c("Hemi-Cellulolytic", "Pectinolytic"), "CE15" = c("Hemi-Cellulolytic", "Lignolytic"),
            
            "AA14" = c("Hemi-Cellulolytic"),
            
            # Lignolytic enzymes
            "AA1" = c("Lignolytic"), "AA2" = c("Lignolytic"), "AA4" = c("Lignolytic"), "AA5" = c("Lignolytic"),
            "AA6" = c("Lignolytic"), "AA8" = c("Lignolytic"), "AA12" = c("Lignolytic"),
            "CE10" = c("Lignolytic"),
            
            # Pectinolytic enzymes
            "GH4" = c("Pectinolytic", "Starch Degrading"), "GH28" = c("Pectinolytic"), "GH33" = c("Pectinolytic"),
            "GH35" = c("Pectinolytic"), "GH42" = c("Pectinolytic"), "GH50" = c("Pectinolytic"),
            "GH59" = c("Pectinolytic"), "GH78" = c("Pectinolytic"), "GH93" = c("Pectinolytic"),
            "GH106" = c("Pectinolytic"), "GH147" = c("Pectinolytic"),
            
            "PL1" = c("Pectinolytic"), "PL2" = c("Pectinolytic"), "PL3" = c("Pectinolytic"), "PL4" = c("Pectinolytic"),
            "PL9" = c("Pectinolytic"), "PL10" = c("Pectinolytic"), "PL11" = c("Pectinolytic"), "PL22" = c("Pectinolytic"),
            "PL26" = c("Pectinolytic"),
            
            "CE8" = c("Pectinolytic"), "CE13" = c("Pectinolytic"),
            "CBM32" = c("Pectinolytic"), "CBM41" = c("Pectinolytic"), "CBM51" = c("Pectinolytic"),
            "CBM61" = c("Pectinolytic"), "CBM67" = c("Pectinolytic"), "CBM77" = c("Pectinolytic"),
            
            # Starch degrading enzymes
            "GH13" = c("Starch Degrading"), "GH14" = c("Starch Degrading"), "GH57" = c("Starch Degrading"),
            "GH119" = c("Starch Degrading"), "GH126" = c("Starch Degrading"), "GH31" = c("Starch Degrading"),
            "GH63" = c("Starch Degrading"), "GH97" = c("Starch Degrading"), "GH122" = c("Starch Degrading"),
            "GT35" = c("Starch Degrading"),
            
            "CBM20" = c("Starch Degrading"), "CBM21" = c("Starch Degrading"), "CBM25" = c("Starch Degrading"),
            "CBM26" = c("Starch Degrading"), "CBM34" = c("Starch Degrading"), "CBM45" = c("Starch Degrading"),
            "CBM53" = c("Starch Degrading"), "CBM69" = c("Starch Degrading"), "CBM74" = c("Starch Degrading"),
            "CBM82" = c("Starch Degrading"), "CBM83" = c("Starch Degrading"),
            
            "AA13" = c("Starch Degrading"),
            
            # Inulin degrading enzymes
            "GH32" = c("Inulin Degrading"), "GH91" = c("Inulin Degrading"),
            "CBM38" = c("Inulin Degrading")
          )
          
          # Pathway color scheme
          pathway_colors <- list(
            "Cellulolytic" = "#3b82f6",
            "Hemi-Cellulolytic" = "#10b981", 
            "Lignolytic" = "#ef4444",
            "Pectinolytic" = "#8b5cf6",
            "Starch Degrading" = "#f59e0b",
            "Inulin Degrading" = "#06b6d4"
          )
          
          # Initialize data structures
          all_nodes <- data.frame()
          all_edges <- data.frame()
          pathway_nodes <- data.frame()
          
          # Define different layout themes
          layout_theme <- "circular"  # Default theme - could be made interactive later
          
          # Create pathway positions based on theme
          if(layout_theme == "circular") {
            # Circular arrangement
            pathway_positions <- list(
              "Cellulolytic" = list(x = 9 + 5*cos(0), y = 6 + 4*sin(0)),
              "Hemi-Cellulolytic" = list(x = 9 + 5*cos(pi/3), y = 6 + 4*sin(pi/3)),
              "Lignolytic" = list(x = 9 + 5*cos(2*pi/3), y = 6 + 4*sin(2*pi/3)),
              "Pectinolytic" = list(x = 9 + 5*cos(pi), y = 6 + 4*sin(pi)),
              "Starch Degrading" = list(x = 9 + 5*cos(4*pi/3), y = 6 + 4*sin(4*pi/3)),
              "Inulin Degrading" = list(x = 9 + 5*cos(5*pi/3), y = 6 + 4*sin(5*pi/3))
            )
          } else if(layout_theme == "hierarchical") {
            # Hierarchical branching layout
            pathway_positions <- list(
              "Cellulolytic" = list(x = 9, y = 10),
              "Hemi-Cellulolytic" = list(x = 6, y = 8),
              "Lignolytic" = list(x = 12, y = 8),
              "Pectinolytic" = list(x = 3, y = 6),
              "Starch Degrading" = list(x = 9, y = 6),
              "Inulin Degrading" = list(x = 15, y = 6)
            )
          } else if(layout_theme == "radial") {
            # Radial spoke pattern
            pathway_positions <- list(
              "Cellulolytic" = list(x = 9, y = 11),
              "Hemi-Cellulolytic" = list(x = 14, y = 9),
              "Lignolytic" = list(x = 14, y = 3),
              "Pectinolytic" = list(x = 9, y = 1),
              "Starch Degrading" = list(x = 4, y = 3),
              "Inulin Degrading" = list(x = 4, y = 9)
            )
          }
          
          # Find which pathways are active (have enzymes present)
          active_pathways <- list()
          pathway_enzyme_counts <- list()
          
          # Debug: Print found enzymes and their families
          cat("Processing", nrow(strain_enzymes), "enzymes for strain", input$pathwaySelectedStrain, "\n")
          
          # Process each enzyme in the strain
          for(i in 1:nrow(strain_enzymes)) {
            enzyme_name <- strain_enzymes[[enzyme_col]][i]
            enzyme_count <- strain_enzymes[[input$pathwaySelectedStrain]][i]
            
            # Extract CAZyme family (e.g., GH5, AA9, etc.) - improved extraction
            enzyme_family <- enzyme_name
            
            # Try different patterns to extract family
            if(grepl("^[A-Z]+[0-9]+", enzyme_name)) {
              enzyme_family <- gsub("^([A-Z]+[0-9]+).*", "\\1", enzyme_name)
            } else if(grepl("^[A-Z]+_[0-9]+", enzyme_name)) {
              enzyme_family <- gsub("^([A-Z]+)_([0-9]+).*", "\\1\\2", enzyme_name)
            } else {
              # Try to extract from beginning of string
              enzyme_family <- gsub("([A-Z]+[0-9]+).*", "\\1", enzyme_name)
            }
            
            cat("Processing enzyme:", enzyme_name, "-> Family:", enzyme_family, "\n")
            
            # Check if this enzyme family maps to any pathways
            if(enzyme_family %in% names(cazyme_pathway_map)) {
              pathways <- cazyme_pathway_map[[enzyme_family]]
              cat("Found pathways for", enzyme_family, ":", paste(pathways, collapse = ", "), "\n")
              
              for(pathway in pathways) {
                if(!pathway %in% names(active_pathways)) {
                  active_pathways[[pathway]] <- list()
                  pathway_enzyme_counts[[pathway]] <- 0
                }
                
                # Add enzyme to pathway
                active_pathways[[pathway]][[enzyme_name]] <- list(
                  count = enzyme_count,
                  family = enzyme_family
                )
                pathway_enzyme_counts[[pathway]] <- pathway_enzyme_counts[[pathway]] + enzyme_count
              }
            } else {
              cat("No pathway mapping found for:", enzyme_family, "\n")
            }
          }
          
          cat("Active pathways found:", length(active_pathways), "\n")
          
          if(length(active_pathways) == 0) {
            return(plot_ly() %>%
                     add_text(x = 0.5, y = 0.5, text = paste("No recognized degradation pathways for", input$pathwaySelectedStrain), 
                              textfont = list(color = "orange", size = 16)) %>%
                     layout(
                       xaxis = list(showgrid = FALSE, showticklabels = FALSE, zeroline = FALSE),
                       yaxis = list(showgrid = FALSE, showticklabels = FALSE, zeroline = FALSE),
                       plot_bgcolor = 'rgba(0,0,0,0)',
                       paper_bgcolor = 'rgba(0,0,0,0)',
                       font = list(color = 'white')
                     ))
          }
          
          # Create pathway nodes
          for(pathway_name in names(active_pathways)) {
            pos <- pathway_positions[[pathway_name]]
            total_count <- pathway_enzyme_counts[[pathway_name]]
            
            pathway_nodes <- rbind(pathway_nodes, data.frame(
              id = paste0("pathway_", gsub("[^A-Za-z0-9]", "_", pathway_name)),
              label = pathway_name,
              short_label = gsub("[-\\s].*", "", pathway_name),
              type = "pathway",
              count = total_count,
              x = pos$x,
              y = pos$y,
              size = pmax(50, pmin(80, 50 + (total_count / 10))),
              color = pathway_colors[[pathway_name]],
              pathway = pathway_name,
              stringsAsFactors = FALSE
            ))
          }
          
          # Create enzyme nodes positioned between pathways they belong to
          processed_enzymes <- character()
          
          for(i in 1:nrow(strain_enzymes)) {
            enzyme_name <- strain_enzymes[[enzyme_col]][i]
            enzyme_count <- strain_enzymes[[input$pathwaySelectedStrain]][i]
            
            if(enzyme_name %in% processed_enzymes) next
            
            enzyme_family <- gsub("_.*", "", enzyme_name)
            enzyme_family <- gsub("([A-Z]+)([0-9]+).*", "\\1\\2", enzyme_family)
            
            if(enzyme_family %in% names(cazyme_pathway_map)) {
              pathways <- cazyme_pathway_map[[enzyme_family]]
              pathways_active <- pathways[pathways %in% names(active_pathways)]
              
              if(length(pathways_active) > 0) {
                # Calculate position based on layout theme
                if(layout_theme == "circular") {
                  # Position enzymes in concentric circles around pathways
                  primary_pathway <- pathways_active[1]
                  center_x <- pathway_positions[[primary_pathway]]$x
                  center_y <- pathway_positions[[primary_pathway]]$y
                  
                  # Create concentric positioning
                  angle_step <- 2*pi/max(6, length(active_pathways[[primary_pathway]]))
                  enzyme_idx <- which(names(active_pathways[[primary_pathway]]) == enzyme_name)
                  angle <- (enzyme_idx - 1) * angle_step
                  
                  radius <- if(length(pathways_active) > 1) 2.5 else 2.0
                  avg_x <- center_x + radius * cos(angle)
                  avg_y <- center_y + radius * sin(angle)
                  
                } else if(layout_theme == "hierarchical") {
                  # Position based on pathway hierarchy
                  primary_pathway <- pathways_active[1]
                  center_x <- pathway_positions[[primary_pathway]]$x
                  center_y <- pathway_positions[[primary_pathway]]$y
                  
                  # Create branching pattern
                  avg_x <- center_x + runif(1, -1.5, 1.5)
                  avg_y <- center_y - 1.5 + runif(1, -0.5, 0.5)
                  
                } else {
                  # Default: average position of all pathways
                  avg_x <- mean(sapply(pathways_active, function(p) pathway_positions[[p]]$x))
                  avg_y <- mean(sapply(pathways_active, function(p) pathway_positions[[p]]$y))
                }
                
                # Reduce jitter for cleaner appearance
                jitter_x <- runif(1, -0.3, 0.3)
                jitter_y <- runif(1, -0.3, 0.3)
                
                # Enhanced color scheme for multi-pathway enzymes
                if(length(pathways_active) > 1) {
                  # Gradient color for multi-functional enzymes
                  primary_color <- "#e2e8f0"  # Light gray for multi-functional
                } else {
                  # Single pathway - use pathway color but lighter
                  primary_color <- adjustcolor(pathway_colors[[pathways_active[1]]], alpha = 0.9)
                }
                
                enzyme_size <- pmax(15, pmin(35, 15 + enzyme_count * 2))
                
                enzyme_node <- data.frame(
                  id = paste0("enzyme_", gsub("[^A-Za-z0-9]", "_", enzyme_name)),
                  label = enzyme_name,
                  short_label = enzyme_family,
                  type = "enzyme",
                  count = enzyme_count,
                  x = avg_x + jitter_x,
                  y = avg_y + jitter_y,
                  size = enzyme_size,
                  color = adjustcolor(primary_color, alpha = 0.9),
                  pathway = paste(pathways_active, collapse = ", "),
                  stringsAsFactors = FALSE
                )
                
                all_nodes <- rbind(all_nodes, enzyme_node)
                
                # Create edges from enzyme to all its pathways
                for(pathway in pathways_active) {
                  pathway_pos <- pathway_positions[[pathway]]
                  
                  edge_width <- pmax(1, pmin(4, enzyme_count / 8))
                  
                  edge <- data.frame(
                    from_id = paste0("enzyme_", gsub("[^A-Za-z0-9]", "_", enzyme_name)),
                    to_id = paste0("pathway_", gsub("[^A-Za-z0-9]", "_", pathway)),
                    from_x = avg_x + jitter_x,
                    from_y = avg_y + jitter_y,
                    to_x = pathway_pos$x,
                    to_y = pathway_pos$y,
                    pathway = pathway,
                    color = adjustcolor(pathway_colors[[pathway]], alpha = 0.6),
                    width = edge_width,
                    stringsAsFactors = FALSE
                  )
                  
                  all_edges <- rbind(all_edges, edge)
                }
                
                processed_enzymes <- c(processed_enzymes, enzyme_name)
              }
            }
          }
          
          # Add pathway nodes to all_nodes
          all_nodes <- rbind(all_nodes, pathway_nodes)
          
          # Create the network visualization
          p <- plot_ly()
          
          # Enhanced edge styling
          if(nrow(all_edges) > 0) {
            for(i in 1:nrow(all_edges)) {
              edge <- all_edges[i, ]
              
              # Dynamic edge styling
              line_width <- pmax(1.5, pmin(5, edge$width))
              
              # Curved edges for better visualization
              mid_x <- (edge$from_x + edge$to_x) / 2
              mid_y <- (edge$from_y + edge$to_y) / 2
              
              p <- p %>% add_trace(
                x = c(edge$from_x, mid_x, edge$to_x),
                y = c(edge$from_y, mid_y, edge$to_y),
                type = "scatter",
                mode = "lines",
                line = list(
                  color = edge$color,
                  width = line_width,
                  shape = "spline",
                  smoothing = 0.8
                ),
                opacity = 0.8,
                hoverinfo = "none",
                showlegend = FALSE
              )
            }
          }
          
          # Enhanced enzyme nodes with better visibility
          enzyme_nodes <- all_nodes[all_nodes$type == "enzyme", ]
          if(nrow(enzyme_nodes) > 0) {
            p <- p %>% add_trace(
              data = enzyme_nodes,
              x = ~x, y = ~y,
              type = "scatter",
              mode = "markers+text",
              marker = list(
                size = ~size,
                color = ~color,
                line = list(color = "rgba(255,255,255,0.9)", width = 2.5),
                opacity = 1.0
              ),
              text = ~short_label,
              textposition = "middle center",
              textfont = list(
                color = "white",
                size = 10,
                family = "Arial Black"
              ),
              hovertemplate = ~paste(
                "<b>🧬 ", short_label, "</b><br>",
                "Full Name: ", label, "<br>",
                "Pathways: ", pathway, "<br>",
                "Count: ", count, "<br>",
                "<extra></extra>"
              ),
              name = "CAZymes",
              showlegend = FALSE
            )
          }
          
          # Enhanced pathway nodes (central hubs) with glow effect
          pathway_nodes_only <- all_nodes[all_nodes$type == "pathway", ]
          if(nrow(pathway_nodes_only) > 0) {
            p <- p %>% add_trace(
              data = pathway_nodes_only,
              x = ~x, y = ~y,
              type = "scatter",
              mode = "markers+text",
              marker = list(
                size = ~size,
                color = ~color,
                line = list(color = "white", width = 4),
                opacity = 1.0
              ),
              text = ~short_label,
              textposition = "middle center",
              textfont = list(
                color = "white",
                size = 13,
                family = "Arial Black"
              ),
              hovertemplate = ~paste(
                "<b>🎯 ", label, "</b><br>",
                "Total CAZyme Count: ", count, "<br>",
                "<extra></extra>"
              ),
              name = "Pathways",
              showlegend = FALSE
            )
          }
          
          # Calculate statistics
          total_pathways <- nrow(pathway_nodes_only)
          total_cazymes <- sum(pathway_nodes_only$count)
          # Count multifunctional CAZymes by checking pathway strings
          multifunctional_cazymes <- sum(grepl(",", enzyme_nodes$pathway))
          
          # Enhanced layout
          p %>% layout(
            title = list(
              text = paste("<b>🔗 Integrated CAZyme Network:</b>", input$pathwaySelectedStrain),
              font = list(color = "white", size = 20, family = "Arial Black"),
              x = 0.5,
              y = 0.95
            ),
            xaxis = list(
              range = c(-1, 19),
              showgrid = FALSE,
              showticklabels = FALSE,
              zeroline = FALSE,
              showline = FALSE
            ),
            yaxis = list(
              range = c(0, 13),
              showgrid = FALSE,
              showticklabels = FALSE,
              zeroline = FALSE,
              showline = FALSE
            ),
            plot_bgcolor = 'rgba(15,23,42,0.95)',
            paper_bgcolor = 'rgba(15,23,42,1)',
            font = list(color = 'white', family = "Arial"),
            showlegend = FALSE,
            annotations = list(
              list(
                x = 9,
                y = 0.5,
                text = paste("🎯 Pathways:", total_pathways, "| 🧪 CAZymes:", nrow(enzyme_nodes), "| 🔀 Multi-functional:", multifunctional_cazymes),
                showarrow = FALSE,
                font = list(color = "#60a5fa", size = 13, family = "Arial Bold"),
                xanchor = "center"
              )
            ),
            margin = list(l = 50, r = 50, t = 100, b = 80)
          )%>%
            config(
              toImageButtonOptions = list(
                format = 'png',
                filename = 'Integrated_CAZyme_Network',
                height = 1200,
                width = 1200,
                scale = 4  
              )
            )
          
        }, error = function(e) {
          cat("ERROR in singleStrainPathway:", e$message, "\n")
          print(traceback())
          plot_ly() %>%
            add_text(x = 0.5, y = 0.5, text = paste("Network Error:", e$message), 
                     textfont = list(color = "red", size = 14)) %>%
            layout(
              xaxis = list(showgrid = FALSE, showticklabels = FALSE, zeroline = FALSE),
              yaxis = list(showgrid = FALSE, showticklabels = FALSE, zeroline = FALSE),
              plot_bgcolor = 'rgba(0,0,0,0)',
              paper_bgcolor = 'rgba(0,0,0,0)',
              font = list(color = 'white')
            )
        })
        
      } else {
        # Strain not found in data
        plot_ly() %>%
          add_text(x = 0.5, y = 0.5, text = paste("Strain", input$pathwaySelectedStrain, "not found in data"), 
                   textfont = list(color = "red", size = 16)) %>%
          layout(
            xaxis = list(showgrid = FALSE, showticklabels = FALSE, zeroline = FALSE),
            yaxis = list(showgrid = FALSE, showticklabels = FALSE, zeroline = FALSE),
            plot_bgcolor = 'rgba(0,0,0,0)',
            paper_bgcolor = 'rgba(0,0,0,0)',
            font = list(color = 'white')
          )
      }
    } else {
      # No data available
      plot_ly() %>%
        add_text(x = 0.5, y = 0.5, text = "Please upload data and select a strain", 
                 textfont = list(color = "white", size = 16)) %>%
        layout(
          xaxis = list(showgrid = FALSE, showticklabels = FALSE, zeroline = FALSE),
          yaxis = list(showgrid = FALSE, showticklabels = FALSE, zeroline = FALSE),
          plot_bgcolor = 'rgba(0,0,0,0)',
          paper_bgcolor = 'rgba(0,0,0,0)',
          font = list(color = 'white')
        )
    }
  })
  
  # Enhanced Pathway Enzyme Distribution with matching theme
  output$pathwayEnzymeBreakdown <- renderPlotly({
    req(input$pathwaySelectedStrain, values$processed_data)
    
    tryCatch({
      enzyme_col <- "CAZymes"  
      if(!"CAZymes" %in% names(values$processed_data)) {
        enzyme_col <- names(values$processed_data)[1]
      }
      
      strain_enzymes <- values$processed_data %>%
        filter(!!sym(input$pathwaySelectedStrain) > 0) %>%
        select(all_of(c(enzyme_col, input$pathwaySelectedStrain)))
      
      if(nrow(strain_enzymes) == 0) {
        return(plot_ly() %>%
                 add_text(x = 0.5, y = 0.5, text = "No enzyme data available", 
                          textfont = list(color = "orange", size = 16)) %>%
                 layout(
                   xaxis = list(showgrid = FALSE, showticklabels = FALSE, zeroline = FALSE),
                   yaxis = list(showgrid = FALSE, showticklabels = FALSE, zeroline = FALSE),
                   plot_bgcolor = 'rgba(0,0,0,0)',
                   paper_bgcolor = 'rgba(0,0,0,0)',
                   font = list(color = 'white')
                 ))
      }
      
      # Enhanced distribution plot with gradient colors
      plot_ly(
        x = strain_enzymes[[input$pathwaySelectedStrain]],
        type = "histogram",
        marker = list(
          color = "#3b82f6", 
          opacity = 0.8,
          line = list(color = "white", width = 1.5)
        ),
        nbinsx = min(20, length(unique(strain_enzymes[[input$pathwaySelectedStrain]])))
      ) %>%
        layout(
          title = list(
            text = paste("📊 Enzyme Count Distribution -", input$pathwaySelectedStrain), 
            font = list(color = "white", size = 16, family = "Arial Bold")
          ),
          xaxis = list(
            title = list(text = "Enzyme Count", font = list(color = "white", size = 14)), 
            color = "white",
            gridcolor = "rgba(255,255,255,0.2)",
            gridwidth = 1
          ),
          yaxis = list(
            title = list(text = "Frequency", font = list(color = "white", size = 14)), 
            color = "white",
            gridcolor = "rgba(255,255,255,0.2)",
            gridwidth = 1
          ),
          plot_bgcolor = 'rgba(15,23,42,0.95)',
          paper_bgcolor = 'rgba(15,23,42,1)',
          font = list(color = 'white', family = "Arial"),
          margin = list(l = 60, r = 40, t = 80, b = 60)
        )
      
    }, error = function(e) {
      plot_ly() %>%
        add_text(x = 0.5, y = 0.5, text = paste("Distribution Error:", e$message), 
                 textfont = list(color = "red", size = 14)) %>%
        layout(
          xaxis = list(showgrid = FALSE, showticklabels = FALSE, zeroline = FALSE),
          yaxis = list(showgrid = FALSE, showticklabels = FALSE, zeroline = FALSE),
          plot_bgcolor = 'rgba(0,0,0,0)',
          paper_bgcolor = 'rgba(0,0,0,0)',
          font = list(color = 'white')
        )
    })
  })
  
  # Enhanced Pathway Enzyme Distribution (keeping this unchanged)
  output$pathwayEnzymeBreakdown <- renderPlotly({
    req(input$pathwaySelectedStrain, values$processed_data)
    
    tryCatch({
      enzyme_col <- "CAZymes"  
      if(!"CAZymes" %in% names(values$processed_data)) {
        enzyme_col <- names(values$processed_data)[1]
      }
      
      strain_enzymes <- values$processed_data %>%
        filter(!!sym(input$pathwaySelectedStrain) > 0) %>%
        select(all_of(c(enzyme_col, input$pathwaySelectedStrain)))
      
      if(nrow(strain_enzymes) == 0) {
        return(plot_ly() %>%
                 add_text(x = 0.5, y = 0.5, text = "No enzyme data available", 
                          textfont = list(color = "orange", size = 16)) %>%
                 layout(
                   xaxis = list(showgrid = FALSE, showticklabels = FALSE, zeroline = FALSE),
                   yaxis = list(showgrid = FALSE, showticklabels = FALSE, zeroline = FALSE),
                   plot_bgcolor = 'rgba(0,0,0,0)',
                   paper_bgcolor = 'rgba(0,0,0,0)',
                   font = list(color = 'white')
                 ))
      }
      
      # Create distribution plot with network theme
      plot_ly(
        x = strain_enzymes[[input$pathwaySelectedStrain]],
        type = "histogram",
        marker = list(
          color = "#60a5fa", 
          opacity = 0.8,
          line = list(color = "white", width = 1)
        ),
        nbinsx = 20
      ) %>%
        layout(
          title = list(
            text = "Enzyme Count Distribution", 
            font = list(color = "white", size = 16)
          ),
          xaxis = list(
            title = "Enzyme Count", 
            color = "white",
            gridcolor = "rgba(255,255,255,0.1)"
          ),
          yaxis = list(
            title = "Frequency", 
            color = "white",
            gridcolor = "rgba(255,255,255,0.1)"
          ),
          plot_bgcolor = 'rgba(15,23,42,1)',
          paper_bgcolor = 'rgba(15,23,42,1)',
          font = list(color = 'white', family = "Arial")
        )
      
    }, error = function(e) {
      plot_ly() %>%
        add_text(x = 0.5, y = 0.5, text = paste("Distribution Error:", e$message), 
                 textfont = list(color = "red", size = 14)) %>%
        layout(
          xaxis = list(showgrid = FALSE, showticklabels = FALSE, zeroline = FALSE),
          yaxis = list(showgrid = FALSE, showticklabels = FALSE, zeroline = FALSE),
          plot_bgcolor = 'rgba(0,0,0,0)',
          paper_bgcolor = 'rgba(0,0,0,0)',
          font = list(color = 'white')
        )
    })
  })
  
  # Enhanced Pathway Enzyme Distribution (keeping this unchanged)
  output$pathwayEnzymeBreakdown <- renderPlotly({
    req(input$pathwaySelectedStrain, values$processed_data)
    
    tryCatch({
      enzyme_col <- "CAZymes"  
      if(!"CAZymes" %in% names(values$processed_data)) {
        enzyme_col <- names(values$processed_data)[1]
      }
      
      strain_enzymes <- values$processed_data %>%
        filter(!!sym(input$pathwaySelectedStrain) > 0) %>%
        select(all_of(c(enzyme_col, input$pathwaySelectedStrain)))
      
      if(nrow(strain_enzymes) == 0) {
        return(plot_ly() %>%
                 add_text(x = 0.5, y = 0.5, text = "No enzyme data available", 
                          textfont = list(color = "orange", size = 16)) %>%
                 layout(
                   xaxis = list(showgrid = FALSE, showticklabels = FALSE, zeroline = FALSE),
                   yaxis = list(showgrid = FALSE, showticklabels = FALSE, zeroline = FALSE),
                   plot_bgcolor = 'rgba(0,0,0,0)',
                   paper_bgcolor = 'rgba(0,0,0,0)',
                   font = list(color = 'white')
                 ))
      }
      
      # Create distribution plot with network theme
      plot_ly(
        x = strain_enzymes[[input$pathwaySelectedStrain]],
        type = "histogram",
        marker = list(
          color = "#60a5fa", 
          opacity = 0.8,
          line = list(color = "white", width = 1)
        ),
        nbinsx = 20
      ) %>%
        layout(
          title = list(
            text = "Enzyme Count Distribution", 
            font = list(color = "white", size = 16)
          ),
          xaxis = list(
            title = "Enzyme Count", 
            color = "white",
            gridcolor = "rgba(255,255,255,0.1)"
          ),
          yaxis = list(
            title = "Frequency", 
            color = "white",
            gridcolor = "rgba(255,255,255,0.1)"
          ),
          plot_bgcolor = 'rgba(15,23,42,1)',
          paper_bgcolor = 'rgba(15,23,42,1)',
          font = list(color = 'white', family = "Arial")
        )
      
    }, error = function(e) {
      plot_ly() %>%
        add_text(x = 0.5, y = 0.5, text = paste("Distribution Error:", e$message), 
                 textfont = list(color = "red", size = 14)) %>%
        layout(
          xaxis = list(showgrid = FALSE, showticklabels = FALSE, zeroline = FALSE),
          yaxis = list(showgrid = FALSE, showticklabels = FALSE, zeroline = FALSE),
          plot_bgcolor = 'rgba(0,0,0,0)',
          paper_bgcolor = 'rgba(0,0,0,0)',
          font = list(color = 'white')
        )
    })
  })
  
  # Enhanced Pathway Enzyme Distribution (keeping this unchanged)
  output$pathwayEnzymeBreakdown <- renderPlotly({
    req(input$pathwaySelectedStrain, values$processed_data)
    
    tryCatch({
      enzyme_col <- "CAZymes"  
      if(!"CAZymes" %in% names(values$processed_data)) {
        enzyme_col <- names(values$processed_data)[1]
      }
      
      strain_enzymes <- values$processed_data %>%
        filter(!!sym(input$pathwaySelectedStrain) > 0) %>%
        select(all_of(c(enzyme_col, input$pathwaySelectedStrain)))
      
      if(nrow(strain_enzymes) == 0) {
        return(plot_ly() %>%
                 add_text(x = 0.5, y = 0.5, text = "No enzyme data available", 
                          textfont = list(color = "orange", size = 16)) %>%
                 layout(
                   xaxis = list(showgrid = FALSE, showticklabels = FALSE, zeroline = FALSE),
                   yaxis = list(showgrid = FALSE, showticklabels = FALSE, zeroline = FALSE),
                   plot_bgcolor = 'rgba(0,0,0,0)',
                   paper_bgcolor = 'rgba(0,0,0,0)',
                   font = list(color = 'white')
                 ))
      }
      
      # Create distribution plot with network theme
      plot_ly(
        x = strain_enzymes[[input$pathwaySelectedStrain]],
        type = "histogram",
        marker = list(
          color = "#60a5fa", 
          opacity = 0.8,
          line = list(color = "white", width = 1)
        ),
        nbinsx = 20
      ) %>%
        layout(
          title = list(
            text = "Enzyme Count Distribution", 
            font = list(color = "white", size = 16)
          ),
          xaxis = list(
            title = "Enzyme Count", 
            color = "white",
            gridcolor = "rgba(255,255,255,0.1)"
          ),
          yaxis = list(
            title = "Frequency", 
            color = "white",
            gridcolor = "rgba(255,255,255,0.1)"
          ),
          plot_bgcolor = 'rgba(15,23,42,1)',
          paper_bgcolor = 'rgba(15,23,42,1)',
          font = list(color = 'white', family = "Arial")
        )
      
    }, error = function(e) {
      plot_ly() %>%
        add_text(x = 0.5, y = 0.5, text = paste("Distribution Error:", e$message), 
                 textfont = list(color = "red", size = 14)) %>%
        layout(
          xaxis = list(showgrid = FALSE, showticklabels = FALSE, zeroline = FALSE),
          yaxis = list(showgrid = FALSE, showticklabels = FALSE, zeroline = FALSE),
          plot_bgcolor = 'rgba(0,0,0,0)',
          paper_bgcolor = 'rgba(0,0,0,0)',
          font = list(color = 'white')
        )
    })
  })
  
  # Industrial Pathway Assessment
  output$pathwayIndustrialAssessment <- renderText({
    req(input$pathwaySelectedStrain, values$processed_data)
    
    tryCatch({
      enzyme_col <- if("CAZymes" %in% names(values$processed_data)) "CAZymes" else names(values$processed_data)[1]
      
      strain_enzymes <- values$processed_data %>%
        filter(!!sym(input$pathwaySelectedStrain) > 0)
      
      total_enzymes <- nrow(strain_enzymes)
      total_activity <- sum(strain_enzymes[[input$pathwaySelectedStrain]])
      
      # Count pathway-relevant enzymes using individual family matching
      cellulose_families <- c("GH5", "GH6", "GH7", "GH8", "GH9", "GH12", "GH44", "GH45", "GH48")
      hemicellulose_families <- c("GH10", "GH11", "GH26", "GH30", "GH39", "GH43", "GH51", "GH52", "CE1", "CE2", "CE3", "CE4", "CE5")
      lignin_families <- c("AA1", "AA2", "AA3", "AA4", "AA5", "AA6", "AA8")
      
      # Count enzymes for each pathway
      cellulose_enzymes <- 0
      hemicellulose_enzymes <- 0
      lignin_enzymes <- 0
      
      for(family in cellulose_families) {
        cellulose_enzymes <- cellulose_enzymes + sum(grepl(paste0("^", family, "($|_|[0-9])"), strain_enzymes[[enzyme_col]]))
      }
      
      for(family in hemicellulose_families) {
        hemicellulose_enzymes <- hemicellulose_enzymes + sum(grepl(paste0("^", family, "($|_|[0-9])"), strain_enzymes[[enzyme_col]]))
      }
      
      for(family in lignin_families) {
        lignin_enzymes <- lignin_enzymes + sum(grepl(paste0("^", family, "($|_|[0-9])"), strain_enzymes[[enzyme_col]]))
      }
      
      total_biorefinery <- cellulose_enzymes + hemicellulose_enzymes + lignin_enzymes
      
      # Simple assessment logic
      if(total_activity > 100) {
        assessment <- "🟢 HIGH POTENTIAL: Excellent enzyme diversity and activity"
      } else if(total_activity > 50) {
        assessment <- "🟡 MODERATE POTENTIAL: Good enzyme complement"
      } else {
        assessment <- "🔴 LOW POTENTIAL: Limited enzyme activity"
      }
      
      potential_score <- if(total_biorefinery > 20) "EXCELLENT" else if(total_biorefinery > 10) "GOOD" else "LIMITED"
      
      paste0(
        "<div style='background: rgba(79, 172, 254, 0.1); padding: 20px; border-radius: 10px; border-left: 4px solid #4facfe;'>",
        "<h4 style='color: #4facfe; margin-top: 0;'>🏭 Industrial Assessment for ", input$pathwaySelectedStrain, "</h4>",
        "<p style='font-size: 16px; color: white; margin: 10px 0;'><strong>", assessment, "</strong></p>",
        "<div style='display: grid; grid-template-columns: 1fr 1fr; gap: 15px; margin: 15px 0;'>",
        "<div style='background: rgba(255,255,255,0.1); padding: 10px; border-radius: 5px;'>",
        "<strong style='color: #00d4ff;'>Total Enzymes:</strong><br>", total_enzymes, "</div>",
        "<div style='background: rgba(255,255,255,0.1); padding: 10px; border-radius: 5px;'>",
        "<strong style='color: #38ef7d;'>Activity Score:</strong><br>", total_activity, "</div>",
        "</div>",
        "<h5 style='color: #ff6b35; margin: 15px 0 10px 0;'>🧬 Biorefinery Potential: ", potential_score, "</h5>",
        "<div style='display: grid; grid-template-columns: 1fr 1fr 1fr; gap: 10px;'>",
        "<div style='text-align: center; padding: 8px; background: rgba(0,212,255,0.1); border-radius: 5px;'>",
        "<small style='color: #00d4ff;'>Cellulose</small><br><strong style='color: white;'>", cellulose_enzymes, "</strong></div>",
        "<div style='text-align: center; padding: 8px; background: rgba(56,239,125,0.1); border-radius: 5px;'>",
        "<small style='color: #38ef7d;'>Hemicellulose</small><br><strong style='color: white;'>", hemicellulose_enzymes, "</strong></div>",
        "<div style='text-align: center; padding: 8px; background: rgba(255,107,53,0.1); border-radius: 5px;'>",
        "<small style='color: #ff6b35;'>Lignin</small><br><strong style='color: white;'>", lignin_enzymes, "</strong></div>",
        "</div></div>"
      )
      
    }, error = function(e) {
      paste("Assessment Error:", e$message)
    })
  })
  
  # Pathway Statistics
  output$pathwayStats <- renderText({
    req(input$pathwaySelectedStrain, values$processed_data)
    
    tryCatch({
      enzyme_col <- if("CAZymes" %in% names(values$processed_data)) "CAZymes" else names(values$processed_data)[1]
      
      strain_enzymes <- values$processed_data %>%
        filter(!!sym(input$pathwaySelectedStrain) > 0)
      
      total_enzymes <- nrow(strain_enzymes)
      total_activity <- sum(strain_enzymes[[input$pathwaySelectedStrain]])
      
      # Count different enzyme families using simple pattern matching
      gh_count <- sum(grepl("^GH", strain_enzymes[[enzyme_col]]))
      ce_count <- sum(grepl("^CE", strain_enzymes[[enzyme_col]]))
      pl_count <- sum(grepl("^PL", strain_enzymes[[enzyme_col]]))
      aa_count <- sum(grepl("^AA", strain_enzymes[[enzyme_col]]))
      cbm_count <- sum(grepl("^CBM", strain_enzymes[[enzyme_col]]))
      
      paste0(
        "<div style='color: white; font-family: Inter, sans-serif;'>",
        "<div style='display: grid; grid-template-columns: 1fr 1fr; gap: 10px; margin-bottom: 15px;'>",
        "<div style='background: rgba(79,172,254,0.1); padding: 10px; border-radius: 5px; text-align: center;'>",
        "<strong style='color: #4facfe; font-size: 18px;'>", total_enzymes, "</strong><br>",
        "<small style='color: #cccccc;'>Total Enzymes</small></div>",
        "<div style='background: rgba(0,242,254,0.1); padding: 10px; border-radius: 5px; text-align: center;'>",
        "<strong style='color: #00f2fe; font-size: 18px;'>", total_activity, "</strong><br>",
        "<small style='color: #cccccc;'>Activity Score</small></div>",
        "</div>",
        "<h6 style='color: #4facfe; margin: 15px 0 10px 0;'>🧪 Enzyme Family Distribution</h6>",
        "<div style='font-size: 12px; line-height: 1.8;'>",
        "<div style='display: flex; justify-content: space-between; margin: 5px 0;'>",
        "<span>GH (Glycoside Hydrolases):</span><strong style='color: #38ef7d;'>", gh_count, "</strong></div>",
        "<div style='display: flex; justify-content: space-between; margin: 5px 0;'>",
        "<span>CE (Carbohydrate Esterases):</span><strong style='color: #ff6b35;'>", ce_count, "</strong></div>",
        "<div style='display: flex; justify-content: space-between; margin: 5px 0;'>",
        "<span>PL (Polysaccharide Lyases):</span><strong style='color: #9c59d1;'>", pl_count, "</strong></div>",
        "<div style='display: flex; justify-content: space-between; margin: 5px 0;'>",
        "<span>AA (Auxiliary Activities):</span><strong style='color: #f7931e;'>", aa_count, "</strong></div>",
        "<div style='display: flex; justify-content: space-between; margin: 5px 0;'>",
        "<span>CBM (Carb. Binding Modules):</span><strong style='color: #1abc9c;'>", cbm_count, "</strong></div>",
        "</div></div>"
      )
      
    }, error = function(e) {
      paste("Statistics Error:", e$message)
    })
  })
  
  # Pathway Enzyme Table
  output$pathwayEnzymeTable <- DT::renderDataTable({
    req(input$pathwaySelectedStrain, values$processed_data)
    
    tryCatch({
      enzyme_col <- if("CAZymes" %in% names(values$processed_data)) "CAZymes" else names(values$processed_data)[1]
      
      strain_data <- values$processed_data %>%
        filter(!!sym(input$pathwaySelectedStrain) > 0) %>%
        select(all_of(c(enzyme_col, input$pathwaySelectedStrain))) %>%
        arrange(desc(!!sym(input$pathwaySelectedStrain)))
      
      # Add pathway classification with individual family matching
      classify_enzyme_pathway <- function(enzyme) {
        # Define families for each pathway
        cellulolytic <- c("CBM1", "CBM2", "CBM3", "CBM4", "CBM6", "CBM8", "CBM9", "CBM10", "CBM16", "CBM17", "CBM28", "CBM30", "CBM37", "CBM44", "CBM46", "CBM49", "CBM59", "CBM63", "CBM64", "CBM72","GH5", "GH6", "GH7", "GH8", "GH9", "GH10", "GH12", "GH26", "GH44", "GH45", "GH48", "GH51", "GH74", "GH124", "GH1", "GH2", "GH3", "GH30", "GH39", "GH116",
                          "AA9", "AA10", "AA15", "AA3")
        
        hemi_cellulolytic <- c("GH1", "GH2", "GH3", "GH5", "GH8", "GH9", "GH10", "GH11", "GH12", "GH16", "GH17", "GH26", "GH30", "GH39", "GH43", "GH44", "GH51", "GH52", "GH54", "GH55", "GH62","GH74", "GH98", "GH113", "GH116", "GH120", "GH127", "GH134", "GH137", "GH141", "GH142", "GH146","CBM2", "CBM4", "CBM6", "CBM9", "CBM13", "CBM15", "CBM22", "CBM31", "CBM35", "CBM36", "CBM37", "CBM42", "CBM54", "CBM59", "CBM60", "CBM72", "CBM16", "CBM23", "CBM27", "CBM29", "CBM62", "CBM76", "CBM80", "CBM44", "CBM65", "CBM75", "CBM78", "CBM81",
                               "CE1", "CE2", "CE3", "CE4", "CE5", "CE6", "CE7", "CE12", "CE15",
                               "AA9", "AA14")
        
        lignolytic <- c("AA1", "AA2", "AA3", "AA4", "AA5", "AA6", "AA8", "AA9", "AA12","CE1", "CE15", "CE10")
        
        pectinolytic <- c("GH1", "GH2", "GH3", "GH4", "GH10", "GH28", "GH33", "GH35", "GH39", "GH42", "GH43", "GH50", "GH51", "GH54", "GH59", "GH62", "GH78", "GH93", "GH106", "GH147",
                          "PL1", "PL2", "PL3", "PL4", "PL9", "PL10", "PL11", "PL22", "PL26","CE6", "CE8", "CE12", "CE13","CBM13", "CBM32", "CBM41", "CBM51", "CBM61", "CBM62", "CBM67",
                          "CBM77", "CBM80")
        
        starch_degrading <- c("GH13", "GH14", "GH57", "GH119", "GH126", "GH4", "GH31", "GH63", "GH97", "GH122", "GT35","CBM20", "CBM21", "CBM25", "CBM26", "CBM34", "CBM45", "CBM53", "CBM69","CBM74", "CBM82", "CBM83","AA13")
        
        inulin_degrading <- c("GH32", "GH91","CBM38")
        
        # Collect all matching pathways
        pathways <- c()
        
        # Check each pathway and collect all matches
        for(family in cellulolytic) {
          if(grepl(paste0("^", family, "($|_|[0-9])"), enzyme)) {
            pathways <- c(pathways, "Cellulose")
            break
          }
        }
        for(family in hemi_cellulolytic) {
          if(grepl(paste0("^", family, "($|_|[0-9])"), enzyme)) {
            pathways <- c(pathways, "Hemicellulose")
            break
          }
        }
        for(family in lignolytic) {
          if(grepl(paste0("^", family, "($|_|[0-9])"), enzyme)) {
            pathways <- c(pathways, "Lignin")
            break
          }
        }
        for(family in pectinolytic) {
          if(grepl(paste0("^", family, "($|_|[0-9])"), enzyme)) {
            pathways <- c(pathways, "Pectin")
            break
          }
        }
        for(family in starch_degrading) {
          if(grepl(paste0("^", family, "($|_|[0-9])"), enzyme)) {
            pathways <- c(pathways, "Starch")
            break
          }
        }
        for(family in inulin_degrading) {
          if(grepl(paste0("^", family, "($|_|[0-9])"), enzyme)) {
            pathways <- c(pathways, "Inulin")
            break
          }
        }
        
        # Return all pathways joined by comma, or "Other" if none found
        if(length(pathways) > 0) {
          return(paste(pathways, collapse = ", "))
        } else {
          return("Other")
        }
      }
      
      strain_data$Pathway <- sapply(strain_data[[enzyme_col]], classify_enzyme_pathway)
      
      # Rename columns for display
      colnames(strain_data) <- c("Enzyme", "Count", "Pathway")
      
      DT::datatable(
        strain_data,
        options = list(
          pageLength = 15,
          scrollX = TRUE,
          dom = 'Bfrtip',
          buttons = c('copy', 'csv', 'excel'),
          columnDefs = list(list(className = 'dt-center', targets = c(1, 2)))
        ),
        class = 'cell-border stripe hover',
        rownames = FALSE
      ) %>%
        DT::formatStyle(
          'Pathway',
          backgroundColor = DT::styleEqual(
            c('Cellulose', 'Hemicellulose', 'Lignin', 'Pectin', 'Starch', 'Inulin', 'Other'),
            c('rgba(0,212,255,0.3)', 'rgba(56,239,125,0.3)', 'rgba(255,107,53,0.3)', 
              'rgba(156,89,209,0.3)', 'rgba(247,147,30,0.3)', 'rgba(26,188,156,0.3)', 'rgba(128,128,128,0.3)')
          )
        )
      
    }, error = function(e) {
      DT::datatable(data.frame(Error = paste("Table Error:", e$message)), options = list(dom = 't'))
    })
  })
  
  # Populate the dropdown when data is loaded
  observe({
    if (!is.null(values$processed_data)) {
      # Get strain names (column names excluding the first column which is likely enzyme names)
      strain_choices <- names(values$processed_data)
      
      # Remove common non-strain columns
      strain_choices <- strain_choices[!strain_choices %in% c("Annotations/Genomes", "CAZymes", "Enzyme", "Family", "Description")]
      
      # Update the dropdown choices
      updateSelectInput(session, "pathwaySelectedStrain",
                        choices = strain_choices,
                        selected = if(length(strain_choices) > 0) strain_choices[1] else NULL)
    }
  })
  
  # Action button functionality (if you want the button to trigger updates)
  observeEvent(input$generatePathwayNetwork, {
    # Force refresh of all pathway outputs
    if (!is.null(input$pathwaySelectedStrain) && !is.null(values$processed_data)) {
      # You can add any additional processing here if needed
      showNotification("Pathway network generated successfully!", type = "message")
    } else {
      showNotification("Please select a strain and ensure data is loaded.", type = "warning")
    }
  })
}
```
