## Supplementary material for "CAZyXplorer: A Shiny Application for Cost-Effective Preliminary Screening of Microbial Strains to Advance Enzyme Discovery in Biorefining and Biotechnology": ui.R

```
# Complete UI code with Single Strain Degradation Pathway replacing Export & Reports
ui <- dashboardPage(
  
  # Header with enhanced styling
  dashboardHeader(
    title = tags$span(
      icon("atom", style = "margin-right: 10px; color: #00d4ff;"),
      "CAZyXplorer",
      style = "font-family: 'Inter', sans-serif; font-weight: 750; font-size: 30px; 
             text-transform: none; letter-spacing: 0.3px; 
             background: linear-gradient(45deg, #00d4ff, #ff6b35); 
             -webkit-background-clip: text; -webkit-text-fill-color: transparent;"
    ),
    titleWidth = 322
  ),
  
  # Enhanced Sidebar
  dashboardSidebar(
    width = 320,
    div(style = "background: linear-gradient(135deg, #1a1a2e 0%, #16213e 50%, #0f3460 100%); height: 100vh;",
        sidebarMenu(
          id = "sidebarMenu",
          menuItem("🚀 Data Upload", tabName = "upload", icon = icon("cloud-upload-alt")),
          menuItem("📊 Dashboard Overview", tabName = "overview", icon = icon("tachometer-alt")),
          
          # Enhanced Strain Analysis Menu
          menuItem("🧬 Strain Analysis", icon = icon("dna"),
                   menuSubItem("🔬 Single Strain Deep Dive", tabName = "single_strain"),
                   menuSubItem("⚔️ Multi-Strain Battle", tabName = "multi_strain"),
                   menuSubItem("🏆 Strain Ranking", tabName = "strain_ranking")
          ),
          
          # Enhanced Plant Cell Wall Degradation Menu
          menuItem("🌱 Degradation Arsenal", icon = icon("leaf"),
                   menuSubItem("🌾 Cellulolytic Power", tabName = "cellulolytic"),
                   menuSubItem("🌿 Hemi-Cellulolytic Force", tabName = "hemi_cellulolytic"),
                   menuSubItem("🌳 Lignolytic Potential", tabName = "lignolytic"),
                   menuSubItem("🍎 Pectinolytic Capacity", tabName = "pectinolytic"),
                   menuSubItem("🌽 Starch Degradation", tabName = "starch_degrading"),
                   menuSubItem("🥔 Inulin Processing", tabName = "inulin_degrading")
          ),
          
          menuItem("🎯 Biorefinery Optimizer", tabName = "biorefinery", icon = icon("industry")),
          menuItem("📈 Advanced Analytics", tabName = "analytics", icon = icon("chart-line")),
          menuItem("🧬 Pathway Network", tabName = "pathway_network", icon = icon("project-diagram"))
        ),
        
        # Enhanced Data upload section
        br(),
        div(style = "padding: 20px; 
                   background: linear-gradient(135deg, rgba(0, 212, 255, 0.1) 0%, rgba(255, 107, 53, 0.1) 100%); 
                   border-radius: 15px; 
                   border: 2px solid #00d4ff; 
                   margin: 15px; 
                   box-shadow: 0 0 25px rgba(0, 212, 255, 0.3);",
            h4("⚡ QUANTUM UPLOAD", 
               style = "color: #00d4ff; font-family: 'Inter', sans-serif; 
                        font-weight: 700; text-transform: uppercase; 
                        letter-spacing: 0.5px; margin-bottom: 15px; font-size: 16px;"),
            fileInput("csvFile", 
                      label = NULL,
                      accept = ".csv",
                      placeholder = "Upload CAZy annotation CSV..."),
            
            conditionalPanel(
              condition = "output.dataUploaded",
              actionButton("refreshData", "🔄 REFRESH QUANTUM ANALYSIS", 
                           class = "btn-primary btn-sm",
                           style = "width: 100%; font-family: 'Inter', sans-serif; 
                               font-weight: 600; text-transform: uppercase; 
                               letter-spacing: 0.3px; background: linear-gradient(45deg, #00d4ff, #ff6b35);
                               border: none; box-shadow: 0 0 20px rgba(0, 212, 255, 0.5); font-size: 13px;")
            )
        )
    )
  ),
  
  # Enhanced Body
  dashboardBody(
    
    # Advanced Custom CSS with improved fonts
    tags$head(
      # Updated Google Fonts import
      tags$link(href = "https://fonts.googleapis.com/css2?family=Inter:wght@300;400;500;600;700;900&family=JetBrains+Mono:wght@400;500;700&display=swap", rel = "stylesheet"),
      tags$style(HTML("
        /* Base typography improvements */
        body {
          background: linear-gradient(135deg, #0f0f23 0%, #1a1a2e 50%, #16213e 100%);
          color: #ffffff;
          font-family: 'Inter', -apple-system, BlinkMacSystemFont, 'Segoe UI', sans-serif;
          font-size: 14px;
          line-height: 1.5;
          font-weight: 400;
        }
        
        /* Headers - Use Inter with appropriate weights */
        h1, h2, h3, h4, h5, h6 {
          font-family: 'Inter', sans-serif;
          font-weight: 600;
          letter-spacing: 0.5px;
          color: #ffffff;
        }
        
        h1 { font-size: 28px; font-weight: 700; }
        h2 { font-size: 24px; font-weight: 600; }
        h3 { font-size: 20px; font-weight: 600; }
        h4 { font-size: 18px; font-weight: 600; }
        h5 { font-size: 16px; font-weight: 500; }
        h6 { font-size: 14px; font-weight: 500; }
        
        /* Update title styling to use Inter */
        .main-header .navbar-brand {
          font-family: 'Inter', sans-serif !important;
          font-weight: 700 !important;
          font-size: 20px !important;
          text-transform: none !important;
          letter-spacing: 0.5px !important;
        }
        
        /* Sidebar menu improvements */
        .sidebar-menu > li > a {
          font-family: 'Inter', sans-serif;
          font-size: 14px;
          font-weight: 500;
          color: #e2e8f0;
        }
        
        .sidebar-menu .treeview-menu > li > a {
          font-family: 'Inter', sans-serif;
          font-size: 13px;
          font-weight: 400;
          color: #cbd5e0;
        }
        
        .content-wrapper, .right-side {
          background: linear-gradient(135deg, #0f0f23 0%, #1a1a2e 50%, #16213e 100%);
          color: #ffffff;
          font-family: 'Inter', sans-serif;
        }
        
        /* Box titles */
        .box-title {
          font-family: 'Inter', sans-serif;
          font-weight: 600;
          font-size: 16px;
          letter-spacing: 0.3px;
        }
        
        .box {
          background: linear-gradient(135deg, rgba(26, 26, 46, 0.9) 0%, rgba(22, 33, 62, 0.9) 100%);
          border: 1px solid rgba(0, 212, 255, 0.3);
          border-radius: 15px;
          box-shadow: 0 10px 30px rgba(0, 0, 0, 0.5);
          color: #ffffff;
        }
        
        .box.box-solid.box-primary > .box-header {
          background: linear-gradient(45deg, #00d4ff, #ff6b35);
          color: white;
          border-radius: 15px 15px 0 0;
        }
        
        /* Value boxes */
        .small-box {
          border-radius: 15px;
          box-shadow: 0 10px 25px rgba(0, 0, 0, 0.3);
          border: 1px solid rgba(255, 255, 255, 0.1);
        }
        
        .small-box h3 {
          font-family: 'Inter', sans-serif;
          font-weight: 700;
          font-size: 32px;
        }
        
        .small-box p {
          font-family: 'Inter', sans-serif;
          font-weight: 500;
          font-size: 14px;
        }
        
        .small-box.bg-yellow { 
          background: linear-gradient(135deg, #f39c12 0%, #e67e22 100%) !important; 
        }
        .small-box.bg-green { 
          background: linear-gradient(135deg, #00a65a 0%, #27ae60 100%) !important; 
        }
        .small-box.bg-blue { 
          background: linear-gradient(135deg, #3c8dbc 0%, #2980b9 100%) !important; 
        }
        .small-box.bg-red { 
          background: linear-gradient(135deg, #dd4b39 0%, #e74c3c 100%) !important; 
        }
        .small-box.bg-purple { 
          background: linear-gradient(135deg, #9b59b6 0%, #8e44ad 100%) !important; 
        }
        .small-box.bg-teal { 
          background: linear-gradient(135deg, #1abc9c 0%, #16a085 100%) !important; 
        }
        
        .enzyme-card {
          background: linear-gradient(135deg, #667eea 0%, #764ba2 100%);
          color: white;
          padding: 25px;
          border-radius: 15px;
          margin: 15px 0;
          box-shadow: 0 15px 35px rgba(102, 126, 234, 0.3);
          border: 1px solid rgba(255, 255, 255, 0.1);
          font-family: 'Inter', sans-serif;
        }
        
        .degradation-card {
          background: linear-gradient(135deg, #11998e 0%, #38ef7d 100%);
          color: white;
          padding: 20px;
          border-radius: 15px;
          margin: 15px 0;
          text-align: center;
          box-shadow: 0 15px 35px rgba(17, 153, 142, 0.3);
          border: 1px solid rgba(255, 255, 255, 0.1);
          font-family: 'Inter', sans-serif;
        }
        
        .strain-card {
          background: linear-gradient(135deg, #ff6b6b 0%, #ffa500 100%);
          color: white;
          padding: 20px;
          border-radius: 15px;
          margin: 15px 0;
          box-shadow: 0 15px 35px rgba(255, 107, 107, 0.3);
          border: 1px solid rgba(255, 255, 255, 0.1);
          font-family: 'Inter', sans-serif;
        }
        
        .biorefinery-card {
          background: linear-gradient(135deg, #667eea 0%, #764ba2 100%);
          color: white;
          padding: 25px;
          border-radius: 15px;
          margin: 15px 0;
          box-shadow: 0 20px 40px rgba(102, 126, 234, 0.4);
          border: 2px solid rgba(255, 255, 255, 0.1);
          font-family: 'Inter', sans-serif;
        }
        
        .pathway-card {
          background: linear-gradient(135deg, #4facfe 0%, #00f2fe 100%);
          color: white;
          padding: 25px;
          border-radius: 15px;
          margin: 15px 0;
          box-shadow: 0 20px 40px rgba(79, 172, 254, 0.4);
          border: 2px solid rgba(255, 255, 255, 0.1);
          font-family: 'Inter', sans-serif;
        }
        
        .potential-high { 
          color: #00ff88; 
          font-weight: 600; 
          font-family: 'Inter', sans-serif;
          text-shadow: 0 0 10px rgba(0, 255, 136, 0.5);
        }
        .potential-medium { 
          color: #ffaa00; 
          font-weight: 600; 
          font-family: 'Inter', sans-serif;
          text-shadow: 0 0 10px rgba(255, 170, 0, 0.5);
        }
        .potential-low { 
          color: #ff4444; 
          font-weight: 600; 
          font-family: 'Inter', sans-serif;
          text-shadow: 0 0 10px rgba(255, 68, 68, 0.5);
        }
        
        .quantum-glow {
          animation: quantumGlow 2s ease-in-out infinite alternate;
        }
        
        @keyframes quantumGlow {
          from { box-shadow: 0 0 20px rgba(0, 212, 255, 0.5); }
          to { box-shadow: 0 0 40px rgba(255, 107, 53, 0.7); }
        }
        
        .welcome-screen {
          text-align: center;
          padding: 80px;
          color: #ffffff;
          font-family: 'Inter', sans-serif;
        }
        
        .metric-card {
          background: linear-gradient(135deg, rgba(0, 212, 255, 0.1) 0%, rgba(255, 107, 53, 0.1) 100%);
          border: 2px solid rgba(0, 212, 255, 0.5);
          border-radius: 15px;
          padding: 20px;
          margin: 10px 0;
          color: #ffffff;
          box-shadow: 0 10px 30px rgba(0, 0, 0, 0.3);
          font-family: 'Inter', sans-serif;
        }
        
        .industrial-highlight {
          font-size: 2.2em;
          font-weight: 700;
          color: #00d4ff;
          text-shadow: 0 0 20px rgba(0, 212, 255, 0.8);
          font-family: 'Inter', sans-serif;
          letter-spacing: 0.5px;
        }
        
        /* Tables */
        .dataTables_wrapper {
          color: #ffffff;
          font-family: 'Inter', sans-serif;
        }
        
        .dataTables_wrapper .dataTables_length,
        .dataTables_wrapper .dataTables_filter,
        .dataTables_wrapper .dataTables_info,
        .dataTables_wrapper .dataTables_paginate {
          color: #ffffff;
          font-family: 'Inter', sans-serif;
        }
        
        table.dataTable {
          background: rgba(26, 26, 46, 0.9);
          color: #ffffff;
          font-family: 'Inter', sans-serif;
          font-size: 13px;
        }
        
        table.dataTable thead th {
          background: linear-gradient(45deg, #00d4ff, #ff6b35);
          color: white;
          font-family: 'Inter', sans-serif;
          font-weight: 600;
          font-size: 14px;
        }
        
        table.dataTable tbody td {
          font-family: 'Inter', sans-serif;
          font-weight: 400;
          font-size: 13px;
          color: #f7fafc;
        }
        
        /* Form inputs */
        .form-control, .selectize-input {
          font-family: 'Inter', sans-serif;
          font-size: 14px;
          color: #ffffff;
          background: rgba(26, 26, 46, 0.9);
          border: 1px solid rgba(0, 212, 255, 0.5);
        }
        
        .selectize-dropdown {
          background: rgba(26, 26, 46, 0.95);
          color: #ffffff;
          border: 1px solid rgba(0, 212, 255, 0.5);
          font-family: 'Inter', sans-serif;
        }
        
        /* Buttons */
        .btn {
          font-family: 'Inter', sans-serif;
          font-weight: 600;
          font-size: 14px;
          text-transform: none;
          letter-spacing: 0.3px;
        }
        
        /* Code/monospace elements */
        code, pre, .code-text {
          font-family: 'JetBrains Mono', 'Monaco', 'Menlo', 'Ubuntu Mono', monospace;
          font-size: 13px;
          font-weight: 400;
        }
      "))
    ),
    
    tabItems(
      
      # Enhanced Data Upload Tab
      tabItem(tabName = "upload",
              fluidRow(
                box(
                  title = tags$span(
                    icon("rocket", style = "margin-right: 10px; color: #00d4ff;"),
                    "QUANTUM DATA INTEGRATION CENTER",
                    style = "font-family: 'Inter', sans-serif; font-weight: 700; text-transform: uppercase; letter-spacing: 0.5px;"
                  ), 
                  status = "primary", 
                  solidHeader = TRUE,
                  width = 12,
                  
                  fluidRow(
                    column(6,
                           fileInput("mainCsvFile", 
                                     tags$span("📁 UPLOAD CAZy ANNOTATION MATRIX", 
                                               style = "font-family: 'Inter', sans-serif; font-weight: 700; text-transform: uppercase; letter-spacing: 0.5px; color: #00d4ff;"),
                                     accept = ".csv",
                                     width = "100%"),
                           
                           conditionalPanel(
                             condition = "output.dataUploaded",
                             div(class = "enzyme-card quantum-glow",
                                 h4("✅ QUANTUM DATA MATRIX LOADED"),
                                 textOutput("dataInfo"),
                                 br(),
                                 div(class = "metric-card",
                                     h5("🧬 DATA QUALITY METRICS"),
                                     textOutput("dataQuality")
                                 )
                             )
                           )
                    ),
                    
                    column(6,
                           div(class = "biorefinery-card quantum-glow",
                               h4("📋 EXPECTED BIOREFINERY DATA FORMAT", 
                                  style = "font-family: 'Inter', sans-serif; font-weight: 700; color: #00d4ff; text-transform: uppercase; letter-spacing: 0.5px;"),
                               
                               div(style = "background: rgba(0, 212, 255, 0.1); padding: 15px; border-radius: 10px; margin: 10px 0; border-left: 4px solid #00d4ff;",
                                   p(tags$strong("🎯 REQUIRED COLUMNS:", style = "color: #00d4ff; font-family: 'Inter', sans-serif; font-weight: 700;")),
                                   tags$ul(
                                     tags$li("Annotations/Genomes (CAZy families)", style = "color: #cccccc; font-family: 'Inter', sans-serif; font-weight: 500;"),
                                     tags$li("Genome columns (enzyme counts)", style = "color: #cccccc; font-family: 'Inter', sans-serif; font-weight: 500;"),
                                     tags$li("Total column (optional)", style = "color: #cccccc; font-family: 'Inter', sans-serif; font-weight: 500;")
                                   )
                               ),
                               
                               div(style = "background: rgba(255, 107, 53, 0.1); padding: 15px; border-radius: 10px; margin: 10px 0; border-left: 4px solid #ff6b35;",
                                   p(tags$strong("🏭 BIOREFINERY ENZYME CLASSES:", style = "color: #ff6b35; font-family: 'Inter', sans-serif; font-weight: 700;")),
                                   tags$code("GH, AA, CBM, PL, CE families for plant biomass degradation", 
                                             style = "background: rgba(26, 26, 46, 0.8); color: #38ef7d; padding: 8px; border-radius: 5px; font-family: 'JetBrains Mono', monospace;")
                               )
                           )
                    )
                  ),
                  
                  conditionalPanel(
                    condition = "output.dataUploaded",
                    hr(style = "border-color: #00d4ff; border-width: 2px;"),
                    h4("🔍 QUANTUM DATA PREVIEW", 
                       style = "font-family: 'Inter', sans-serif; font-weight: 700; color: #00d4ff; text-transform: uppercase; letter-spacing: 0.5px;"),
                    withSpinner(DT::dataTableOutput("dataPreview"), color = "#00d4ff")
                  )
                )
              )
      ),
      
      # Enhanced Overview Dashboard Tab
      tabItem(tabName = "overview",
              conditionalPanel(
                condition = "!output.dataUploaded",
                fluidRow(
                  box(
                    title = tags$span(
                      icon("atom", style = "margin-right: 10px; color: #00d4ff;"),
                      "WELCOME TO THE BIOREFINERY QUANTUM LAB",
                      style = "font-family: 'Inter', sans-serif; font-weight: 700; text-transform: uppercase; letter-spacing: 0.5px;"
                    ),
                    status = "info",
                    solidHeader = TRUE,
                    width = 12,
                    height = "600px",
                    div(class = "welcome-screen",
                        icon("atom", style = "font-size: 8em; color: #00d4ff; margin-bottom: 30px; 
                                   text-shadow: 0 0 50px rgba(0, 212, 255, 0.8);"),
                        h2("🧬 INDUSTRIAL ENZYME DISCOVERY PLATFORM", 
                           style = "background: linear-gradient(45deg, #00d4ff, #ff6b35); 
                            -webkit-background-clip: text; -webkit-text-fill-color: transparent;
                            font-family: 'Inter', sans-serif; font-weight: 700;"),
                        h4("Advanced bioinformatics platform for bacterial plant cell wall degrading enzymes", 
                           style = "font-size: 24px; margin-bottom: 30px; color: #cccccc; font-family: 'Inter', sans-serif;"),
                        div(style = "display: grid; grid-template-columns: 1fr 1fr 1fr; gap: 30px; margin: 40px 0;",
                            div(class = "metric-card",
                                icon("dna", style = "font-size: 3em; color: #00d4ff;"),
                                h5("CAZy Analysis", style = "color: #00d4ff; font-weight: 700; font-family: 'Inter', sans-serif;"),
                                p("Comprehensive enzyme classification", style = "font-family: 'Inter', sans-serif;")
                            ),
                            div(class = "metric-card",
                                icon("industry", style = "font-size: 3em; color: #ff6b35;"),
                                h5("Biorefinery Optimization", style = "color: #ff6b35; font-weight: 700; font-family: 'Inter', sans-serif;"),
                                p("Industrial strain selection", style = "font-family: 'Inter', sans-serif;")
                            ),
                            div(class = "metric-card",
                                icon("chart-line", style = "font-size: 3em; color: #38ef7d;"),
                                h5("Advanced Analytics", style = "color: #38ef7d; font-weight: 700; font-family: 'Inter', sans-serif;"),
                                p("AI-powered insights", style = "font-family: 'Inter', sans-serif;")
                            )
                        ),
                        actionButton("goToUpload", "🚀 INITIATE QUANTUM ANALYSIS", 
                                     class = "btn-primary btn-lg quantum-glow",
                                     style = "padding: 20px 50px; font-size: 22px; font-weight: 700; font-family: 'Inter', sans-serif;
                                     background: linear-gradient(45deg, #00d4ff, #ff6b35);
                                     border: none; border-radius: 15px; text-transform: uppercase; letter-spacing: 0.5px;")
                    )
                  )
                )
              ),
              
              conditionalPanel(
                condition = "output.dataUploaded",
                # Enhanced Summary metrics
                fluidRow(
                  valueBoxOutput("totalEnzymes", width = 2),
                  valueBoxOutput("totalGenomes", width = 2),
                  valueBoxOutput("avgEnzymesPerGenome", width = 2),
                  valueBoxOutput("topEnzymeFamily", width = 2),
                  valueBoxOutput("biodiversityIndex", width = 2),
                  valueBoxOutput("industrialPotential", width = 2)
                ),
                
                # Enhanced Main charts
                fluidRow(
                  box(
                    title = tags$span(
                      icon("chart-area", style = "margin-right: 10px;"),
                      "ENZYME DISTRIBUTION QUANTUM MATRIX",
                      style = "font-family: 'Inter', sans-serif; font-weight: 700; text-transform: uppercase; letter-spacing: 0.5px;"
                    ),
                    status = "primary",
                    solidHeader = TRUE,
                    width = 8,
                    withSpinner(plotlyOutput("enzymeDistribution", height = "450px"), color = "#00d4ff")
                  ),
                  
                  box(
                    title = tags$span(
                      icon("trophy", style = "margin-right: 10px;"),
                      "ELITE ENZYME ARSENAL",
                      style = "font-family: 'Inter', sans-serif; font-weight: 700; text-transform: uppercase; letter-spacing: 0.5px;"
                    ),
                    status = "success",
                    solidHeader = TRUE,
                    width = 4,
                    withSpinner(plotlyOutput("topFamilies", height = "450px"), color = "#11998e")
                  )
                ),
                
                fluidRow(
                  box(
                    title = tags$span(
                      icon("fire", style = "margin-right: 10px;"),
                      "INDUSTRIAL BIOREFINERY HEATMAP",
                      style = "font-family: 'Inter', sans-serif; font-weight: 700; text-transform: uppercase; letter-spacing: 0.5px;"
                    ),
                    status = "warning",
                    solidHeader = TRUE,
                    width = 6,
                    withSpinner(plotlyOutput("enzymeHeatmap", height = "500px"), color = "#f7931e")
                  ),
                  
                  box(
                    title = tags$span(
                      icon("network-wired", style = "margin-right: 10px;"),
                      "DEGRADATION PATHWAY NETWORK",
                      style = "font-family: 'Inter', sans-serif; font-weight: 700; text-transform: uppercase; letter-spacing: 0.5px;"
                    ),
                    status = "info",
                    solidHeader = TRUE,
                    width = 6,
                    withSpinner(forceNetworkOutput("enzymeNetwork", height = "500px"), color = "#3c8dbc")
                  )
                )
              )
      ),
      
      # Enhanced Single Strain Analysis Tab
      tabItem(tabName = "single_strain",
              conditionalPanel(
                condition = "output.dataUploaded",
                fluidRow(
                  box(
                    title = tags$span(
                      icon("microscope", style = "margin-right: 10px;"),
                      "SINGLE STRAIN QUANTUM ANALYZER",
                      style = "font-family: 'Inter', sans-serif; font-weight: 700; text-transform: uppercase; letter-spacing: 0.5px;"
                    ),
                    status = "primary",
                    solidHeader = TRUE,
                    width = 4,
                    
                    selectInput("selectedStrain", 
                                tags$span("🦠 SELECT TARGET BIOREFINERY STRAIN:", 
                                          style = "font-family: 'Inter', sans-serif; font-weight: 700; color: #00d4ff;"),
                                choices = NULL,
                                selected = NULL),
                    
                    div(class = "strain-card quantum-glow",
                        h4("🧬 STRAIN BIOACTIVITY MATRIX"),
                        htmlOutput("strainSummary")
                    ),
                    
                    div(class = "metric-card",
                        h5("🎯 INDUSTRIAL METRICS"),
                        htmlOutput("strainMetrics")
                    ),
                    
                    actionButton("analyzeSingleStrain", "🔬 INITIATE DEEP QUANTUM SCAN",
                                 class = "btn-primary btn-lg quantum-glow",
                                 style = "width: 100%; margin-top: 15px; font-family: 'Inter', sans-serif; 
                                   font-weight: 700; text-transform: uppercase; letter-spacing: 0.5px;
                                   background: linear-gradient(45deg, #00d4ff, #ff6b35); border: none;")
                  ),
                  
                  box(
                    title = tags$span(
                      icon("dna", style = "margin-right: 10px;"),
                      "STRAIN ENZYME BIOACTIVITY SPECTRUM",
                      style = "font-family: 'Inter', sans-serif; font-weight: 700; text-transform: uppercase; letter-spacing: 0.5px;"
                    ),
                    status = "success",
                    solidHeader = TRUE,
                    width = 8,
                    
                    tabsetPanel(
                      tabPanel(tags$span("📊 ENZYME DISTRIBUTION", style = "font-family: 'Inter', sans-serif; font-weight: 700;"), 
                               withSpinner(plotlyOutput("singleStrainDistribution", height = "450px"), color = "#11998e")),
                      tabPanel(tags$span("🎯 DEGRADATION ARSENAL", style = "font-family: 'Inter', sans-serif; font-weight: 700;"), 
                               withSpinner(plotlyOutput("singleStrainCapabilities", height = "450px"), color = "#11998e")),
                      tabPanel(tags$span("🔬 ENZYME CLASSES", style = "font-family: 'Inter', sans-serif; font-weight: 700;"), 
                               withSpinner(plotlyOutput("enzymeClassDistribution", height = "450px"), color = "#11998e")),
                      tabPanel(tags$span("📋 DETAILED MATRIX", style = "font-family: 'Inter', sans-serif; font-weight: 700;"), 
                               withSpinner(DT::dataTableOutput("singleStrainTable"), color = "#11998e"))
                    )
                  )
                ),
                
                fluidRow(
                  box(
                    title = tags$span(
                      icon("industry", style = "margin-right: 10px;"),
                      "PLANT CELL WALL DEGRADATION - INDUSTRIAL QUANTUM ASSESSMENT",
                      style = "font-family: 'Inter', sans-serif; font-weight: 700; text-transform: uppercase; letter-spacing: 0.5px;"
                    ),
                    status = "warning",
                    solidHeader = TRUE,
                    width = 12,
                    
                    htmlOutput("singleStrainDegradationSummary")
                  )
                )
              )
      ),
      
      # Multi-Strain Battle Tab
      tabItem(tabName = "multi_strain",
              conditionalPanel(
                condition = "output.dataUploaded",
                fluidRow(
                  box(
                    title = tags$span(
                      icon("users", style = "margin-right: 10px;"),
                      "MULTI-STRAIN BATTLE ARENA",
                      style = "font-family: 'Inter', sans-serif; font-weight: 700; text-transform: uppercase; letter-spacing: 0.5px;"
                    ),
                    status = "primary",
                    solidHeader = TRUE,
                    width = 4,
                    
                    checkboxGroupInput("selectedStrains", 
                                       tags$span("🦠 SELECT COMPETING STRAINS:", 
                                                 style = "font-family: 'Inter', sans-serif; font-weight: 700; color: #00d4ff;"),
                                       choices = NULL),
                    
                    selectInput("battleMetric", "Battle Metric:",
                                choices = list(
                                  "Overall Enzyme Count" = "total",
                                  "Cellulolytic Power" = "cellulolytic", 
                                  "Hemi-Cellulolytic" = "hemi_cellulolytic",
                                  "Lignolytic Potential" = "lignolytic"
                                )),
                    
                    actionButton("startBattle", "⚔️ INITIATE STRAIN BATTLE",
                                 class = "btn-danger btn-lg quantum-glow",
                                 style = "width: 100%; margin-top: 15px; font-family: 'Inter', sans-serif; 
                             font-weight: 700; text-transform: uppercase; letter-spacing: 0.5px;
                             background: linear-gradient(45deg, #ff4444, #ff6b35); border: none;")
                  ),
                  
                  box(
                    title = "BATTLE RESULTS",
                    status = "success",
                    solidHeader = TRUE,
                    width = 8,
                    
                    withSpinner(plotlyOutput("multiBattleChart", height = "450px"), color = "#11998e")
                  )
                ),
                
                fluidRow(
                  box(
                    title = "STRAIN COMPARISON MATRIX",
                    status = "info",
                    solidHeader = TRUE,
                    width = 12,
                    
                    withSpinner(DT::dataTableOutput("multiBattleTable"), color = "#3c8dbc")
                  )
                )
              )
      ),
      
      # Strain Ranking Tab  
      tabItem(tabName = "strain_ranking",
              conditionalPanel(
                condition = "output.dataUploaded",
                fluidRow(
                  box(
                    title = tags$span(
                      icon("trophy", style = "margin-right: 10px;"),
                      "STRAIN RANKING LEADERBOARD",
                      style = "font-family: 'Inter', sans-serif; font-weight: 700; text-transform: uppercase; letter-spacing: 0.5px;"
                    ),
                    status = "primary", 
                    solidHeader = TRUE,
                    width = 12,
                    
                    withSpinner(DT::dataTableOutput("strainRankingTable"), color = "#00d4ff")
                  )
                ),
                
                fluidRow(
                  box(
                    title = "TOP PERFORMERS VISUALIZATION",
                    status = "success",
                    solidHeader = TRUE,
                    width = 12,
                    
                    withSpinner(plotlyOutput("topPerformersChart", height = "500px"), color = "#11998e")
                  )
                )
              )
      ),
      
      # Enhanced Cellulolytic Potential Tab with Comprehensive Analysis
      tabItem(tabName = "cellulolytic",
              conditionalPanel(
                condition = "output.dataUploaded",
                # Enhanced value boxes
                fluidRow(
                  valueBoxOutput("cellulaseCount", width = 2),
                  valueBoxOutput("celluloseBMCount", width = 2),
                  valueBoxOutput("topCellulaseStrain", width = 2),
                  valueBoxOutput("cellulaseEfficiency", width = 2),
                  valueBoxOutput("endoglucanaseCount", width = 2),
                  valueBoxOutput("exoglucanaseCount", width = 2)
                ),
                
                fluidRow(
                  box(
                    title = tags$span(
                      icon("seedling", style = "margin-right: 10px;"),
                      "CELLULOLYTIC ENZYME QUANTUM MATRIX",
                      style = "font-family: 'Inter', sans-serif; font-weight: 700; text-transform: uppercase; letter-spacing: 0.5px;"
                    ),
                    status = "primary",
                    solidHeader = TRUE,
                    width = 8,
                    
                    div(class = "degradation-card quantum-glow",
                        h4("🌱 CELLULOSE DEGRADATION - COMPREHENSIVE ANALYSIS"),
                        div(style = "display: grid; grid-template-columns: 1fr 1fr 1fr; gap: 15px; margin-top: 15px;",
                            div(style = "background: rgba(0, 212, 255, 0.1); padding: 10px; border-radius: 8px; border-left: 3px solid #00d4ff;",
                                p(tags$strong("🔬 ENDOGLUCANASES:", style = "color: #00d4ff; font-size: 14px; font-family: 'Inter', sans-serif;"), br(),
                                  "GH5, GH8, GH9, GH12, GH44, GH45, GH74", style = "font-size: 12px; color: #cccccc; font-family: 'Inter', sans-serif;")
                            ),
                            div(style = "background: rgba(255, 107, 53, 0.1); padding: 10px; border-radius: 8px; border-left: 3px solid #ff6b35;",
                                p(tags$strong("⚡ EXOGLUCANASES:", style = "color: #ff6b35; font-size: 14px; font-family: 'Inter', sans-serif;"), br(),
                                  "GH6, GH7, GH48", style = "font-size: 12px; color: #cccccc; font-family: 'Inter', sans-serif;")
                            ),
                            div(style = "background: rgba(56, 239, 125, 0.1); padding: 10px; border-radius: 8px; border-left: 3px solid #38ef7d;",
                                p(tags$strong("🎯 β-GLUCOSIDASES:", style = "color: #38ef7d; font-size: 14px; font-family: 'Inter', sans-serif;"), br(),
                                  "GH1, GH3, GH116", style = "font-size: 12px; color: #cccccc; font-family: 'Inter', sans-serif;")
                            )
                        )
                    ),
                    
                    withSpinner(plotlyOutput("cellulaseDistribution", height = "400px"), color = "#11998e")
                  ),
                  
                  box(
                    title = tags$span(
                      icon("award", style = "margin-right: 10px;"),
                      "CELLULASE BIOREFINERY RANKING",
                      style = "font-family: 'Inter', sans-serif; font-weight: 700; text-transform: uppercase; letter-spacing: 0.5px;"
                    ),
                    status = "success",
                    solidHeader = TRUE,
                    width = 4,
                    
                    withSpinner(DT::dataTableOutput("cellulasePotentialTable"), color = "#11998e")
                  )
                ),
                
                fluidRow(
                  box(
                    title = tags$span(
                      icon("cogs", style = "margin-right: 10px;"),
                      "CELLULOLYTIC MECHANISM - INDUSTRIAL QUANTUM INSIGHTS",
                      style = "font-family: 'Inter', sans-serif; font-weight: 700; text-transform: uppercase; letter-spacing: 0.5px;"
                    ),
                    status = "info",
                    solidHeader = TRUE,
                    width = 6,
                    
                    htmlOutput("cellulaseInsights")
                  ),
                  
                  box(
                    title = tags$span(
                      icon("chart-pie", style = "margin-right: 10px;"),
                      "CELLULASE SYNERGY MATRIX",
                      style = "font-family: 'Inter', sans-serif; font-weight: 700; text-transform: uppercase; letter-spacing: 0.5px;"
                    ),
                    status = "warning",
                    solidHeader = TRUE,
                    width = 6,
                    
                    withSpinner(plotlyOutput("cellulaseSynergy", height = "400px"), color = "#f7931e")
                  )
                )
              )
      ),
      
      # Enhanced Hemi-Cellulolytic Potential Tab
      tabItem(tabName = "hemi_cellulolytic",
              conditionalPanel(
                condition = "output.dataUploaded",
                fluidRow(
                  valueBoxOutput("hemiCellulaseCount", width = 2),
                  valueBoxOutput("xylanaseCount", width = 2),
                  valueBoxOutput("mannanaseCount", width = 2),
                  valueBoxOutput("topHemiStrain", width = 2),
                  valueBoxOutput("hemiEfficiency", width = 2),
                  valueBoxOutput("hemiDiversity", width = 2)
                ),
                
                fluidRow(
                  box(
                    title = tags$span(
                      icon("leaf", style = "margin-right: 10px;"),
                      "HEMI-CELLULOLYTIC ENZYME ARSENAL",
                      style = "font-family: 'Inter', sans-serif; font-weight: 700; text-transform: uppercase; letter-spacing: 0.5px;"
                    ),
                    status = "primary",
                    solidHeader = TRUE,
                    width = 12,
                    
                    div(class = "degradation-card quantum-glow",
                        h4("🌿 HEMICELLULOSE DEGRADATION - COMPREHENSIVE MATRIX"),
                        div(style = "display: grid; grid-template-columns: 1fr 1fr 1fr 1fr; gap: 15px; margin-top: 15px;",
                            div(style = "background: rgba(0, 212, 255, 0.1); padding: 12px; border-radius: 8px; border-left: 3px solid #00d4ff;",
                                p(tags$strong("🔬 XYLANASES:", style = "color: #00d4ff; font-size: 13px; font-family: 'Inter', sans-serif;"), br(),
                                  "GH10, GH11, GH43", style = "font-size: 11px; color: #cccccc; font-family: 'Inter', sans-serif;")
                            ),
                            div(style = "background: rgba(255, 107, 53, 0.1); padding: 12px; border-radius: 8px; border-left: 3px solid #ff6b35;",
                                p(tags$strong("⚡ MANNANASES:", style = "color: #ff6b35; font-size: 13px; font-family: 'Inter', sans-serif;"), br(),
                                  "GH5, GH26, GH113", style = "font-size: 11px; color: #cccccc; font-family: 'Inter', sans-serif;")
                            ),
                            div(style = "background: rgba(56, 239, 125, 0.1); padding: 12px; border-radius: 8px; border-left: 3px solid #38ef7d;",
                                p(tags$strong("🎯 ARABINASES:", style = "color: #38ef7d; font-size: 13px; font-family: 'Inter', sans-serif;"), br(),
                                  "GH51, GH54, GH62", style = "font-size: 11px; color: #cccccc; font-family: 'Inter', sans-serif;")
                            ),
                            div(style = "background: rgba(156, 89, 209, 0.1); padding: 12px; border-radius: 8px; border-left: 3px solid #9c59d1;",
                                p(tags$strong("💜 ACCESSORY:", style = "color: #9c59d1; font-size: 13px; font-family: 'Inter', sans-serif;"), br(),
                                  "CE1-15, AA9, AA14", style = "font-size: 11px; color: #cccccc; font-family: 'Inter', sans-serif;")
                            )
                        )
                    ),
                    
                    withSpinner(plotlyOutput("hemiCellulaseDistribution", height = "400px"), color = "#11998e")
                  )
                )
              )
      ),
      
      # Enhanced Lignolytic Potential Tab
      tabItem(tabName = "lignolytic",
              conditionalPanel(
                condition = "output.dataUploaded",
                fluidRow(
                  valueBoxOutput("ligninaseCount", width = 2),
                  valueBoxOutput("peroxidaseCount", width = 2),
                  valueBoxOutput("laccaseCount", width = 2),
                  valueBoxOutput("topLigninStrain", width = 2),
                  valueBoxOutput("ligninEfficiency", width = 2),
                  valueBoxOutput("ligninDiversity", width = 2)
                ),
                
                fluidRow(
                  box(
                    title = tags$span(
                      icon("tree", style = "margin-right: 10px;"),
                      "LIGNOLYTIC ENZYME ARSENAL",
                      style = "font-family: 'Inter', sans-serif; font-weight: 700; text-transform: uppercase; letter-spacing: 0.5px;"
                    ),
                    status = "primary",
                    solidHeader = TRUE,
                    width = 12,
                    
                    div(class = "degradation-card quantum-glow",
                        h4("🌳 LIGNIN MODIFICATION - COMPREHENSIVE MATRIX"),
                        div(style = "display: grid; grid-template-columns: 1fr 1fr 1fr; gap: 15px; margin-top: 15px;",
                            div(style = "background: rgba(139, 69, 19, 0.1); padding: 12px; border-radius: 8px; border-left: 3px solid #8B4513;",
                                p(tags$strong("🔬 LACCASES:", style = "color: #8B4513; font-size: 13px; font-family: 'Inter', sans-serif;"), br(),
                                  "AA1 multi-copper oxidases", style = "font-size: 11px; color: #cccccc; font-family: 'Inter', sans-serif;")
                            ),
                            div(style = "background: rgba(205, 133, 63, 0.1); padding: 12px; border-radius: 8px; border-left: 3px solid #CD853F;",
                                p(tags$strong("⚡ PEROXIDASES:", style = "color: #CD853F; font-size: 13px; font-family: 'Inter', sans-serif;"), br(),
                                  "AA2, AA3 heme peroxidases", style = "font-size: 11px; color: #cccccc; font-family: 'Inter', sans-serif;")
                            ),
                            div(style = "background: rgba(160, 82, 45, 0.1); padding: 12px; border-radius: 8px; border-left: 3px solid #A0522D;",
                                p(tags$strong("🎯 GMC OXIDASES:", style = "color: #A0522D; font-size: 13px; font-family: 'Inter', sans-serif;"), br(),
                                  "AA3, AA5, AA8 dehydrogenases", style = "font-size: 11px; color: #cccccc; font-family: 'Inter', sans-serif;")
                            )
                        )
                    ),
                    
                    withSpinner(plotlyOutput("ligninaseDistribution", height = "400px"), color = "#8B4513")
                  )
                ),
                
                fluidRow(
                  box(
                    title = tags$span(
                      icon("cogs", style = "margin-right: 10px;"),
                      "LIGNOLYTIC MECHANISM - INDUSTRIAL QUANTUM INSIGHTS",
                      style = "font-family: 'Inter', sans-serif; font-weight: 700; text-transform: uppercase; letter-spacing: 0.5px;"
                    ),
                    status = "info",
                    solidHeader = TRUE,
                    width = 6,
                    
                    htmlOutput("ligninaseInsights")
                  ),
                  
                  box(
                    title = tags$span(
                      icon("award", style = "margin-right: 10px;"),
                      "LIGNIN DEGRADATION RANKING",
                      style = "font-family: 'Inter', sans-serif; font-weight: 700; text-transform: uppercase; letter-spacing: 0.5px;"
                    ),
                    status = "success",
                    solidHeader = TRUE,
                    width = 6,
                    
                    withSpinner(DT::dataTableOutput("ligninasePotentialTable"), color = "#8B4513")
                  )
                )
              )
      ),
      
      # Enhanced Pectinolytic Capacity Tab
      tabItem(tabName = "pectinolytic",
              conditionalPanel(
                condition = "output.dataUploaded",
                fluidRow(
                  valueBoxOutput("pectinaseCount", width = 2),
                  valueBoxOutput("polygalacturonaseCount", width = 2),
                  valueBoxOutput("pectinLyaseCount", width = 2),
                  valueBoxOutput("topPectinStrain", width = 2),
                  valueBoxOutput("pectinEfficiency", width = 2),
                  valueBoxOutput("pectinDiversity", width = 2)
                ),
                
                fluidRow(
                  box(
                    title = tags$span(
                      icon("apple-alt", style = "margin-right: 10px;"),
                      "PECTINOLYTIC ENZYME ARSENAL",
                      style = "font-family: 'Inter', sans-serif; font-weight: 700; text-transform: uppercase; letter-spacing: 0.5px;"
                    ),
                    status = "primary",
                    solidHeader = TRUE,
                    width = 12,
                    
                    div(class = "degradation-card quantum-glow",
                        h4("🍎 PECTIN DEGRADATION - COMPREHENSIVE MATRIX"),
                        div(style = "display: grid; grid-template-columns: 1fr 1fr 1fr 1fr; gap: 15px; margin-top: 15px;",
                            div(style = "background: rgba(255, 20, 147, 0.1); padding: 12px; border-radius: 8px; border-left: 3px solid #FF1493;",
                                p(tags$strong("🔬 POLYGALACTURONASES:", style = "color: #FF1493; font-size: 13px; font-family: 'Inter', sans-serif;"), br(),
                                  "GH28, GH33 hydrolases", style = "font-size: 11px; color: #cccccc; font-family: 'Inter', sans-serif;")
                            ),
                            div(style = "background: rgba(255, 105, 180, 0.1); padding: 12px; border-radius: 8px; border-left: 3px solid #FF69B4;",
                                p(tags$strong("⚡ PECTIN LYASES:", style = "color: #FF69B4; font-size: 13px; font-family: 'Inter', sans-serif;"), br(),
                                  "PL1, PL3, PL9 lyases", style = "font-size: 11px; color: #cccccc; font-family: 'Inter', sans-serif;")
                            ),
                            div(style = "background: rgba(255, 182, 193, 0.1); padding: 12px; border-radius: 8px; border-left: 3px solid #FFB6C1;",
                                p(tags$strong("🎯 PECTATE LYASES:", style = "color: #FFB6C1; font-size: 13px; font-family: 'Inter', sans-serif;"), br(),
                                  "PL1, PL2, PL3 eliminases", style = "font-size: 11px; color: #cccccc; font-family: 'Inter', sans-serif;")
                            ),
                            div(style = "background: rgba(219, 112, 147, 0.1); padding: 12px; border-radius: 8px; border-left: 3px solid #DB7093;",
                                p(tags$strong("💜 ACCESSORY:", style = "color: #DB7093; font-size: 13px; font-family: 'Inter', sans-serif;"), br(),
                                  "CE8, CE12 esterases", style = "font-size: 11px; color: #cccccc; font-family: 'Inter', sans-serif;")
                            )
                        )
                    ),
                    
                    withSpinner(plotlyOutput("pectinaseDistribution", height = "400px"), color = "#FF1493")
                  )
                )
              )
      ),
      
      # Enhanced Starch Degradation Tab
      tabItem(tabName = "starch_degrading",
              conditionalPanel(
                condition = "output.dataUploaded",
                fluidRow(
                  valueBoxOutput("amylaseCount", width = 2),
                  valueBoxOutput("glucoamylaseCount", width = 2),
                  valueBoxOutput("pullulanaseCount", width = 2),
                  valueBoxOutput("topStarchStrain", width = 2),
                  valueBoxOutput("starchEfficiency", width = 2),
                  valueBoxOutput("starchDiversity", width = 2)
                ),
                
                fluidRow(
                  box(
                    title = tags$span(
                      icon("seedling", style = "margin-right: 10px;"),
                      "STARCH DEGRADING ENZYME ARSENAL",
                      style = "font-family: 'Inter', sans-serif; font-weight: 700; text-transform: uppercase; letter-spacing: 0.5px;"
                    ),
                    status = "primary",
                    solidHeader = TRUE,
                    width = 12,
                    
                    div(class = "degradation-card quantum-glow",
                        h4("🌽 STARCH PROCESSING - COMPREHENSIVE MATRIX"),
                        div(style = "display: grid; grid-template-columns: 1fr 1fr 1fr 1fr; gap: 15px; margin-top: 15px;",
                            div(style = "background: rgba(255, 215, 0, 0.1); padding: 12px; border-radius: 8px; border-left: 3px solid #FFD700;",
                                p(tags$strong("🔬 α-AMYLASES:", style = "color: #FFD700; font-size: 13px; font-family: 'Inter', sans-serif;"), br(),
                                  "GH13 endoamylases", style = "font-size: 11px; color: #cccccc; font-family: 'Inter', sans-serif;")
                            ),
                            div(style = "background: rgba(255, 140, 0, 0.1); padding: 12px; border-radius: 8px; border-left: 3px solid #FF8C00;",
                                p(tags$strong("⚡ β-AMYLASES:", style = "color: #FF8C00; font-size: 13px; font-family: 'Inter', sans-serif;"), br(),
                                  "GH14 exoamylases", style = "font-size: 11px; color: #cccccc; font-family: 'Inter', sans-serif;")
                            ),
                            div(style = "background: rgba(255, 165, 0, 0.1); padding: 12px; border-radius: 8px; border-left: 3px solid #FFA500;",
                                p(tags$strong("🎯 GLUCOAMYLASES:", style = "color: #FFA500; font-size: 13px; font-family: 'Inter', sans-serif;"), br(),
                                  "GH15 exoglucosidases", style = "font-size: 11px; color: #cccccc; font-family: 'Inter', sans-serif;")
                            ),
                            div(style = "background: rgba(218, 165, 32, 0.1); padding: 12px; border-radius: 8px; border-left: 3px solid #DAA520;",
                                p(tags$strong("💫 DEBRANCHING:", style = "color: #DAA520; font-size: 13px; font-family: 'Inter', sans-serif;"), br(),
                                  "GH57, GH133 pullulanases", style = "font-size: 11px; color: #cccccc; font-family: 'Inter', sans-serif;")
                            )
                        )
                    ),
                    
                    withSpinner(plotlyOutput("starchDegradingDistribution", height = "400px"), color = "#FFD700")
                  )
                )
              )
      ),
      
      # Enhanced Inulin Processing Tab
      tabItem(tabName = "inulin_degrading",
              conditionalPanel(
                condition = "output.dataUploaded",
                fluidRow(
                  valueBoxOutput("inulinaseCount", width = 2),
                  valueBoxOutput("fructanaseCount", width = 2),
                  valueBoxOutput("levanaseCount", width = 2),
                  valueBoxOutput("topInulinStrain", width = 2),
                  valueBoxOutput("inulinEfficiency", width = 2),
                  valueBoxOutput("inulinDiversity", width = 2)
                ),
                
                fluidRow(
                  box(
                    title = tags$span(
                      icon("carrot", style = "margin-right: 10px;"),
                      "INULIN DEGRADING ENZYME ARSENAL",
                      style = "font-family: 'Inter', sans-serif; font-weight: 700; text-transform: uppercase; letter-spacing: 0.5px;"
                    ),
                    status = "primary",
                    solidHeader = TRUE,
                    width = 12,
                    
                    div(class = "degradation-card quantum-glow",
                        h4("🥔 INULIN PROCESSING - COMPREHENSIVE MATRIX"),
                        div(style = "display: grid; grid-template-columns: 1fr 1fr 1fr; gap: 15px; margin-top: 15px;",
                            div(style = "background: rgba(128, 0, 128, 0.1); padding: 12px; border-radius: 8px; border-left: 3px solid #800080;",
                                p(tags$strong("🔬 INULINASES:", style = "color: #800080; font-size: 13px; font-family: 'Inter', sans-serif;"), br(),
                                  "GH32 fructan hydrolases", style = "font-size: 11px; color: #cccccc; font-family: 'Inter', sans-serif;")
                            ),
                            div(style = "background: rgba(147, 112, 219, 0.1); padding: 12px; border-radius: 8px; border-left: 3px solid #9370DB;",
                                p(tags$strong("⚡ LEVANASES:", style = "color: #9370DB; font-size: 13px; font-family: 'Inter', sans-serif;"), br(),
                                  "GH32 levan degrading", style = "font-size: 11px; color: #cccccc; font-family: 'Inter', sans-serif;")
                            ),
                            div(style = "background: rgba(138, 43, 226, 0.1); padding: 12px; border-radius: 8px; border-left: 3px solid #8A2BE2;",
                                p(tags$strong("🎯 β-FRUCTOSIDASES:", style = "color: #8A2BE2; font-size: 13px; font-family: 'Inter', sans-serif;"), br(),
                                  "GH32 invertases", style = "font-size: 11px; color: #cccccc; font-family: 'Inter', sans-serif;")
                            )
                        )
                    ),
                    
                    withSpinner(plotlyOutput("inulinDegradingDistribution", height = "400px"), color = "#800080")
                  )
                )
              )
      ),
      
      # Biorefinery Optimizer Tab
      tabItem(tabName = "biorefinery",
              conditionalPanel(
                condition = "output.dataUploaded",
                fluidRow(
                  box(
                    title = tags$span(
                      icon("industry", style = "margin-right: 10px;"),
                      "BIOREFINERY STRAIN OPTIMIZATION CENTER",
                      style = "font-family: 'Inter', sans-serif; font-weight: 700; text-transform: uppercase; letter-spacing: 0.5px;"
                    ),
                    status = "primary",
                    solidHeader = TRUE,
                    width = 12,
                    
                    fluidRow(
                      column(4,
                             div(class = "biorefinery-card quantum-glow",
                                 uiOutput("enzymeFamilyAnalytics")  # This single line replaces everything!
                             )
                      ),
                      
                      column(8,
                             div(class = "metric-card",
                                 h4("🏆 BIOREFINERY STRAIN RECOMMENDATIONS"),
                                 htmlOutput("biorefineryRecommendations")
                             )
                      )
                    )
                  )
                ),
                
                fluidRow(
                  box(
                    title = "Optimal Strain Performance Matrix",
                    status = "success",
                    solidHeader = TRUE,
                    width = 8,
                    withSpinner(plotlyOutput("biorefineryMatrix", height = "500px"), color = "#00a65a")
                  ),
                  
                  box(
                    title = "Economic Viability Assessment",
                    status = "warning",
                    solidHeader = TRUE,
                    width = 4,
                    htmlOutput("economicAssessment")
                  )
                )
              )
      ),
      
      # Advanced Analytics Tab
      tabItem(tabName = "analytics",
              conditionalPanel(
                condition = "output.dataUploaded",
                fluidRow(
                  box(
                    title = "Advanced Statistical Analysis",
                    status = "primary",
                    solidHeader = TRUE,
                    width = 6,
                    withSpinner(plotlyOutput("correlationMatrix", height = "400px"), color = "#3c8dbc")
                  ),
                  
                  box(
                    title = "Principal Component Analysis",
                    status = "info",
                    solidHeader = TRUE,
                    width = 6,
                    withSpinner(plotlyOutput("pcaAnalysis", height = "400px"), color = "#17a2b8")
                  )
                ),
                
                fluidRow(
                  box(
                    title = "Machine Learning Predictions",
                    status = "success",
                    solidHeader = TRUE,
                    width = 12,
                    htmlOutput("mlPredictions")
                  )
                )
              )
      ),
      
      # ENHANCED UI CODE - Single Strain Degradation Pathway Network Tab
      tabItem(tabName = "pathway_network",
              conditionalPanel(
                condition = "output.dataUploaded",
                fluidRow(
                  box(
                    title = tags$span(
                      icon("project-diagram", style = "margin-right: 10px;"),
                      "SINGLE STRAIN PATHWAY ANALYZER",
                      style = "font-family: 'Inter', sans-serif; font-weight: 700; text-transform: uppercase; letter-spacing: 0.5px;"
                    ),
                    status = "primary",
                    solidHeader = TRUE,
                    width = 4,
                    
                    div(class = "pathway-card quantum-glow",
                        h4("🧬 PATHWAY NETWORK CONTROL CENTER"),
                        
                        selectInput("pathwaySelectedStrain", 
                                    tags$span("🦠 SELECT TARGET STRAIN:", 
                                              style = "font-family: 'Inter', sans-serif; font-weight: 700; color: #4facfe;"),
                                    choices = NULL,
                                    selected = NULL),
                        
                        div(class = "metric-card",
                            h5("🎯 PATHWAY CATEGORIES"),
                            div(style = "display: grid; grid-template-columns: 1fr 1fr; gap: 10px; margin-top: 10px;",
                                div(style = "background: rgba(0, 212, 255, 0.1); padding: 8px; border-radius: 6px; text-align: center;",
                                    tags$small("🌾 CELLULOSE", style = "color: #00d4ff; font-weight: 600; font-family: 'Inter', sans-serif;")
                                ),
                                div(style = "background: rgba(56, 239, 125, 0.1); padding: 8px; border-radius: 6px; text-align: center;",
                                    tags$small("🌿 HEMICELLULOSE", style = "color: #38ef7d; font-weight: 600; font-family: 'Inter', sans-serif;")
                                ),
                                div(style = "background: rgba(255, 107, 53, 0.1); padding: 8px; border-radius: 6px; text-align: center;",
                                    tags$small("🌳 LIGNIN", style = "color: #ff6b35; font-weight: 600; font-family: 'Inter', sans-serif;")
                                ),
                                div(style = "background: rgba(156, 89, 209, 0.1); padding: 8px; border-radius: 6px; text-align: center;",
                                    tags$small("🍎 PECTIN", style = "color: #9c59d1; font-weight: 600; font-family: 'Inter', sans-serif;")
                                ),
                                div(style = "background: rgba(247, 147, 30, 0.1); padding: 8px; border-radius: 6px; text-align: center;",
                                    tags$small("🌽 STARCH", style = "color: #f7931e; font-weight: 600; font-family: 'Inter', sans-serif;")
                                ),
                                div(style = "background: rgba(26, 188, 156, 0.1); padding: 8px; border-radius: 6px; text-align: center;",
                                    tags$small("🥔 INULIN", style = "color: #1abc9c; font-weight: 600; font-family: 'Inter', sans-serif;")
                                )
                            )
                        ),
                        
                        actionButton("generatePathwayNetwork", "🚀 GENERATE PATHWAY NETWORK",
                                     class = "btn-info btn-lg quantum-glow",
                                     style = "width: 100%; margin-top: 15px; font-family: 'Inter', sans-serif; 
                               font-weight: 700; text-transform: uppercase; letter-spacing: 0.5px;
                               background: linear-gradient(45deg, #4facfe, #00f2fe); border: none;"),
                        
                        br(),
                        div(class = "metric-card",
                            h5("📊 PATHWAY STATISTICS"),
                            htmlOutput("pathwayStats")
                        )
                    )
                  ),
                  
                  box(
                    title = tags$span(
                      icon("network-wired", style = "margin-right: 10px;"),
                      "DEGRADATION PATHWAY QUANTUM NETWORK",
                      style = "font-family: 'Inter', sans-serif; font-weight: 700; text-transform: uppercase; letter-spacing: 0.5px;"
                    ),
                    status = "info",
                    solidHeader = TRUE,
                    width = 8,
                    
                    withSpinner(plotlyOutput("singleStrainPathway", height = "750px"), color = "#4facfe")  # Increased height
                  )
                ),
                
                fluidRow(
                  box(
                    title = tags$span(
                      icon("microscope", style = "margin-right: 10px;"),
                      "INDUSTRIAL PATHWAY ASSESSMENT",
                      style = "font-family: 'Inter', sans-serif; font-weight: 700; text-transform: uppercase; letter-spacing: 0.5px;"
                    ),
                    status = "warning",
                    solidHeader = TRUE,
                    width = 12,
                    
                    div(class = "pathway-card",
                        h4("🏭 BIOREFINERY PATHWAY POTENTIAL"),
                        htmlOutput("pathwayIndustrialAssessment")
                    )
                  )
                ),
                
                fluidRow(
                  box(
                    title = tags$span(
                      icon("table", style = "margin-right: 10px;"),
                      "DETAILED PATHWAY ENZYME MATRIX",
                      style = "font-family: 'Inter', sans-serif; font-weight: 700; text-transform: uppercase; letter-spacing: 0.5px;"
                    ),
                    status = "primary",
                    solidHeader = TRUE,
                    width = 12,
                    
                    withSpinner(DT::dataTableOutput("pathwayEnzymeTable"), color = "#3c8dbc")
                  )
                )
              ),
              
              conditionalPanel(
                condition = "!output.dataUploaded",
                fluidRow(
                  box(
                    title = tags$span(
                      icon("project-diagram", style = "margin-right: 10px; color: #4facfe;"),
                      "PATHWAY NETWORK ANALYSIS CENTER",
                      style = "font-family: 'Inter', sans-serif; font-weight: 700; text-transform: uppercase; letter-spacing: 0.5px;"
                    ),
                    status = "info",
                    solidHeader = TRUE,
                    width = 12,
                    height = "500px",
                    div(class = "welcome-screen",
                        icon("project-diagram", style = "font-size: 6em; color: #4facfe; margin-bottom: 30px; 
                             text-shadow: 0 0 50px rgba(79, 172, 254, 0.8);"),
                        h2("🧬 SINGLE STRAIN DEGRADATION PATHWAY NETWORK", 
                           style = "background: linear-gradient(45deg, #4facfe, #00f2fe); 
                      -webkit-background-clip: text; -webkit-text-fill-color: transparent;
                      font-family: 'Inter', sans-serif; font-weight: 700;"),
                        h4("Visualize comprehensive enzyme degradation pathways for individual bacterial strains", 
                           style = "font-size: 20px; margin-bottom: 30px; color: #cccccc; font-family: 'Inter', sans-serif;"),
                        div(style = "display: grid; grid-template-columns: 1fr 1fr; gap: 30px; margin: 40px 0;",
                            div(class = "pathway-card",
                                icon("sitemap", style = "font-size: 2.5em; color: #4facfe;"),
                                h5("Network Visualization", style = "color: #4facfe; font-weight: 700; font-family: 'Inter', sans-serif;"),
                                p("Interactive pathway networks showing enzyme relationships", style = "font-family: 'Inter', sans-serif;")
                            ),
                            div(class = "pathway-card",
                                icon("cogs", style = "font-size: 2.5em; color: #00f2fe;"),
                                h5("Industrial Assessment", style = "color: #00f2fe; font-weight: 700; font-family: 'Inter', sans-serif;"),
                                p("Biorefinery potential analysis for strain optimization", style = "font-family: 'Inter', sans-serif;")
                            )
                        ),
                        p("Please upload your CAZy annotation data to access pathway network analysis.", 
                          style = "font-size: 18px; color: #ff6b35; font-family: 'Inter', sans-serif; font-weight: 500;")
                    )
                  )
                )
              )
      )
    )
  )
      
)
```
