## Supplementary Material for "CAZyXplorer: A Shiny Application for Cost-Effective Preliminary Screening of Microbial Strains to Advance Enzyme Discovery in Biorefining and Biotechnology"

**Data Retrieval**: CAZyme annotation data can be obtained from multiple sources including: (1) JGI MycoCosm web repository (<https://mycocosm.jgi.doe.gov/>) for fungal genomes, (2) JGI IMG (Integrated Microbial Genomes) database for bacterial genomes, or (3) custom annotations generated from genome sequences using the dbCAN annotation pipeline (<https://bcb.unl.edu/dbCAN2/>). The CAZyXplorer platform accepts CAZyme annotation data from any of these sources in CSV format, provided the data is formatted with CAZyme family annotations in the first column and genome-specific enzyme counts in subsequent columns. An example data retrieval workflow from JGI MycoCosm is provided in the Supporting Materials.

Preparation of Input file for loading the data in **CAZyXplorer**

1. The CAZymes data can be downloaded (or) retrieved from MycoCosm <https://mycocosm.jgi.doe.gov/mycocosm/home> web repository. For the Present tutorial I have retrieved the CAZy annotations for the subdivision *Agaricomycotina* fungi.
2. JGI-Mycocosm (webpage) 🡪 Agaricomycotina 🡪 Tree 🡪 click on 585 genomes 🡪 Annotations 🡪 CAZYMES 🡪 copy the data to a excel file 🡪 Replace empty values with 0.
3. Remove consolidated CAZy (gene counts) in the first row, also the group level consolidated gene counts for the GH, GT, PL, CE, CBM, AA for reflecting correct CAZy based analysis and analysis of microbial strains.
4. Similarly, the CAZyme data can be downloaded from the CAZy database <https://www.cazy.org/b.html> (Genomes section) prepare it as a CSV format to process it in CAZyXplorer

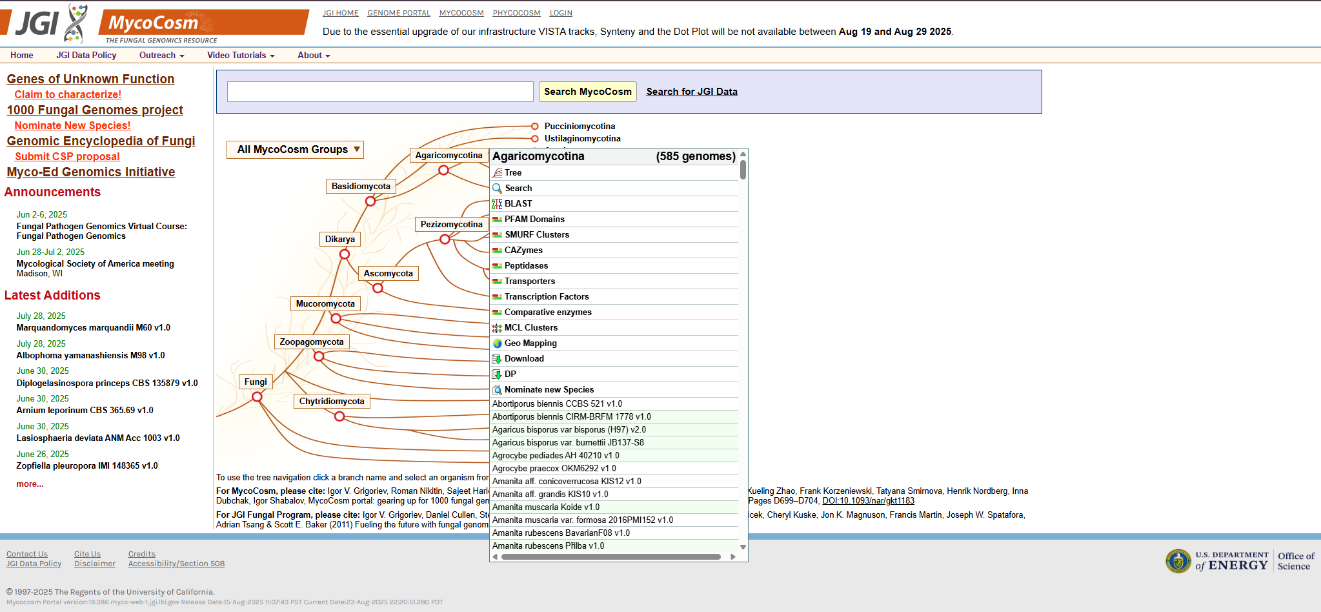

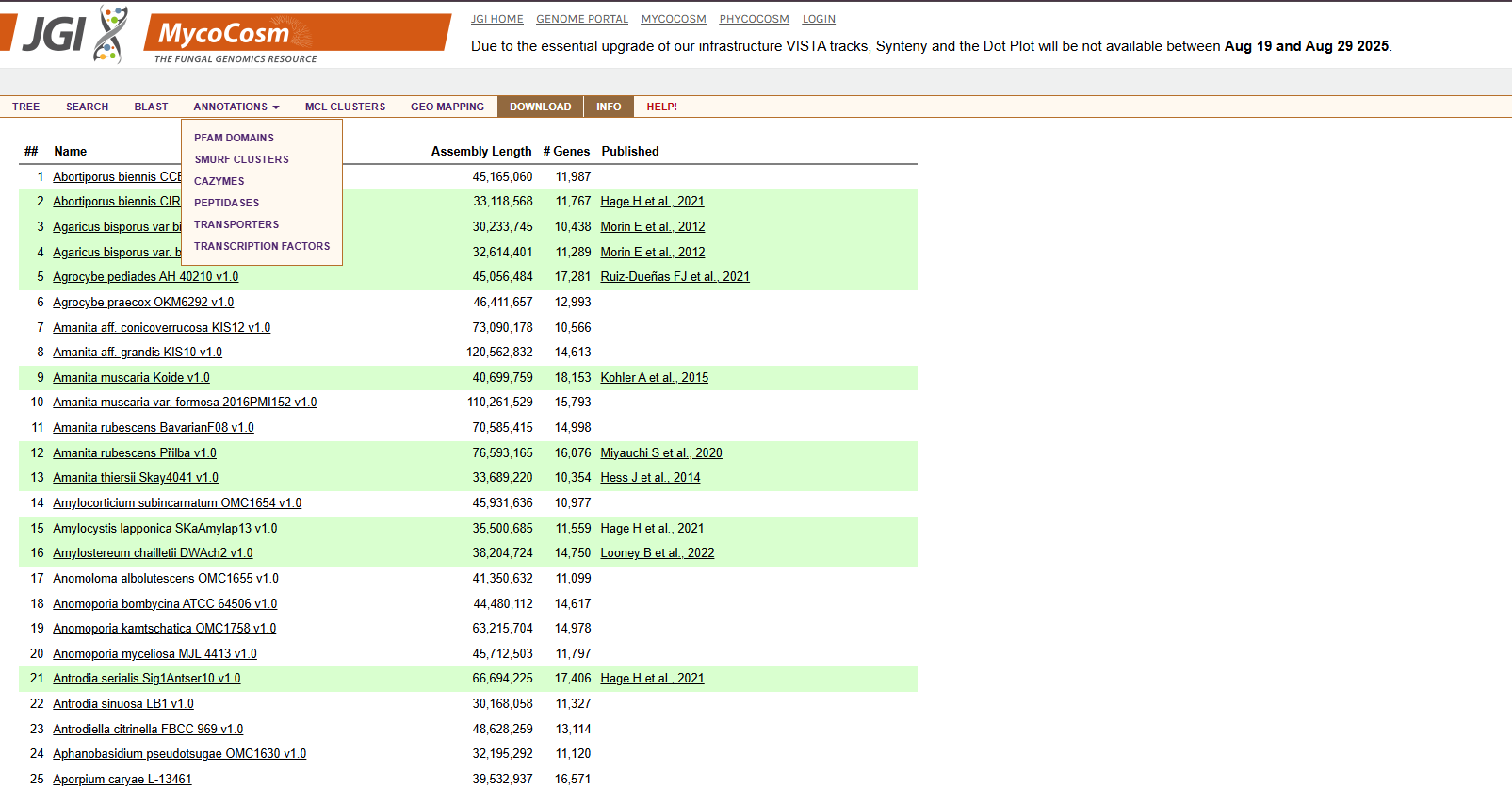

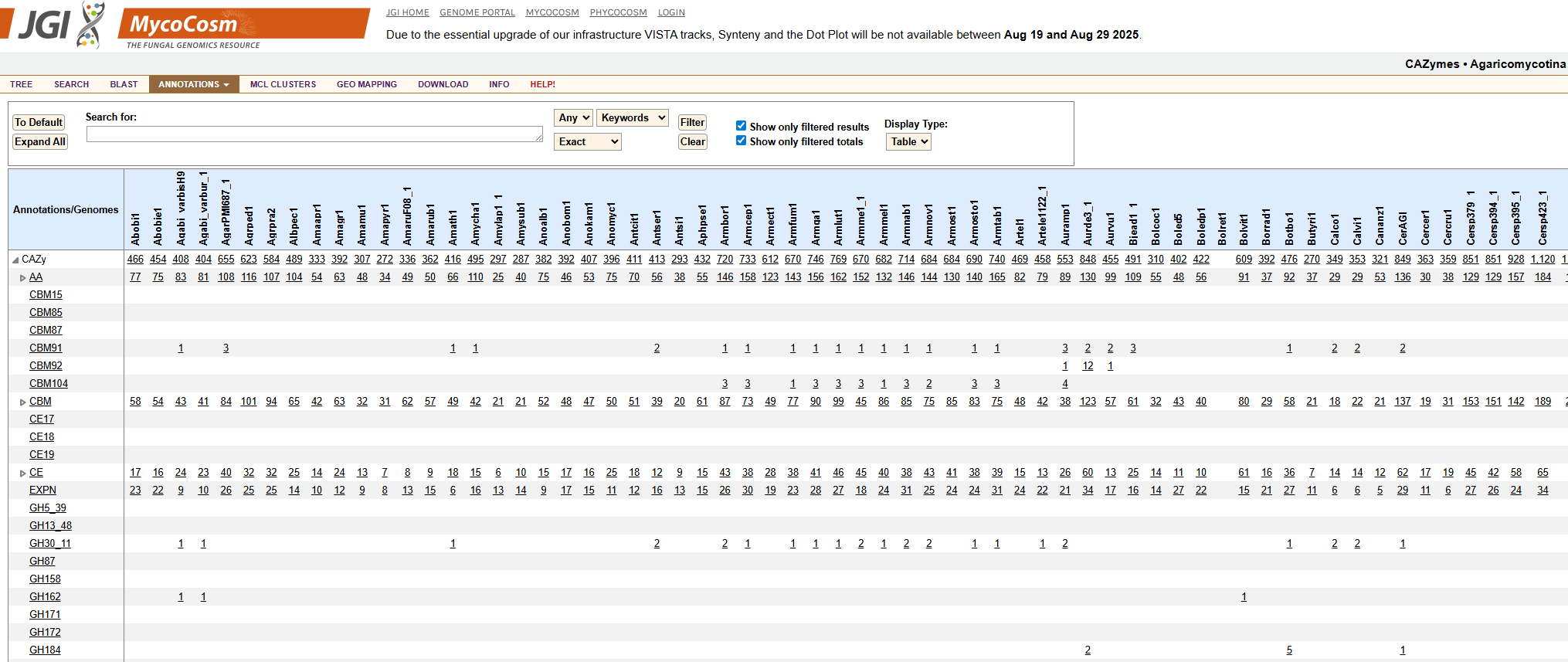

**A Step-by-Step Guide to Using CAZyXplorer**

**A. Preparation**

**Reading Assignment**

Prior to conducting the analysis, users are encouraged to review the following articles to familiarize themselves with carbohydrate-active enzymes and biorefinery applications:

1. **The carbohydrate-active enzymes database (CAZy) in 2013.** Lombard V et al. Nucleic Acids Res. 2014;42(D1):D490-5. DOI: 10.1093/nar/gkt1178
2. **Carbohydrate-active enzymes: structure, function and applications.** Davies G and Henrissat B. Biochem J. 1995;321(2):557-9.
3. **Fungal lignocellulose degradation: fundamentals and applications.** van den Brink J and de Vries RP. Microbiol Mol Biol Rev. 2011;75(2):259-77.

**Downloading and Installing the App**

1. Install R (https://www.r-project.org/) and RStudio (https://posit.co/download/rstudio-desktop/)
2. Download CAZyXplorer from the provided source or GitHub repository
3. Open the CAZyXplorer.R file with RStudio and start CAZyXplorer to install all necessary packages by clicking the “Run App” button
4. Install all necessary packages when prompted (required only on first use)

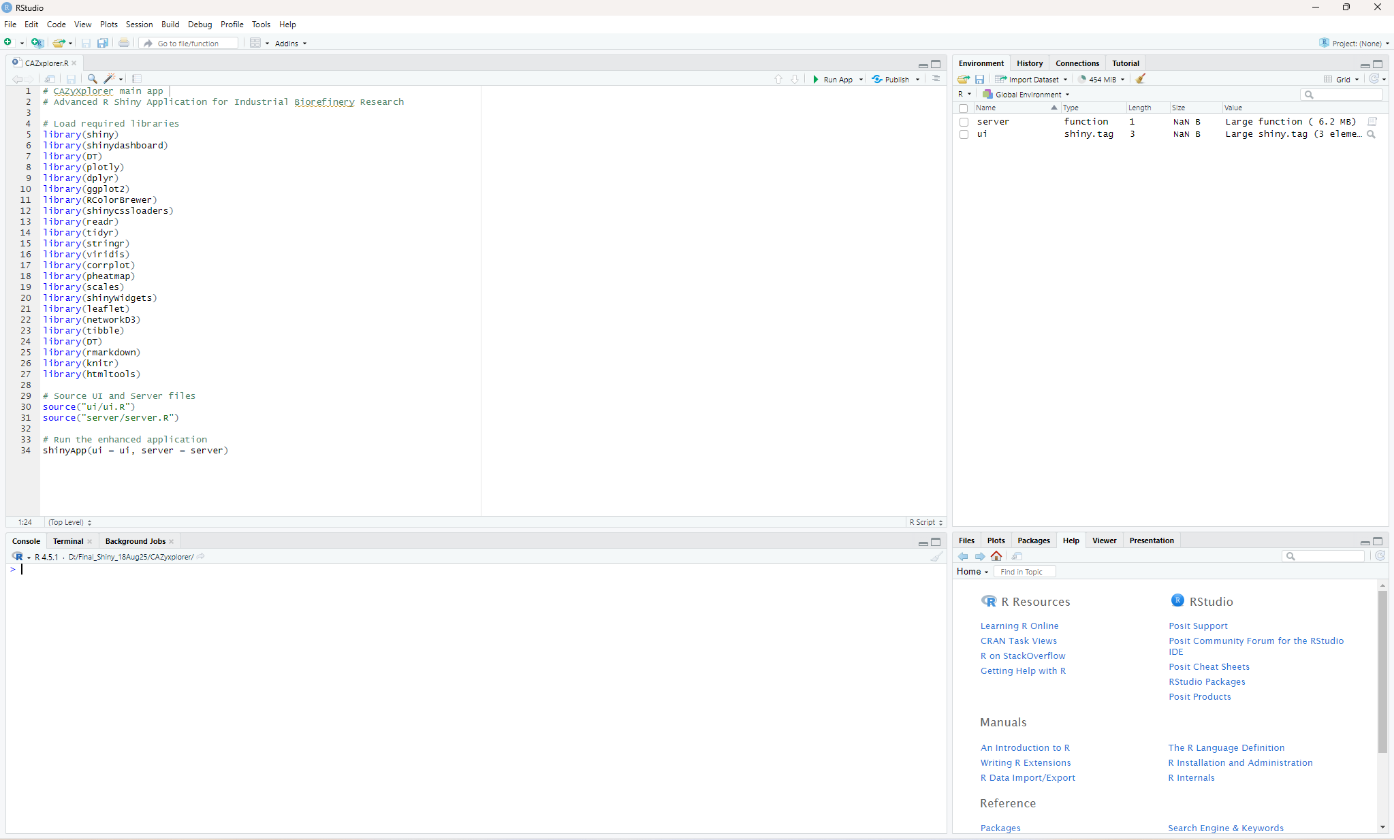

Click here to start CAZyXplorer

**Preparing Your CAZy Annotation Data**

**File Requirements:**

- **Format:** .csv (comma-separated values)
- **Structure:** Matrix format with:
  - First column: CAZy family annotations (e.g., GH1, AA3, CE4, PL1, CBM6)
  - Subsequent columns: Individual genome/strain names
  - Cell values: Copy numbers of each enzyme family per genome

**Data Quality Checklist:**

1. Ensure column names contain no spaces or special characters
2. CAZy family names must follow standard nomenclature (GH, AA, PL, CE, CBM, GT)
3. All values should be numeric (enzyme copy counts)
4. Missing values will be automatically converted to zero

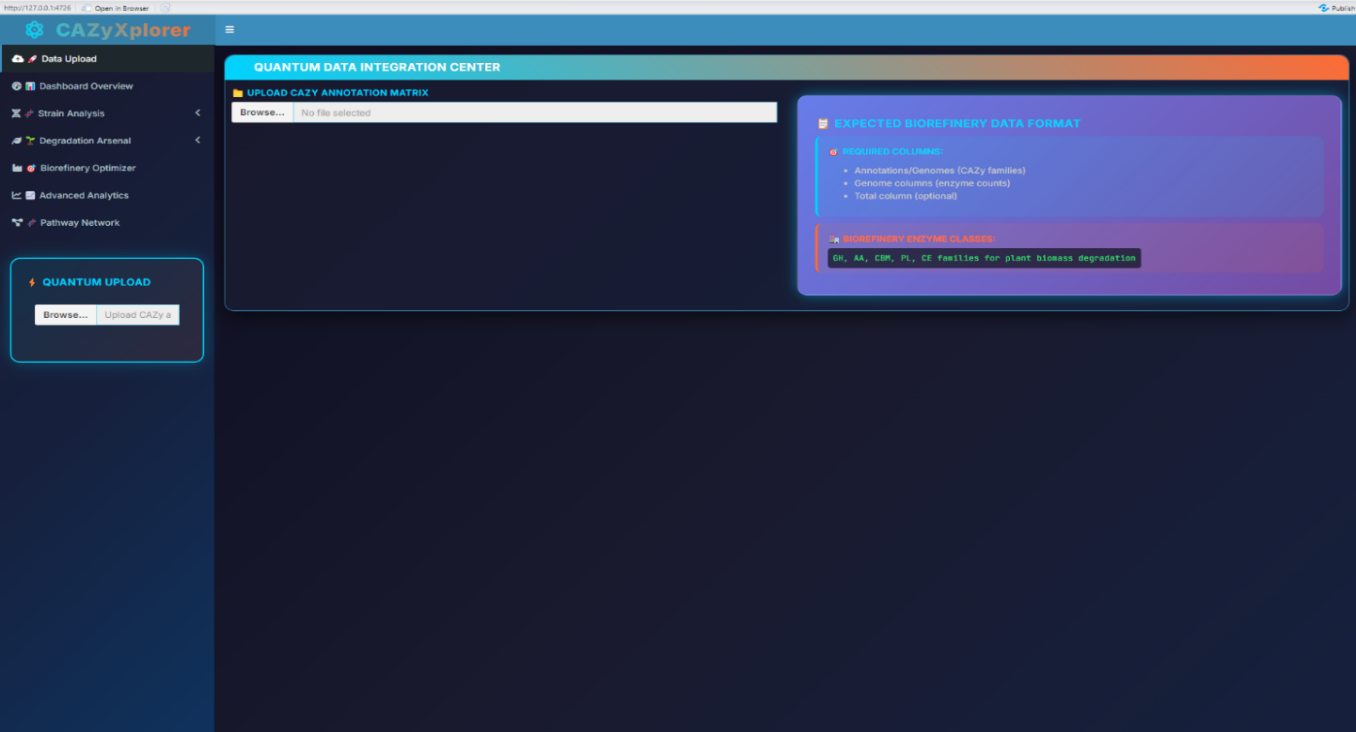

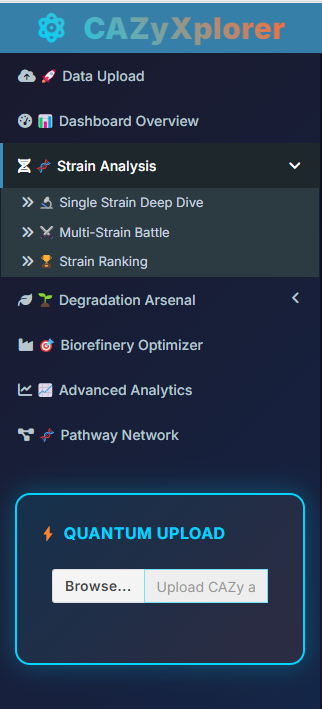

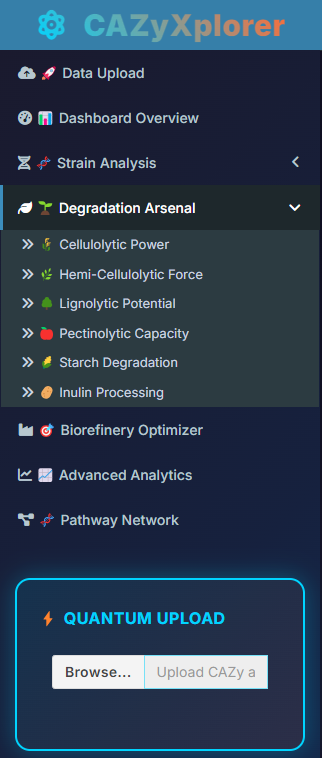

A

B

C

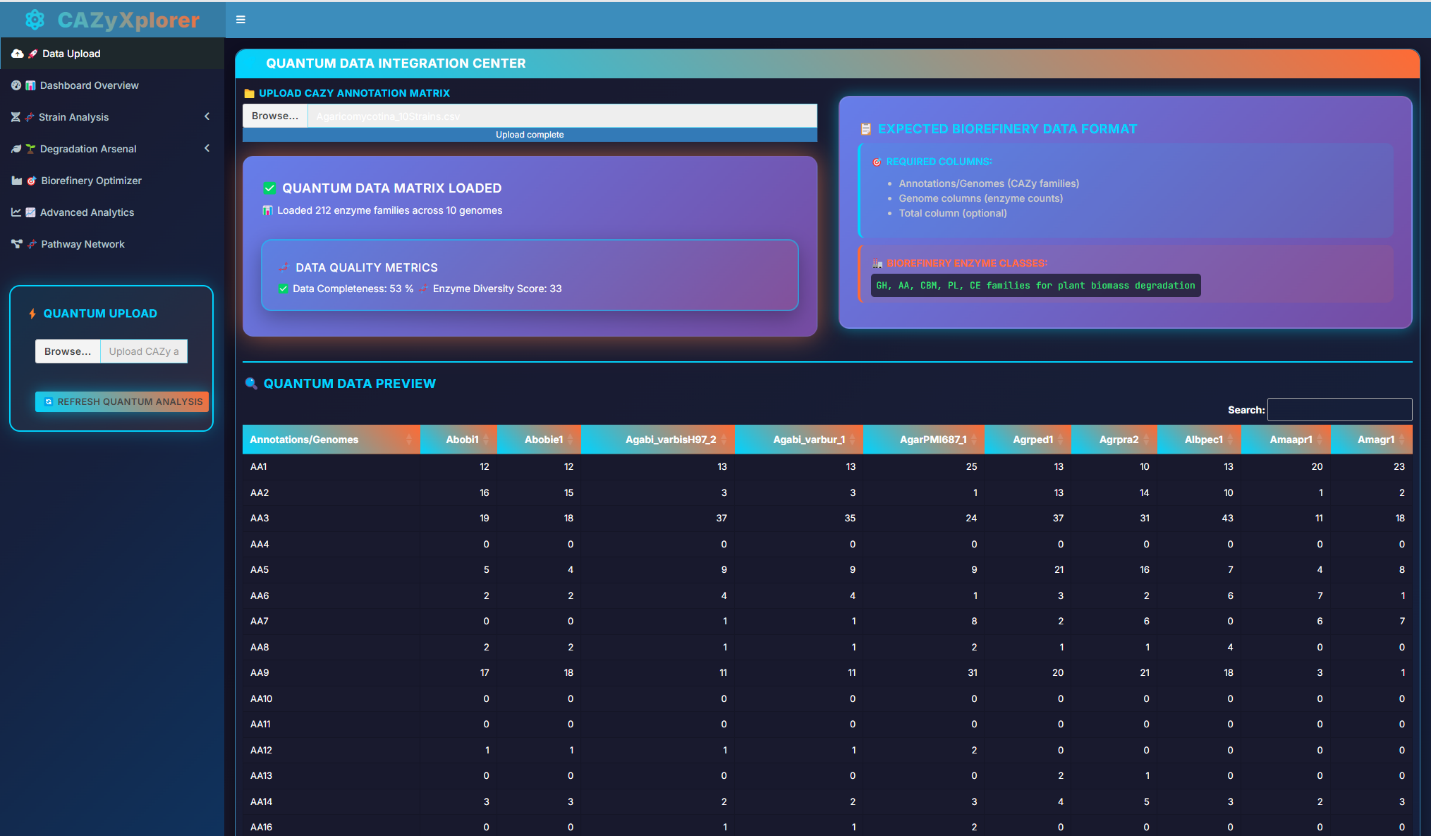

**B. Analytical Workflow**

**1. Data Upload and Validation**

**Step 1.1: Navigate to Data Upload**

- Launch CAZyXplorer
- Click on "Data Upload" in the sidebar menu
- You will see the "QUANTUM DATA INTEGRATION CENTER"

**Step 1.2: Upload Your CAZy Matrix**

- Click "Browse" under "UPLOAD CAZy ANNOTATION MATRIX"
- Select your prepared .csv file
- Wait for the upload confirmation message

**Step 1.3: Data Quality Assessment** After successful upload, CAZyXplorer will display:

- **Data Quality Metrics:** Overview of your dataset completeness
- **Data Preview:** Interactive table showing your uploaded data
- **Genome Count:** Total number of analyzed strains
- **Enzyme Family Count:** Total CAZy families detected

**2. Dashboard Overview - Initial Data Exploration**

**Step 2.1: Navigate to Dashboard Overview**

- Click "Dashboard Overview" in the sidebar
- View the comprehensive summary metrics:
  - Total Enzymes Detected
  - Total Genomes Analyzed
  - Average Enzymes Per Genome
  - Top Enzyme Family
  - Biodiversity Index
  - Industrial Potential Score

**Step 2.2: Explore Distribution Patterns**

- **Enzyme Distribution Matrix:** Stacked bar chart showing enzyme family distribution across genomes
- **Elite Enzyme Arsenal:** Top 15 most abundant enzyme families
- **Industrial Biorefinery Heatmap:** Comprehensive enzyme abundance visualization
- **Degradation Pathway Network:** Force-directed network showing enzyme relationships

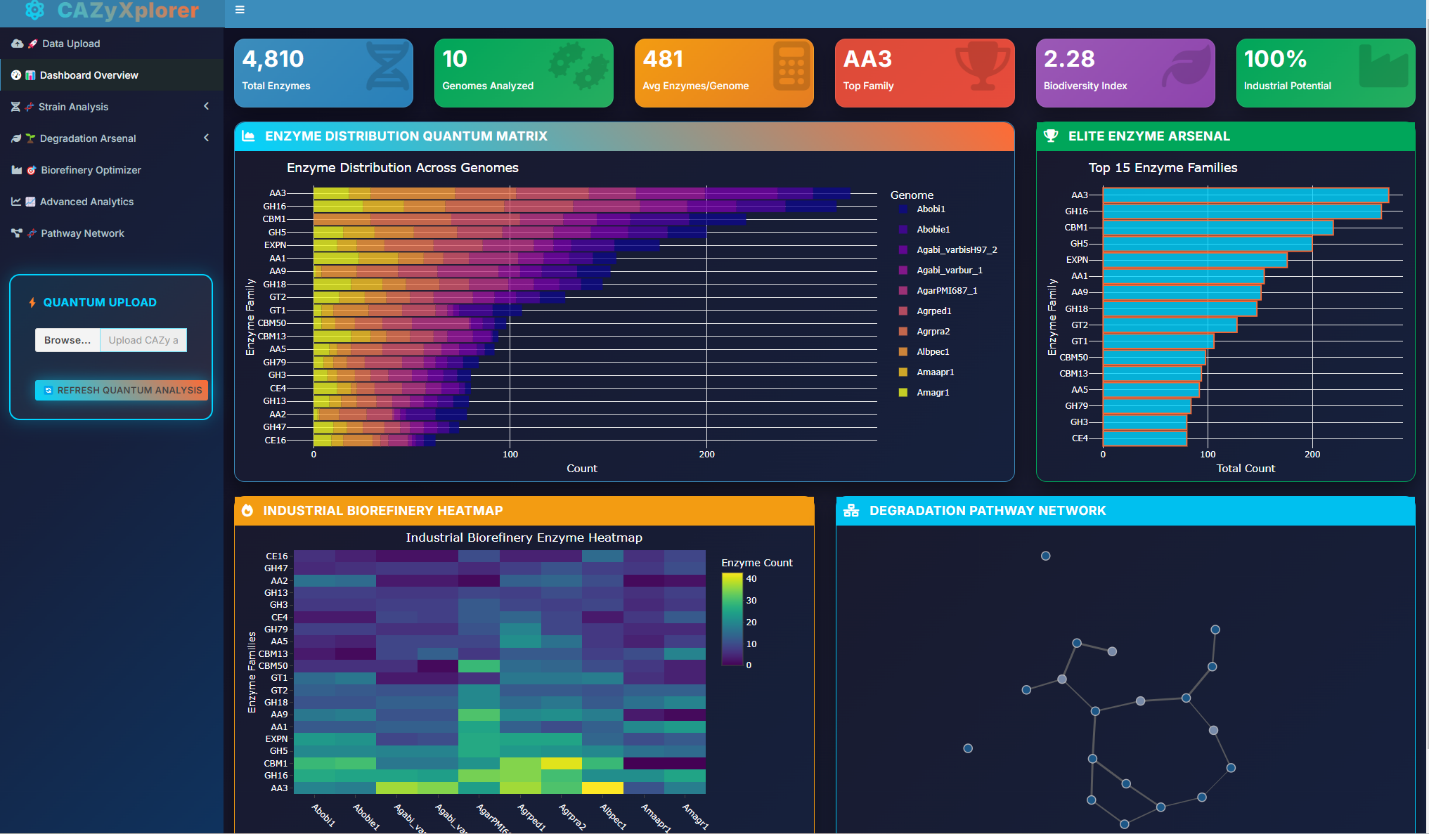

**3. Single Strain Deep Dive Analysis**

**Step 3.1: Select Target Strain**

- Navigate to "Strain Analysis" → "Single Strain Deep Dive"
- Use the dropdown menu to select your strain of interest
- View the real-time strain bioactivity matrix

**Step 3.2: Initiate Deep Analysis**

- Click "INITIATE DEEP QUANTUM SCAN"
- Explore multiple visualization tabs:
  - **Enzyme Distribution:** Bar chart of enzyme family abundance
  - **Degradation Arsenal:** Radar chart of degradation capabilities
  - **Enzyme Classes:** Pie chart of CAZy class distribution
  - **Detailed Matrix:** Complete enzyme inventory table

**Step 3.3: Industrial Assessment** Review the "Plant Cell Wall Degradation" assessment showing:

- Cellulolytic potential and efficiency rating
- Hemi-cellulolytic capabilities
- Ligninolytic activity levels
- Pectinolytic and starch degradation capacity

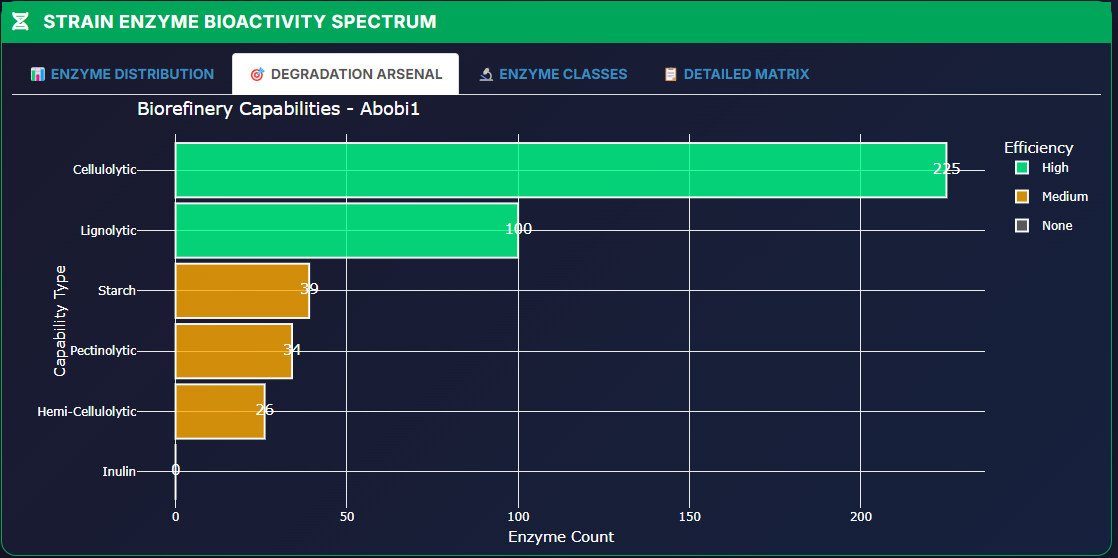

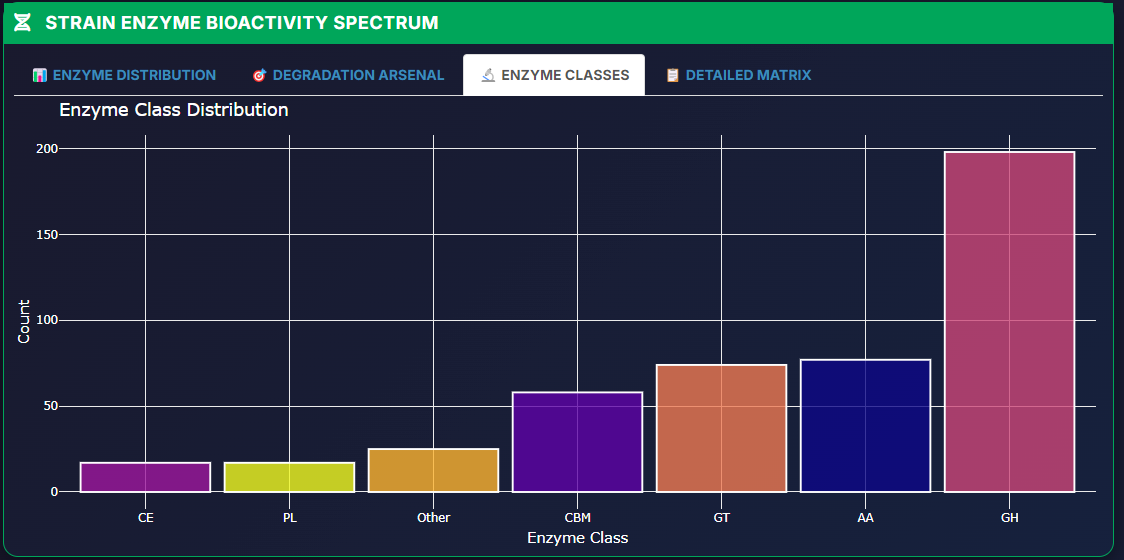

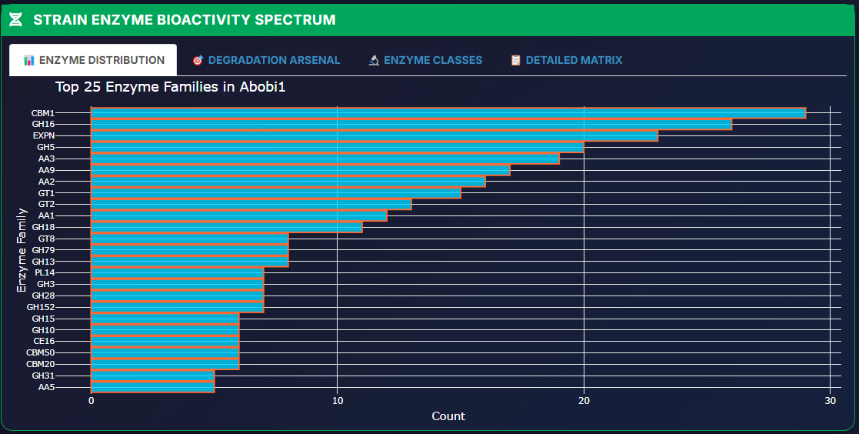

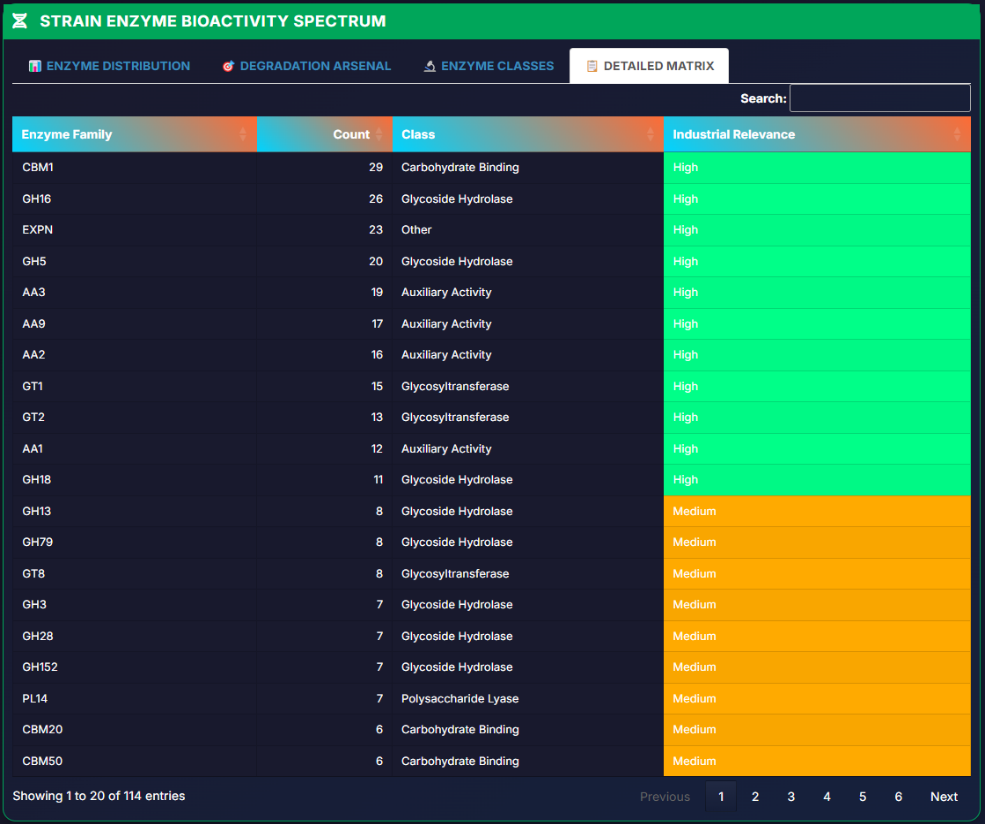

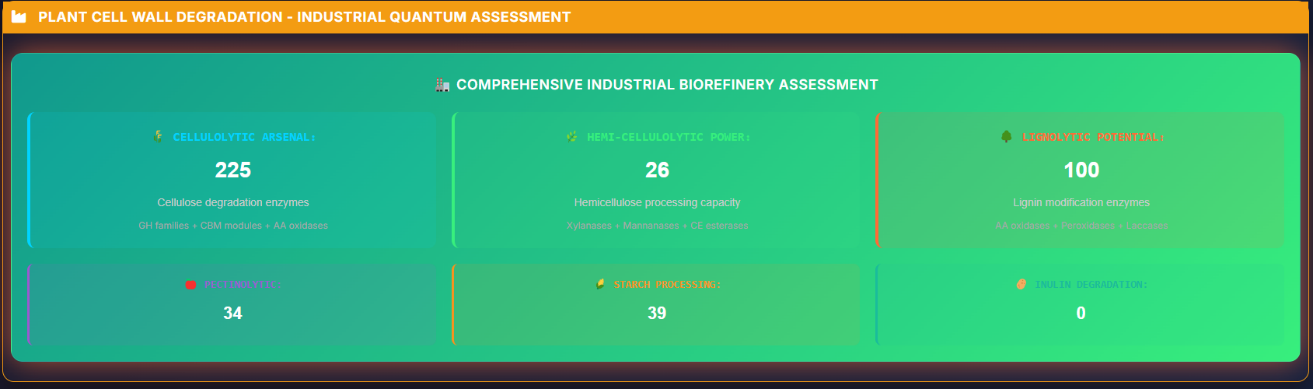

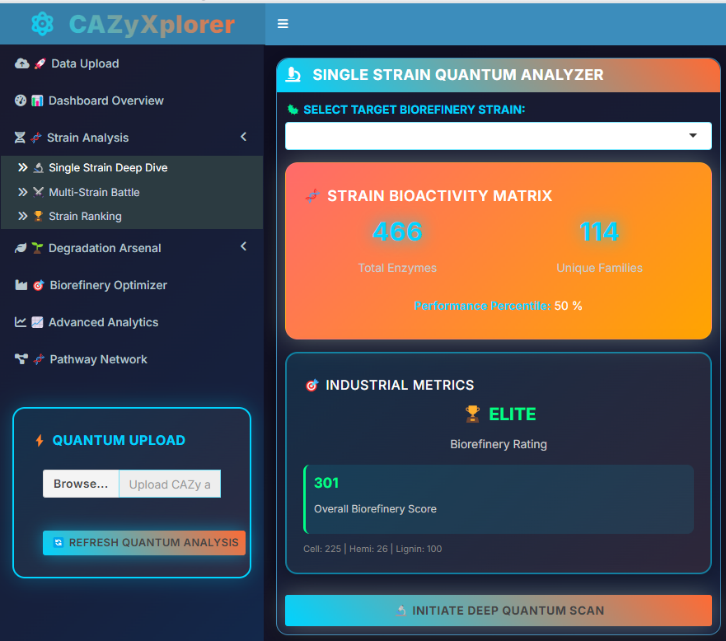

**4. Multi-Strain Comparative Analysis**

**Step 4.1: Strain Battle Arena**

- Navigate to "Strain Analysis" → "Multi-Strain Battle"
- Select 2-8 competing strains using checkboxes
- Choose battle metric (Overall, Cellulolytic, Hemi-cellulolytic, or Lignolytic)
- Click "INITIATE STRAIN BATTLE"

**Step 4.2: Battle Results Interpretation**

- **Battle Chart:** Visual comparison of selected strains
- **Comparison Matrix:** Detailed numerical comparison table
- Identify superior performers for specific applications

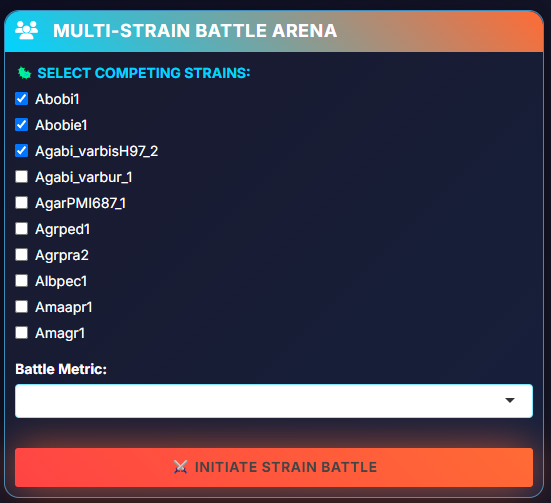

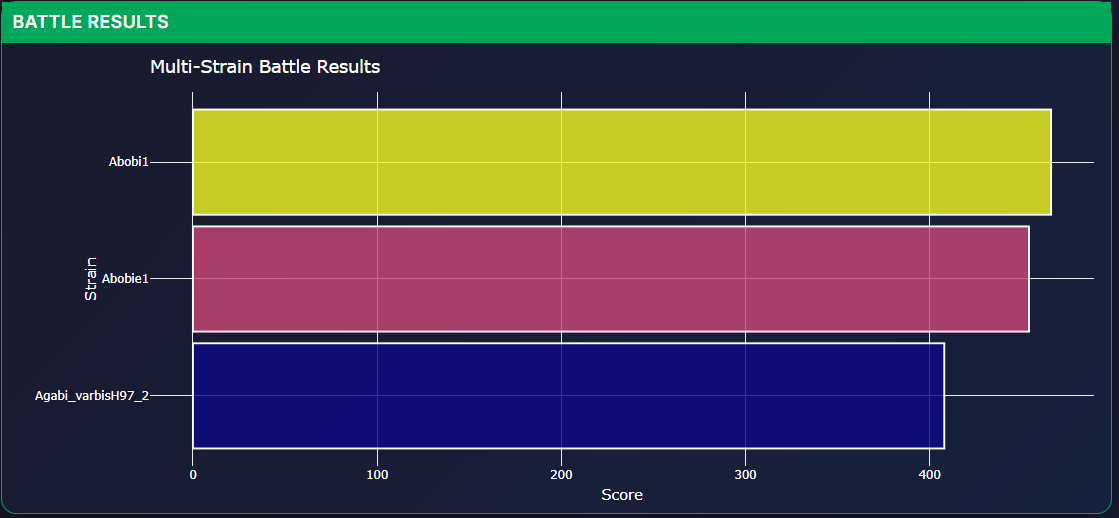

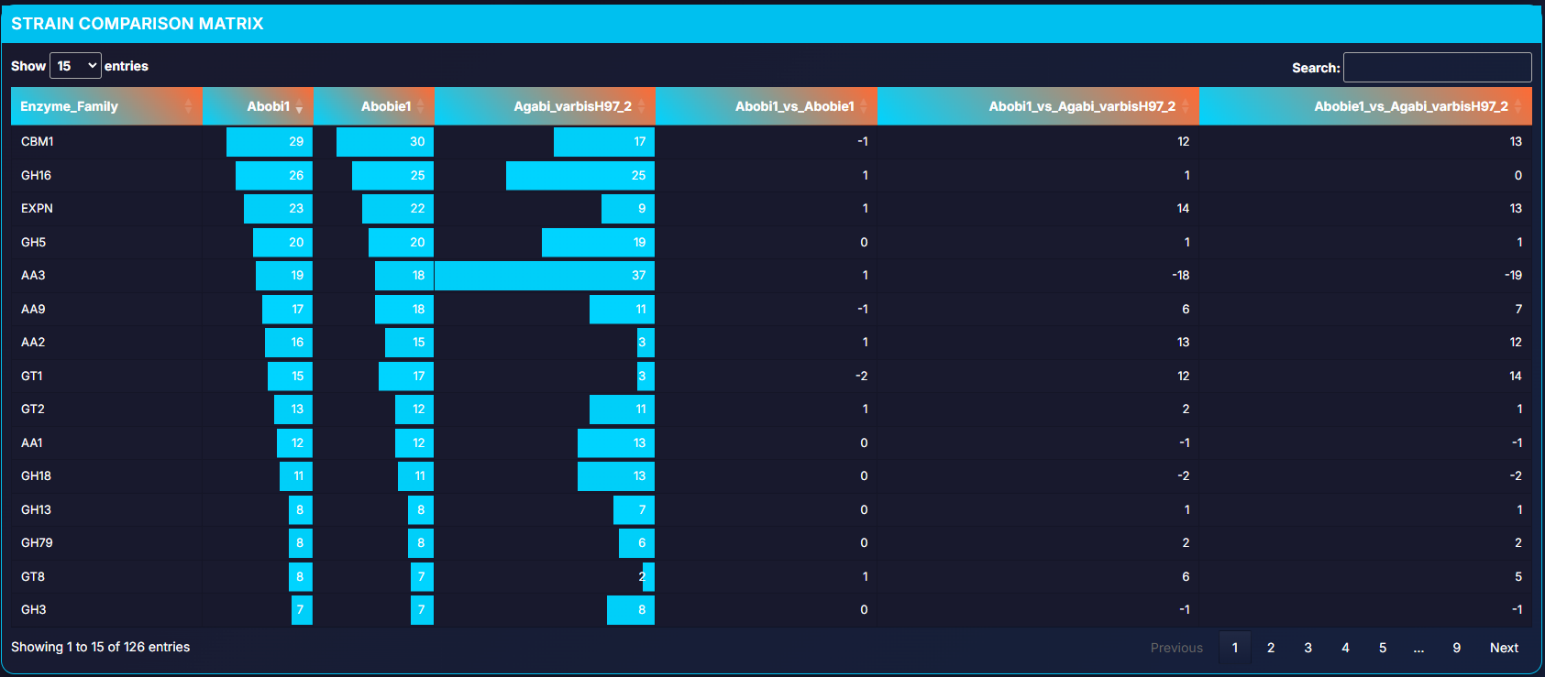

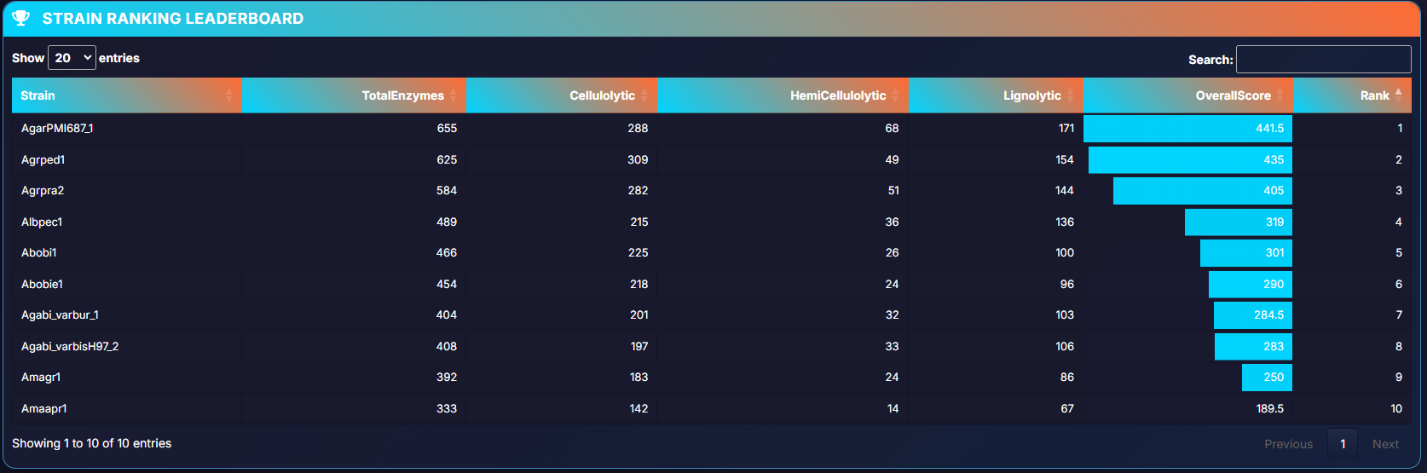

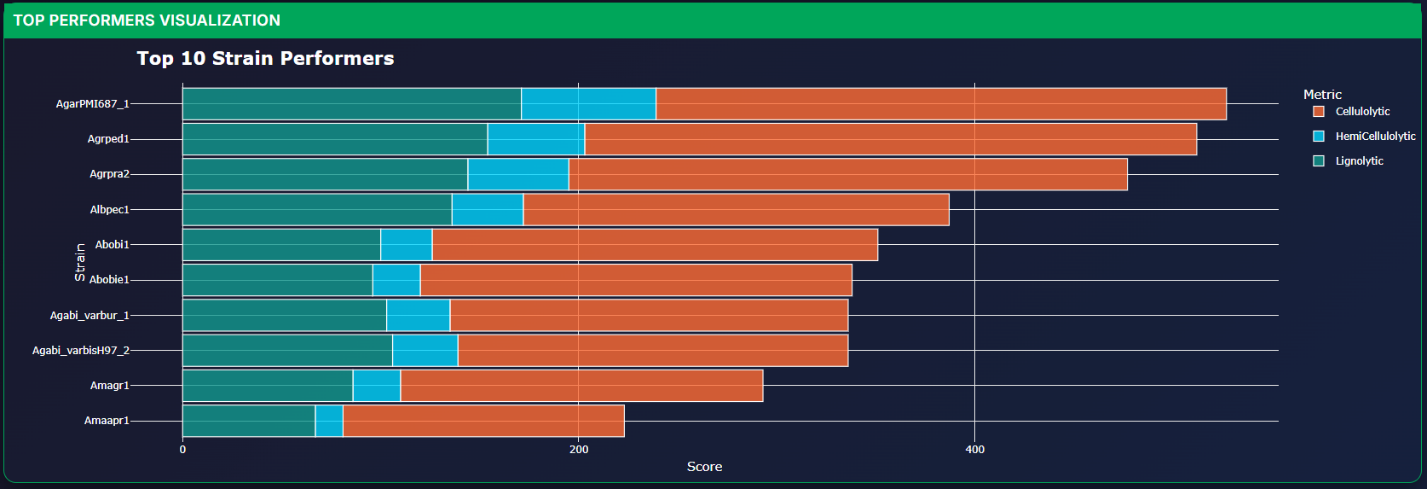

**5. Strain Ranking and Performance Assessment**

**Step 5.1: Comprehensive Ranking**

- Navigate to "Strain Analysis" → "Strain Ranking"
- View the complete leaderboard of all analyzed strains
- Rankings include:
  - Overall enzymatic performance scores
  - Pathway-specific rankings
  - Industrial potential assessments

**Step 5.2: Top Performers Visualization**

- Interactive charts showing performance hierarchies
- Filter by specific degradation pathways
- Export rankings for further analysis

**6. Degradation Arsenal Analysis**

**Step 6.1: Pathway-Specific Analysis** Navigate to "Degradation Arsenal" and explore each pathway:

**Cellulolytic Power:**

- Comprehensive cellulase analysis (endoglucanases, exoglucanases, β-glucosidases)
- Cellulase synergy matrix
- Industrial cellulase ranking

**Hemi-Cellulolytic Force:**

- Xylanase, mannanase, and arabinase profiles
- Accessory enzyme analysis
- Hemicellulose degradation efficiency

**Lignolytic Potential:**

- Laccase, peroxidase, and GMC oxidase analysis
- Lignin modification mechanisms
- White-rot vs brown-rot capabilities

**Pectinolytic Capacity:**

- Polygalacturonase and pectin lyase profiles
- Pectin degradation pathway analysis

**Starch Degradation:**

- α-amylase, β-amylase, and glucoamylase analysis
- Debranching enzyme capabilities

**Inulin Processing:**

- Inulinase and fructanase analysis
- Fructan degradation pathways

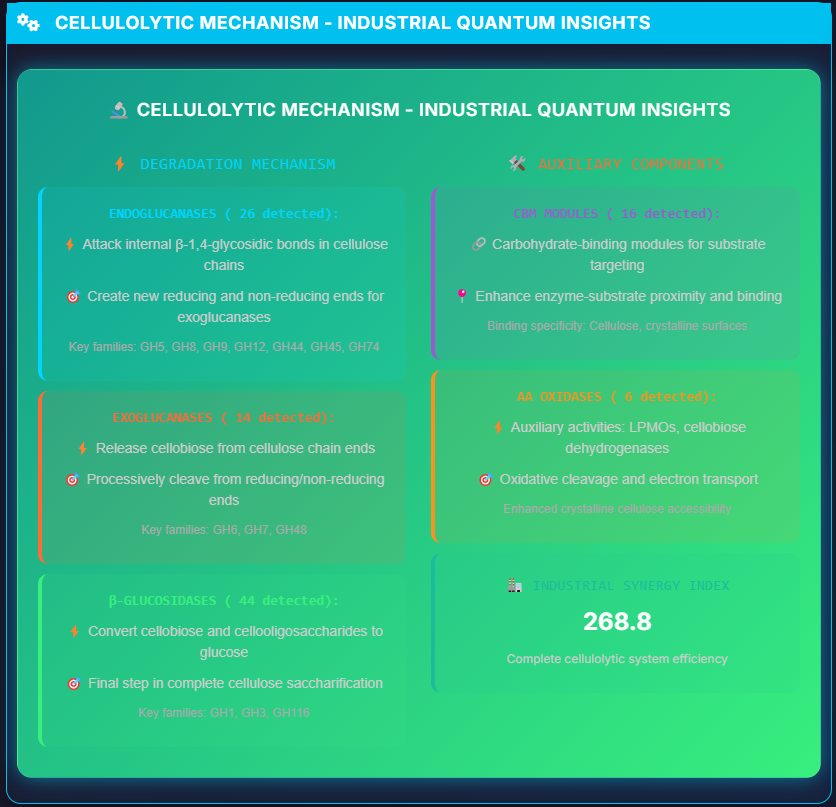

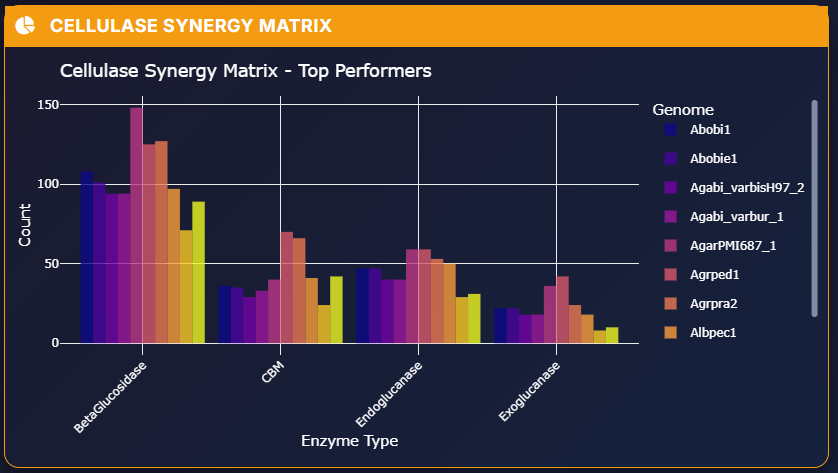

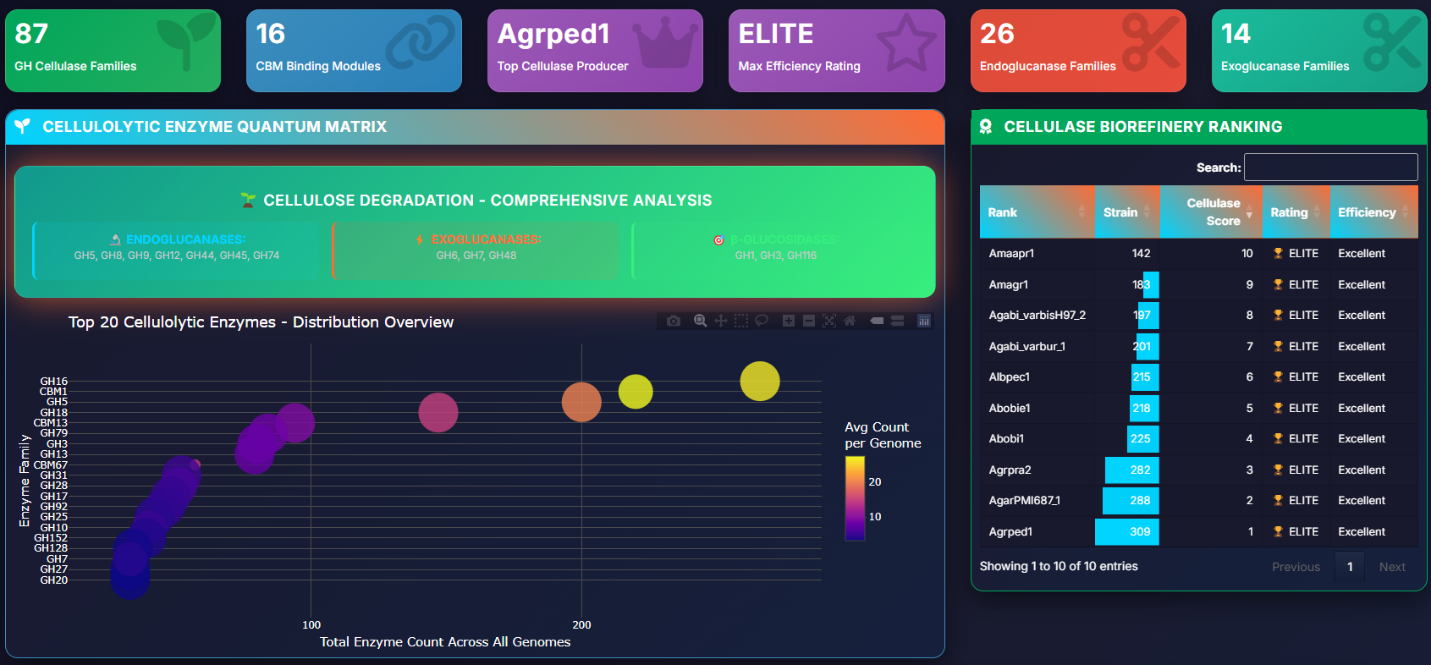

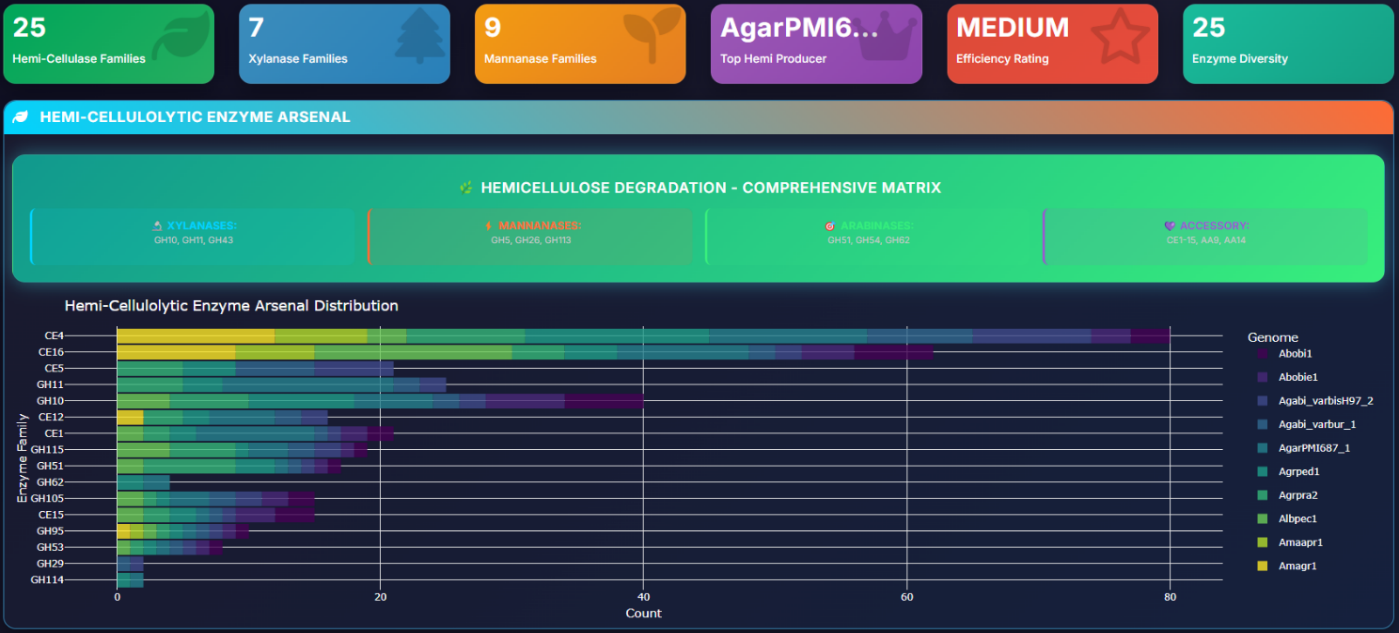

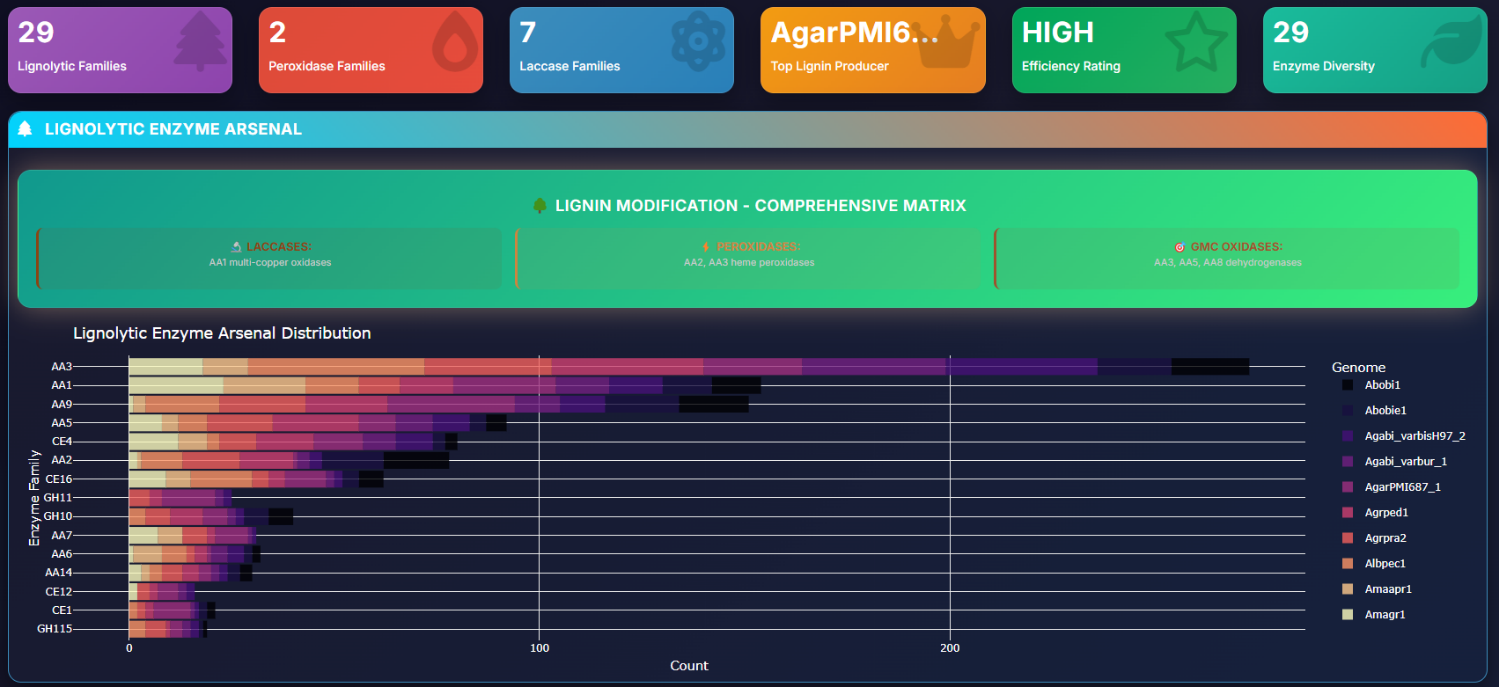

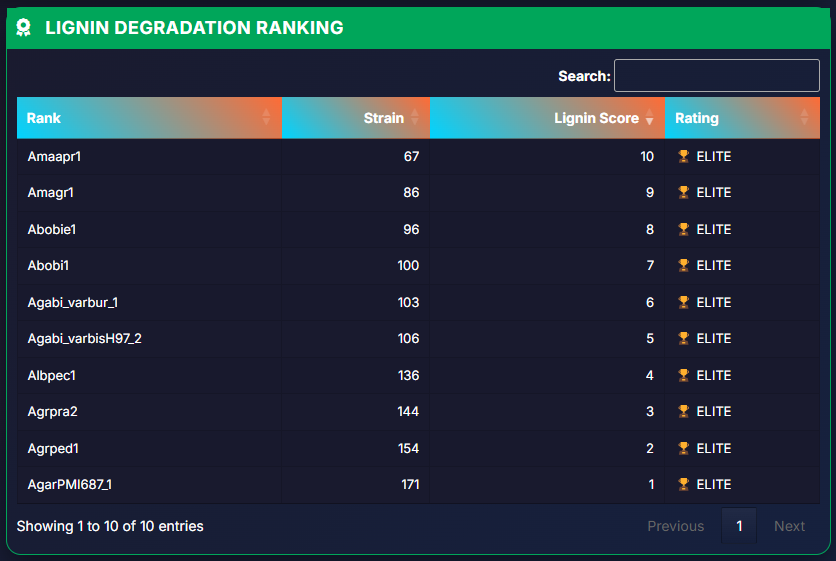

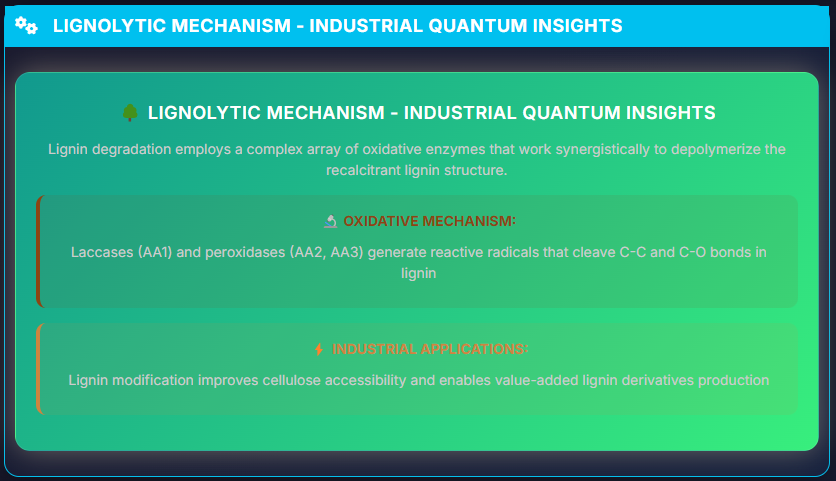

**7. Biorefinery Optimization**

**Step 7.1: Industrial Strain Selection**

- Navigate to "Biorefinery Optimizer"
- Review automated strain recommendations:
  - **Best Overall:** Complete biorefinery potential
  - **Cellulase Champion:** Superior cellulose processing
  - **Hemicellulose Expert:** Advanced hemicellulose degradation

**Step 7.2: Performance Matrix Analysis**

- Optimal strain performance visualization
- Economic viability assessment
- Industrial application recommendations

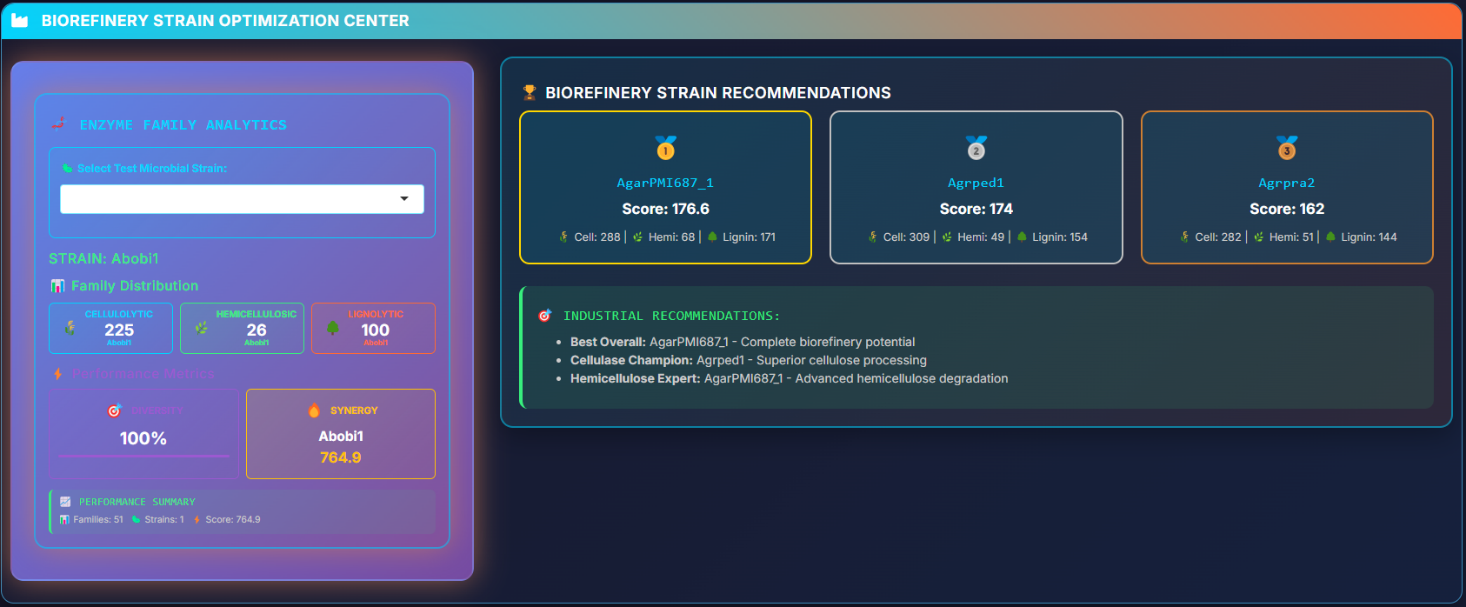

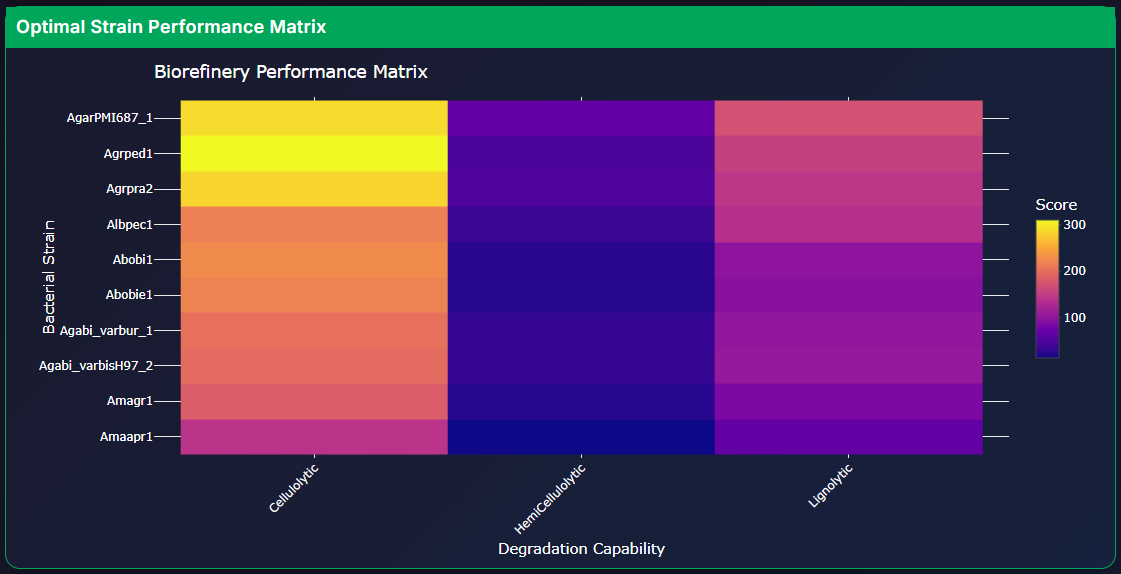

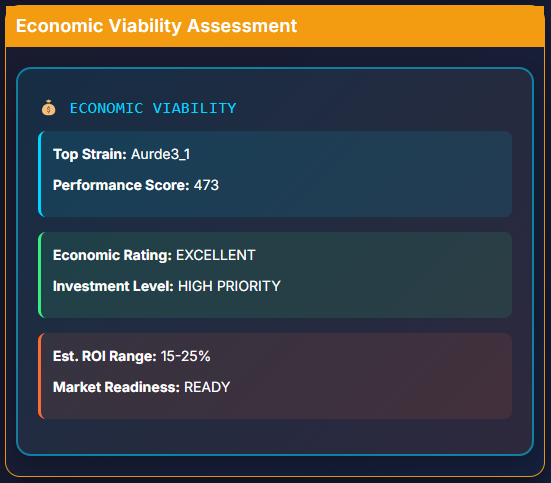

**8. Advanced Analytics**

**Step 8.1: Statistical Analysis**

- Navigate to "Advanced Analytics"
- Explore correlation matrices between enzyme families
- Principal Component Analysis (PCA) for dimensionality reduction
- Identify co-occurring enzyme patterns

**Step 8.2: Machine Learning Insights**

- Predictive modeling results
- Enzyme synergy predictions
-
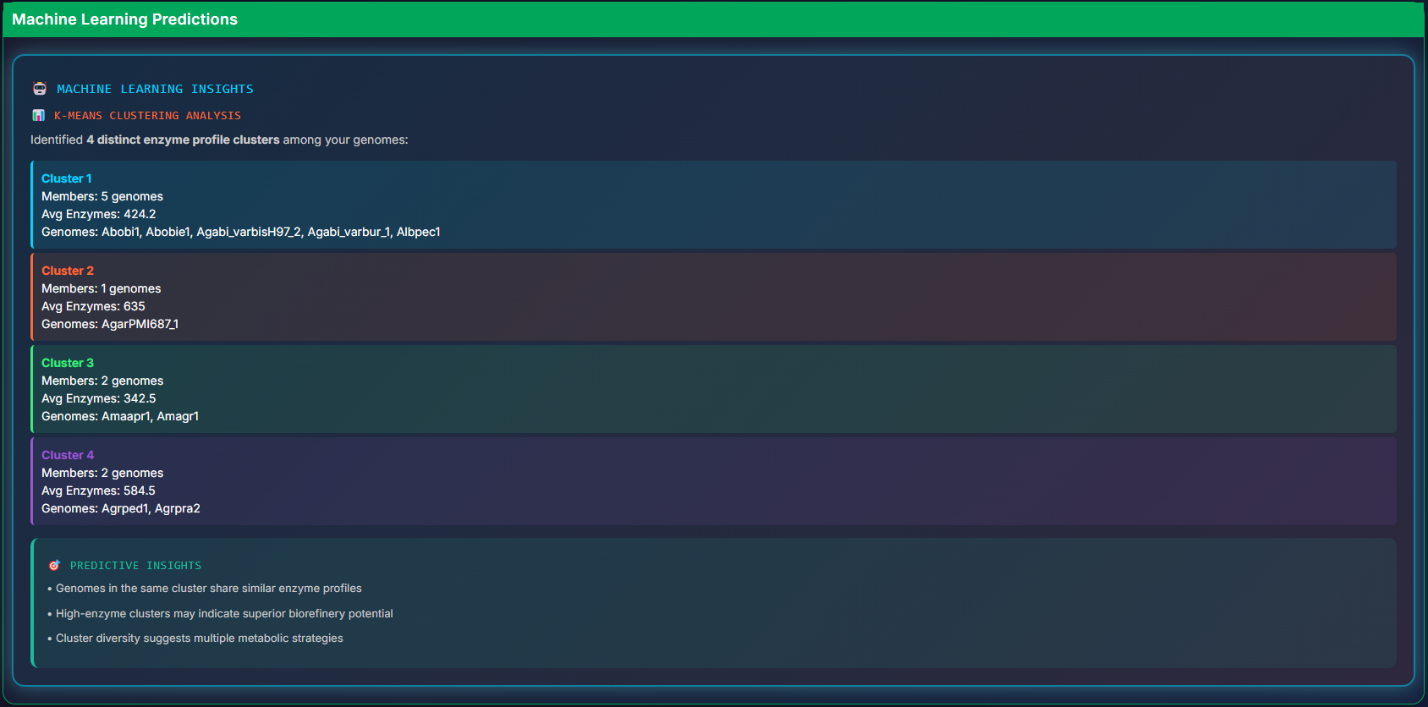
Biorefinery performance forecasting

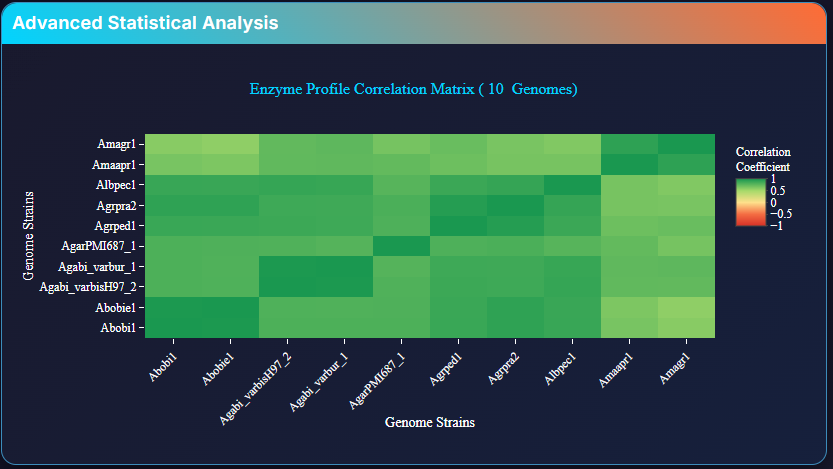

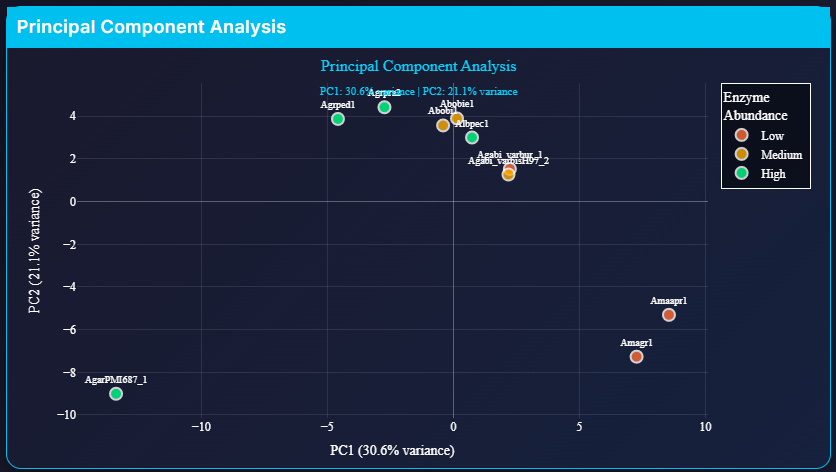

**9. Pathway Network Analysis**

**Step 9.1: Single Strain Network Visualization**

- Navigate to "Pathway Network"
- Select target strain for network analysis
- Click "GENERATE PATHWAY NETWORK"

**Step 9.2: Network Interpretation**

- Interactive network showing enzyme-pathway relationships
- Node sizes represent enzyme abundance
- Color coding indicates pathway categories:
  - Blue: Cellulolytic
  - Green: Hemi-cellulolytic
  - Red: Ligninolytic
  - Purple: Pectinolytic
  - Orange: Starch-degrading
  - Teal: Inulin-degrading

**Step 9.3: Industrial Assessment**

- Multi-functional enzyme identification
- Pathway integration analysis
- Biorefinery potential scoring

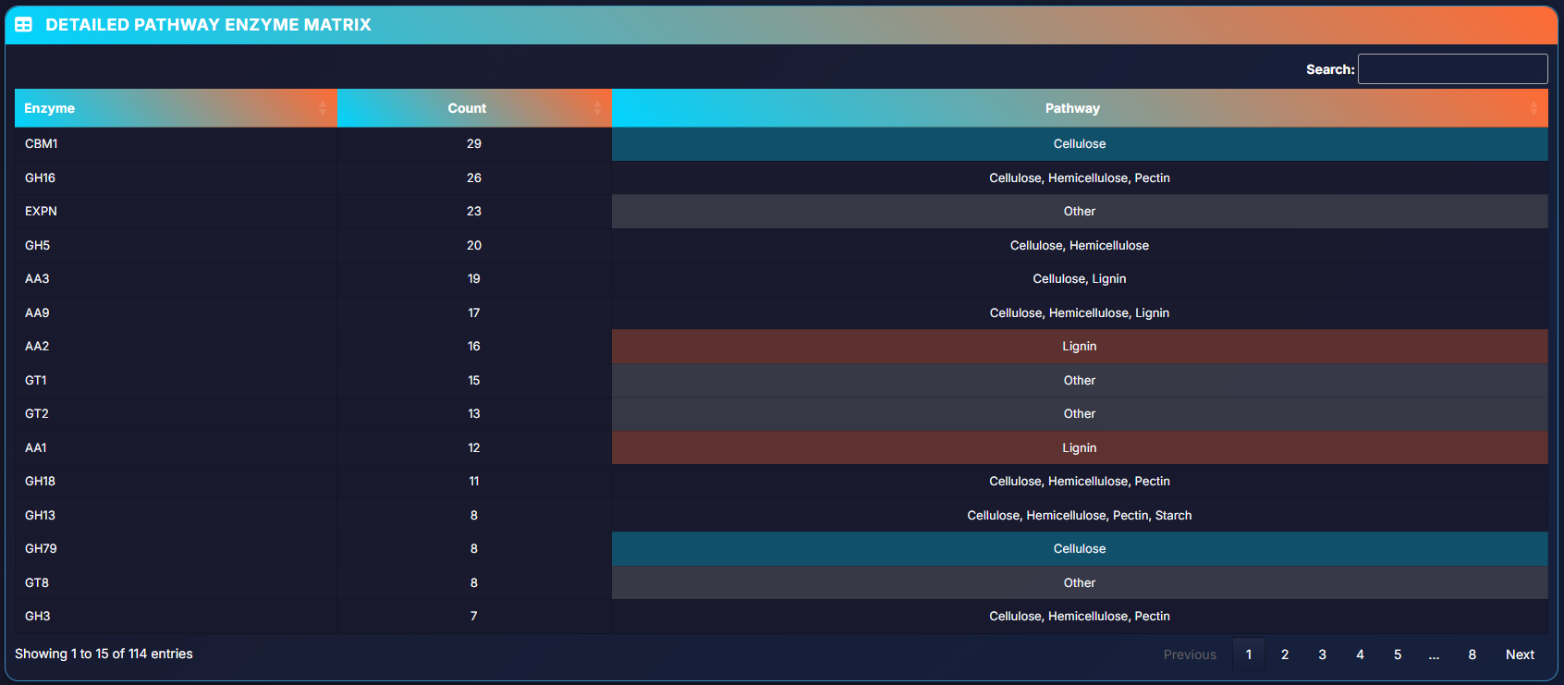

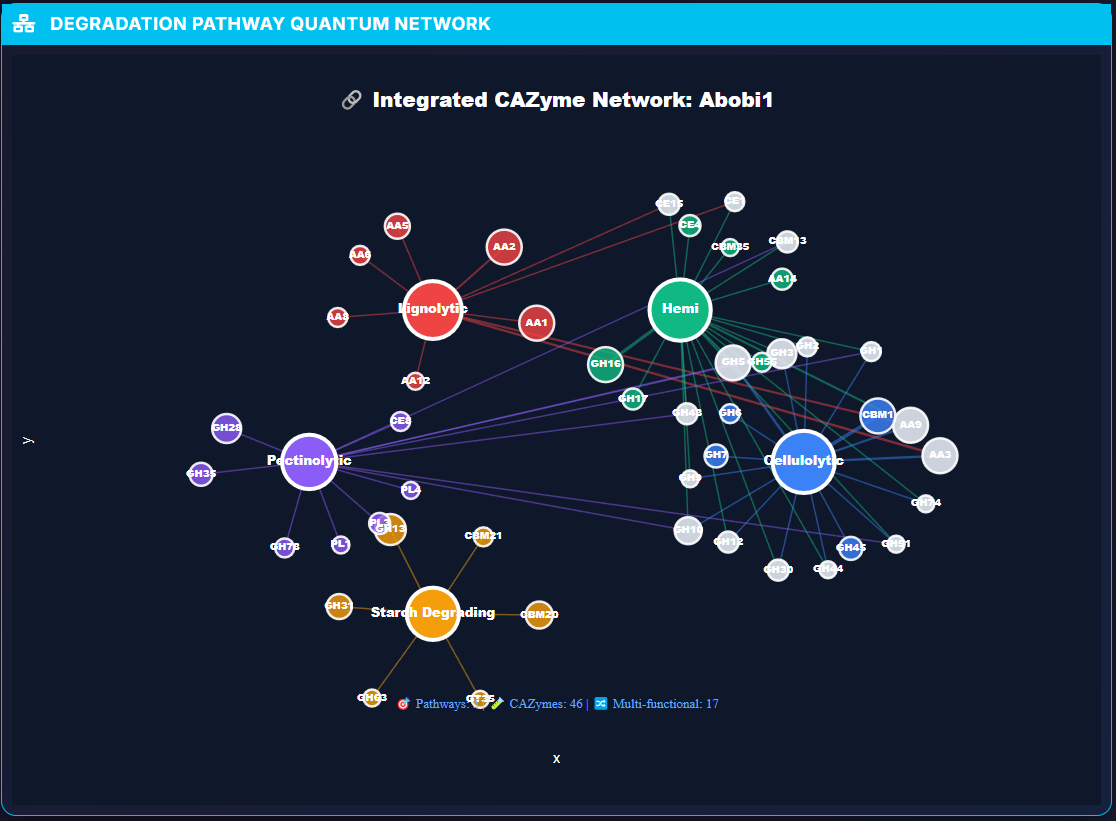

This comprehensive guide enables users to fully leverage CAZyXplorer's capabilities for identifying optimal microbial catalysts in biorefinery applications.
